# High-Resolution Genome Assembly and Population Genetic Study of the Endangered Maple *Acer pentaphyllum* (Sapindaceae): Implications for Conservation Strategies

**DOI:** 10.1101/2024.08.06.606808

**Authors:** Xiong Li, Li-Sha Jiang, Heng-Ning Deng, Qi Yu, Wen-Bin Ju, Yu Feng, Bo Xu

## Abstract

*Acer pentaphyllum* Diels (Sapindaceae), a highly threatened maple endemic to dry-hot valleys of the Yalong River in western Sichuan, China, requires urgent conservation efforts due to its extremely rarity and restricted distribution. In this study, we present a high-quality chromosome-level reference genome of *A. pentaphyllum* (626 Mb, 2n = 26), comprising 70.64% repetitive sequences and 38,540 protein-coding genes. Phylogenetic analysis shows that *A. pentaphyllum* diverged from a clade consisting of *A. yangbiense* and *A. saccharum* in the late Eocene (∼37.83 Ma). In the genome of *A. pentaphyllum*, genes involved in responding to external environmental change have undergone expansion and positive selection, potentially reflecting its adaptive strategies. While no recent independent whole-genome duplication events were detected, a recent rapid amplification of transposable elements was observed. Population genomic analysis of 227 individuals from 28 populations revealed low genetic diversity (1.04 ± 0.97 × 10^−3^) compared to other woody species. Phylogeographic patterns suggest an upstream colonization along the Yalong River, with two severe population bottlenecks correlating with major Pleistocene climatic transitions. In addition, most populations exhibit high inbreeding and genetic load, particularly those at range edges (TKX, CDG, TES). Based on these genomic insights, we propose targeted conservation strategies, including genetic rescue measures, to safeguard this unique maple species. These findings not only contribute to the preservation of *A. pentaphyllum* but also enhance our understanding of plant adaptation to extreme environments and the impacts of climate change on species with restricted distributions.

## Introduction

The genus *Acer* L., commonly known as maples, is placed in the soapberry family, along with lychee and horse chestnut (Stevens, 2016). Comprising approximately 200 species, *Acer* is predominantly distributed across the temperate regions of the Northern Hemisphere, with a particular concentration in Asia. China stands as the modern center of *Acer* diversity, hosting over 100 species (Xu *et al*., 2008). Maples are renowned for their distinctive palmate leaves and winged fruits (samaras), and hold significant ecological, economic, and cultural importance. Their applications span timber production, ethnomedicine, ecosystem services, and ornamental horticulture. Traditional medicinal uses include the consumption of young buds and leaves in teas for circulatory benefits, and the use of fruits for their purported anti-inflammatory and oral health properties (González-Sarrías *et al*., 2012, Bi *et al*., 2016).

*Acer pentaphyllum* Diels (2n=2x=26) is a unique species within the genus, being the sole member of the series *Pentaphylla*. Endemic to the dry-hot valley of the Yalong River in western Sichuan Province, China, it inhabits elevations between 2,100 and 3,100 meters. After a half-century absence from botanical records and presumed extinction, *A. pentaphyllum* was rediscovered in 1982. Recent surveys by our team have documented its presence in several Sichuan counties, including Yajiang, Kangding, Jiulong, and Muli. However, the species faces severe conservation challenges due to its narrow distribution range and habitat fragmentation caused by human activities such as road construction and grazing. *A. pentaphyllum* is classified as critically endangered (CR) in both the Chinese Red List of Biodiversity and the IUCN Red List of Threatened Species. Furthermore, the Sichuan government has designated it as a “plant species with extremely small population” (PSESP) (Pan *et al*., 2014). Our field investigations have revealed alarmingly small population sizes in some locations, emphasizing the urgent need for comprehensive research and effective conservation measures.

*A. pentaphyllum*, a deciduous tree reaching up to 10 meters in height, is acclaimed as one of the most ornamentally appealing maples. Its leaves undergo a striking autumnal color transformation from green to yellow and finally to a vivid golden-red, making it highly desirable for horticultural and landscape applications. Despite its aesthetic appeal and scientific significance for studying *Acer* evolution, particularly in the context of adaptation to dry-hot valleys, *A. pentaphyllum* remains critically endangered. While numerous studies have been conducted on *A. pentaphyllum* in recent years (Roh *et al*., 2010, Luo *et al*., 2017, Hao *et al*., 2019, Li *et al*., 2019), its evolutionary history and the genetic mechanisms underlying its endangered status remain poorly understood. This knowledge gap is primarily due to a lack of comprehensive genomic information, which has limited conservation efforts to traditional methods such as translocation.

Recent advancements in sequencing technologies have facilitated whole-genome studies across numerous *Acer* species, including *A. yangbiense* (Yang *et al*., 2019), *A. truncatum* (Liang *et al*., 2019), *A. catalpifolium* (Yu *et al*., 2021), *A. saccharum, A. negundo* (McEvoy *et al*., 2022), *A. pseudosieboldianum* (Li *et al*., 2022), *A. rubrum* (Lu *et al*., 2022) and *A. palmatum* (Chen *et al*., 2023). These genomic resources have provided unprecedented insights into the evolution and diversification of the *Acer* genus, shedding light on their adaptive strategies and phylogenetic relationships. Concurrently, conservation genomic studies combining whole-genome sequencing and population-level resequencing have significantly enhanced our understanding of the genetic mechanisms underlying population decline in many endangered plant species. Notable examples include studies on *Ostrya* (Yang *et al*., 2018), *Dipteronia* (Feng *et al*., 2024), and *Davidia* (Chen *et al*., 2020). These investigations have revealed critical information about genetic diversity, inbreeding depression, and adaptive potential, which are essential for developing effective conservation strategies.

Here, we present the first high-quality, chromosome-level genome for *A. pentaphyllum* using the PacBio Single Molecule Real Time (SMRT) and Next Generation Sequencing (NGS) technologies. To comprehensively elucidate the species’ genetic landscape, we employed an integrative approach combining comparative genomics with population genetics analyses. Our dataset encompasses 227 resequenced individuals collected across 28 remaining wild populations, representing an unprecedented sampling effort for this critically endangered maple species. Utilizing this extensive genomic data, we conducted a series of in-depth analyses to address several key questions pertaining to the evolutionary history, adaptation, and conservation of *A. pentaphyllum*: 1) How has adaptation to the dry-hot valley environment been reflected in the genome of *A. pentaphyllum*? 2) How have past climate changes influenced the colonization and demographic history of *A. pentaphyllum*? 3) What are the most effective conservation strategies for this critically endangered species? By addressing these questions, our study not only advances our understanding of *A. pentaphyllum*’s biology and evolution but also provides crucial insights for developing evidence-based conservation strategies. Our findings will contribute to the scientific basis for preserving *A. pentaphyllum* and maintaining the rich biodiversity of its unique ecosystem.

## Results

### Genome sequencing, assembly, and assessment

We obtained a total of approximately 126.96 Gb (188.45×) sequencing data for the *de novo* assembly of *A. pentaphyllum*, including 30.12 Gb (44.47×) PacBio HiFi reads, 35.98 Gb (53.13×) Illumina short reads, and 60.86 Gb (89.85×) Hi-C sequencing data (Table S1). GenomeScope results suggested low heterozygosity rate (0.156%) and high repeat contents (69.59%) in the *A. pentaphyllum* genome. Utilizing the optimal assembly method, the draft assembly of *A. pentaphyllum* was ∼662.38 Mb, which was similar to the 630 Mb genome size estimated by the flow cytometry analysis and the 677 Mb genome size calculated using 17-mers (Figures S1–S2, Table S2, Appendix S2). In total 65 contigs (626.26 Mb, accounting for 94.55% of the total assembly) were anchored to the 13 pseudo-chromosomes of *A. pentaphyllum,* with a contig N50 of 25.06 Mb and a scaffold N50 of 48.21 Mb (Figure 2a, Figure S3, Table S3). Assessment of the conservation, completeness, and consistency suggested that 99.99% of HiFi sequences could be mapped to the final assembly and 98.7% of complete benchmarking universal single-copy orthologs (BUSCO) groups (2,294, including 2,195 complete and single-copy groups and 99 complete and duplicated groups) were detected (Table S4), suggesting a high-quality reference genome for *A. pentaphyllum*.

**Figure 1.**
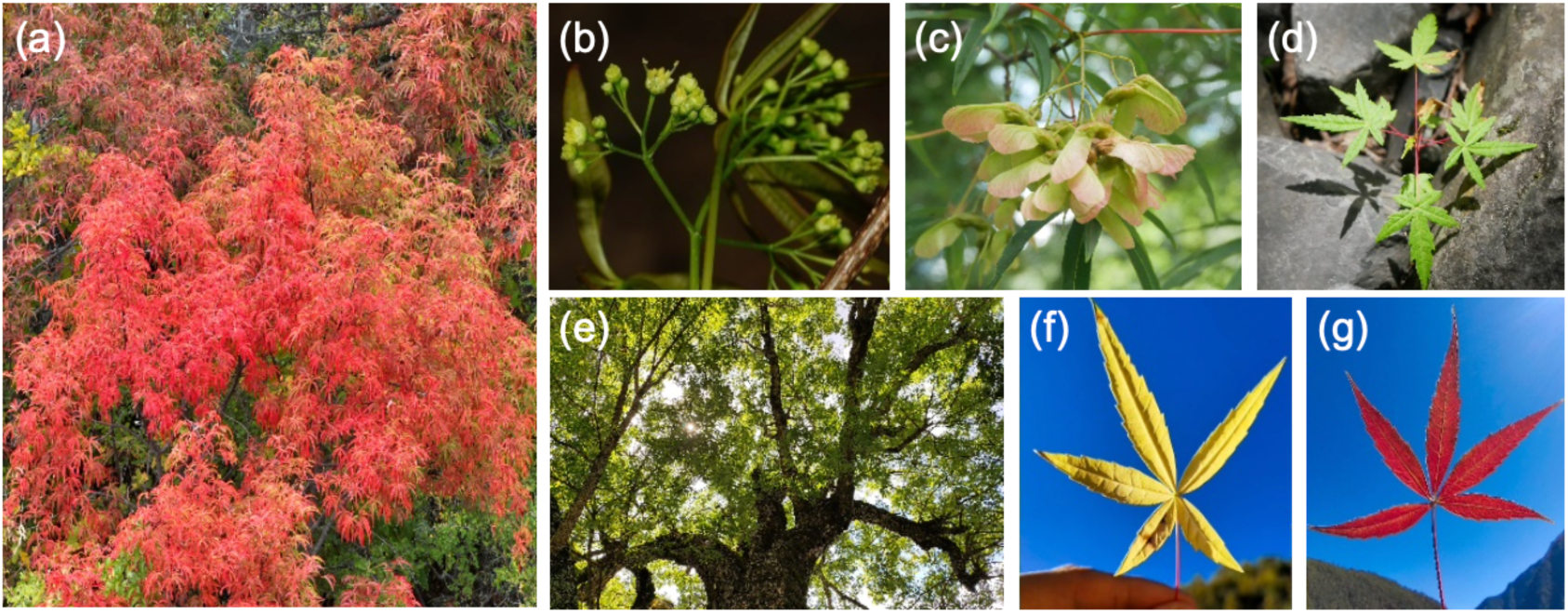
Photographs of *A. pentaphyllum*. (a) Adult tree, (b) flower, (c) samara, (d) seedling, (e) canopy, and leaves in different colors during autumn (f) (g).

**Figure 2.**
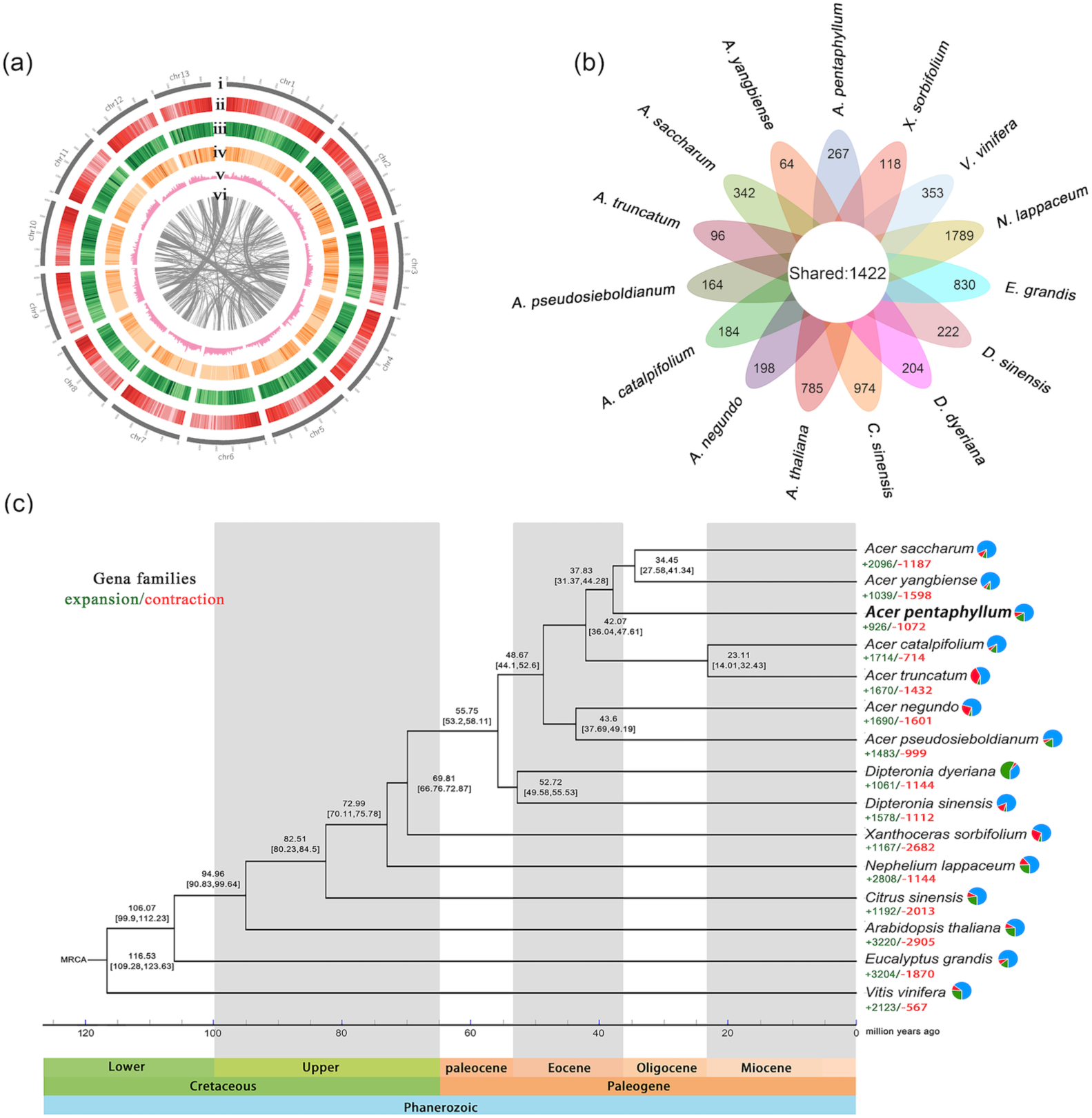
Genomic characteristics and genome evolution of *A. pentaphyllum*. (a) Circos plot of the genomic features. The circles, from outermost to innermost, show (i) thirteen pseudochromosomes, (ⅱ) gene density, (ⅲ) repeat element density, (ⅳ) single nucleotide polymorphism (SNP) density for population genomic analysis, (ⅴ) GC content, and (ⅵ) syntenic blocks. (b) Petal map of unique gene families and shared single-copy genes in 15 woody species. (c) A maximum likelihood tree, including *A. pentaphyllum* and other 6 *Acer* species, was constructed based on 1422 single-copy orthologous genes. Gene families that expanded (green) and contracted (red) are displayed as numbers below each species of the tree, and values at the nodes represent the divergence times and 95% CI (confidence interval).

### Genome annotation

We identified and annotated a total of 1,283,444 repetitive sequences, comprising 70.64% (∼467.93 Mb) of the whole genome of *A. pentaphyllum*. Among these, the most abundant elements were long terminal repeat retrotransposons (LTR-RTs), making up 45.93% (∼304.22 Mb) of the genomic content, and *Copia* and *Gypsy* superfamilies were the primary components, accounting for 20.80% (∼137.77 Mb) and 13.89% (∼92 Mb) of the assembled genome, respectively (Table S5). The distribution of LTR insertion times showed that most LTR-RTs have gradually accumulated in the *A. pentaphyllum* genome over the past 10 million years, with a peak of insertion activity occurring around 1.3 million years ago (Mya, Figure S4). Additionally, DNA transposons were also plentiful, spanning 80.05 Mb (12.09%) on the genome (Table S5). A total of 38,540 protein-coding genes were successfully annotated in the genome of *A. pentaphyllum* based on structural predictions and transcriptome data, spanning 146.18 Mb and accounting for 23.34% of the total genome. The mean lengths of gene region, transcript, and protein-coding region (CDS) were 3,792 bp, 1,689 bp, and 1,425 bp, respectively. Each transcript contains 5 exons on average, with an average length of 355 base pairs (Table S6). Among the 38,540 genes predicted in the *A. pentaphyllum* genome, functions of 36,336 genes (accounted for approximately 94.28%) were annotated using Non-Reduntant Protein Database (NR, 36,329, 94.26%), Swissport (24,865, 64.52%), Kyoto Encyclopedia of Genes and Genomes (KEGG, 12,411, 32.20%), Gene Ontology (GO, 13,798, 35.80%), and evolutionary genealogy of genes: Non-supervised Orthologous Groups (eggNOG, 32,251, 83.68%) databases (Figure S5, Table S7). Moreover, we identified 10,972 non-coding RNA (ncRNA), including 4,396 rRNA (∼2372.25 kb), 5492 tRNA (∼41.92 kb), 128 miRNA (∼16.61 kb), 151 snRNA (∼22.22 kb), and 805 snoRNA (83.99 kb), counting for 0.44% (∼2914.25 kb) of the *A. pentaphyllum* genome (Table S8).

### Genomic **p**hylogenetic evolution and **s**ignatures of **p**ositive **s**election

Overall, we identified 460,262 homologous genes belonging to 30,421 gene families from 15 woody plant genomes, among which 1,422 were single-copy orthologous genes shared by all species, and 267 gene families containing 1,572 genes were species-specific to *A. pentaphyllum* (Figure 2b, Figure S6, Table S9). *A. pentaphyllum* shared 11,661 orthogroups with other 6 *Acer* species while shearing 15,014 orthogroups with *D. sinensis* and *D. dyeriana*. Results from the dated phylogenetic tree showed that *A. pentaphyllum* is sister to a clade consists of *A. saccharum* and *A. yangbiense*, with the divergence time estimated at approximately 37.83 Mya (95% HPD: 44.28–31.37 Mya) during the Eocene-Oligocene transition (EOT), slightly later than the divergence of *A. negundo* and *A. pseudosieboldianum* (∼43.6 Mya, 95% HPD: 49.19–37.69 Mya). Furthermore, the *Acer* and the *Dipteronia* species were predicted to have diverged from each other in the late Paleocene–early Eocene boundary (∼55.75 Mya, 95% HPD: 53.2–58.11, Figure 2c).

A total of 926 expanded and 1,072 contracted gene families in *A. pentaphyllum* were identified and annotated (Figure 2c). GO enrichment analysis of these expanded genes revealed significant enrichment (Q <0.05) in macromolecular metabolism (GO:0019318, GO:0044238), cytochrome synthesis (GO:0016168), photosynthesis (GO:0009767, GO:0009772), ATP synthesis (GO:0015986), heme transport (GO:0015886, GO0015232), and ion transport (GO:1902600) etc (Figure S7a, Table S10). Further KEGG analysis identified 13 significantly enriched pathways (Q <0.05), primarily associated with photosynthesis (ko00195), plant hormone signal transduction (ko04075), glyoxylic acid and dicarboxylic acid metabolism (ko00630), plant and pathogen interaction (ko04626), monoterpenoid biosynthesis (ko00902), and other metabolic pathways (Figure S7b, Table S11). In addition, we also identified 108 putative positively selected genes (PSGs) (Q < 0.05) in the *A. pentaphyllum* genome through the calculation of the ratio of non-synonymous substitutions to synonymous substitutions (*Ka/Ks*). These positively selected genes are significantly enriched (Q <0.15) in biological processes related to abiotic stress, response to plant hormones, and catalytic activity (Figure S7, Table S12).

### Analysis of collinearity and **w**hole-genome **d**uplication

Intragenomic collinearity analysis detected a total of 3,381 colinear gene pairs on 290 high-quality colinear blocks (gene pairs ≥4) within the *A. pentaphyllum* genome using JCVI, accounting for 8.77% of the total genes. When compared to the genomes of the two *Acer* species and one *Dipteronia* species, *A. pentaphyllum* contained fewer proportion of intragenomic collinear gene pairs (*A. yangbiense*: 9.65% vs, *A. pseudosieboldianum*: 11.93%, and *D. sinensis*: 10.46%). Further interspecific collinearity analysis between *Acer* and *Dipteronia* revealed that *A. pentaphyllum* shares 33,342 homologous genes with *A. yangbiense*, slightly higher than the number with *D. sinensis* (32,687). The distribution and arrangement of longer collinear blocks among the three *Acer* species are relatively consistent across 13 chromosomes, revealing a nearly 1:1 collinearity depth ratio (Figure 3a, Figure S9). This indicated that there was no significant structural or copy number variation in the chromosomes of *A. pentaphyllum* since diverging with other *Acer* species. In contrast, the arrangement of collinear blocks between *A. pentaphyllum* and *D. sinensis* is more disordered (Figure 3a, Figure S9), which aligns with their more distant positions in the phylogenetic tree (Figure 2c). Additionally, the *Ks* distribution map for 4 species (*A. pentaphyllum, A. yangbiense*, *D. dyeriana*, and *V. vinifera*) showed only one distinct peak (Figure 3b, Figure S10), indicating similar evolution histories of these species without recent whole-genome duplications (WGDs) other than the typical core eudicot common hexaploidization (Li and Barker, 2020).

**Figure 3.**
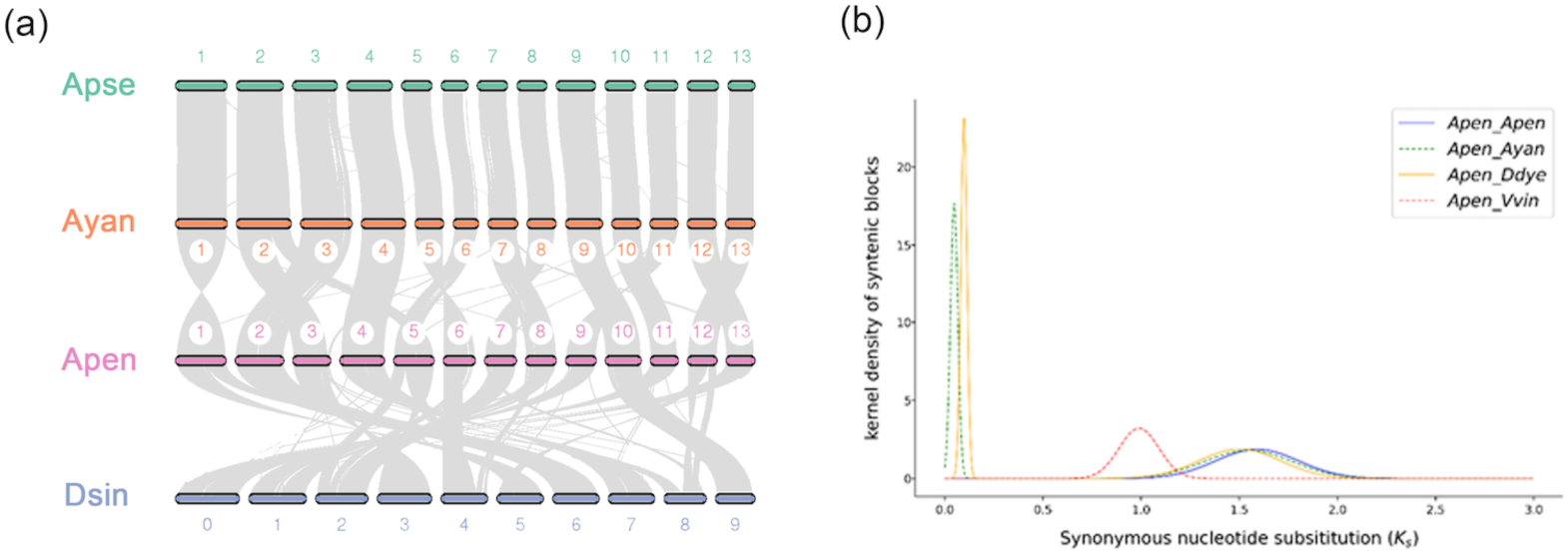
(a) Syntenic blocks among *Acer* and *Dipteronia* species. (b) Frequency distribution of *K*_s_ for paralogous and orthologous genes of *A. pentaphyllum* (Apen), *A. yangbiense* (Ayan), and *V. vinifera* (Vvin).

### Field studies, population structure and **p**hylogenetic **r**elationships

Through field investigations, we found a total of ∼1.65×10^4^ *A. pentaphyllum* individuals within the entire distribution areas, including ∼7,200 adult trees (Table S22). The TKX, MDG, TZC, ZQG, CDG, and MLC populations show extremely small numbers of individuals (<100) and mature trees (<50), whereas the AJ and ZD populations in Muli County are relatively large, with over 2,000 individuals and more than 1,000 mature trees (Table S22). A total of ∼3.01 Tb resequencing reads were generated for 227 *A. pentaphyllum* accessions from 28 wild populations, with an average sequencing depth of approximately 26 × and an average heterozygosity of 0.16% (Table S13). After SNP calling and filtering, 2,757,997 high-quality SNPs were retained (Dataset 3). An annotation of SNPs showed that ∼61.57% of SNPs were located in the intergenic regions while the remaining ∼38.43% were found in genic regions, which encompass exonic (7.27%), splicing (0.04%), stop-gain (0.23%), nonsynonymous (4.42%), UTR5/3 (0.84%), intronic (10.83%), and upstream/downstream (21.02%) regions (Table S14).

The admixture analysis based on unlinked and intergenic sites (Dataset 4) revealed clear subspecies genetic structure among populations of *A. pentaphyllum*, revealing seven clusters (*K* = 7) at the lowest cross-validation (CV) error (Figure 4a, Table S15). Individuals from cluster 1 (CDG), cluster 2 (TKX), cluster 5 (LM), and cluster 7 showed relatively purer ancestry compared to other *A. pentaphyllum* clusters, especially those from cluster 6. This group is distributed at the center of the overall distribution across the river and showed higher levels of mixed genetic ancestry (Figure 4a, Figure 4d). Additional subclusters were detected at higher *K* values (Figure S11). Interestingly, individuals from TES clustered with those from Yajiang and Kangding counties when *K*<8, but as *K* increases, TES shared a similar genetic ancestry to AJ (Figure S10), despite the considerable geographic distance between these two populations (∼74 km). PCA analysis revealed clear separations of the CDG and TKX populations from all others based on PC1 and PC2, which accounted for 15.26% and 11.55% of the total variance, respectively (Figure 4b).

**Figure 4.**
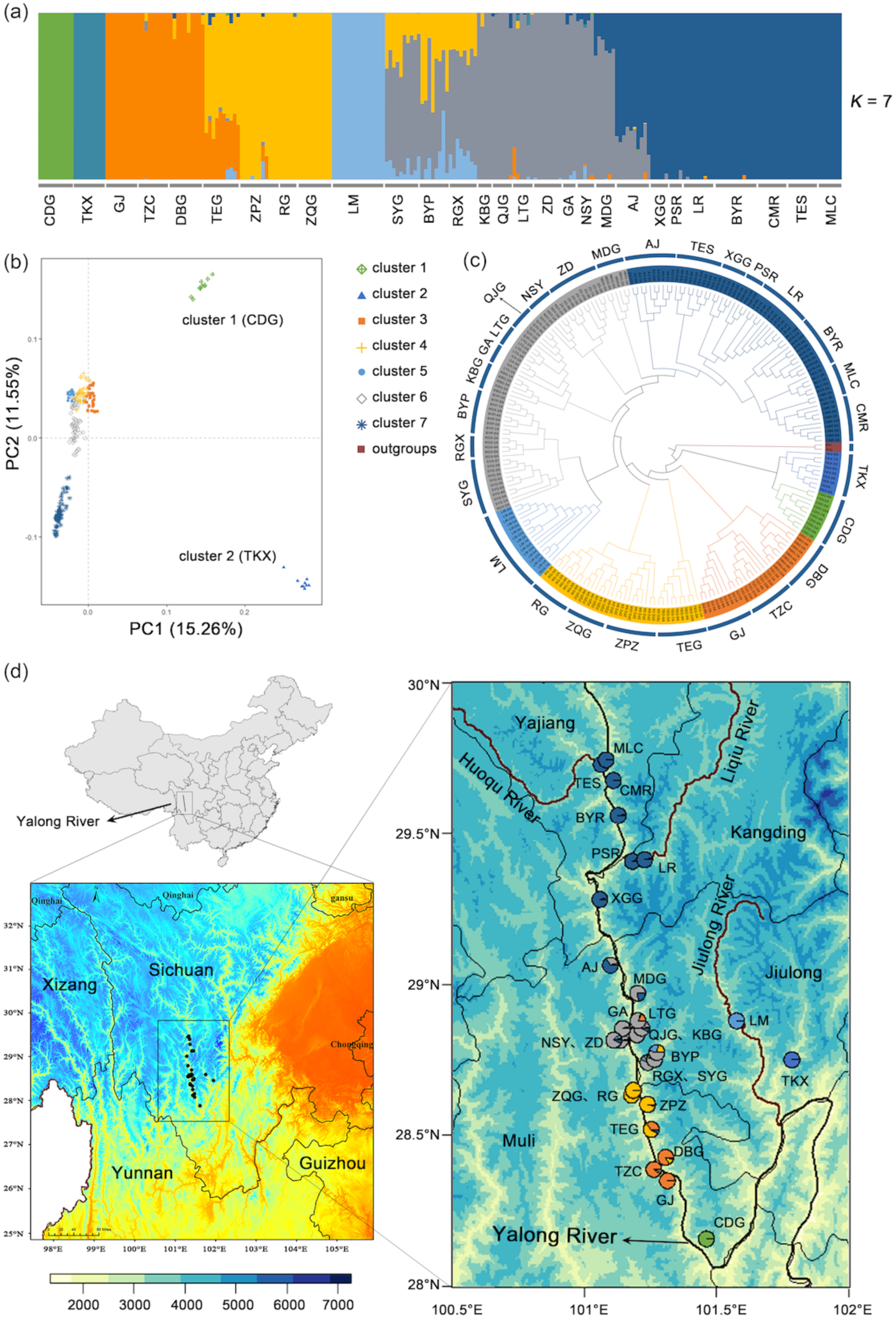
Genetic structure and phylogenetic relationships of the 28 populations of *A. pentaphyllum*. (a) Population genetic structure at *K* = 7. (b) PCA plot of *A. pentaphyllum* based on neutral loci. (c) The phylogenetic tree of 227 *A. pentaphyllum* individuals was constructed based on SNP Dataset 4. (d) Geographic distribution maps of ADMIXTURE clusters for the 28 populations of *A. pentaphyllum* at *K* = 7. The colors of the populations in (b), (c), and (d) correspond to the genetic clusters in (a).

Notably, the phylogenetic positions of the 28 populations on the ML tree were geographically located almost completely opposite to the flow direction of the Yalong River (Figure 4c, Figure 4d, Figure S12). Specifically, individuals from CDG and TKX, situated in the lower reaches of *A. pentaphyllum*’s entire distribution range, diverged first at the base of the tree. This was followed by the divergence of individuals from Muli and Jiulong counties, while individuals from Kangding and Yajiang counties diverged most recently (Figure 4c, Figure 4d). However, the TES population from Yajiang County, located geographically closest to MLC at the northern edge of the overall distribution, diverged earlier compared to the other populations from Yajiang and Kangding counties, except the AJ population which is the basal group of cluster 7 (Figure 4c, Figure 4d, Figure S12). Additionally, several populations located on opposite sides of the river, such as DBG-TZC-GJ, ZPZ-RG-ZQG, and GA-LTG-QJG, were also clustered together on the tree (Figure 4d), consistent with the low *F*_ST_ (Figure S15, Table S17) statistics observed between nearby populations of *A. pentaphyllum*.

### Genome-wide diversity and **i**nbreeding

Genome-wide linkage disequilibrium (LD) decay analysis revealed remarkable variation among the *A. pentaphyllum* populations The decay of LD reached half of its maximum average r^2^ at distances ranging from ∼16.78 kb in the AJ population to ∼191.84 kb in the GJ population (Figure S13). This substantial variation in LD decay rates reflects the diverse demographic histories of these populations. The genetic diversity of this narrow endemic species was found to be relatively low (1.212 ± 1.058 × 10^−3^) compared to other threatened woody plant species with available genetic diversity information (Figure S14, Table S16). It was lower than *A. yangbiense* (3.13 × 10^−3^) and *Rhododendron griersonianum* (1.94 × 10^−3^), but slightly higher than *Cercidiphyllum japonicum* (1.1 × 10^−3^), *Manglietiastrum sinicumand* (1.16 × 10^−3^), and *Populus ilicifolia* (8.3 × 10^−4^) (Figure S14, Table S16). Among the populations within *A. pentaphyllum,* the RG population had the highest genetic diversity (1.083 ± 1.071 × 10^−3^) while the TKX population had the lowest genetic diversity (5.74 ± 7.65 × 10^−4^) (Table 1). The overall *F*_ST_ analysis results indicated a correlation between geographical distance and genetic distance among population pairs of *A. pentaphyllum* (Pearson’s r = 0.4495, Figure S15). Neighboring populations, even those located on opposite sides of the Yalong River, often showed lower *F*_ST_ (Figure S16, Table S17, Table S18), suggesting that gene flow tended to be more frequent between these populations. Moreover, consistent with the results of population genetic structure and PCA, TKX (mean pairwise *F*_ST_ = 0.48) and CDG (mean pairwise *F*_ST_ = 0.28) populations showed a significant differentiation from the other populations of *A. pentaphyllum* (Figure S16; Table S17).

**Table 1.**
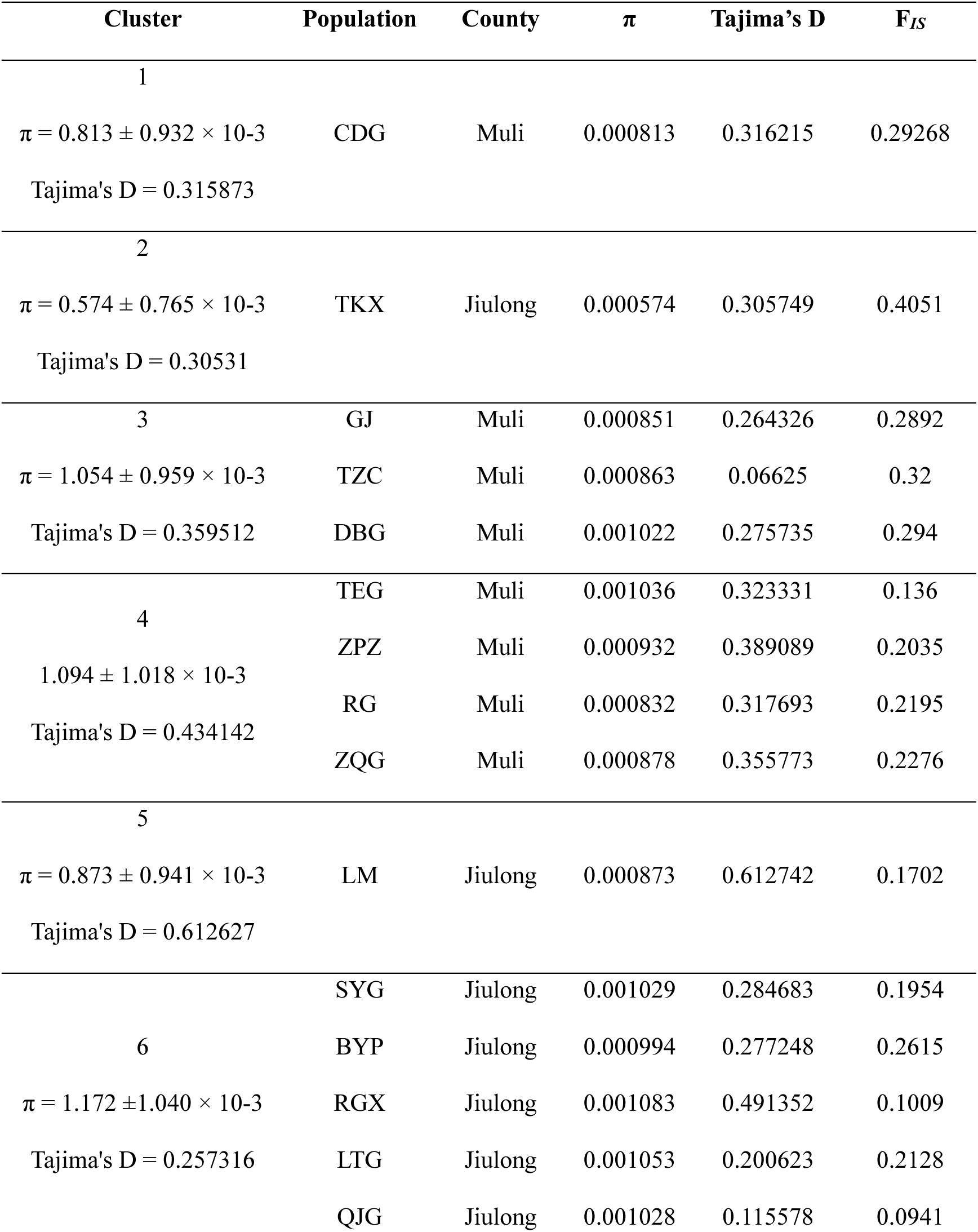
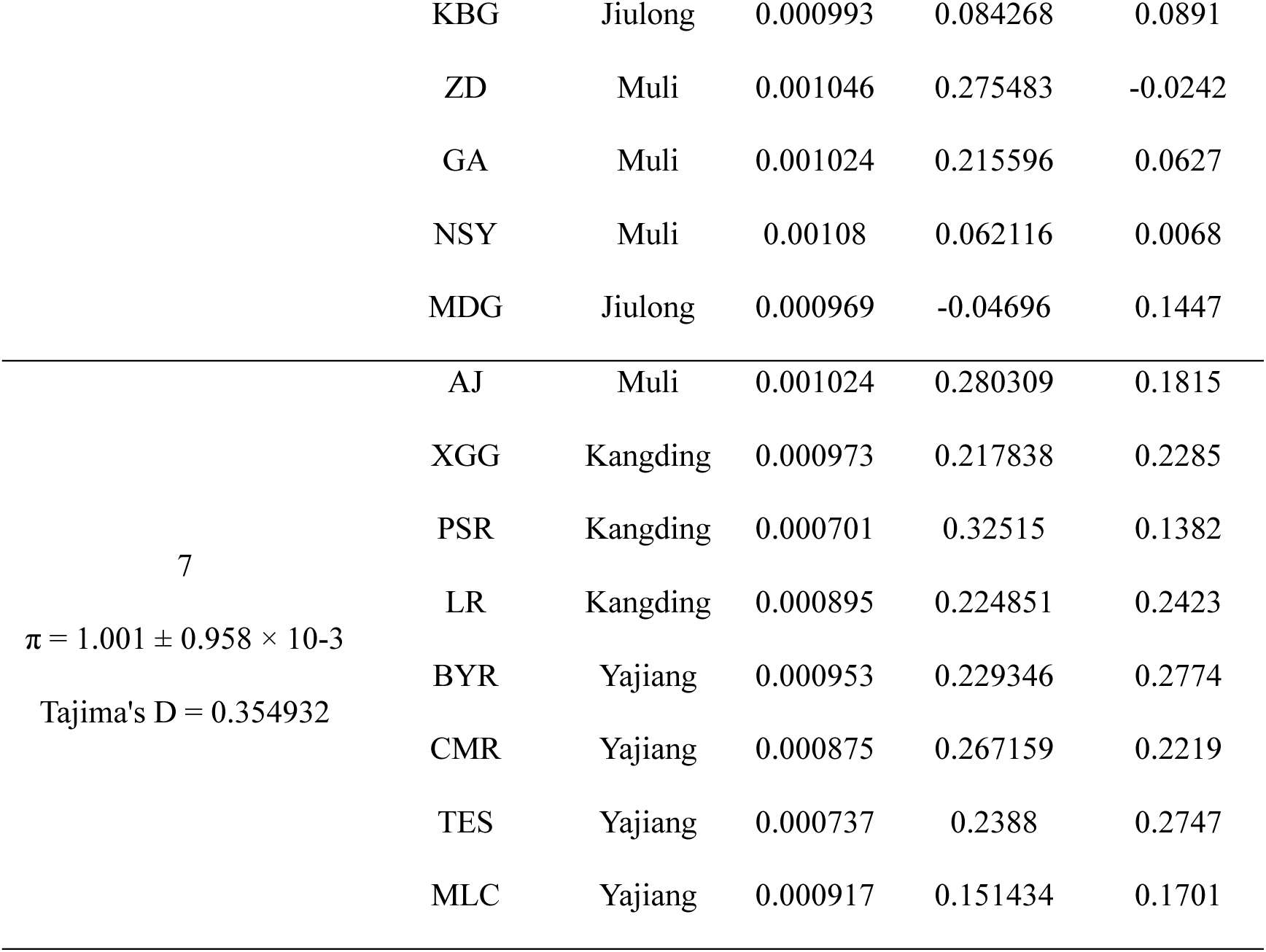
The genetic diversity (π), Tajima’s D, and inbreeding coefficients (F_IS_) for each cluster and population of *A. pentaphyllum*.

The estimation of kinship coefficient among individuals identified a total of 67 pairs of individuals as less than 3rd-degree relatives from the 7 clusters of *A. pentaphyllum* (Table S19). This included 11 pairs categorized as duplicate or monozygotic twin (Dup/MZ Twin), 22 pairs as 2nd-degree, 33 pairs as parent-offspring (PO), and 1 pair of full siblings (FS). Notably, the small-size populations, including the TKX, CDG, TZX, LM, and TES, had a higher proportion (64.18%) of closely related individual pairs, most of which were estimated to be 2nd-degree or PO relatives (Table S15). Similar, higher inbreeding coefficients (*F*_IS_) were observed in the TKX (0.4051), TZC (0.32), and TES (0.29), while most populations of lineage 6 exhibited lower *F*_IS_ (0.096, Table 1).

PLINK identified a total of 192,587 runs of homozygosity (ROHs) across all 227 individuals of *A. pentaphyllum,* averaging approximately 848 ROHs per individual. The mean ROH (*L*_ROH_) length was 168.39 kb, with the longest segment measuring 1.583 Mb (7,240 SNPs) found on XGG-1 (Table S20). The fraction of ROH in the genome (*F*_ROH_) was significantly higher in the BYR (0.35 ± 0.15), TZC (0.32 ± 0.05), GJ (0.30 ± 0.06), and TKX (0.29 ± 0.07) populations, each characterized by a high number of short segments (*L*_ROH_ < 200 kb). These short segments accounted for approximately 76.08% of all detected ROHs and contributed the highest proportion (60.52%) of the cumulative ROH length. This exceeded the contributions of large (*L*_ROH_ > 400 kb, 6.48%) and medium segments (200 kb < *L*_ROH_ ≤ 400 kb, 33%), indicating high levels of inbreeding (Table S21). However, consistent with the *F*_IS_ estimation, the *F*_ROH_ of individuals from cluster 6 was very low (mean *F*_ROH_ < 0.15), indicating reduced inbreeding within this cluster, which is located in the central region of the *A. pentaphyllum* distribution area (Table 1, Table S20, Fig. 6a).

**Figure 5.**
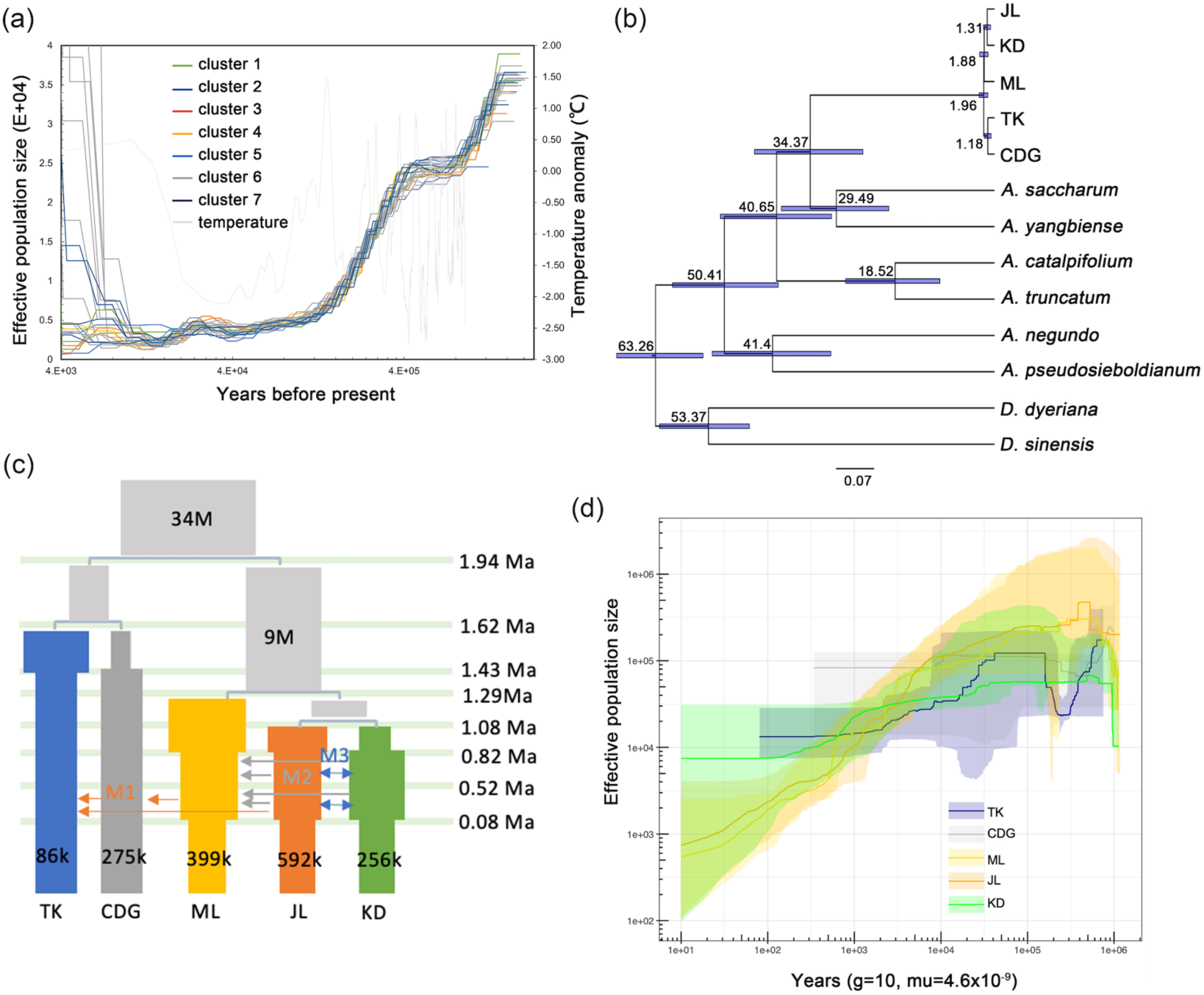
Demographic history of *A. pentaphyllum* inferred by different methods. (a) Historical changes in effective population size (*N*_e_) through time plots for *A. pentaphyllum* based on PSMC modeling, with a generation time of 10 years and a mutation rate of 4.7e-9. (b) Dated phylogeny of clusters within *A. pentaphyllum* and six *Acer* species, with two *Dipteronia* species as outgroups. (c) Illustration of the best-fit model with point estimations of the parameters. (d) Estimation of changes in effective population size changes over time for each cluster using Stairway Plot2. Colored shadings represent 95% confidence intervals calculated from 200 bootstrap replicates.

**Figure 6.**
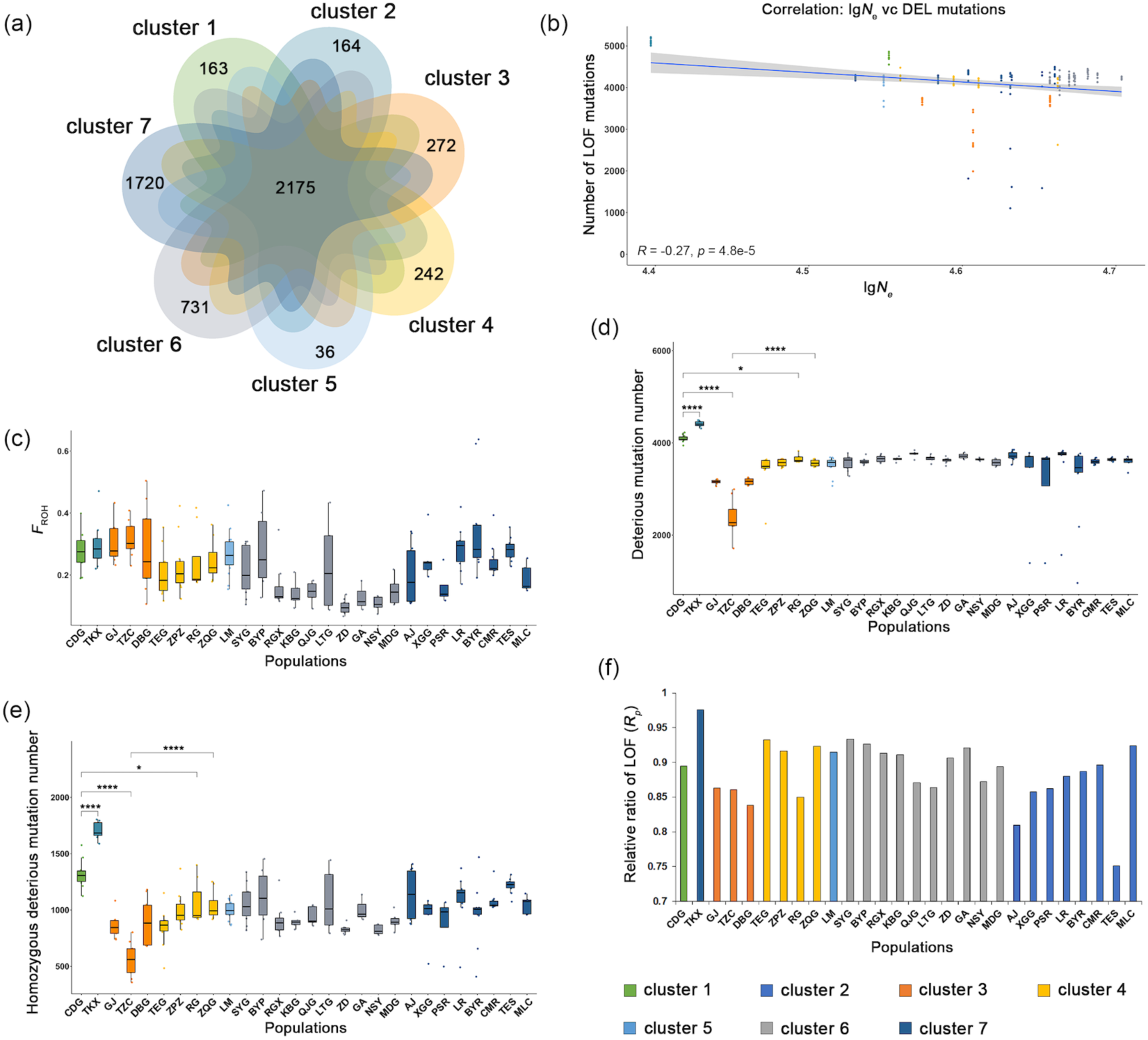
Assessment of inbreeding and genetic load in 28 populations of *A. pentaphyllum*. (a) Venn diagram of private or shared deleterious mutations among clusters of *A. pentaphyllum* at *K* = 7. (b) Correlation between the number of extreme deleterious mutations (LOF) and effective population size (*N*_e_) calculated by the formula *N*_e_ = θ/(4μ) across the 28 populations of *A. pentaphyllum*. (c) Boxplot showing *F*_ROH_ of individuals grouped by populations. (d) Number of deleterious mutations, including DEL, RADICAL, and LOF sites. (e) Number of homozygous deleterious mutations, which are calculated as 2× homozygous deleterious genotypes. (f) The relative ratio (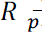) of the mean derived alleles frequency (*p*) for LOF mutations.

### Demographic history **i**nference

The demographic history inferred by PSMC suggested that *A. pentaphyllum* experienced two severe population declines within the past million years. The earliest population decline began around 1.1 Ma and lasted for approximately 0.3 million years, while the more recent decline started around 0.36 Ma and continued until approximately 0.16 Ma (Figure 5a). Following the most recent bottleneck, *A. pentaphyllum* has maintained a stable effective population size until recently. The *N*_e_ of *A. pentaphyllum* was estimated to be 5.78×10^4^ using the formula *N*_e_ = θ/(4μ) (Table S22).

To explore the comprehensive demographic histories including lineage divergence, gene flow, and population size changes through time, we employed coalescent population simulation and composite likelihood method in fastsimcoal2 (Excoffier *et al*., 2013). The populations were grouped into five major clusters according to their geographic distribution and results of phylogenetic analysis (CDG: cluster 1; TKX: cluster 2; ML: clusters 3-5; JL: cluster 6, KD, cluster 7). The priority distribution of divergence times were estimated from the 1,422 single copy orthologous genes in the comparative genomic analysis. Results showed that the crown age of *A. pentaphyllum* was dated to the early/middle Pleistocene, ca. 1.96 (95% highest posterior density, HPD: 2.82–1.18) Mya, with the clusters diverged from ca. 1.88–1.18 Mya (Figure 5b). Comparison of likelihood values among eight demographic scenarios (Figure S17) revealed that Model 8 exhibited the highest likelihood (Figures 5c, Figure S18, Table S23). According to this model, the most recent common ancestor (MRCA) of CDG and TKX diverged from the MRCA of the three other clusters approximately 1.94 Mya. Subsequently, CDG and TKX separated around 1.62 Mya, and thereafter experienced population contraction and expansion around 1.43 Mya, respectively. The remaining three clusters diverged around 1.29 Mya and 1.08 Mya. Following divergence, ML and JL underwent two distinct population contractions at approximately 0.82 Mya and 0.08 Mya, respectively. Meanwhile, the effective population size of KD initially expanded around 0.82 Mya before contracting around 0.08 Mya (Figure 5c). Gene flow was detected between JL and KD, and from KD to JL and ML, spanning from 0.82 Mya to 0.08 Mya. Additionally, gene flow from ML to TKX and CDG, as well as from JL to ML, and from CDG to TKX was observed between 0.52 Mya and 0.08 Mya (Figure 5c, Figure S18). In addition, population size changes through time inferred based on SFS by StairwayPlot2 revealed similar patterns for these clusters (Figure 5d).

### Evaluation of mutation load

After filtering out SNPs with reference alleles inconsistent with ancestral states in the *A. pentaphyllum* genome, 67,769 sites from protein-coding regions were retained for the evaluation of deleterious mutations. A total of 23,290 mutations were identified and classified into three groups, of which 19,699 (84.58%) were “deleterious” (DEL) annotated by SIFT4G, 5,600 (24.04%) were detected as “radical missense substitution” (RADICAL) with Grantham score >150, and 1499 (6.44%) were loss-of-function (LOF) mutations identified through SNPEFF (Table S25). The seven clusters shared 2,175 deleterious mutant sites (9.34% of the total), with at least one individual from each cluster carrying deleterious derived alleles at these sites. Cluster 5 (LM, 16 individuals) exhibited the fewest unique deleterious mutations (36, 0.15%), while cluster 7 (64 individuals) had accumulated the most unique deleterious mutations (1,720, 7.39%) (Figure 6a). Moreover, among the 23,290 mutations, 10,364 (44.50%) were homozygous in at least one individual (Table S26). A considerable number of observed homozygous deleterious mutations (1,712, 16.85% of the total homozygous sites) were shared by more than 50 individuals (Table S26).

We found a negative correlation between the number of deleterious mutations and effective population size among the 28 populations of *A. pentaphyllum* (DEL vs *N*_e_: Pearson’s *R* = –0.27, *P* < 4.8e-5; LOF vs *N*_e_: Pearson’s *R* = –0.2, *P* < 2.5e-3; Figure 6b, Figure S19), suggesting that smaller populations tend to accumulate more deleterious mutations. A similar trend was observed with census population size, although it was less pronounced and not as significant (DEL vs *N*_c_: Pearson’s R = –0.077, P = 0.25; LOF vs *N*_c_: Pearson’s R = –0.15, P = 0.023; Figure S20). For example, samples from CDG and TKX had higher inbreeding (*F*_ROH_: 27.64% and 29.87%, Figure 6c) than the average of all populations, and carried significantly more deleterious derived alleles (DEL: 3518 and 3800; RADICAL: 913 and 967; LOF: 236 and 277) compared to the remaining individuals of *A. pentaphyllum* (Figure 6c, Figure 6d, Figure S21, Table S21, Table S27). The large number of derived alleles was usually caused by a high number of homozygous sites (Chen *et al*., 2020). TKX and CDG carried 18–86% more homozygous DEL alleles, more homozygous RADICAL alleles, and 22% more homozygous LOF alleles than other populations, indicating high genetic load in these two populations of *A. pentaphyllum* (Figure 6e, Figure S22, Table S27). However, despite having a high *F*_ROH_ (31.84%), with small population size (only 30 mature individuals surveyed) and low nucleotide diversity (8.63 ± 8.88 × 10^−4^, Table 1, Table S22), the TZC population from cluster 3 exhibited the fewest number of deleterious mutations (DEL: 2045; RADICAL: = 561; LOF: 133) and most of them were present in a heterozygous state (DEL: 1092, 53.40%; RADICAL: 301, 53.65%; LOF: 71, 53.38%) (Figure 6c, Figure 6d, Figure S23, Table S27). However, in cluster 6, some populations (e.g., LTG) showed high *F*_ROH_ and numerous deleterious mutations, while others (e.g., BYP) displayed high *F*_ROH_ but few deleterious mutations. Additionally, some populations (e.g., GA) had low *F*_ROH_ but many deleterious mutations (Figure 6c, Figure S21, Table S27). These varying patterns indicate that different demographic histories and isolation levels among these populations have resulted in contrasting levels of inbreeding and genetic load, despite their central distribution within the overall range.

The relative ratio (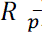) of the mean derived alleles frequency reflects the strength of deleterious mutation purging for each population (Figure S24a, Table S28). By calculating 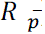 for three types of mutation, all populations of *A. pentaphyllum* exhibited similarly low efficiency (mean 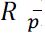 =1.01) in purging DEL deleterious mutations (Figure S24a, Table S28). TKX showed the strongest purging ability against RADICAL (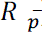 = 0.91, Figure S24b) mutations but the weakest ability to remove extreme deleterious mutations (LOF: 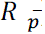 = 0.98, Figure 6f), while the TES population excelled in removing RADICAL (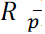 = 0.93) and LOF (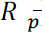 = 0.75) sites (Figure 6f, Figure S24b, Table S28).

GO enrichment analysis of genes containing deleterious mutations in *A. pentaphyllum* revealed significant enrichment (Q <0.05) in the structure and function of membranes (GO: 0016021, GO: 0016020, GO: 0022857), ATP binding (GO: 0005524), protein activity (GO: 0004672, GO: 0004674, GO: 0016887, GO: 0016301, GO: 0004252), molecular binding (GO: 0008270, GO: 0000166, GO: 0003676, GO: 0046872), response to heat (GO:0009408), recognition of pollen (GO: 0048544), among others (Figure 7a, Table S29). Further KEGG pathway analysis for these loci showed that they were mainly enriched (Q <0.5) in carbohydrate and fatty acid metabolisms, such as the glycolysis/gluconeogenesis (ko00010), galactose metabolism (ko00052), fatty acid elongation (ko00062) and TCA cycle (ko00020) (Figure 7b, Table S30).

**Figure 7.**
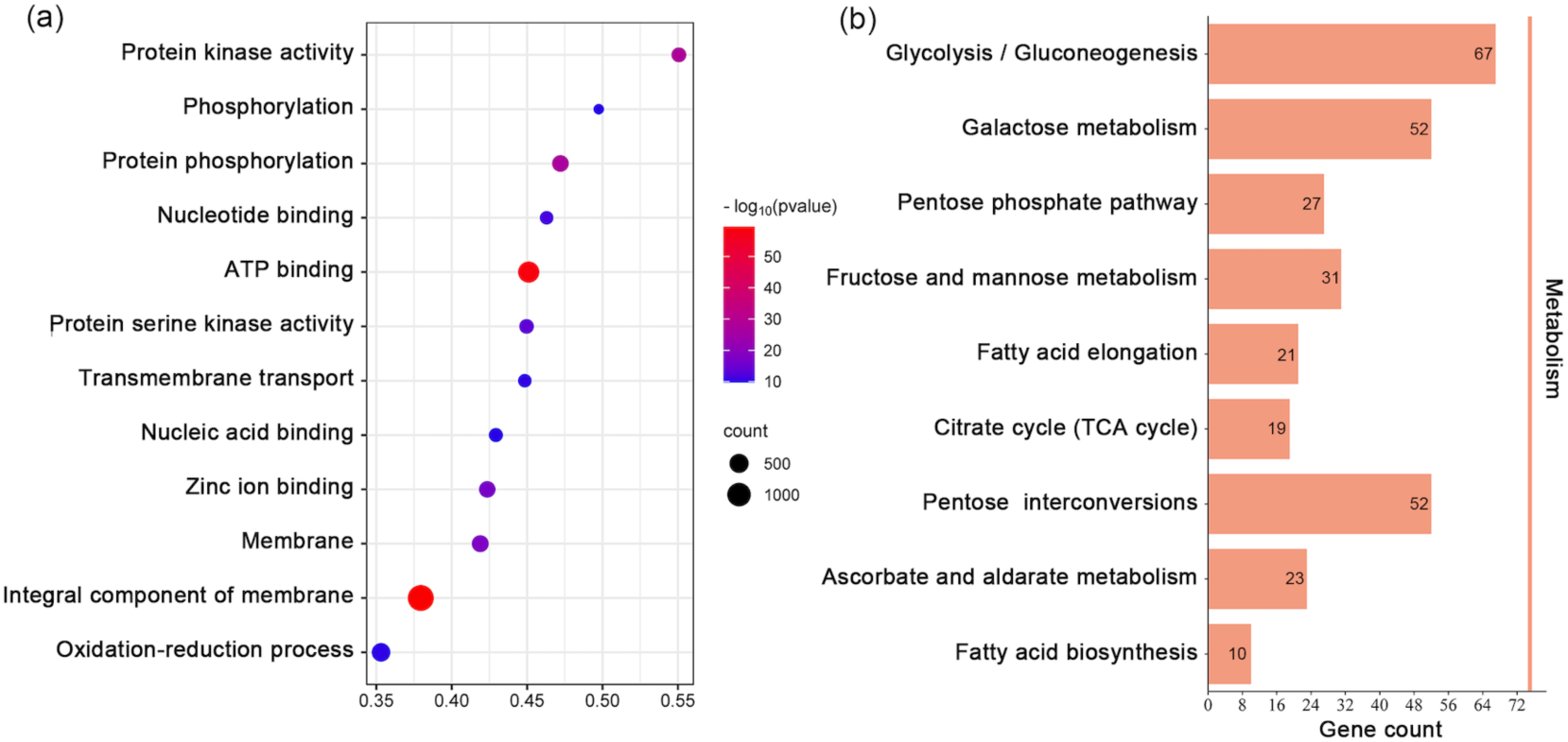
GO (a) and KEGG (b)enrichment diagram of genes containing deleterious mutations.

## Discussions

### The evolving genome of the endangered maple tree

*A. pentaphyllum* is one of the world’s most elegant maples, possessing significant potential for both horticultural applications and conservation efforts. However, its small population size and endangered status underscore the urgent need for comprehensive research. Until now, the lack of genomic data has impeded progress in conservation genetics for this species. In this study, we presented for the first time a high-quality chromosomal-scale genome assembly for the narrow endemic *A. pentaphyllum* (∼626 Mb) constructed using PacBio long HiFi reads and Hi-C high-accuracy sequencing. Taken together previously published *Acer* genomes, we found that abundant repetitive elements are a shared characteristic within the genus. For instance, repetitive sequences comprise 70.64% of the *A. pentaphyllum* genome (Table S5), similar to *A. yangbiense* (68.9% in a ∼666 Mb genome; Yang *et al*., 2019) and *A. pseudosieboldianum* (76.84% in a ∼690.24 Mb genome; Li *et al*., 2022). Notably, all these species have undergone only the ancient γ hexaploidization event shared by core eudicots, with no evidence of recent whole-genome duplications.

However, the *A. pentaphyllum* genome exhibits evidence of significant recent genomic changes through rapid amplification of transposable elements (TEs), particularly long terminal repeat retrotransposons (LTR-RTs). This amplification is evident from the presence of short syntenic blocks in the genome over the past 10 million years (Figure S4). Notably, this timeframe coincides with a period of extensive *in situ* speciation in the Hengduan Mountains (Xing and Ree, 2017). This burst of speciation was likely triggered by the rapid and recent uplift of the Hengduan Mountains and the formation of its distinctive landscape, characterized by deep valleys and steep slopes, occurring between the late Miocene and late Pliocene (Xing and Ree, 2017, Ding *et al*., 2020). The dramatic changes in climate and environment associated with this geological activity may have been a driving force behind the burst of transposable element activity in the *A. pentaphyllum* genome. This recent surge in transposon proliferation has played a crucial role in shaping the current genomic landscape of *A. pentaphyllum* (Belyayev, 2014, Canapa *et al*., 2015). Such genomic restructuring may have significantly contributed to increased genomic variation and enhanced adaptability, underscoring the importance of transposable elements in the species’ evolutionary dynamics.

*A. pentaphyllum* thrives in the dry-hot valley of the middle Yalong River, experiencing distinct wet and dry seasons, with low rainfall (less than 6% of the annual total) and cold temperatures (–15.6°C) during the dry season (Luo *et al*., 2017). Comparative genomic analyses revealed that several genes involved in photosynthesis, plant hormone signal transduction, and membrane transport were selectively expanded and/or positively selected in the genome during the processes of evolution (Figure S8, Table S10, Table S11, Table S12), potentially as candidates for adaptation of *A. pentaphyllum* to high-temperature and low-humidity valley environment.

### The special evolutionary history of *A. pentaphyllum*

Distributing in the deep valleys surrounded by high mountains, a subconscious colonization route of *A. pentaphyllum* is thought to be from the upstream to the downstream of the Yalong River, because wind may not be able to facilitate its seeds to fly over the high mountains but water is able to disperse the seeds for at least 250 km (Tessier, 2019). Interestingly, our population genomic analysis revealed a contrast pattern: CDG and TKX, representing the most downstream populations of *A. pentaphyllum* positioned the basal of the phylogenetic tree, while other upstream populations possessed derived lineages, suggesting an upstream colonization before/during the lineage diversification of this species (Figure 4c). The lineage diversification was dated back to 1–2 Mya (Figure 5c), indicating that before this, the intensify of the East Asia monsoon during Late Pliocene/Early Pleistocene (2.89–2.34 Mya, Yao *et al*., 2012) may drive the formation and/or northward expansion of the hot-dry valley (Ma and McConchie, 2001, Li *et al*., 2019, Liu *et al*., 2020). Consequently, this local climate change likely triggered a progressive northward shift in suitable habitat for *A. pentaphyllum*, facilitating its gradual spread into the Kangding and Yajiang county regions.

Interestingly, many populations on opposite sides of the river often exhibited similar and mixed genetic compositions. This was particularly evident in populations from Sanyanlong and Bawolong townships of Jiulong County (e.g., BYP, QJG) and Bowo Township of Muli County (e.g., ZD), which clustered together on the phylogenetic tree. This pattern could be attributed to the paired samara structure of *A. pentaphyllum* seeds, which, facilitated by valley winds, may enable cross-river dispersal (Lee *et al*., 2014, Wu *et al*., 2021). These findings suggest that despite its distribution along the valley, the Yalong River appears neither to facilitate long-distance dispersal nor to significantly impede the spread of *A. pentaphyllum* seeds. Instead, the species’ colonization pattern seems more influenced by historical climate changes and local wind-mediated dispersal.

The demographic history inferred by PSMC detected two genetic bottlenecks within the past million years. The earliest occurred at approximately 1.1 to 0.8 Mya and may have been caused by the Early to Middle Pleistocene transition, a period of extremely low temperatures marked by a series of glacial and interglacial cycles, resulting in increased global ice volume and further intensified aridity and monsoon strength in Asia (Legrain *et al*., 2023). Population declines at a similar time are also reported in many species (Yang *et al*., 2019, Ma *et al*., 2021, Liu *et al*., 2024) including human ancestors, 98.7% of whom were lost at the beginning of this bottleneck, thus threatening our ancestors with extinction (Hu *et al*, 2023). The most recent bottleneck occurred at the Penultimate Glacial Period (∼0.36-0.13 Mya), with the population size and genetic diversity of *A. pentaphyllum* experiencing sharp reductions. After this population decline, the *Ne* of *A. pentaphyllum* inferred in the Stairway plot (Figure 5d) and fastsimcoal2 (Figure 5b) recovered and maintained a stable size until recently, which is coincident with the gradual increase in the temperature and humidity of the global climate since the Holocene (∼12 kya). Multiple population declines may have resulted in the narrow distribution pattern of *A. pentaphyllum*. In addition, the significantly lower census population size (*Nc* ≈ 1.65×10^4^) investigated in the field than the contemporary *Ne* (∼1.19×10^6^) indicated that strong pressures have recently been imposed on *A. pentaphyllum* (Table S22, Ferchaud *et al*., 2016). Contemporary intensifying anthropogenic activities (e.g., logging), which accelerated habitat fragmentation and population isolation, are likely to have been responsible for the current endangered status of *A. pentaphyllum*.

### Impact of inbreeding on genetic load in small populations

Small populations are more susceptible to inbreeding depression due to their low numbers of individuals, leading to an increased probability of fixing homozygous deleterious mutations in the genome and resulting in an overall loss of diversity (Lynch *et al*., 1993, Höglund 2009, Agrawal and Whitlock, 2012, Teixeira and Huber, 2021, Robinson *et al*., 2023). While seriously deleterious alleles can be effectively purged during population contractions, prolonged bottlenecking leads to their gradual accumulation (Xie *et al*., 2021). This demographic stress, combined with inbreeding, elevates the overall genetic load and reduces genomic variation, ultimately impairing the species’ ability to adapt to environmental changes. The higher genetic load observed in small populations such as TKX, CDG, LM, and TES may be attributed to higher level of recent inbreeding, as evidenced by the higher total length of ROH and more related individual pairs (Figure 6c, Table S19, Table S20). Furthermore, genetic isolation among adjacent populations, even within the core distribution area (e.g., cluster 6), is evident from the distinct genetic structure (Figure S11) and varying patterns of inbreeding and mutation load across clusters (Figure 6). This phenomenon suggests that habitat fragmentation may have resulted in small, isolated populations with reduced capacity to purge deleterious mutations and decreased adaptive potential, potentially leading to their gradual extinction (e.g., Feng et al., 2024). Functional annotation indicated that genes containing deleterious mutations were enriched in energy synthesis and transformation primarily affects processes, which encompasses nearly all biological processes of an organism. The accumulation of these deleterious mutations would potentially reduce the adaptive potential of *A. pentaphyllum* and further affecting its long-term survival.

### Strategies for conservation management units and genetic rescue

The identification of Management Units (MUs) is crucial for the management of natural populations. Obviously, populations TKX and CDG, located lowest reaches of the entire distribution range of *A. pentaphyllum*, exhibited the lowest nucleotide diversity (Table 1), with relatively high *F*_ROH_ and more homozygous deleterious mutations (Figure 6). These two populations showed pure ancestry with no admixture at all values of *K* in the admixture analysis (Figure S11), likely due to long-distance isolation from other populations (TKX vs others: ∼25-129 km; CDG vs others: ∼26-180 km, Table S18), which may decrease opportunities for pollen and seed dispersal. Furthermore, TKX and CDG represent two ancient diverged lineages of *A. pentaphyllum*, necessitating consideration as two separate evolutionarily significant units (ESUs). Similarly, LM was genetically isolated (Figure 4d, Table S18), with low nucleotide diversity and high recent inbreeding (Figure 6a, Table 1, Table S19), and deserves to be treated as another ESU. The TES population is rather peculiar, exhibiting different genetic components from neighboring populations when K > 8 and forming a lineage with AJ. In addition, this population is distributed on a small hillside, with few individuals, low genetic diversity (Table 1, Table S27), high inbreeding (Figure 6c, Table S19, Table S20), and large number of homozygous DEL variants (Figure 6e), which needs further conservation action. The nearest MLC seemed to be the ideal source population for TES, due to their similar genetic background (*F*_ST_ = 0.148) and relatively lower number of shared deleterious mutations. We recommend introducing seedlings germinated from seeds produced by hand-cross pollination in MLC into the TES population to increase genetic diversity in this small population.

## Conclusions

In conclusion, this study presents the first chromosome-level reference genome for *A. pentaphyllum*, critically endangered but adapted to dry-hot valleys. Population genomic analyses uncovered the genetic mechanisms endangering *A. pentaphyllum*. Genetic bottlenecks resulting from historical climate changes and geological events, as well as habitat fragmentation due to anthropogenic activities are likely the main reasons for the current occurrence of small, isolated populations and low genetic diversity in this species. The high inbreeding and high genetic load indicate low adaptive potential in the wild populations, which provides guidance for the conservation of this endangered species. Our high-quality reference genome and extensive population data of *A. pentaphtyllum* will facilitate its conservation by providing valuable genetic information and efficient genetic markers for future horticultural development.

## Materials and Methods

### Plant materials and field investigations

An *ex situ* conserved plant of *A. pentaphyllum* growing at the Chengdu Institute of Biology (CIB), Chinese Academy of Sciences, was used for reference genome construction. This tree was grown from a seed in 2018, which was originally collected from Muli County, Liangshan, Sichuan. Fresh young leaves were collected for DNA extraction, library preparation, and whole genome *de novo* sequencing. Two tissues (leaves and stems) from this individual were sampled for transcriptome sequencing. Fresh young flowers were obtained from Pusharong Town, Kangding County, Sichuan in 2022. The materials were collected and immediately frozen using liquid nitrogen and then stored at –80 ℃ for subsequent DNA and RNA extraction.

For the population material, our team conducted a comprehensive survey for all known distribution areas (including Yajiang, Kangding, Jiulong, and Muli counties) of *A. pentaphyllum* from 2021 to 2022 to estimate their census size. We recorded the number of individuals for each population found based on direct measurement methods, finally identifying 28 populations from which 227 individuals were collected and preserved using silica gel for resequencing (Table S13).

### Library preparation and sequencing

High-quality genomic DNA was extracted from the tender leaves of *A. pentaphyllum* using a modified CTAB method (Porebski *et al*., 1997). To acquire an accurate genome assembly, several sequencing platforms were utilized, including Sequel II (Pacific Biosciences, Menlo Park, CA, USA) and Novaseq 6000 (Illumina, San Diego, CA, USA). For chromosome conformation capture sequencing, Hi-C libraries were constructed from tender leaves and sequenced using the MGISEQ-2000 system (BGI, Shenzhen, China). RNA was extracted from different tissues to create cDNA libraries using TRIzol reagent (Invitrogen, Carlsbad, CA, United States). The libraries were further purified and quality control (QC) to obtain Iso-Seq SMRT bell libraries, which were sequenced using Novaseq 6000. Raw sequencing data were filtered with fastp v0.23.2 (Chen *et al*., 2018) to remove low-quality reads, adapter contamination, and PCR duplicates before further analysis.

### *De novo* assembly, assessment, and annotation of reference genome

In this study, we employed a strategy of ‘assembling primarily with third-generation data and error correction with second-generation data’ to assemble the genome of *A. pentaphyllum*. Initially, HiFiasm v0.16.1 (Cheng *et al*., 2021) software was used to assemble filtered PacBio long reads with the parameter ‘––hom-cov 46’, yielding a primary assembly. The primary assembly was then polished with the Illumina short reads using Pilon v1.24 (Walker *et al*., 2014) for three iterations to obtain a draft genome of *A. pentaphyllum*. Finally, Hi-C reads were aligned to the draft genome using BWA-MEM v0.7.17 (Li, 2013), and contigs were clustered at the chromosome level using Lachesis (Burton *et al*., 2013) based on interaction information recorded in the data, generating the chromosome-level reference genome of *A. pentaphyllum*.

To assess the conservation, completeness, and consistency of the reference genome of *A. pentaphyllum*, we employed the Minimap v2.1 (Li, 2016) to align the Illumina short reads to the genome, and SAMTools v1.6 (-flagstat) (Li *et al*., 2009) was used to statistics the mapping rate, coverage, and average sequencing depth. Concurrently, the Benchmarking Universal Single-Copy Orthologs5 (BUSCO5) test was performed to evaluate the quality of the assembly using the Eudicots_odb10 database (https://buscodata.ezlab.org/v5/data/lineages/), which contains 2326 conserved eukaryotic genes (Simão *et al*., 2015).

The genomic components, including repetitive elements, gene structures, and non-coding RNAs (ncRNA), were annotated through a combination of *ab initio* prediction, homology-based alignment, and transcripts from different tissues (see details in Appendix S2).

### Comparative genome analysis

We performed comparative genomic analyses using the genomic sequences of *A. pentaphyllum* and 14 related published plant species, which represent 5 orders, including Sapindales, Rhoeadales, Rutales, Myrtales, and Rhamnales: *A. yangbiense*, A*. truncatum, A. catalpifolium*, *A. saccharum, A. negundo*, *A. pseudosieboldianum*, *Dipteronia dyeriana*, *Dipteronia sinensis* (Feng *et al*., 2024), *Arabidopsis thaliana* (Cheng *et al*., 2017), *Citrus sinensis* (Wang *et al*., 2017), *Eucalyptus grandis* (Myburg *et al*., 2014), *Nephelium lappaceum* (Zhang *et al*., 2021), *Vitis vinifera* (Canaguier *et al*., 2017), and *Xanthoceras sorbifolium* (Liang *et al*., 2019). OrthoFinder v2.5.4 (Emms and Kelly, 2015) was used to group orthologous and paralogous genes among the 15 genomes based on a Markov Cluster Algorithm (MCL). A total of 1422 single-copy orthologous genes were shared by all 15 species and used to construct a maximum likelihood tree using RAxML (Stamatakis, 2014) and IQ-TREE v2 (Minh *et al*., 2020) software. The specific genes of *A. pentaphyllum* were annotated and functionally enriched by GO and KEGG analyses.

To estimate the divergence time between *Acer* plants, the MCMCtree program in PAML (Yang, 2007) was utilized, with 4 corrected divergence time points from the TimeTree website (http://www.timetree.org/): *A. yangbiense* vs *C. sinensis* (68 – 82.8 Ma), *A. yangbiense* vs *A. thaliana* (90 – 99.9 Ma), *A. yangbiense* vs *E. grandis* (100 – 128 Ma), and *A. yangbiense* vs *V. vinifera* (109 – 123.5 Ma), and one corrected divergence time point from the fossil records (McClain and Manchester, 2001, Feng *et al*., 2024): *Acer* vs *Dipteronia* (56 – 73 Ma, based on the earliest fossil records of the genus *Dipteronia* dating back to 63 Ma).

Based on the result of gene family clustering and ultrametric tree containing the divergence times, CAFÉ (De Bie *et al*., 2006) and CODEML (Yang, 2007) programs were used to calculate the number of gene families undergoing expansion, contraction, and positive selection. Genes that exhibited significant expansion, contraction, and strong positive selection in the *A. pentaphyllum* genome were further annotated and functionally enriched by GO and KEGG analyses (see details in Appendix S3).

### Inference of WGD events and LTR insertions

Whole-genome duplication (WGD) events are a ubiquitous phenomenon in plants (especially in angiosperms), contributing to variations in both the size and structure of the genome and providing an important driver for species evolution and speciation. We conducted all-vs-all BLAST (E-value = 1e-5) on the protein sequences of *A. pentaphyllum*, *A. yangbiense*, *A. pseudosieboldianum*, and *D. sinensis*. Next, JCVI v1.2.7 (Tang *et al*., 2008) software was employed to perform collinearity analyses among the 4 Sapindaceae species with the grape genome as a reference. Additionally, WGDI v0.5.9 (Sun *et al*., 2022) was utilized to calculate the synonymous substitution rates (*Ks*) of collinear genes, aiming to infer potential WGD events in the *A. pentaphyllum* genome.

To infer the history of TE activity and selective pressures against LTR-RTs in the *A. pentaphyllum* genome, we calculated the insertion time for each of the LTR-RTs. Firstly, the mutation rate (μ) of *A. pentaphyllum* was estimated using the formula μ = *K*_*s*_/2*T*, where *K*_*s*_ represents the synonymous substitution rate peak value for all paralogous genes and *T* is the estimated divergence time between two species that had been evaluated in the phylogenomic analyses outlined above, and thus μ was calculated to be 4.66 × 10^-9^ per site per year. Secondly, LRT pairs from the results of LTRharvest and LTR_FINDER were aligned using MUSCLE v5 (Edgar, 2022). After trimming gaps on the ends of aligned sequences, we assessed the distance between the two LTR-RTs for all LRT pairs, and the insertion times could be calculated using the inverse formula *T* = *K*_*s*_/2μ (Hu *et al*., 2011, Slotte *et al*., 2013).

### Genome mapping and SNP calling

The original resequencing data, including 227 individuals of *A. pentaphyllum* and 2 closely related species: *A. yangbiense* (PRJNA524417) and *D. sinensis* (PRJNA796066.), underwent filtering using fastp to discard the adaptors, duplicates, and low-quality sequences, with parameters of “––length_required 50 –-cut_window_size 4 –-cut_mean_quality 15”. After quality control, paired-end clean data were mapped to the reference genome of *A. pentaphyllum* using BWA-MEM (Li, 2013). SAMTools (Li *et al*., 2019) was then used to filter out the unaligned and low-quality reads. Only the bases with a quality score ≥ 20 and reads with a mapping quality score ≥ 20 were considered in the subsequent analyses. The MarkDuplicates module in Picard v1.1 (http://broadinstitute.github.io/picard/) was employed to mark and remove the duplicate reads generated during the sequencing process with default parameters.

The Genome Analysis Toolkit (GATK) pipeline v4.2.0.0 (Van der Auwera *et al*., 2013) was performed for variants calling and preliminary filtering with parameters of “QD < 2.0, MQ < 40.0, FS > 60.0, SOR > 3.0, MQRankSum < –12.5, ReadPosRankSum < –8.0”, obtaining a total of 4,775,335 high-quality SNPs (Dataset 1) with an average of 21,036 SNPs per individual. The resulting dataset was further filtered by utilizing VCFtools v0.1.15 (Danecek *et al*., 2011) software, involving the following steps: (1) removal of SNPs with a quality score < 30 (––minQ 30); (2) removal of non-bi-allelic SNPs and non-SNPs (––max-alleles 2 –-min-alleles 2); (3) removal of SNPs with a depth ≥ 2 times or ≤ 1/3 of the average sequencing depth (––maxDP 47 –-minDP 8); and (4) removal of SNPs with a missing rate > 10% (––max-missing 0.9), generating 3,818,766 SNPs (Dataset 2). Furthermore, SNPs with minor allele frequency lower than 0.01 (–– maf 0.01) in Dataset 2 were removed, and the remaining loci (2,757,997, Dataset 3) were used for downstream analysis. In addition, variable sites in Dataset 3 were annotated using ANNOVAR (Wang *et al*., 2010) software.

### Population genetic diversity and estimation of inbreeding

Lineage disequilibrium (LD) decay analysis was performed across the *A. pentaphyllum* genome using PopLDdecay v3.42 (Zhang *et al*., 2019) software, which allow us to obtain LD statistics results directly from SNP Dataset1. Different population genetic estimators, including nucleotide diversity (*π*), Tajima’s *D*, and fixation index of subdivision (*F*_ST_), were calculated using VCFtools with a sliding window size of 100 Kb.

To assess the inbreeding level among populations, based on SNP Dataset 2 generated previously, we first calculate the kinship coefficient between individuals of *A. pentaphyllum* to identify potential relatives or clones using a robust relationship inference algorithm KING v2.3.2 (Manichaikul *et al*., 2010). VCFtools (––het) was then used to calculate each individual’s inbreeding coefficients (FIS) based on their heterozygosity. Furthermore, we estimated the runs of homozygosity (ROHs) for each sampled tree using PLINK software with parameters of “––homozyg-density 10 –-homozyg-gap 100 –-homozyg-kb 100 –-homozyg-snp 10”. We assessed the fraction of ROHs on the genome (F_ROH_) using the formula 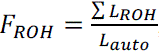, where 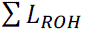 is the total length of ROHs that are longer than 100 Kb and have at least 10 SNPs in the genome and *L_auto_* is the total length of the *A. pentaphyllum* reference genome. F_ROH_ was calculated and were visualized via Manhattan plots using the qqman v0.1.9 (https://cran.rproject.org/web/packages/qq-man/) package.

### Population structure

PLINK v1.9 (Purcell *et al*., 2007) was used to filter out linked sites in Dataset 3 with the parameter of “––indep-pairwise 50 10 0.2”. After filtering, a total of 548,609 loci (Dataset 4) were obtained for inferring the population genetic structure of *A. pentaphyllum* using ADMIXTURE v1.3 (Alexander *et al*., 2009) software. The number of assumed ancestral populations (*K*) varied from 2 to 20 and the optimal *K* value was determined by the lowest cross-validation (CV) error. Then, GCTA v1.93.2 (Yang *et al*., 2011) was employed for performing principal component analysis (PCA) to detect population stratification and evolutionary relationships. Furthermore, we reconstructed an ML phylogenetic tree using RAxML with *A. yangbiense* and *D. sinensis* as outgroups.

### Lineage divergence and inference of demographic history

We employed the pairwise sequentially Markovian coalescent (PSMC) method (Li and Durbin, 2011) to reconstruct historical changes in effective population size (*Ne*) based on representative individuals for each cluster and population. The mutation rate was set as 4.66 × 10^-8^ per site per 10 years based on the previous calculation and the generation time of *A. pentaphyllum* was set as 10 years according to field observations (from seed to seed). To recover the demographic history and explore the role of gene flow in the process of speciation and lineage divergence of *A. pentaphyllum*, we conducted composite maximum likelihood (ML) inference in FASTSIMCOAL2 v2.6 (Excoffier et al. 2013) as complementary based on site frequency spectrum (SFS, see Appendix S4 for details). In addition, Stairway Plot 2 (Liu and Fu, 2020) was used to infer changes in population size through time based on the unfolded SFS of each group.

### Estimation of genetic load and deleterious mutations

To assess the potential influence of genetic load in *A. pentaphyllum*, we employed 3 different methods to detect deleterious mutations using polarized SNPs (filtering out sites where reference alleles inconsistent with ancestral states). We first calculated the Grantham distance (Grantham, 1974) for each nonsynonymous SNP, and radical missense substitution (RADICAL) was identified when the Grantham score >150. SIFT4G v2.0.0117 (Vaser *et al*., 2016) software was then performed to annotate the variant sites from vcf files using SIFT4G_Annotator.jar (https://github.com/pauline-ng/SIFT4G_Annotator). SNPs with a score <0.05 were assumed to be deleterious (DEL). In addition, we searched through the whole genome to predict severely deleterious mutations that caused start loss, stop gain, stop loss, and changes in splice sites with the help of annotation information. To further access the accumulation of severely deleterious mutations among the different populations and groups of *A. pentaphyllum*, SNPEFF v4.5 (Cingolani *et al*., 2012) was used to annotate SNPs leading to gene loss of function (LOF) with the parameter “-lof”.

Additionally, to assess the purging efficiency of deleterious mutations among different populations and clusters of *A. pentaphyllum,* we first calculated the mean ratio of derived alleles (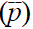) for each population using 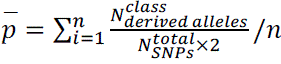 for all types of deleterious mutations, where 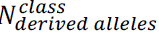 is the number of derived allelic genes for three types of mutations (DEL, RADICAL, and LOF), which is based on counting each heterozygous site once and each homozygous derived site twice for each individual, 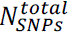 is the total number of SNPs used for deleterious mutations annotation, and *n* represents the number of individuals in a population of *A. pentaphyllum*. The mean ratio of derived alleles for all SNPs 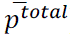 was used for correction, and the relative ratio of mutant alleles (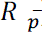) was calculated as 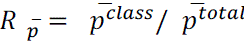 to account for inconsistencies in derived allele accumulation across different clusters. Here, 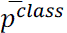 is the average derived alleles ratio for a specific type of harmful mutations, and 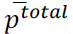 is the average derived allele ratio of the total number of SNPs. A lower relative ratio indicates a stronger purging efficiency of deleterious mutations.

## Acknowledgements.

We thank Min Liao, Junyi Zhang, Yushan Zhou, Shuo Tan, Cier Zhongyang, and Pianchu for assistance with sample collection. We thank Yongling Qiu for her help with writing. We also gratefully thank Wentai Dai for providing transcriptome data of leaves. This work was supported by the Wild Plants Sharing and Service Platform of Sichuan Province, the Western China Youth Scholars Project, and the China Postdoctoral Science Foundation under Grant Number 2022M713072.

## Author contributions

B.X. conceived the project. B.X. and Y.F. designed the research. X.L., Q.Y., H.N.D., L.S.J., Y.F. and W.B.J. collected the materials. X.L. and Y.F. analysed the data. X.L., Y.F., B.X., and W.B.J. contributed to the interpretation of results. X.L., Y.F. and L.S.J. wrote the manuscript, B.X. and Y.F. revised the manuscript. All authors read and approved the manuscript.

## Data availability

All data that support the findings of this study, including sequencing data, reference genome, and gene annotations, have been deposited into CNGB Sequence Archive (CNSA, Guo *et al*., 2020) of China National GeneBank DataBase (CNGBdb) with accession number CNP0006021.

## Supporting Information

### Supplementary methods

**Appendix S1** Estimation of genome features

**Appendix S2** Genome annotation

**Appendix S3** Positive selection analysis

**Appendix S4** Lineage divergence and inference of demographic history

### Supplementary Figures

**Figure S1** Flow cytometry analysis of *A. pentaphyllum*.

**Figure S2** Estimation of genome features of *A. pentaphyllum* based on 17-mer analysis.

**Figure S3** Hi-C intrachromosomal contact map of 13 chromosomes for *A. pentaphyllum*.

**Figure S4** Distribution of LTR transposons retrotransposons insertion times on the *A. pentaphyllum* genome *A. pentaphyllum*.

**Figure S5** Venn diagram of functional annotation for *A. pentaphyllum* based on 5 databases.

**Figure S6** Number of genes in 15 woody species

**Figure S7** Visualization of results from GO and KEGG enrichment analysis of 2254 significantly expanded genes in *A. pentaphyllum*.

**Figure S8** GO enrichment analysis of positive selected genes in *A. pentaphyllum*.

**Figure S9** Dot plots of synteny blocks among *A. pentaphyllum*, *A. yangbiense*, and *D*. *sinensis*.

**Figure S10** Synonymous substitution rate (*Ks*) distribution maps of synteny blocks between and within species.

**Figure S11** Population structure of 227 *A. pentaphyllum* individuals at *K* = 2 to *K* =9.

**Figure S12** The phylogenetic tree of *A. pentaphyllum* based on SNP Dataset 4 (different colors represent different clusters at *K* = 7).

**Figure S13** Genome-wide linkage disequilibrium (LD) decay in *A. pentaphyllum* among 28 separate populations and considering the ten populations as a whole.

**Figure S14** Genome-wide sequence diversity (π) for 17 threatened woody species.

**Figure S15** Mental test plot of genetic and geographic distances for *A. pentaphyllum*.

**Figure S16** Genetic differentiation level (*F*_st_) of each population for *A. pentaphyllum*

**Figure S17** Representative scenarios tested in this study. Asterisks besides parameters represent changes in population sizes.

**Figure S18** Illustration of the best-fit model with parameters names.

**Figure S19** Correlation analysis between the number of extreme deleterious mutations (LOF) and the effective population size (*N*_e_) across the 28 populations of *A. pentaphyllum*.

**Figure S20** Correlation analysis between the census population size (*N*_c_) and the number of deleterious mutations of DEL and LOF across the 28 populations of *A. pentaphyllum*.

**Figure S21** Number of DEL, LOF, and RADICAL mutations among 28 populations of *A. pentaphyllum*.

**Figure S22** Number of homozygous DEL, LOF, and RADICAL mutations among 28 populations of *A. pentaphyllum*.

**Figure S23** Number of heterozygous DEL, LOF, and SYN mutations among 28 populations of *A. pentaphyllum*.

**Figure S24** The relative ratio (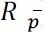) of the mean derived alleles frequency (*p*) for DEL and RADICAL mutations.

## Supplementary tables

**Table S1** Sequencing data used for the de novo assembly of *A. pentaphyllum*.

**Table S2** Flow cytometry analysis of *A. pentaphyllum*.

**Table S3** Statistics of genome assembly for *A. pentaphyllum* (chromosome level).

**Table S4** Summary of BUSCO evaluation of the *A. pentaphyllum* genome.

**Table S5** Classification statistics of repeat contents for *A. pentaphyllum*.

**Table S6** Integration statistics of structure annotation of *A. pentaphyllum*.

**Table S7** Classification statistics of non-coding RNA for *A. pentaphyllum*.

**Table S8** Results of gene functional annotation of *A. pentaphyllum* based on 5 databases.

**Table S9** Summary of the gene family clustering analyses for 15 woody species.

**Table S10** GO terms significantly enriched in significantly expanded gene families of *A. pentaphyllum* (Q < 0.05).

**Table S11** KEGG terms significantly enriched in significantly expanded gene families of *A. pentaphyllum* (Q < 0.05).

**Table S12** GO terms significantly enriched in positive selected genes of *A. pentaphyllum* (Q < 0.05).

**Table S13** Sampling information and genome resequencing statistics of the 227 *A. pentaphyllum* individuals.

**Table S14** Annotation results of SNP dataset 3 using Annovar.

**Table S15** CV errors for K value from 1 to 20.

**Table S16** Nucleic acid diversity of threatened plant species.

**Table S17** Genetic differentiation level (*F*_st_) matrix for 27 populations of *A. pentaphyllum*.

**Table S18** Geographic distance matrix for 27 populations of *A. pentaphyllum*.

**Table S19** Kinship coefficient between individual pairs.

**Table S20** Descriptive statistics of runs of homozygosity number (ROH) for each individual of *A. pentaphyllum*.

**Table S21** Descriptive statistics of runs of homozygosity number (ROH) for each population of *A. pentaphyllum*.

**Table S22** Table S22 Census population size (*N*_c_) and effective population size (*N*_e_) for each population of A. pentaphyllum

**Table S23** Likelihood comparison of all the demographic scenarios. Schematic illustrations of the scenarios are shown in Figure S18.

**Table S24** Likelihood comparison of all the demographic scenarios. Schematic illustrations of the scenarios are shown in Figure 5b.

**Table S25** Statistics on the number of three deleterious mutations.

**Table S26** The number of homozygous deleterious mutations shared by different numbers of individuals of *A. pentaphyllum*.

**Table S27** Statistical information on deleterious mutations in 227 individuals of *A. pentaphyllum*.

**Table S28** Statistical information on deleterious mutations among 28 populations of *A. pentaphyllum*.

**Table S29** GO terms significantly enriched in positive selected genes of *A.pentaphyllum* (Q < 0.05).

**Table S30** GO terms significantly enriched in positive selected genes of *A.pentaphyllum* (Q < 0.05).

## References

1. Agrawal, A.F. and Whitlock, M.C. (2012) Mutation Load: The Fitness of Individuals in Populations Where Deleterious Alleles Are Abundant. Annual Review of Ecology, Evolution, and Systematics, 43, 115-+.

2. Alexander, D.H., Novembre, J. and Lange, K. (2009) Fast model-based estimation of ancestry in unrelated individuals. Genome Research, 19, 1655–1664.

3. Belyayev, A. (2014) Bursts of transposable elements as an evolutionary driving force. Journal of Evolutionary Biology, 27, 2573–2584.

4. Bi, W., Gao, Y., Shen, J., He, C.N., Liu, H.B., Peng, Y., Zhang, C.H. and Xiao, P.G. (2016) Traditional uses, phytochemistry, and pharmacology of the genus *Acer* (maple): A review. Journal of Ethnopharmacology, 189, 31–60.

5. Burton, J.N., Adey, A., Patwardhan, R.P., Qiu, R., Kitzman, J.O. and Shendure, J. (2013) Chromosome-scale scaffolding of de novo genome assemblies based on chromatin interactions. Nature Biotechnology, 31, 1119–1125.

6. Canaguier, A., Grimplet, J., Di Gaspero, G., Scalabrin, S., Duchene, E., Choisne, N., Mohellibi, N., Guichard, C., Rombauts, S., Le Clainche, I., Berard, A., Chauveau, A., Bounon, R., Rustenholz, C., Morgante, M., Le Paslier, M.C., Brunel, D. and Adam-Blondon, A.F. (2017) A new version of the grapevine reference genome assembly (12X.v2) and of its annotation (VCost.v3). Genomics Data, 14, 56–62.

7. Canapa, A., Barucca, M., Biscotti, M.A., Forconi, M. and Olmo, E. (2015) Transposons, Genome Size, and Evolutionary Insights in Animals. Cytogenetic and Genome Research, 147, 217–239.

8. Chen, S.F., Zhou, Y.Q., Chen, Y.R. and Gu, J. (2018) fastp: an ultra-fast all-in-one FASTQ preprocessor. Bioinformatics, 34, i884–i890.

9. Chen, Y., Ma, T., Zhang, L.S., Kang, M.H., Zhang, Z.Y., Zheng, Z.Y., Sun, P.C., Shrestha, N., Liu, J.Q. and Yang, Y.Z. (2020) Genomic analyses of a “living fossil”: The endangered dove-tree. Molecular Ecology Resources, 20.

10. Chen, Z.Y., Ai, F.D., Zhang, J.L., Ma, X.Z., Yang, W.L., Wang, W.W., Su, Y.T., Wang, M.C., Yang, Y.Z., Mao, K.S., Wang, Q.F., Lascoux, M., Liu, J.Q. and Ma, T. (2020) Survival in the Tropics despite isolation, inbreeding and asexual reproduction: insights from the genome of the world’s southernmost poplar (*Populus ilicifolia*). The Plant Journal, 103, 430–442.

11. Chen, Z., Lu, X.Y., Zhu, L., Afzal, S.F., Zhou, J.B., Ma, Q.Y., Li, Q.Z., Chen, J.H. and Ren, J. (2023) Chromosomal-level genome and multi-omics dataset provides new insights into leaf pigmentation in *Acer palmatum*. International Journal of Biological Macromolecules, 227, 93–104.

12. Cheng, C.Y., Krishnakumar, V., Chan, A.P., Thibaud-Nissen, F., Schobel, S. and Town, C.D. (2017) Araport11: a complete reannotation of the *Arabidopsis thaliana* reference genome. The Plant Journal, 89, 789–804.

13. Cheng, H., Concepcion, G.T., Feng, X., Zhang, H. and Li, H. (2021) Haplotype-resolved de novo assembly using phased assembly graphs with hifiasm. Nature Methods, 18, 1–6.

14. Cingolani, P., Platts, A., Wang le, L., Coon, M., Nguyen, T., Wang, L., Land, S.J., Lu, X. and Ruden, D.M. (2012) A program for annotating and predicting the effects of single nucleotide polymorphisms, SnpEff: SNPs in the genome of Drosophila melanogaster strain w1118; iso-2; iso-3. Fly (Austin), 6, 80–92.

15. Danecek, P., Auton, A., Abecasis, G., Albers, C.A., Banks, E., DePristo, M.A., Handsaker, R.E., Lunter, G., Marth, G.T., Sherry, S.T., McVean, G., Durbin, R. and Genomes Project Analysis, G. (2011) The variant call format and VCFtools. Bioinformatics, 27, 2156–2158.

16. De Bie, T., Cristianini, N., Demuth, J.P. and Hahn, M.W. (2006) CAFE: a computational tool for the study of gene family evolution. Bioinformatics, 22, 1269–1271.

17. Ding, W.N., Ree, R.H., Spicer, R.A. and Xing, Y.W. (2020) Ancient orogenic and monsoon-driven assembly of the world’s richest temperate alpine flora. Science, 369, 578–581.

18. Do, R., Balick, D., Li, H., Adzhubei, I., Sunyaev, S. and Reich, D. (2015) No evidence that selection has been less effective at removing deleterious mutations in Europeans than in Africans. Nature Genetics, 47, 126–131.

19. Edgar, R.C. (2022) Muscle5: High-accuracy alignment ensembles enable unbiased assessments of sequence homology and phylogeny. Nature Communications, 13, 6968.

20. Emms, D.M. and Kelly, S. (2015) OrthoFinder: solving fundamental biases in whole genome comparisons dramatically improves orthogroup inference accuracy. Genome Biology, 16, 157.

21. Excoffier, L., Dupanloup, I., Huerta-Sánchez, E., Sousa, V.C. and Foll, M. (2013) Robust Demographic Inference from Genomic and SNP Data. PLOS Genetics, 9.

22. Feng, Y., Comes, H.P., Chen, J., Zhu, S.S., Lu, R.S., Zhang, X.Y., Li, P., Qiu, J., Olsen, K.M. and Qiu, Y.X. (2024) Genome sequences and population genomics provide insights into the demographic history, inbreeding, and mutation load of two ‘living fossil’ tree species of *Dipteronia*. The Plant Journal, 117, 177–192.

23. Ferchaud, A.L., Perrier, C., April, J., Hernandez, C., Dionne, M. and Bernatchez, L. (2016) Making sense of the relationships between *N*e, *N*b and *N*c towards defining conservation thresholds in Atlantic salmon (*Salmo salar*). Heredity, 117, 268–278.

24. González-Sarrías, A., Li, L. and Seeram, N.P. (2012) Effects of maple (Acer) plant part extracts on proliferation, apoptosis and cell cycle arrest of human tumorigenic and non-tumorigenic colon cells. Phytotherapy Research, 26, 995–1002.

25. Grantham, R. (1974) Amino acid difference formula to help explain protein evolution. Science, 185, 862–864.

26. Guo, X.Q., Chen, F.Z., Gao, F., Li, L., Liu, K., You, L.J., Hua, C., Yang, F., Liu, W.L., Peng, C.H., Wang, L., Yang, X.X., Zhou, F.Y., Tong, J.W., Cai, J., Li, Z.Y., Wan, B., Zhang, L., Yang, T., Zhang, M.W., Yang, L.L., Yang, Y.W., Zeng, W.J., Wang, B., Wei, X.F. and Xu, X. (2020) CNSA: a data repository for archiving omics data. Database (Oxford).

27. Hao, Y.Q, Luo, X.B. and Wang, X.L. (2019) Genetic diversity of the endangered *Acer pentaphyllum* Diels by ISSR analysis. Journal of Sichuan University (Natural Science Edition), 56, 161–166.

28. Höglund J. (2009) Evolutionary conservation genetics. Oxford University Press.

29. Hu, T.T., Pattyn, P., Bakker, E.G., Cao, J., Cheng, J.F., Clark, R.M., Fahlgren, N., Fawcett, J.A., Grimwood, J., Gundlach, H., Haberer, G., Hollister, J.D., Ossowski, S., Ottilar, R.P., Salamov, A.A., Schneeberger, K., Spannagl, M., Wang, X., Yang, L., Nasrallah, M.E., Bergelson, J., Carrington, J.C., Gaut, B.S., Schmutz, J., Mayer, K.F., Van de Peer, Y., Grigoriev, I.V., Nordborg, M., Weigel, D. and Guo, Y.L. (2011) The Arabidopsis lyrata genome sequence and the basis of rapid genome size change. Nature Genetics, 43, 476–481.

30. Lee, S.J., Lee, E.J. and Sohn, M.H. (2014) Mechanism of autorotation flight of maple samaras (*Acer palmatum*). Experiments in Fluids, 55, 1–9.

31. Legrain, E., Parrenin, F. and Capron, E. (2023) A gradual change is more likely to have caused the Mid-Pleistocene Transition than an abrupt event. Communications Earth & Environment, 4.

32. Li, D., Peng, Y., Zhang, Z., Hu, Y., Chen, L. and Hu, Y. (2019) Research Overview and Development Strategy Analysis of Rare and Endangered Plant *Acer pentaphyllum* Diels. Jiangxi Science, 37, 190–192+220.

33. Li, H. (2013) Aligning sequence reads, clone sequences and assembly contigs with BWA-MEM. arXiv e-prints.

34. Li, H. (2016) Minimap and miniasm: fast mapping and de novo assembly for noisy long sequences. Bioinformatics, 32, 2103–2110.

35. Li, H. and Durbin, R. (2011) Inference of human population history from individual whole-genome sequences. Nature, 475, 493–496.

36. Li, H., Handsaker, B., Wysoker, A., Fennell, T., Ruan, J., Homer, N., Marth, G., Abecasis, G., Durbin, R. and Genome Project Data Processing, S. (2009) The Sequence Alignment/Map format and SAMtools. Bioinformatics, 25, 2078–2079.

37. Li, X., Cai, K., Han, Z.M., Zhang, S.K., Sun, A.R., Xie, Y., Han, R., Guo, R.X., Tigabu, M., Sederoff, R., Pei, X.N., Zhao, C.L. and Zhao, X.Y. (2022) Chromosome-Level Genome Assembly for *Acer pseudosieboldianum* and Highlights to Mechanisms for Leaf Color and Shape Change. Frontiers in Plant Science, 13, 850054.

38. Li, Y.C., Zhang, Z., Ding, G.Q., Xu, Q.H., Wang, Y., Chi, Z.Q., Dong, J. and Zhang, L. (2019) Late Pliocene and early pleistocene vegetation and climate change revealed by a pollen record from Nihewan Basin, North China. Quaternary Science Reviews, 222.

39. Li, Z. and Barker, M.S. (2020) Inferring putative ancient whole-genome duplications in the 1000 Plants (1KP) initiative: access to gene family phylogenies and age distributions. Gigascience, 9, giaa004.

40. Liang, Q., Li, H., Li, S., Yuan, F., Sun, J., Duan, Q., Li, Q., Zhang, R., Sang, Y.L., Wang, N., Hou, X., Yang, K.Q., Liu, J.N. and Yang, L. (2019) The genome assembly and annotation of yellowhorn (*Xanthoceras sorbifolium* Bunge). Gigascience, 8.

41. Liu, F.L., Gao, H.S., Pan, B.T. and Li, Z.M. (2020) The genesis, age and its paleoclimatic significance of loess-like sediments in the Huatan section of the dry-hot valley of the Jinsha River. Journal of Desert Research, 2022, 42: 60–70.

42. Liu, Y., Cai, L. and Sun, W.B. (2024) Transcriptome data analysis provides insights into the conservation of a plant species with extremely small populations distributed in Yunnan province, China. BMC Plant Biology, 24.

43. Liu, X. and Fu, Y.X. (2020) Stairway Plot 2: demographic history inference with folded SNP frequency spectra. Genome Biology, 21, 280.

44. Lu, X.Y., Chen, Z., Liao, B.Y., Han, G.M., Shi, D., Li, Q.Z., Ma, Q.Y., Zhu, L., Zhu, Z.Y., Luo, X.M., Fu, S.L. and Ren, J. (2022) The chromosome-scale genome provides insights into pigmentation in *Acer rubrum*. Plant Physiology and Biochemistry, 186, 322–333.

45. Luo, X.B., Wang, X.L., Hao, Y.Q., Xiong, J. and Zeng, D.G. (2017) A Study of Species Diversity and Dominant Species Niche Characteristics of *Acer pentaphyllum* Community. Journal of Sichuan Forestry Science and Technology, 38, 79–83.

46. Lynch, M., Burger, R., Butcher, D. and Gabriel, W. (1993) The mutational meltdown in asexual populations. Journal of Heredity, 84, 339–344.

47. Ma, H., Liu, Y.B., Liu, D.T., Sun, W.B., Liu, X.F., Wan, Y.M., Zhang, X.J., Zhang, R.G., Yun, Q.Z., Wang, J.H., Li, Z.H. and Ma, Y.P. (2021) Chromosome-level genome assembly and population genetic analysis of a critically endangered rhododendron provide insights into its conservation. The Plant Journal, 107, 1533–1545.

48. Ma, H.C. and McConchie, J.A. (2001) The dry-hot valleys and forestation in southwest China. Journal of Forestry Research, 12, 35–39.

49. Manichaikul, A., Mychaleckyj, J.C., Rich, S.S., Daly, K., Sale, M. and Chen, W.M. (2010) Robust relationship inference in genome-wide association studies. Bioinformatics, 26, 2867–2873.

50. McClain, A.M. and Manchester, S.R. (2001) *Dipteronia* (Sapindaceae) from the Tertiary of North America and implications for the phytogeographic history of the Aceroideae. American Journal of Botany, 88, 1316–1325.

51. McEvoy, S.L., Sezen, U.U., Trouern-Trend, A., McMahon, S.M., Schaberg, P.G., Yang, J., Wegrzyn, J.L. and Swenson, N.G. (2022) Strategies of tolerance reflected in two North American maple genomes. The Plant Journal, 109, 1591–1613.

52. Minh, B.Q., Schmidt, H.A., Chernomor, O., Schrempf, D., Woodhams, M.D., von Haeseler, A. and Lanfear, R. (2020) IQ-TREE 2: New Models and Efficient Methods for Phylogenetic Inference in the Genomic Era. Molecular Biology and Evolution, 37, 1530–1534.

53. Myburg, A.A., Grattapaglia, D., Tuskan, G.A., Hellsten, U., Hayes, R.D., Grimwood, J., Jenkins, J., Lindquist, E., Tice, H., Bauer, D., Goodstein, D.M., Dubchak, I., Poliakov, A., Mizrachi, E., Kullan, A.R., Hussey, S.G., Pinard, D., van der Merwe, K., Singh, P., van Jaarsveld, I., Silva-Junior, O.B., Togawa, R.C., Pappas, M.R., Faria, D.A., Sansaloni, C.P., Petroli, C.D., Yang, X., Ranjan, P., Tschaplinski, T.J., Ye, C.Y., Li, T., Sterck, L., Vanneste, K., Murat, F., Soler, M., Clemente, H.S., Saidi, N., Cassan-Wang, H., Dunand, C., Hefer, C.A., Bornberg-Bauer, E., Kersting, A.R., Vining, K., Amarasinghe, V., Ranik, M., Naithani, S., Elser, J., Boyd, A.E., Liston, A., Spatafora, J.W., Dharmwardhana, P., Raja, R., Sullivan, C., Romanel, E., Alves-Ferreira, M., Kulheim, C., Foley, W., Carocha, V., Paiva, J., Kudrna, D., Brommonschenkel, S.H., Pasquali, G., Byrne, M., Rigault, P., Tibbits, J., Spokevicius, A., Jones, R.C., Steane, D.A., Vaillancourt, R.E., Potts, B.M., Joubert, F., Barry, K., Pappas, G.J., Strauss, S.H., Jaiswal, P., Grima-Pettenati, J., Salse, J., Van de Peer, Y., Rokhsar, D.S. and Schmutz, J. (2014) The genome of *Eucalyptus grandis*. Nature, 510, 356–362.

54. Pan, H.L., Feng, Q.H., L, T.L., He, F. and L, X.L. (2014) Discussion on Resource Condition and Protection Technique for Rare Endangered Species in Sichuan Province. Journal of Sichuan Forestry Science and Technology, 35, 41–46.

55. Porebski, S., Bailey, L.G. and Baum, B.R. (1997) Modification of a CTAB DNA extraction protocol for plants containing high polysaccharide and polyphenol components. Plant Molecular Biology Reporter, 15, 8–15.

56. Purcell, S., Neale, B., Todd-Brown, K., Thomas, L., Ferreira, M.A., Bender, D., Maller, J., Sklar, P., de Bakker, P.I., Daly, M.J. and Sham, P.C. (2007) PLINK: a tool set for whole-genome association and population-based linkage analyses. The American Journal of Human Genetics, 81, 559–575.

57. Robinson, J., Kyriazis, C.C., Yuan, S.C. and Lohmueller, K.E. (2023) Deleterious Variation in Natural Populations and Implications for Conservation Genetics. Annual Review of Animal Biosciences, 11, 93–114.

58. Roh, M.S., McNamara, W.A., Barnes, C., Yin, K. and Wang, Q. (2010) Genetic variations of *Acer pentaphyllum* based on AFLP analysis, seed germination, and seed morphology Kaipu Yin and Qian Wang Chengdu institute of biology. Acta Horticulturae, 885, 305–312.

59. Simão, F., Waterhouse, R.M., Panagiotis, I., Kriventseva, E.V. and Zdobnov, E.M. (2015) BUSCO: assessing genome assembly and annotation completeness with single-copy orthologs. Bioinformatics, 31, 3210–3212.

60. Slotte, T., Hazzouri, K.M., Agren, J.A., Koenig, D., Maumus, F., Guo, Y.L., Steige, K., Platts, A.E., Escobar, J.S., Newman, L.K., Wang, W., Mandakova, T., Vello, E., Smith, L.M., Henz, S.R., Steffen, J., Takuno, S., Brandvain, Y., Coop, G., Andolfatto, P., Hu, T.T., Blanchette, M., Clark, R.M., Quesneville, H., Nordborg, M., Gaut, B.S., Lysak, M.A., Jenkins, J., Grimwood, J., Chapman, J., Prochnik, S., Shu, S., Rokhsar, D., Schmutz, J., Weigel, D. and Wright, S.I. (2013) The Capsella rubella genome and the genomic consequences of rapid mating system evolution. Nature Genetics, 45, 831–835.

61. Stamatakis, A. (2014) RAxML version 8: a tool for phylogenetic analysis and post-analysis of large phylogenies. Bioinformatics, 30, 1312–1313.

62. Stevens, P.F. (2016) Angiosperm Phylogeny Website. Version 14. *Angiosperm Phylogeny Website. Version 13*.

63. Sun, P.C., Jiao, B.B., Yang, Y.Z., Shan, L.X., Li, T., Li, X.N., Xi, Z.X., Wang, X.Y. and Liu, J.Q. (2022) WGDI: A user-friendly toolkit for evolutionary analyses of whole-genome duplications and ancestral karyotypes. Molecular Plant, 15, 1841–1851.

64. Tang, H., Bowers, J.E., Wang, X., Ming, R., Alam, M. and Paterson, A.H. (2008) Synteny and collinearity in plant genomes. Science, 320, 486–488.

65. Teixeira, J.C. and Huber, C.D. (2021) The inflated significance of neutral genetic diversity in conservation genetics. Proceedings of the National Academy of Sciences of the United States of America, 118.

66. Tessier, J.T. (2019) Evidence of capacity for water dispersal in. Ecosphere, 10.

67. Van der Auwera, G.A., Carneiro, M.O., Hartl, C., Poplin, R., Del Angel, G., Levy-Moonshine, A., Jordan, T., Shakir, K., Roazen, D., Thibault, J., Banks, E., Garimella, K.V., Altshuler, D., Gabriel, S. and DePristo, M.A. (2013) From FastQ data to high confidence variant calls: the Genome Analysis Toolkit best practices pipeline. Curr Protoc Bioinformatics, 43, 11 10 11–11 10 33.

68. Vaser, R., Adusumalli, S., Leng, S.N., Sikic, M. and Ng, P.C. (2016) SIFT missense predictions for genomes. Nature Protocols, 11, 1–9.

69. Walker, B.J., Abeel, T., Shea, T., Priest, M., Abouelliel, A., Sakthikumar, S., Cuomo, C.A., Zeng, Q., Wortman, J., Young, S.K. and Earl, A.M. (2014) Pilon: an integrated tool for comprehensive microbial variant detection and genome assembly improvement. PLoS One, 9, e112963.

70. Wang, K., Li, M. and Hakonarson, H. (2010) ANNOVAR: functional annotation of genetic variants from high-throughput sequencing data. Nucleic Acids Research, 38, e164.

71. Wang, X., Xu, Y.T., Zhang, S.Q., Cao, L., Huang, Y., Cheng, J.F., Wu, G.Z., Tian, S.L., Chen, C.L., Liu, Y., Yu, H.W., Yang, X.M., Lan, H., Wang, N., Wang, L., Xu, J.D., Jiang, X.L., Xie, Z.Z., Tan, M.L., Larkin, R.M., Chen, L.L., Ma, B.G., Ruan, Y.J., Deng, X.X. and Xu, Q. (2017) Genomic analyses of primitive, wild and cultivated citrus provide insights into asexual reproduction. Nature Genetics, 49, 765–772.

72. Wu, H., Yan, L.P., Li, C.Z., Xia, Q., Zhou, X. and Zhao, B.Y. (2021) Morphological characteristics and wind dispersal characteristics of samara of common Acer species. Journal of Nanjing Forest University, 45, 103.

73. Xie, H.X., Liang, X.X., Chen, Z.Q., Li, W.M., Mi, C.R., Li, M., Wu, Z.J., Zhou, X.M. and Du, W.G. (2021) Ancient Demographics Determine the Effectiveness of Genetic Purging in Endangered Lizards. Molecular Biology and Evolution, 39.

74. Xing, Y. and Ree, R.H. (2017) Uplift-driven diversification in the Hengduan Mountains, a temperate biodiversity hotspot. Proceedings of the National Academy of Sciences of the United States of America, 114, E3444–E3451.

75. Xu, T.Z., Chen, Y.S., de Jong, P.C., Oterdoom, H.J. and Chang, C.S. (2008) Aceraceae. In: Wu, Z.Y., Raven, P.H. & Hong, D.Y. (Eds.) Flora of China. Beijing, China: Science Press; St. Louis, USA: Missouri Botanical Garden Press, pp. 515–553.

76. Yang, J., Wariss, H.M., Tao, L.D., Zhang, R.G., Yun, Q.Z., Hollingsworth, P., Dao, Z.L., Luo, G.F., Guo, H.J., Ma, Y.P. and Sun, W.B. (2019) De novo genome assembly of the endangered Acer yangbiense, a plant species with extremely small populations endemic to Yunnan Province, China. Gigascience, 8. giz085

77. Yang, J.A., Lee, S.H., Goddard, M.E. and Visscher, P.M. (2011) GCTA: A Tool for Genome-wide Complex Trait Analysis. The American Journal of Human Genetics, 88, 76–82.

78. Yang, Y., Ma, T., Wang, Z.F., Lu, Z.Q., Li, Y., Fu, C.X., Chen, X.Y., Zhao, M.S., Olson, M.S. and Liu, J.Q. (2018) Genomic effects of population collapse in a critically endangered ironwood tree *Ostrya rehderiana*. Nature Communications, 9, 5449.

79. Yang, Z. (2007) PAML 4: phylogenetic analysis by maximum likelihood. Molecular Biology and Evolution, 24, 1586–1591.

80. Yao, Y.F., Bruch, A.A., Cheng, Y.M., Mosbrugger, V., Wang, Y.F. and Li, C.S. (2012) Monsoon versus uplift in southwestern China––Late Pliocene climate in Yuanmou Basin, Yunnan. PLoS One, 7, e37760.

81. Yu, T., Hu, Y.H., Zhang, Y.Y., Zhao, R., Yan, X.Q., Dayananda, B., Wang, J.P., Jiao, Y.N., Li, J.Q. and Yi, X. (2021) Whole-Genome Sequencing of *Acer catalpifolium* Reveals Evolutionary History of Endangered Species. Genome Biology and Evolution, 13. evab271.

82. Zhang, C., Dong, S.S., Xu, J.Y., He, W.M. and Yang, T.L. (2019) PopLDdecay: a fast and effective tool for linkage disequilibrium decay analysis based on variant call format files. Bioinformatics, 35, 1786–1788.

83. Zhang, W.P., Lin, J.S., Li, J.G., Zheng, S.Q., Zhang, X.T., Chen, S., Ma, X.K., Dong, F., Jia, H.F., Xu, X.M., Yang, Z.Q., Ma, P.P., Deng, F., Deng, B., Huang, Y.J., Li, Z.J., Lv, X.Z., Ma, Y.Y., Liao, Z.Y., Lin, Z.C., Lin, J., Zhang, S.C., Matsumoto, T., Xia, R., Zhang, J.S. and Ming, R. (2021) Rambutan genome revealed gene networks for spine formation and aril development. The Plant Journal, 108, 1037–1052.

