## Supplementary material for "High-Resolution Genome Assembly and Population Genetic Study of the Endangered Maple *Acer pentaphyllum* (Sapindaceae): Implications for Conservation Strategies": Supplementary_figures_20240801.docx

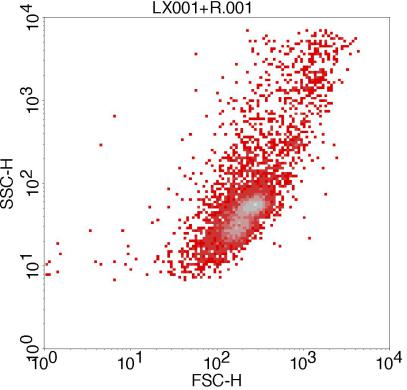

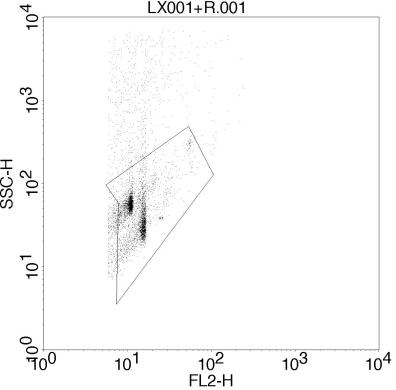

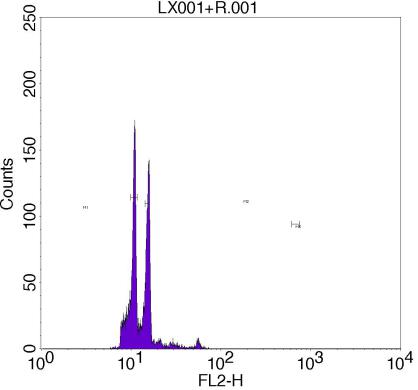

(a)

(c)

(b)

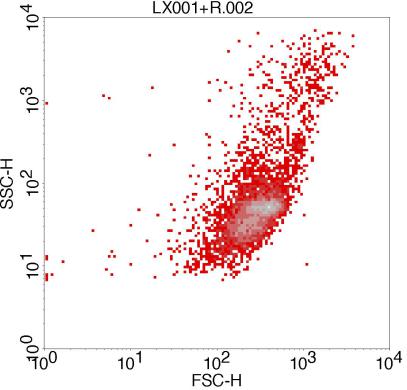

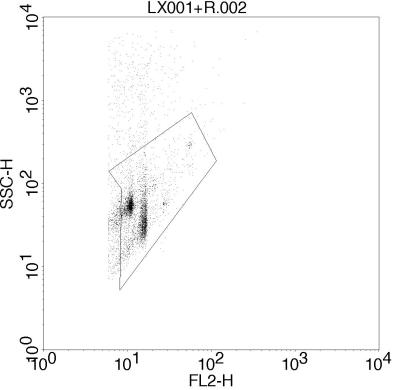

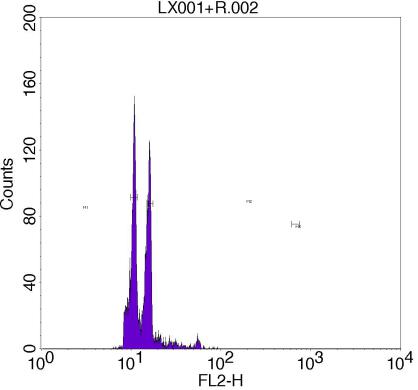

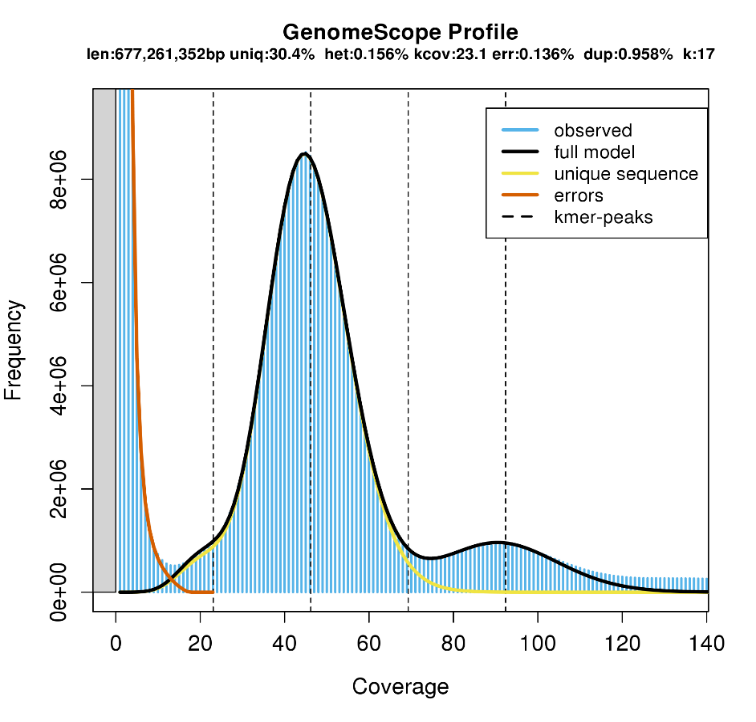
**Figure S1** Flow cytometry analysis of *A. pentaphyllum.* (a) Scatter plot of side scatter (SSC-H) versus forward scatter (FSC-H), indicating the size and granularity of the cells. (b) The gating strategy based on side scatter (SSC-H) versus fluorescence intensity (FL2-H), is used to identify specific cell populations. (c) Histogram of fluorescence intensity (FL2-H), illustrating the distribution of fluorescence within the gated cell population.

**Figure S2** Estimation of genome features of *A. pentaphyllum* based on 17-mer analysis

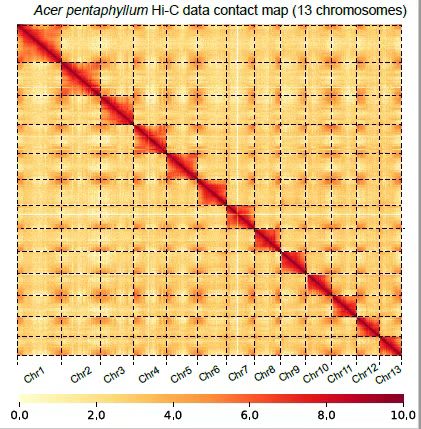

**Figure S3** A Hi-C intrachromosomal contact map of 13 chromosomes for *A. pentaphyllum.*

peak = 1.321

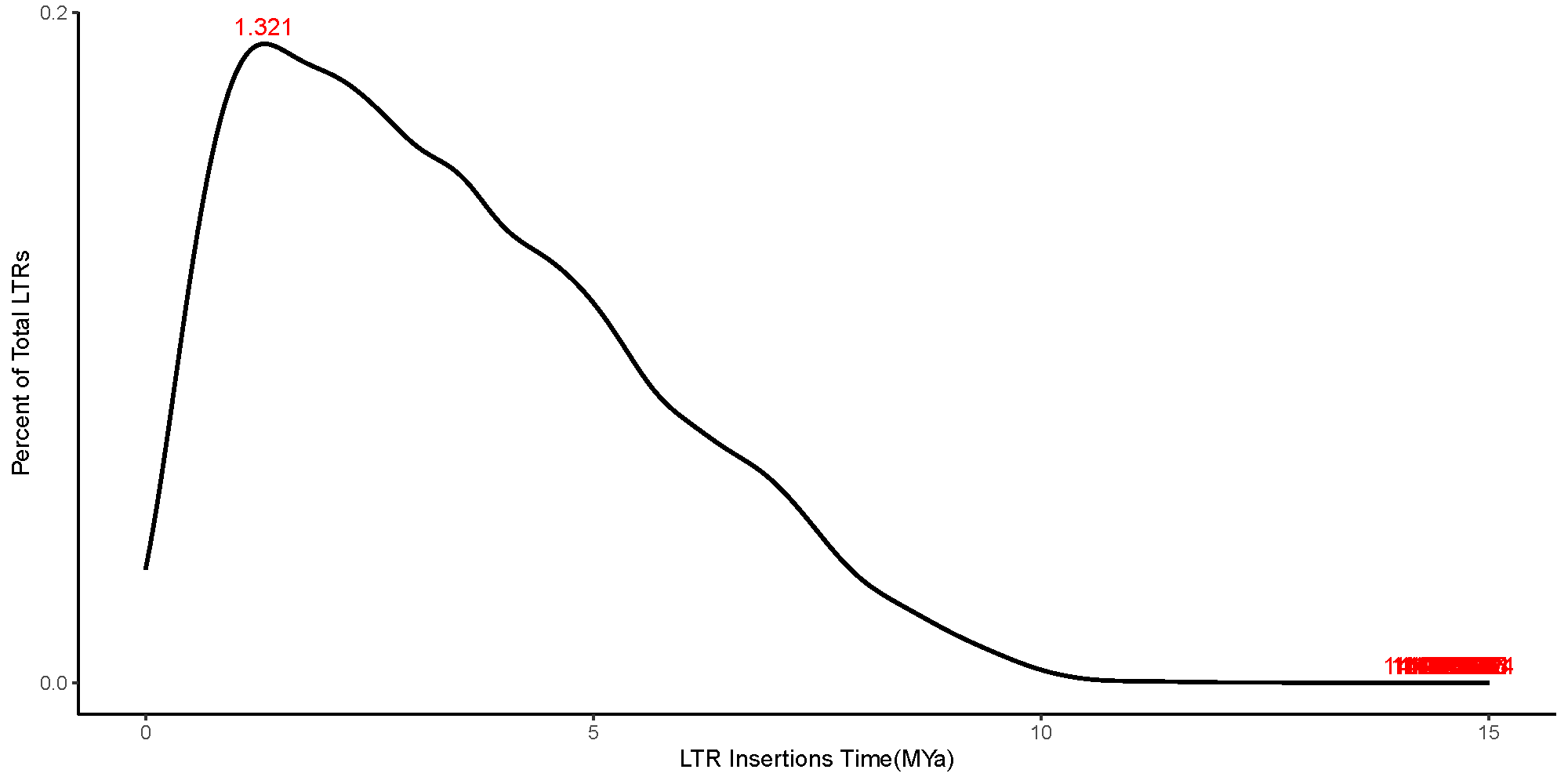

**Figure S4** Distribution of LTR transposons retrotransposons insertion times on the *A. pentaphyllum* genome *A. pentaphyllum.*

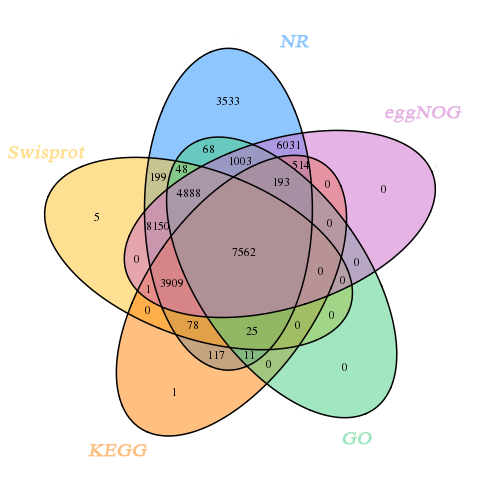

**Figure S5** Venn diagram of functional annotation for *A. pentaphyllum* based on 5 databases.

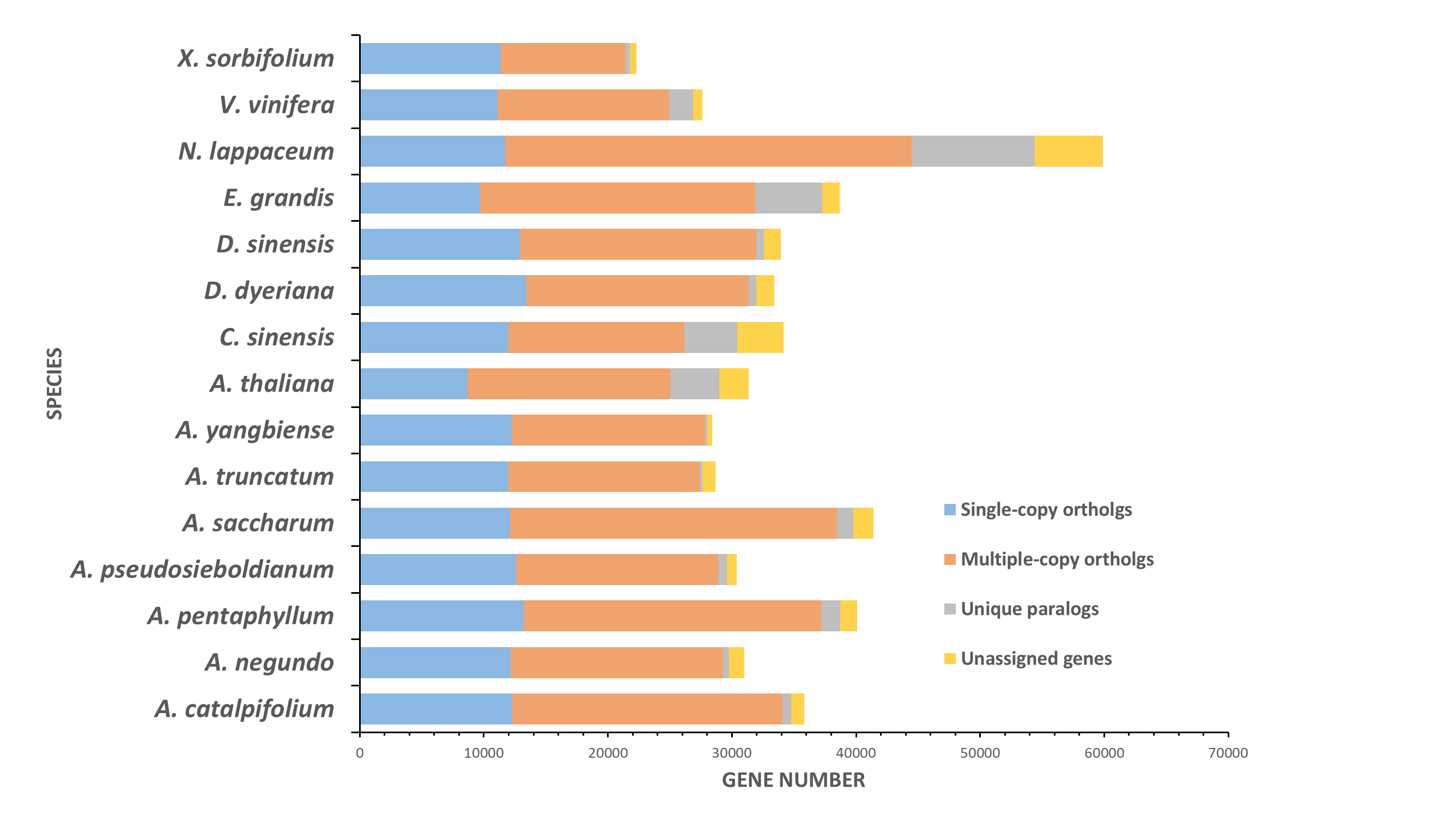

**Figure S6** Number of genes in 15 woody species

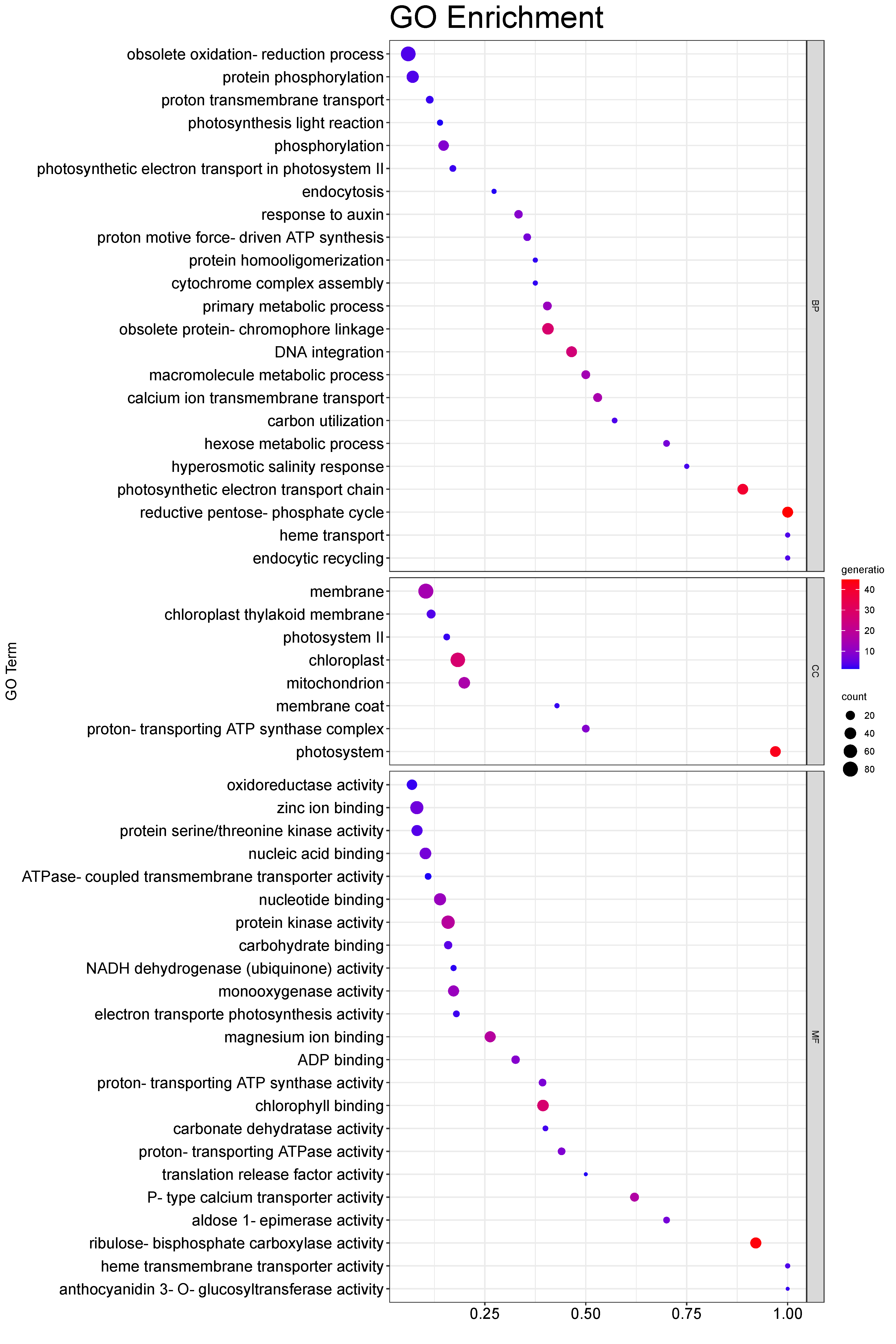

(a)

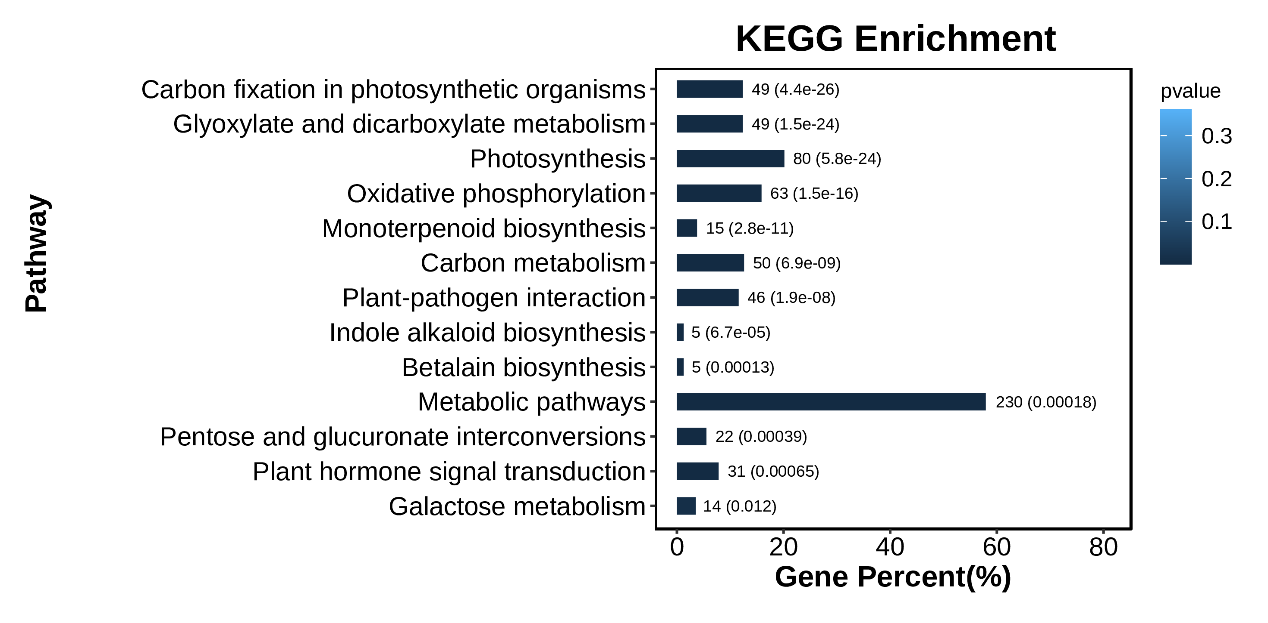

(b)

**Figure S7** Visualization of results from (a) GO and (b) KEGG enrichment analysis of 2254 significantly expanded genes in *A. pentaphyllum*. BP = biological process, CC = cellular component, MF = molecular function.

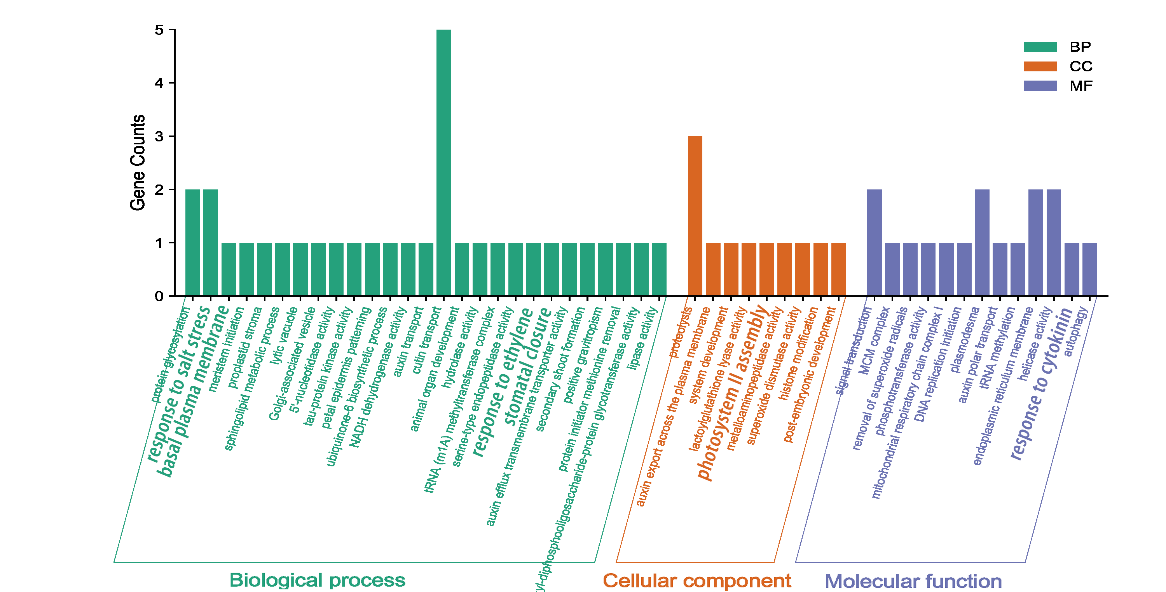

**Figure S8** GO enrichment analysis of positive selected genes in *A. pentaphyllum*.

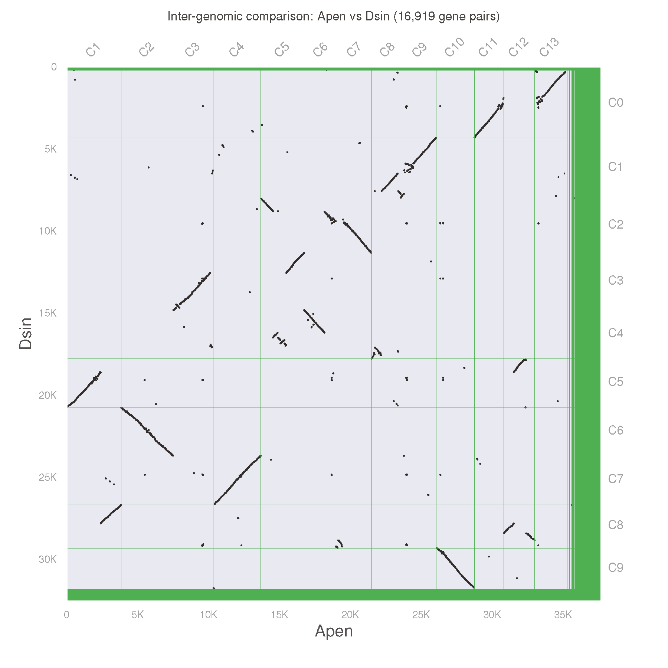

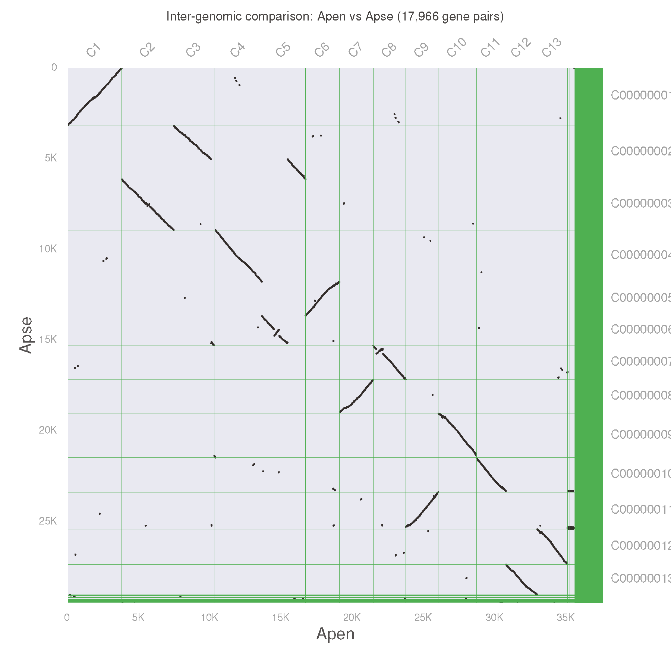
**Figure S9** Dot plots of synteny blocks among *A. pentaphyllum*, (a) *A. yangbiense*, and *D. sinensis*

(b)

(a)

b

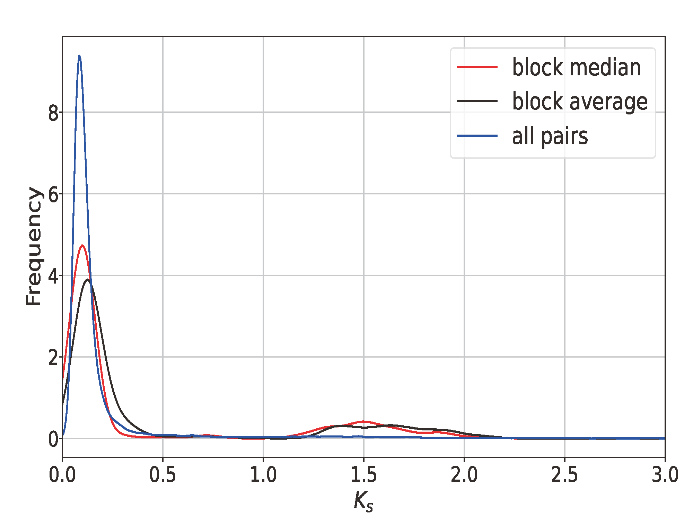

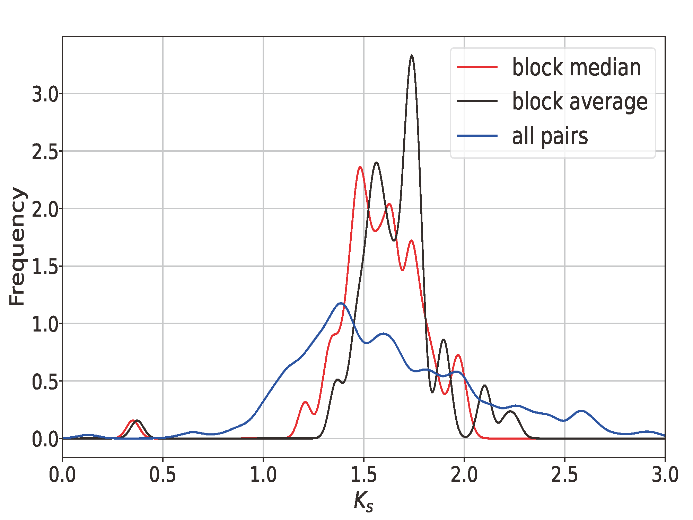

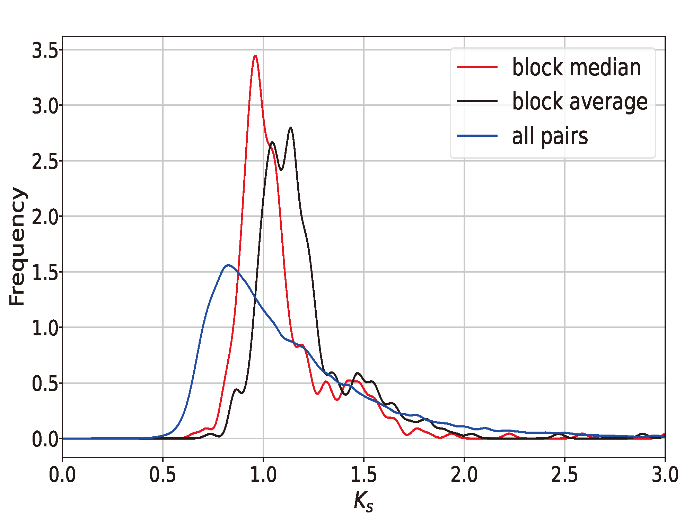

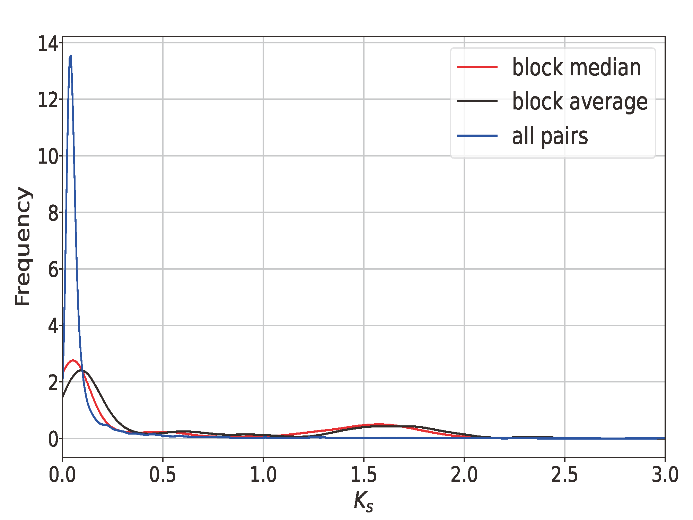
**Figure S10** Synonymous substitution rate (*Ks*) distribution maps of synteny blocks between and within species. (a) *A. pentaphyllum vs A. pentaphyllum*, (b) *A. pentaphyllum vs A. yangbiense*, (c) *A. pentaphyllum vs D. sinensis*, (d) *A. pentaphyllum vs V. vinifera*

(d)

(c)

(b)

(a)

K = 3

|  |  |  |  |
| --- | --- | --- | --- |
| CDG-1A | 1e-05 | 0.99998 | 1e-05 |
| CDG-2A | 1e-05 | 0.99998 | 1e-05 |
| CDG-3A | 1e-05 | 0.99998 | 1e-05 |
| CDG-4A | 1e-05 | 0.99998 | 1e-05 |
| CDG-5A | 1e-05 | 0.99998 | 1e-05 |
| CDG-6A | 1e-05 | 0.99998 | 1e-05 |
| CDG-7A | 1e-05 | 0.99998 | 1e-05 |
| CDG-8A | 1e-05 | 0.99998 | 1e-05 |
| CDG-9A | 1e-05 | 0.99998 | 1e-05 |
| CDG-10A | 1e-05 | 0.99998 | 1e-05 |
| TKX-1A | 1e-05 | 0.99998 | 1e-05 |
| TKX-2A | 1e-05 | 0.99998 | 1e-05 |
| TKX-3A | 1e-05 | 0.99998 | 1e-05 |
| TKX-4A | 1e-05 | 0.99998 | 1e-05 |
| TKX-5A | 1e-05 | 0.99998 | 1e-05 |
| TKX-6A | 1e-05 | 0.99998 | 1e-05 |
| TKX-7A | 1e-05 | 0.99998 | 1e-05 |
| TKX-8A | 1e-05 | 0.99998 | 1e-05 |
| TKX-9A | 1e-05 | 0.99998 | 1e-05 |
| GJ-1A | 0.051158 | 0.403193 | 0.545649 |
| GJ-2A | 0.053497 | 0.407612 | 0.538891 |
| GJ-3A | 1e-05 | 0.418796 | 0.581194 |
| GJ-4A | 0.002753 | 0.389096 | 0.608151 |
| GJ-5A | 1e-05 | 0.416757 | 0.583233 |
| GJ-6A | 1e-05 | 0.402995 | 0.596995 |
| GJ-7A | 0.029013 | 0.40602 | 0.564967 |
| GJ-8A | 1e-05 | 0.416368 | 0.583622 |
| GJ-9A | 1e-05 | 0.468239 | 0.531751 |
| TZC-1A | 1e-05 | 0.344703 | 0.655287 |
| TZC-2A | 1e-05 | 0.345475 | 0.654515 |
| TZC-3A | 0.023029 | 0.26789 | 0.709081 |
| TZC-4A | 1e-05 | 0.350194 | 0.649796 |
| TZC-01A | 0.051026 | 0.346648 | 0.602327 |
| TZC-04A | 0.068711 | 0.349128 | 0.582161 |
| TZC-07A | 0.092875 | 0.362315 | 0.54481 |
| TZC-11A | 0.009027 | 0.363337 | 0.627636 |
| TZC-13A | 0.008023 | 0.394318 | 0.597659 |
| DBG-10A | 1e-05 | 0.25045 | 0.74954 |
| DBG-1A | 0.11008 | 0.266671 | 0.623249 |
| DBG-2A | 0.068091 | 0.306416 | 0.625493 |
| DBG-3A | 0.055485 | 0.241859 | 0.702656 |
| DBG-4A | 0.048562 | 0.220369 | 0.731069 |
| DBG-5A | 0.025331 | 0.300155 | 0.674513 |
| DBG-6A | 0.080759 | 0.372615 | 0.546625 |
| DBG-7A | 0.061963 | 0.307991 | 0.630045 |
| DBG-8A | 0.023688 | 0.298636 | 0.677675 |
| DBG-9A | 0.009618 | 0.298943 | 0.691439 |
| TEG-10A | 0.003199 | 0.009762 | 0.987038 |
| TEG-1A | 0.094384 | 0.027481 | 0.878135 |
| TEG-2A | 0.005115 | 0.006105 | 0.98878 |
| TEG-3A | 1e-05 | 0.003456 | 0.996534 |
| TEG-4A | 1e-05 | 0.048938 | 0.951052 |
| TEG-5A | 1e-05 | 0.025801 | 0.974189 |
| TEG-6A | 1e-05 | 0.035832 | 0.964158 |
| TEG-7A | 0.057492 | 0.04973 | 0.892779 |
| TEG-8A | 1e-05 | 0.030003 | 0.969987 |
| TEG-9A | 0.038014 | 1e-05 | 0.961976 |
| ZPZ-10A | 1e-05 | 1e-05 | 0.99998 |
| ZPZ-1A | 1e-05 | 0.014844 | 0.985146 |
| ZPZ-2A | 1e-05 | 0.002254 | 0.997736 |
| ZPZ-3A | 1e-05 | 1e-05 | 0.99998 |
| ZPZ-4A | 1e-05 | 1e-05 | 0.99998 |
| ZPZ-5A | 1e-05 | 1e-05 | 0.99998 |
| ZPZ-6A | 1e-05 | 0.004106 | 0.995884 |
| ZPZ-7A | 1e-05 | 1e-05 | 0.99998 |
| ZPZ-8A | 1e-05 | 0.014519 | 0.985471 |
| ZPZ-9A | 1e-05 | 1e-05 | 0.99998 |
| YLJ1-1A | 1e-05 | 1e-05 | 0.99998 |
| YLJ1-2A | 1e-05 | 1e-05 | 0.99998 |
| YLJ1-3A | 1e-05 | 1e-05 | 0.99998 |
| YLJ1-4A | 1e-05 | 1e-05 | 0.99998 |
| YLJ1-5A | 1e-05 | 1e-05 | 0.99998 |
| YLJ1-6A | 1e-05 | 1e-05 | 0.99998 |
| YLJ1-7A | 1e-05 | 1e-05 | 0.99998 |
| YLJ1-8A | 1e-05 | 1e-05 | 0.99998 |
| YLJ2-1A | 1e-05 | 1e-05 | 0.99998 |
| YLJ2-2A | 1.7e-05 | 1e-05 | 0.999973 |
| YLJ2-3A | 1e-05 | 1e-05 | 0.99998 |
| YLJ2-4A | 1e-05 | 1e-05 | 0.99998 |
| YLJ2-5A | 1e-05 | 0.018862 | 0.981128 |
| YLJ2-6B | 1e-05 | 1e-05 | 0.99998 |
| YLJ2-7A | 1.2e-05 | 1e-05 | 0.999978 |
| YLJ2-8A | 1e-05 | 1e-05 | 0.99998 |
| LM-10A | 1e-05 | 1e-05 | 0.99998 |
| LM-1A | 1.3e-05 | 1e-05 | 0.999977 |
| LM-2A | 1e-05 | 1e-05 | 0.99998 |
| LM-3A | 1e-05 | 1e-05 | 0.99998 |
| LM-4A | 1e-05 | 1e-05 | 0.99998 |
| LM-5A | 1e-05 | 1e-05 | 0.99998 |
| LM-6A | 1e-05 | 1e-05 | 0.99998 |
| LM-7A | 1e-05 | 1e-05 | 0.99998 |
| LM-8A | 1e-05 | 1e-05 | 0.99998 |
| LM-9A | 1e-05 | 1e-05 | 0.99998 |
| LM-11A | 1e-05 | 1e-05 | 0.99998 |
| LM-12A | 1e-05 | 1e-05 | 0.99998 |
| LM-13A | 1e-05 | 1e-05 | 0.99998 |
| LM-14A | 1e-05 | 1e-05 | 0.99998 |
| LM-15A | 1e-05 | 1e-05 | 0.99998 |
| BYP1-1A | 1e-05 | 1e-05 | 0.99998 |
| BYP1-2A | 0.016873 | 1e-05 | 0.983117 |
| BYP1-3A | 1e-05 | 1e-05 | 0.99998 |
| BYP1-4A | 1.7e-05 | 1e-05 | 0.999973 |
| BYP1-5A | 1e-05 | 1e-05 | 0.99998 |
| BYP1-6A | 1e-05 | 1e-05 | 0.99998 |
| BYP1-7A | 1e-05 | 1e-05 | 0.99998 |
| BYP1-8A | 1e-05 | 1e-05 | 0.99998 |
| BYP1-9A | 1e-05 | 1e-05 | 0.99998 |
| BYP1-10A | 0.058164 | 1e-05 | 0.941826 |
| BYP2-1A | 1e-05 | 1e-05 | 0.99998 |
| BYP2-2A | 1e-05 | 1e-05 | 0.99998 |
| BYP2-3A | 0.002488 | 1e-05 | 0.997502 |
| BYP2-4A | 1e-05 | 1e-05 | 0.99998 |
| BYP2-5A | 1e-05 | 1e-05 | 0.99998 |
| BYP2-6A | 1.1e-05 | 1e-05 | 0.999979 |
| BYP2-7A | 1e-05 | 1e-05 | 0.99998 |
| BYP2-8A | 1e-05 | 1e-05 | 0.99998 |
| BYP3-1A | 1.2e-05 | 1e-05 | 0.999978 |
| BYP3-2A | 1e-05 | 1e-05 | 0.99998 |
| BYP3-3A | 1e-05 | 1e-05 | 0.99998 |
| BYP3-4A | 0.044721 | 1e-05 | 0.955269 |
| BYP3-5A | 0.063872 | 1e-05 | 0.936118 |
| BYP3-6A | 1e-05 | 1e-05 | 0.99998 |
| BYP3-7A | 0.028302 | 1e-05 | 0.971688 |
| BYP3-8A | 1e-05 | 1e-05 | 0.99998 |
| KBG-1A | 0.219232 | 1e-05 | 0.780758 |
| KBG-2A | 0.213147 | 1e-05 | 0.786843 |
| KBG-3A | 0.171721 | 1e-05 | 0.828269 |
| KBG-4A | 0.204146 | 1e-05 | 0.795844 |
| KBG-5A | 0.264651 | 1e-05 | 0.735339 |
| QJG-1A | 0.294423 | 1e-05 | 0.705567 |
| QJG-2A | 0.248444 | 1e-05 | 0.751546 |
| QJG-3A | 0.269611 | 1e-05 | 0.730379 |
| QJG-4A | 0.258331 | 1e-05 | 0.741659 |
| QJG-5A | 0.334415 | 0.02246 | 0.643126 |
| LTG-1A | 0.121736 | 1e-05 | 0.878254 |
| LTG-2A | 0.214665 | 0.038829 | 0.746507 |
| LTG-3A | 0.243885 | 1e-05 | 0.756105 |
| LTG-4A | 0.280535 | 1e-05 | 0.719455 |
| LTG-5A | 0.165136 | 0.030289 | 0.804575 |
| BW-10B | 0.14115 | 1e-05 | 0.85884 |
| BW-1B | 0.180583 | 1e-05 | 0.819407 |
| BW-2B | 0.173091 | 1e-05 | 0.826899 |
| BW-3B | 0.203161 | 1e-05 | 0.796829 |
| BW-4B | 0.181516 | 1e-05 | 0.818474 |
| BW-5B | 0.168584 | 1e-05 | 0.831406 |
| BW-6B | 0.169383 | 1e-05 | 0.830607 |
| BW-7A | 0.130984 | 1e-05 | 0.869006 |
| BW-8B | 0.17495 | 1e-05 | 0.82504 |
| BW-9A | 0.188006 | 1e-05 | 0.811984 |
| GA-1A | 0.264944 | 1e-05 | 0.735046 |
| GA-2A | 0.177888 | 1e-05 | 0.822102 |
| GA-3A | 0.120942 | 0.006471 | 0.872587 |
| MR-1A | 0.219604 | 1e-05 | 0.780386 |
| MR-2A | 0.228963 | 0.021732 | 0.749306 |
| MR-3A | 0.228723 | 0.02342 | 0.747857 |
| MR-4A | 0.276481 | 1e-05 | 0.723509 |
| MR-5A | 0.163596 | 0.041197 | 0.795207 |
| MDG-6A | 0.51381 | 1e-05 | 0.48618 |
| MDG-1A | 0.459007 | 1e-05 | 0.540983 |
| MDG-2A | 0.458761 | 1e-05 | 0.541229 |
| MDG-3A | 0.473554 | 1e-05 | 0.526436 |
| MDG-4A | 0.474145 | 0.023953 | 0.501901 |
| MDG-5A | 0.429895 | 1e-05 | 0.570095 |
| AJ-10A | 0.864232 | 1e-05 | 0.135758 |
| AJ-12A | 0.947036 | 1e-05 | 0.052954 |
| AJ-13A | 0.892579 | 1e-05 | 0.107411 |
| AJ-14A | 0.860101 | 1e-05 | 0.139889 |
| AJ-1A | 0.843725 | 1e-05 | 0.156265 |
| AJ-3A | 0.780835 | 1e-05 | 0.219155 |
| AJ-4A | 0.936786 | 0.01466 | 0.048555 |
| AJ-5A | 0.86491 | 1e-05 | 0.13508 |
| AJ-7A | 0.776758 | 1e-05 | 0.223232 |
| AJ-9A | 0.99998 | 1e-05 | 1e-05 |
| XGG-1A | 0.99998 | 1e-05 | 1e-05 |
| XGG-2A | 0.99998 | 1e-05 | 1e-05 |
| XGG-3A | 0.99998 | 1e-05 | 1e-05 |
| XGG-4A | 0.99998 | 1e-05 | 1e-05 |
| XGG-5A | 0.99998 | 1e-05 | 1e-05 |
| PSR-01A | 0.99998 | 1e-05 | 1e-05 |
| PSR-04A | 0.99998 | 1e-05 | 1e-05 |
| PSR-10A | 0.99998 | 1e-05 | 1e-05 |
| PSR-13A | 0.995004 | 1e-05 | 0.004986 |
| LR-10A | 0.99998 | 1e-05 | 1e-05 |
| LR-1A | 0.99998 | 1e-05 | 1e-05 |
| LR-2A | 0.99998 | 1e-05 | 1e-05 |
| LR-3A | 0.99998 | 1e-05 | 1e-05 |
| LR-4A | 0.99998 | 1e-05 | 1e-05 |
| LR-5A | 0.99998 | 1e-05 | 1e-05 |
| LR-6A | 0.99998 | 1e-05 | 1e-05 |
| LR-7A | 0.99998 | 1e-05 | 1e-05 |
| LR-8A | 0.99998 | 1e-05 | 1e-05 |
| LR-9A | 0.99998 | 1e-05 | 1e-05 |
| BYR-10A | 0.99998 | 1e-05 | 1e-05 |
| BYR-1A | 0.99998 | 1e-05 | 1e-05 |
| BYR-2A | 0.99998 | 1e-05 | 1e-05 |
| BYR-3A | 0.99998 | 1e-05 | 1e-05 |
| BYR-4A | 0.99998 | 1e-05 | 1e-05 |
| BYR-5A | 0.99998 | 1e-05 | 1e-05 |
| BYR-6A | 0.99998 | 1e-05 | 1e-05 |
| BYR-7A | 0.99998 | 1e-05 | 1e-05 |

K = 4

|  |  |  |  |  |
| --- | --- | --- | --- | --- |
| CDG-1A | 1e-05 | 0.99997 | 1e-05 | 1e-05 |
| CDG-2A | 1.1e-05 | 0.999969 | 1e-05 | 1e-05 |
| CDG-3A | 1e-05 | 0.999969 | 1.1e-05 | 1e-05 |
| CDG-4A | 1e-05 | 0.99997 | 1e-05 | 1e-05 |
| CDG-5A | 1e-05 | 0.999969 | 1.1e-05 | 1e-05 |
| CDG-6A | 1e-05 | 0.99997 | 1e-05 | 1e-05 |
| CDG-7A | 1e-05 | 0.99997 | 1e-05 | 1e-05 |
| CDG-8A | 1e-05 | 0.99997 | 1e-05 | 1e-05 |
| CDG-9A | 1e-05 | 0.946777 | 0.053203 | 1e-05 |
| CDG-10A | 1e-05 | 0.99997 | 1e-05 | 1e-05 |
| TKX-1A | 1e-05 | 0.999967 | 1.3e-05 | 1e-05 |
| TKX-2A | 1e-05 | 0.99997 | 1e-05 | 1e-05 |
| TKX-3A | 1e-05 | 0.99997 | 1e-05 | 1e-05 |
| TKX-4A | 1e-05 | 0.99997 | 1e-05 | 1e-05 |
| TKX-5A | 1e-05 | 0.99997 | 1e-05 | 1e-05 |
| TKX-6A | 1e-05 | 0.999969 | 1.1e-05 | 1e-05 |
| TKX-7A | 1e-05 | 0.99997 | 1e-05 | 1e-05 |
| TKX-8A | 1e-05 | 0.99997 | 1e-05 | 1e-05 |
| TKX-9A | 1.5e-05 | 0.999964 | 1.2e-05 | 1e-05 |
| GJ-1A | 1e-05 | 1e-05 | 0.99997 | 1e-05 |
| GJ-2A | 1e-05 | 1e-05 | 0.99997 | 1e-05 |
| GJ-3A | 1.1e-05 | 1e-05 | 0.999969 | 1e-05 |
| GJ-4A | 1e-05 | 1e-05 | 0.99997 | 1e-05 |
| GJ-5A | 1e-05 | 1e-05 | 0.99997 | 1e-05 |
| GJ-6A | 1e-05 | 1.4e-05 | 0.999966 | 1e-05 |
| GJ-7A | 1e-05 | 1e-05 | 0.99997 | 1e-05 |
| GJ-8A | 1.3e-05 | 1e-05 | 0.999967 | 1e-05 |
| GJ-9A | 1e-05 | 1.4e-05 | 0.999966 | 1e-05 |
| TZC-1A | 1e-05 | 1e-05 | 0.99997 | 1e-05 |
| TZC-2A | 1e-05 | 1e-05 | 0.99997 | 1e-05 |
| TZC-3A | 1e-05 | 1e-05 | 0.99997 | 1e-05 |
| TZC-4A | 1e-05 | 1e-05 | 0.99997 | 1e-05 |
| TZC-01A | 1e-05 | 0.039853 | 0.960127 | 1e-05 |
| TZC-04A | 1e-05 | 1e-05 | 0.99997 | 1e-05 |
| TZC-07A | 1e-05 | 0.035854 | 0.964126 | 1e-05 |
| TZC-11A | 1e-05 | 1.3e-05 | 0.999967 | 1e-05 |
| TZC-13A | 1e-05 | 1e-05 | 0.99997 | 1e-05 |
| DBG-10A | 1e-05 | 1e-05 | 0.99997 | 1e-05 |
| DBG-1A | 0.053886 | 1e-05 | 0.946094 | 1e-05 |
| DBG-2A | 0.001818 | 0.018346 | 0.979826 | 1e-05 |
| DBG-3A | 1e-05 | 1e-05 | 0.99997 | 1e-05 |
| DBG-4A | 1.2e-05 | 1e-05 | 0.999968 | 1e-05 |
| DBG-5A | 1e-05 | 1e-05 | 0.99997 | 1e-05 |
| DBG-6A | 1e-05 | 0.060951 | 0.939029 | 1e-05 |
| DBG-7A | 1e-05 | 0.014161 | 0.985819 | 1e-05 |
| DBG-8A | 1e-05 | 1e-05 | 0.99997 | 1e-05 |
| DBG-9A | 1e-05 | 0.035446 | 0.936709 | 0.027836 |
| TEG-10A | 1e-05 | 1e-05 | 0.99997 | 1e-05 |
| TEG-1A | 0.074276 | 1e-05 | 0.891507 | 0.034207 |
| TEG-2A | 1e-05 | 1e-05 | 0.930943 | 0.069037 |
| TEG-3A | 1e-05 | 1.6e-05 | 0.999964 | 1e-05 |
| TEG-4A | 1e-05 | 1e-05 | 0.962657 | 0.037323 |
| TEG-5A | 1e-05 | 1e-05 | 0.99997 | 1e-05 |
| TEG-6A | 1e-05 | 1e-05 | 0.884527 | 0.115453 |
| TEG-7A | 0.05084 | 1e-05 | 0.821625 | 0.127526 |
| TEG-8A | 1e-05 | 1e-05 | 0.943352 | 0.056628 |
| TEG-9A | 0.003027 | 1e-05 | 0.996953 | 1e-05 |
| ZPZ-10A | 1e-05 | 1e-05 | 0.99997 | 1e-05 |
| ZPZ-1A | 1e-05 | 1.1e-05 | 0.999969 | 1e-05 |
| ZPZ-2A | 1e-05 | 1e-05 | 0.99997 | 1e-05 |
| ZPZ-3A | 1.9e-05 | 1e-05 | 0.974446 | 0.025525 |
| ZPZ-4A | 1e-05 | 1e-05 | 0.99997 | 1e-05 |
| ZPZ-5A | 1e-05 | 1e-05 | 0.99997 | 1e-05 |
| ZPZ-6A | 1e-05 | 1e-05 | 0.762322 | 0.237658 |
| ZPZ-7A | 1e-05 | 1e-05 | 0.94908 | 0.0509 |
| ZPZ-8A | 1e-05 | 1e-05 | 0.99997 | 1e-05 |
| ZPZ-9A | 1e-05 | 1e-05 | 0.99997 | 1e-05 |
| YLJ1-1A | 1e-05 | 1e-05 | 0.99997 | 1e-05 |
| YLJ1-2A | 1e-05 | 1e-05 | 0.99997 | 1e-05 |
| YLJ1-3A | 1e-05 | 1e-05 | 0.99997 | 1e-05 |
| YLJ1-4A | 1e-05 | 1e-05 | 0.99997 | 1e-05 |
| YLJ1-5A | 1e-05 | 1.1e-05 | 0.999969 | 1e-05 |
| YLJ1-6A | 1e-05 | 1e-05 | 0.99997 | 1e-05 |
| YLJ1-7A | 1e-05 | 1e-05 | 0.99997 | 1e-05 |
| YLJ1-8A | 1e-05 | 1e-05 | 0.99997 | 1e-05 |
| YLJ2-1A | 1.1e-05 | 1e-05 | 0.999969 | 1e-05 |
| YLJ2-2A | 1e-05 | 1e-05 | 0.99997 | 1e-05 |
| YLJ2-3A | 1e-05 | 1e-05 | 0.99997 | 1e-05 |
| YLJ2-4A | 1e-05 | 1e-05 | 0.864776 | 0.135204 |
| YLJ2-5A | 1e-05 | 1e-05 | 0.99997 | 1e-05 |
| YLJ2-6B | 1e-05 | 1e-05 | 0.99997 | 1e-05 |
| YLJ2-7A | 1.7e-05 | 1e-05 | 0.999963 | 1e-05 |
| YLJ2-8A | 1e-05 | 1e-05 | 0.875569 | 0.124411 |
| LM-10A | 1e-05 | 1e-05 | 1e-05 | 0.99997 |
| LM-1A | 1e-05 | 1e-05 | 1e-05 | 0.99997 |
| LM-2A | 1e-05 | 1e-05 | 1e-05 | 0.99997 |
| LM-3A | 1.1e-05 | 1.1e-05 | 1e-05 | 0.999969 |
| LM-4A | 1e-05 | 1e-05 | 1e-05 | 0.99997 |
| LM-5A | 1e-05 | 1e-05 | 1e-05 | 0.99997 |
| LM-6A | 1e-05 | 1e-05 | 1e-05 | 0.99997 |
| LM-7A | 1e-05 | 1e-05 | 1e-05 | 0.99997 |
| LM-8A | 1e-05 | 1e-05 | 1e-05 | 0.99997 |
| LM-9A | 1e-05 | 1e-05 | 1e-05 | 0.99997 |
| LM-11A | 1e-05 | 1e-05 | 1e-05 | 0.99997 |
| LM-12A | 1e-05 | 1e-05 | 1e-05 | 0.99997 |
| LM-13A | 1e-05 | 1e-05 | 1e-05 | 0.99997 |
| LM-14A | 1e-05 | 1e-05 | 1e-05 | 0.99997 |
| LM-15A | 1e-05 | 1e-05 | 1e-05 | 0.99997 |
| BYP1-1A | 1e-05 | 1e-05 | 0.075134 | 0.924846 |
| BYP1-2A | 1.3e-05 | 1e-05 | 0.046688 | 0.953289 |
| BYP1-3A | 1e-05 | 1e-05 | 0.151045 | 0.848935 |
| BYP1-4A | 1e-05 | 1e-05 | 0.107204 | 0.892776 |
| BYP1-5A | 1e-05 | 1e-05 | 0.123288 | 0.876692 |
| BYP1-6A | 1e-05 | 1e-05 | 1e-05 | 0.99997 |
| BYP1-7A | 1e-05 | 1e-05 | 1e-05 | 0.99997 |
| BYP1-8A | 1e-05 | 1e-05 | 1e-05 | 0.99997 |
| BYP1-9A | 1e-05 | 1e-05 | 1e-05 | 0.99997 |
| BYP1-10A | 0.035998 | 1e-05 | 1e-05 | 0.963982 |
| BYP2-1A | 1e-05 | 1e-05 | 1e-05 | 0.99997 |
| BYP2-2A | 1e-05 | 1e-05 | 0.041988 | 0.957992 |
| BYP2-3A | 1e-05 | 1e-05 | 1e-05 | 0.99997 |
| BYP2-4A | 1e-05 | 1e-05 | 1e-05 | 0.99997 |
| BYP2-5A | 1e-05 | 1e-05 | 0.143323 | 0.856657 |
| BYP2-6A | 1e-05 | 1e-05 | 0.047285 | 0.952695 |
| BYP2-7A | 1e-05 | 1e-05 | 0.097224 | 0.902756 |
| BYP2-8A | 1e-05 | 1e-05 | 1e-05 | 0.99997 |
| BYP3-1A | 1e-05 | 1e-05 | 0.050872 | 0.949108 |
| BYP3-2A | 1e-05 | 1e-05 | 1e-05 | 0.99997 |
| BYP3-3A | 1e-05 | 1e-05 | 0.05797 | 0.94201 |
| BYP3-4A | 1e-05 | 1e-05 | 0.040073 | 0.959907 |
| BYP3-5A | 0.020151 | 1e-05 | 0.030627 | 0.949212 |
| BYP3-6A | 1e-05 | 1e-05 | 0.097169 | 0.902811 |
| BYP3-7A | 0.000299 | 1e-05 | 1e-05 | 0.999681 |
| BYP3-8A | 1e-05 | 1e-05 | 1e-05 | 0.99997 |
| KBG-1A | 0.112917 | 1e-05 | 1e-05 | 0.887063 |
| KBG-2A | 0.119879 | 1e-05 | 0.07874 | 0.801371 |
| KBG-3A | 0.122991 | 1e-05 | 1e-05 | 0.876989 |
| KBG-4A | 0.147085 | 1e-05 | 1e-05 | 0.852895 |
| KBG-5A | 0.188966 | 0.001867 | 1e-05 | 0.809157 |
| QJG-1A | 0.192728 | 1e-05 | 1e-05 | 0.807252 |
| QJG-2A | 0.177461 | 1e-05 | 1e-05 | 0.822519 |
| QJG-3A | 0.156223 | 1e-05 | 1e-05 | 0.843757 |
| QJG-4A | 0.150984 | 1e-05 | 1e-05 | 0.848996 |
| QJG-5A | 0.253037 | 1e-05 | 1e-05 | 0.746943 |
| LTG-1A | 0.050151 | 1e-05 | 0.14701 | 0.802829 |
| LTG-2A | 0.155419 | 0.035283 | 0.027393 | 0.781906 |
| LTG-3A | 0.134881 | 1e-05 | 1e-05 | 0.865099 |
| LTG-4A | 0.190381 | 1e-05 | 1e-05 | 0.809599 |
| LTG-5A | 0.116018 | 0.036566 | 1e-05 | 0.847407 |
| BW-10B | 0.105055 | 1e-05 | 1e-05 | 0.894925 |
| BW-1B | 0.103465 | 1e-05 | 1e-05 | 0.896515 |
| BW-2B | 0.058029 | 1e-05 | 1e-05 | 0.941951 |
| BW-3B | 0.131811 | 1e-05 | 1e-05 | 0.868169 |
| BW-4B | 0.106884 | 1e-05 | 1e-05 | 0.893096 |
| BW-5B | 0.109244 | 1e-05 | 1e-05 | 0.890736 |
| BW-6B | 0.057221 | 1e-05 | 1e-05 | 0.942759 |
| BW-7A | 0.033091 | 1e-05 | 1e-05 | 0.966889 |
| BW-8B | 0.104683 | 1e-05 | 1e-05 | 0.895297 |
| BW-9A | 0.107652 | 1.5e-05 | 1e-05 | 0.892323 |
| GA-1A | 0.184414 | 1e-05 | 1e-05 | 0.815566 |
| GA-2A | 0.111314 | 1e-05 | 1e-05 | 0.888666 |
| GA-3A | 0.03517 | 1e-05 | 1e-05 | 0.96481 |
| MR-1A | 0.180824 | 1e-05 | 1e-05 | 0.819156 |
| MR-2A | 0.176992 | 0.013606 | 1e-05 | 0.809392 |
| MR-3A | 0.191422 | 0.023405 | 1e-05 | 0.785163 |
| MR-4A | 0.197893 | 1e-05 | 1e-05 | 0.802087 |
| MR-5A | 0.129215 | 0.032658 | 0.001947 | 0.83618 |
| MDG-6A | 0.409101 | 1e-05 | 0.007236 | 0.583653 |
| MDG-1A | 0.340417 | 1e-05 | 1e-05 | 0.659563 |
| MDG-2A | 0.344799 | 1e-05 | 1e-05 | 0.655181 |
| MDG-3A | 0.372305 | 1e-05 | 1e-05 | 0.627675 |
| MDG-4A | 0.383107 | 0.013584 | 1e-05 | 0.603299 |
| MDG-5A | 0.371196 | 1e-05 | 0.000685 | 0.628109 |
| AJ-10A | 0.835589 | 1e-05 | 1e-05 | 0.164391 |
| AJ-12A | 0.900099 | 1e-05 | 1e-05 | 0.099881 |
| AJ-13A | 0.863556 | 1e-05 | 1e-05 | 0.136424 |
| AJ-14A | 0.815905 | 1e-05 | 1e-05 | 0.184075 |
| AJ-1A | 0.806326 | 1e-05 | 1e-05 | 0.193654 |
| AJ-3A | 0.758543 | 1e-05 | 0.028425 | 0.213022 |
| AJ-4A | 0.906642 | 0.005087 | 1e-05 | 0.08826 |
| AJ-5A | 0.828711 | 1e-05 | 1e-05 | 0.171269 |
| AJ-7A | 0.750144 | 1e-05 | 0.09187 | 0.157976 |
| AJ-9A | 0.961448 | 1.1e-05 | 1e-05 | 0.038532 |
| XGG-1A | 0.999968 | 1.2e-05 | 1e-05 | 1e-05 |
| XGG-2A | 0.99997 | 1e-05 | 1e-05 | 1e-05 |
| XGG-3A | 0.999961 | 1e-05 | 1.9e-05 | 1e-05 |
| XGG-4A | 0.999969 | 1e-05 | 1.1e-05 | 1e-05 |
| XGG-5A | 0.99997 | 1e-05 | 1e-05 | 1e-05 |
| PSR-01A | 0.99997 | 1e-05 | 1e-05 | 1e-05 |
| PSR-04A | 0.99997 | 1e-05 | 1e-05 | 1e-05 |
| PSR-10A | 0.999967 | 1.2e-05 | 1e-05 | 1e-05 |
| PSR-13A | 0.967614 | 1e-05 | 1e-05 | 0.032366 |
| LR-10A | 0.99997 | 1e-05 | 1e-05 | 1e-05 |
| LR-1A | 0.99997 | 1e-05 | 1e-05 | 1e-05 |
| LR-2A | 0.99997 | 1e-05 | 1e-05 | 1e-05 |
| LR-3A | 0.99997 | 1e-05 | 1e-05 | 1e-05 |
| LR-4A | 0.999964 | 1e-05 | 1.6e-05 | 1e-05 |
| LR-5A | 0.99997 | 1e-05 | 1e-05 | 1e-05 |
| LR-6A | 0.987733 | 1e-05 | 0.012247 | 1e-05 |
| LR-7A | 0.99997 | 1e-05 | 1e-05 | 1e-05 |
| LR-8A | 0.99997 | 1e-05 | 1e-05 | 1e-05 |
| LR-9A | 0.99997 | 1e-05 | 1e-05 | 1e-05 |
| BYR-10A | 0.99997 | 1e-05 | 1e-05 | 1e-05 |
| BYR-1A | 0.99997 | 1e-05 | 1e-05 | 1e-05 |
| BYR-2A | 0.99997 | 1e-05 | 1e-05 | 1e-05 |
| BYR-3A | 0.99997 | 1e-05 | 1e-05 | 1e-05 |
| BYR-4A | 0.999969 | 1.1e-05 | 1e-05 | 1e-05 |
| BYR-5A | 0.999969 | 1e-05 | 1.1e-05 | 1e-05 |
| BYR-6A | 0.99997 | 1e-05 | 1e-05 | 1e-05 |
| BYR-7A | 0.99997 | 1e-05 | 1e-05 | 1e-05 |

K = 5

|  |  |  |  |  |  |
| --- | --- | --- | --- | --- | --- |
| CDG-1A | 0.99996 | 1e-05 | 1e-05 | 1e-05 | 1e-05 |
| CDG-2A | 0.999955 | 1.5e-05 | 1e-05 | 1e-05 | 1e-05 |
| CDG-3A | 0.99996 | 1e-05 | 1e-05 | 1e-05 | 1e-05 |
| CDG-4A | 0.99996 | 1e-05 | 1e-05 | 1e-05 | 1e-05 |
| CDG-5A | 0.99996 | 1e-05 | 1e-05 | 1e-05 | 1e-05 |
| CDG-6A | 0.99996 | 1e-05 | 1e-05 | 1e-05 | 1e-05 |
| CDG-7A | 0.99996 | 1e-05 | 1e-05 | 1e-05 | 1e-05 |
| CDG-8A | 0.99996 | 1e-05 | 1e-05 | 1e-05 | 1e-05 |
| CDG-9A | 0.913461 | 0.086509 | 1e-05 | 1e-05 | 1e-05 |
| CDG-10A | 0.999955 | 1.5e-05 | 1e-05 | 1e-05 | 1e-05 |
| TKX-1A | 0.99996 | 1e-05 | 1e-05 | 1e-05 | 1e-05 |
| TKX-2A | 0.999959 | 1e-05 | 1.1e-05 | 1e-05 | 1e-05 |
| TKX-3A | 0.99996 | 1e-05 | 1e-05 | 1e-05 | 1e-05 |
| TKX-4A | 0.99996 | 1e-05 | 1e-05 | 1e-05 | 1e-05 |
| TKX-5A | 0.99996 | 1e-05 | 1e-05 | 1e-05 | 1e-05 |
| TKX-6A | 0.99996 | 1e-05 | 1e-05 | 1e-05 | 1e-05 |
| TKX-7A | 0.99996 | 1e-05 | 1e-05 | 1e-05 | 1e-05 |
| TKX-8A | 0.99996 | 1e-05 | 1e-05 | 1e-05 | 1e-05 |
| TKX-9A | 0.99996 | 1e-05 | 1e-05 | 1e-05 | 1e-05 |
| GJ-1A | 1e-05 | 0.99996 | 1e-05 | 1e-05 | 1e-05 |
| GJ-2A | 1e-05 | 0.99996 | 1e-05 | 1e-05 | 1e-05 |
| GJ-3A | 1e-05 | 0.99996 | 1e-05 | 1e-05 | 1e-05 |
| GJ-4A | 1.2e-05 | 0.999956 | 1e-05 | 1.2e-05 | 1e-05 |
| GJ-5A | 1e-05 | 0.999957 | 1e-05 | 1.3e-05 | 1e-05 |
| GJ-6A | 1.8e-05 | 0.999952 | 1e-05 | 1e-05 | 1e-05 |
| GJ-7A | 1.5e-05 | 0.999955 | 1e-05 | 1e-05 | 1e-05 |
| GJ-8A | 1e-05 | 0.99996 | 1e-05 | 1e-05 | 1e-05 |
| GJ-9A | 1e-05 | 0.99996 | 1e-05 | 1e-05 | 1e-05 |
| TZC-1A | 1e-05 | 0.99996 | 1e-05 | 1e-05 | 1e-05 |
| TZC-2A | 1e-05 | 0.99996 | 1e-05 | 1e-05 | 1e-05 |
| TZC-3A | 1e-05 | 0.890689 | 1e-05 | 0.045438 | 0.063853 |
| TZC-4A | 1e-05 | 0.99996 | 1e-05 | 1e-05 | 1e-05 |
| TZC-01A | 1e-05 | 0.99996 | 1e-05 | 1e-05 | 1e-05 |
| TZC-04A | 1e-05 | 0.99996 | 1e-05 | 1e-05 | 1e-05 |
| TZC-07A | 1e-05 | 0.99996 | 1e-05 | 1e-05 | 1e-05 |
| TZC-11A | 1e-05 | 0.99996 | 1e-05 | 1e-05 | 1e-05 |
| TZC-13A | 1e-05 | 0.99996 | 1e-05 | 1e-05 | 1e-05 |
| DBG-10A | 1e-05 | 0.99996 | 1e-05 | 1e-05 | 1e-05 |
| DBG-1A | 1e-05 | 0.840558 | 0.031804 | 1e-05 | 0.127617 |
| DBG-2A | 1e-05 | 0.99996 | 1e-05 | 1e-05 | 1e-05 |
| DBG-3A | 1.8e-05 | 0.999952 | 1e-05 | 1e-05 | 1e-05 |
| DBG-4A | 1e-05 | 0.99996 | 1e-05 | 1e-05 | 1e-05 |
| DBG-5A | 1.2e-05 | 0.877782 | 1e-05 | 1e-05 | 0.122187 |
| DBG-6A | 1e-05 | 0.999955 | 1.5e-05 | 1e-05 | 1e-05 |
| DBG-7A | 1e-05 | 0.99996 | 1e-05 | 1e-05 | 1e-05 |
| DBG-8A | 1e-05 | 0.99996 | 1e-05 | 1e-05 | 1e-05 |
| DBG-9A | 1e-05 | 0.948107 | 1e-05 | 1e-05 | 0.051863 |
| TEG-10A | 1e-05 | 0.269817 | 1e-05 | 1e-05 | 0.730153 |
| TEG-1A | 1e-05 | 0.356464 | 0.078401 | 1e-05 | 0.565115 |
| TEG-2A | 1e-05 | 0.24621 | 0.012845 | 1e-05 | 0.740925 |
| TEG-3A | 1e-05 | 0.36734 | 1e-05 | 1e-05 | 0.63263 |
| TEG-4A | 1e-05 | 0.416906 | 1e-05 | 1e-05 | 0.583064 |
| TEG-5A | 1e-05 | 0.393226 | 1e-05 | 1e-05 | 0.606744 |
| TEG-6A | 1e-05 | 0.297295 | 1e-05 | 0.067569 | 0.635116 |
| TEG-7A | 1e-05 | 0.2756 | 0.057141 | 0.063122 | 0.604126 |
| TEG-8A | 1e-05 | 0.351444 | 1e-05 | 1e-05 | 0.648526 |
| TEG-9A | 1e-05 | 0.328178 | 0.012496 | 1e-05 | 0.659306 |
| ZPZ-10A | 1e-05 | 1e-05 | 1e-05 | 1e-05 | 0.99996 |
| ZPZ-1A | 1.1e-05 | 1.2e-05 | 1e-05 | 1e-05 | 0.999957 |
| ZPZ-2A | 1e-05 | 1e-05 | 1e-05 | 1e-05 | 0.99996 |
| ZPZ-3A | 1e-05 | 1e-05 | 1e-05 | 1e-05 | 0.99996 |
| ZPZ-4A | 1e-05 | 1.2e-05 | 1e-05 | 1e-05 | 0.999958 |
| ZPZ-5A | 1e-05 | 1e-05 | 1e-05 | 1e-05 | 0.99996 |
| ZPZ-6A | 1e-05 | 0.072814 | 0.006844 | 0.17822 | 0.742112 |
| ZPZ-7A | 1e-05 | 0.136316 | 1e-05 | 1e-05 | 0.863654 |
| ZPZ-8A | 1e-05 | 1e-05 | 1e-05 | 1e-05 | 0.99996 |
| ZPZ-9A | 1e-05 | 1e-05 | 1e-05 | 1e-05 | 0.99996 |
| YLJ1-1A | 1e-05 | 1e-05 | 1e-05 | 1e-05 | 0.99996 |
| YLJ1-2A | 1e-05 | 1e-05 | 1e-05 | 1e-05 | 0.99996 |
| YLJ1-3A | 1e-05 | 1e-05 | 1e-05 | 1e-05 | 0.99996 |
| YLJ1-4A | 1e-05 | 1e-05 | 1e-05 | 1e-05 | 0.99996 |
| YLJ1-5A | 1e-05 | 1e-05 | 1e-05 | 1e-05 | 0.99996 |
| YLJ1-6A | 1e-05 | 1e-05 | 1e-05 | 1e-05 | 0.99996 |
| YLJ1-7A | 1e-05 | 1e-05 | 1e-05 | 1e-05 | 0.99996 |
| YLJ1-8A | 1e-05 | 1e-05 | 1e-05 | 1e-05 | 0.99996 |
| YLJ2-1A | 1e-05 | 1e-05 | 1e-05 | 1e-05 | 0.99996 |
| YLJ2-2A | 1e-05 | 1e-05 | 1e-05 | 1e-05 | 0.99996 |
| YLJ2-3A | 1e-05 | 1e-05 | 1e-05 | 1e-05 | 0.99996 |
| YLJ2-4A | 1e-05 | 1e-05 | 1e-05 | 1e-05 | 0.99996 |
| YLJ2-5A | 1e-05 | 1e-05 | 1e-05 | 1e-05 | 0.99996 |
| YLJ2-6B | 1e-05 | 1e-05 | 1e-05 | 1e-05 | 0.99996 |
| YLJ2-7A | 1e-05 | 1e-05 | 1e-05 | 1e-05 | 0.99996 |
| YLJ2-8A | 1e-05 | 1e-05 | 1e-05 | 1e-05 | 0.99996 |
| LM-10A | 1e-05 | 1e-05 | 1e-05 | 0.99996 | 1e-05 |
| LM-1A | 1e-05 | 1e-05 | 1e-05 | 0.99996 | 1e-05 |
| LM-2A | 1e-05 | 1e-05 | 1e-05 | 0.99996 | 1e-05 |
| LM-3A | 1.1e-05 | 1e-05 | 1e-05 | 0.999959 | 1e-05 |
| LM-4A | 1e-05 | 1e-05 | 1e-05 | 0.99996 | 1e-05 |
| LM-5A | 1e-05 | 1e-05 | 1e-05 | 0.99996 | 1e-05 |
| LM-6A | 1e-05 | 1e-05 | 1e-05 | 0.99996 | 1e-05 |
| LM-7A | 1e-05 | 1e-05 | 1e-05 | 0.99996 | 1e-05 |
| LM-8A | 1.2e-05 | 1e-05 | 1e-05 | 0.999958 | 1e-05 |
| LM-9A | 1e-05 | 1e-05 | 1e-05 | 0.99996 | 1e-05 |
| LM-11A | 1.2e-05 | 1e-05 | 1e-05 | 0.999958 | 1e-05 |
| LM-12A | 1e-05 | 1e-05 | 1e-05 | 0.99996 | 1e-05 |
| LM-13A | 1e-05 | 1e-05 | 1e-05 | 0.99996 | 1e-05 |
| LM-14A | 1e-05 | 1e-05 | 1e-05 | 0.99996 | 1e-05 |
| LM-15A | 1e-05 | 1e-05 | 1e-05 | 0.99996 | 1e-05 |
| BYP1-1A | 1e-05 | 1e-05 | 1e-05 | 0.705864 | 0.294106 |
| BYP1-2A | 1e-05 | 1e-05 | 1e-05 | 0.806241 | 0.193729 |
| BYP1-3A | 1e-05 | 1e-05 | 1e-05 | 0.712381 | 0.287589 |
| BYP1-4A | 1e-05 | 1e-05 | 1e-05 | 0.730367 | 0.269603 |
| BYP1-5A | 1e-05 | 1e-05 | 1e-05 | 0.656242 | 0.343728 |
| BYP1-6A | 1e-05 | 1e-05 | 1e-05 | 0.9183 | 0.08167 |
| BYP1-7A | 1e-05 | 1e-05 | 1e-05 | 0.867344 | 0.132626 |
| BYP1-8A | 1e-05 | 1e-05 | 1e-05 | 0.915386 | 0.084584 |
| BYP1-9A | 1e-05 | 1e-05 | 1e-05 | 0.866215 | 0.133755 |
| BYP1-10A | 1e-05 | 1e-05 | 0.029697 | 0.860011 | 0.110272 |
| BYP2-1A | 1e-05 | 1e-05 | 1e-05 | 0.739738 | 0.260231 |
| BYP2-2A | 1e-05 | 0.012159 | 1e-05 | 0.718733 | 0.269088 |
| BYP2-3A | 1e-05 | 1e-05 | 1e-05 | 0.90168 | 0.09829 |
| BYP2-4A | 1e-05 | 1e-05 | 1e-05 | 0.687289 | 0.312681 |
| BYP2-5A | 1e-05 | 0.034979 | 1e-05 | 0.749372 | 0.215629 |
| BYP2-6A | 1e-05 | 1e-05 | 1e-05 | 0.772662 | 0.227308 |
| BYP2-7A | 1e-05 | 1e-05 | 1e-05 | 0.808376 | 0.191594 |
| BYP2-8A | 1e-05 | 1e-05 | 1e-05 | 0.735816 | 0.264154 |
| BYP3-1A | 1e-05 | 0.005046 | 1e-05 | 0.824357 | 0.170577 |
| BYP3-2A | 1e-05 | 1e-05 | 1e-05 | 0.825951 | 0.174019 |
| BYP3-3A | 1e-05 | 0.003718 | 1e-05 | 0.822836 | 0.173426 |
| BYP3-4A | 1e-05 | 1e-05 | 0.003752 | 0.785482 | 0.210746 |
| BYP3-5A | 1e-05 | 1e-05 | 0.025309 | 0.788473 | 0.186198 |
| BYP3-6A | 1e-05 | 1e-05 | 1e-05 | 0.740402 | 0.259568 |
| BYP3-7A | 1e-05 | 1e-05 | 0.003552 | 0.835259 | 0.161169 |
| BYP3-8A | 1e-05 | 1e-05 | 1e-05 | 0.793603 | 0.206367 |
| KBG-1A | 1e-05 | 1e-05 | 0.105605 | 0.894365 | 1e-05 |
| KBG-2A | 1e-05 | 1e-05 | 0.108766 | 0.804868 | 0.086347 |
| KBG-3A | 0.005465 | 1e-05 | 0.117161 | 0.877354 | 1e-05 |
| KBG-4A | 1e-05 | 1e-05 | 0.14072 | 0.85925 | 1e-05 |
| KBG-5A | 8.7e-05 | 1e-05 | 0.180895 | 0.818998 | 1e-05 |
| QJG-1A | 1.9e-05 | 1e-05 | 0.179665 | 0.820297 | 1e-05 |
| QJG-2A | 1e-05 | 0.003237 | 0.164535 | 0.788989 | 0.04323 |
| QJG-3A | 1e-05 | 1e-05 | 0.138836 | 0.861134 | 1e-05 |
| QJG-4A | 1e-05 | 1e-05 | 0.137003 | 0.862967 | 1e-05 |
| QJG-5A | 1e-05 | 0.042893 | 0.227301 | 0.729786 | 1e-05 |
| LTG-1A | 1e-05 | 0.195669 | 0.033442 | 0.770869 | 1e-05 |
| LTG-2A | 0.030842 | 0.074695 | 0.137304 | 0.73866 | 0.018499 |
| LTG-3A | 1e-05 | 1e-05 | 0.123747 | 0.876223 | 1e-05 |
| LTG-4A | 1e-05 | 0.023838 | 0.1731 | 0.803042 | 1e-05 |
| LTG-5A | 0.044312 | 0.036414 | 0.094163 | 0.825101 | 1e-05 |
| BW-10B | 1e-05 | 0.011129 | 0.091509 | 0.897343 | 1e-05 |
| BW-1B | 1e-05 | 0.002698 | 0.081945 | 0.915337 | 1e-05 |
| BW-2B | 1e-05 | 1e-05 | 0.028939 | 0.971031 | 1e-05 |
| BW-3B | 1.7e-05 | 1e-05 | 0.115109 | 0.884854 | 1e-05 |
| BW-4B | 1e-05 | 1e-05 | 0.090865 | 0.909105 | 1e-05 |
| BW-5B | 1e-05 | 1e-05 | 0.096881 | 0.903089 | 1e-05 |
| BW-6B | 1e-05 | 1e-05 | 0.028469 | 0.971501 | 1e-05 |
| BW-7A | 1e-05 | 1e-05 | 0.003361 | 0.996609 | 1e-05 |
| BW-8B | 1e-05 | 0.039407 | 0.086611 | 0.873961 | 1e-05 |
| BW-9A | 1e-05 | 1e-05 | 0.090557 | 0.909413 | 1e-05 |
| GA-1A | 1.6e-05 | 1.1e-05 | 0.169616 | 0.830347 | 1e-05 |
| GA-2A | 1e-05 | 0.045916 | 0.091822 | 0.862242 | 1e-05 |
| GA-3A | 1e-05 | 0.003179 | 0.025796 | 0.971005 | 1e-05 |
| MR-1A | 1.1e-05 | 0.014142 | 0.166407 | 0.81943 | 1e-05 |
| MR-2A | 1e-05 | 0.055062 | 0.155709 | 0.789208 | 1e-05 |
| MR-3A | 0.021335 | 0.031569 | 0.168253 | 0.778833 | 1e-05 |
| MR-4A | 1.1e-05 | 0.052578 | 0.166597 | 0.780803 | 1e-05 |
| MR-5A | 0.008486 | 0.099916 | 0.099402 | 0.792187 | 1e-05 |
| MDG-6A | 1.7e-05 | 0.062413 | 0.380217 | 0.557342 | 1e-05 |
| MDG-1A | 1e-05 | 0.032316 | 0.317638 | 0.650026 | 1e-05 |
| MDG-2A | 1e-05 | 0.033143 | 0.321163 | 0.645674 | 1e-05 |
| MDG-3A | 1e-05 | 0.015286 | 0.353232 | 0.631462 | 1e-05 |
| MDG-4A | 0.006289 | 0.063181 | 0.352626 | 0.577894 | 1e-05 |
| MDG-5A | 1e-05 | 0.035505 | 0.346208 | 0.618267 | 1e-05 |
| AJ-10A | 1e-05 | 0.01871 | 0.820547 | 0.160723 | 1e-05 |
| AJ-12A | 1e-05 | 1e-05 | 0.888636 | 0.111334 | 1e-05 |
| AJ-13A | 1e-05 | 1e-05 | 0.855805 | 0.144165 | 1e-05 |
| AJ-14A | 1e-05 | 1e-05 | 0.804751 | 0.195219 | 1e-05 |
| AJ-1A | 1e-05 | 0.000965 | 0.795804 | 0.203211 | 1e-05 |
| AJ-3A | 1e-05 | 1e-05 | 0.752545 | 0.214507 | 0.032928 |
| AJ-4A | 1.2e-05 | 0.037547 | 0.88696 | 0.075471 | 1e-05 |
| AJ-5A | 1e-05 | 1e-05 | 0.820379 | 0.179591 | 1e-05 |
| AJ-7A | 1e-05 | 0.070157 | 0.738094 | 0.164444 | 0.027296 |
| AJ-9A | 1e-05 | 1e-05 | 0.950201 | 0.049769 | 1e-05 |
| XGG-1A | 1e-05 | 1e-05 | 0.99996 | 1e-05 | 1e-05 |
| XGG-2A | 1e-05 | 1e-05 | 0.99996 | 1e-05 | 1e-05 |
| XGG-3A | 1e-05 | 1e-05 | 0.99996 | 1e-05 | 1e-05 |
| XGG-4A | 1e-05 | 1e-05 | 0.99996 | 1e-05 | 1e-05 |
| XGG-5A | 1e-05 | 1e-05 | 0.99996 | 1e-05 | 1e-05 |
| PSR-01A | 1e-05 | 1e-05 | 0.99996 | 1e-05 | 1e-05 |
| PSR-04A | 1e-05 | 1e-05 | 0.99996 | 1e-05 | 1e-05 |
| PSR-10A | 1e-05 | 1e-05 | 0.99996 | 1e-05 | 1e-05 |
| PSR-13A | 1e-05 | 1e-05 | 0.966496 | 0.033474 | 1e-05 |
| LR-10A | 1.2e-05 | 1e-05 | 0.999958 | 1e-05 | 1e-05 |
| LR-1A | 1e-05 | 1e-05 | 0.99996 | 1e-05 | 1e-05 |
| LR-2A | 1e-05 | 1e-05 | 0.99996 | 1e-05 | 1e-05 |
| LR-3A | 1e-05 | 1e-05 | 0.99996 | 1e-05 | 1e-05 |
| LR-4A | 1e-05 | 1e-05 | 0.99996 | 1e-05 | 1e-05 |
| LR-5A | 1e-05 | 1e-05 | 0.99996 | 1e-05 | 1e-05 |
| LR-6A | 1e-05 | 0.009657 | 0.983198 | 1e-05 | 0.007126 |
| LR-7A | 1e-05 | 1e-05 | 0.999958 | 1.2e-05 | 1e-05 |
| LR-8A | 1.1e-05 | 1e-05 | 0.999959 | 1e-05 | 1e-05 |
| LR-9A | 1e-05 | 1e-05 | 0.99996 | 1e-05 | 1e-05 |
| BYR-10A | 1e-05 | 1e-05 | 0.99996 | 1e-05 | 1e-05 |
| BYR-1A | 1e-05 | 1e-05 | 0.99996 | 1e-05 | 1e-05 |
| BYR-2A | 1e-05 | 1.6e-05 | 0.999954 | 1e-05 | 1e-05 |
| BYR-3A | 1e-05 | 1.8e-05 | 0.999952 | 1e-05 | 1e-05 |
| BYR-4A | 1e-05 | 1e-05 | 0.99996 | 1e-05 | 1e-05 |
| BYR-5A | 1e-05 | 1e-05 | 0.99996 | 1e-05 | 1e-05 |
| BYR-6A | 1e-05 | 1e-05 | 0.99996 | 1e-05 | 1e-05 |
| BYR-7A | 1e-05 | 0.01955 | 0.98042 | 1e-05 | 1e-05 |

K = 6

|  |  |  |  |  |  |  |
| --- | --- | --- | --- | --- | --- | --- |
| CDG-1A | 1e-05 | 0.99995 | 1e-05 | 1e-05 | 1e-05 | 1e-05 |
| CDG-2A | 1e-05 | 0.99995 | 1e-05 | 1e-05 | 1e-05 | 1e-05 |
| CDG-3A | 1e-05 | 0.999949 | 1e-05 | 1.1e-05 | 1e-05 | 1e-05 |
| CDG-4A | 1e-05 | 0.99995 | 1e-05 | 1e-05 | 1e-05 | 1e-05 |
| CDG-5A | 1e-05 | 0.99995 | 1e-05 | 1e-05 | 1e-05 | 1e-05 |
| CDG-6A | 1e-05 | 0.99995 | 1e-05 | 1e-05 | 1e-05 | 1e-05 |
| CDG-7A | 1e-05 | 0.99995 | 1e-05 | 1e-05 | 1e-05 | 1e-05 |
| CDG-8A | 1e-05 | 0.99995 | 1e-05 | 1e-05 | 1e-05 | 1e-05 |
| CDG-9A | 1.1e-05 | 0.999949 | 1e-05 | 1e-05 | 1e-05 | 1e-05 |
| CDG-10A | 1e-05 | 0.999946 | 1.4e-05 | 1e-05 | 1e-05 | 1e-05 |
| TKX-1A | 1e-05 | 1e-05 | 1e-05 | 1e-05 | 1e-05 | 0.99995 |
| TKX-2A | 1e-05 | 1e-05 | 1e-05 | 1e-05 | 1e-05 | 0.99995 |
| TKX-3A | 1e-05 | 1e-05 | 1e-05 | 1e-05 | 1e-05 | 0.99995 |
| TKX-4A | 1e-05 | 1e-05 | 1e-05 | 1e-05 | 1e-05 | 0.99995 |
| TKX-5A | 1e-05 | 1e-05 | 1e-05 | 1e-05 | 1e-05 | 0.99995 |
| TKX-6A | 1e-05 | 1e-05 | 1e-05 | 1e-05 | 1e-05 | 0.99995 |
| TKX-7A | 1e-05 | 1e-05 | 1e-05 | 1e-05 | 1e-05 | 0.99995 |
| TKX-8A | 1e-05 | 1e-05 | 1e-05 | 1e-05 | 1e-05 | 0.99995 |
| TKX-9A | 1e-05 | 1.1e-05 | 1e-05 | 1e-05 | 1e-05 | 0.999949 |
| GJ-1A | 1e-05 | 0.055466 | 1e-05 | 1e-05 | 0.944494 | 1e-05 |
| GJ-2A | 1e-05 | 0.056702 | 1e-05 | 1e-05 | 0.943258 | 1e-05 |
| GJ-3A | 1e-05 | 1e-05 | 1e-05 | 1e-05 | 0.99995 | 1e-05 |
| GJ-4A | 1e-05 | 1e-05 | 1e-05 | 1e-05 | 0.99995 | 1e-05 |
| GJ-5A | 1e-05 | 0.061896 | 1e-05 | 1e-05 | 0.938064 | 1e-05 |
| GJ-6A | 1e-05 | 1e-05 | 1e-05 | 1e-05 | 0.99995 | 1e-05 |
| GJ-7A | 1e-05 | 0.004366 | 1e-05 | 1e-05 | 0.995594 | 1e-05 |
| GJ-8A | 1e-05 | 1e-05 | 1e-05 | 1e-05 | 0.99995 | 1e-05 |
| GJ-9A | 1e-05 | 1e-05 | 1e-05 | 1e-05 | 0.99995 | 1e-05 |
| TZC-1A | 1e-05 | 0.002356 | 1e-05 | 1e-05 | 0.997604 | 1e-05 |
| TZC-2A | 1e-05 | 1e-05 | 1e-05 | 1e-05 | 0.99995 | 1e-05 |
| TZC-3A | 0.13262 | 1e-05 | 1e-05 | 1e-05 | 0.86734 | 1e-05 |
| TZC-4A | 1e-05 | 1e-05 | 1e-05 | 1e-05 | 0.99995 | 1e-05 |
| TZC-01A | 1e-05 | 1e-05 | 1e-05 | 1e-05 | 0.965907 | 0.034053 |
| TZC-04A | 1e-05 | 1e-05 | 1e-05 | 1e-05 | 0.99995 | 1e-05 |
| TZC-07A | 1e-05 | 1e-05 | 1e-05 | 1e-05 | 0.96712 | 0.03284 |
| TZC-11A | 1e-05 | 1.1e-05 | 1e-05 | 1e-05 | 0.999949 | 1e-05 |
| TZC-13A | 1e-05 | 1e-05 | 1e-05 | 1e-05 | 0.99995 | 1e-05 |
| DBG-10A | 1e-05 | 1e-05 | 1e-05 | 1e-05 | 0.99995 | 1e-05 |
| DBG-1A | 1e-05 | 0.0093 | 1e-05 | 0.052896 | 0.937774 | 1e-05 |
| DBG-2A | 1e-05 | 0.02456 | 0.059627 | 0.002879 | 0.910632 | 0.002291 |
| DBG-3A | 1e-05 | 1e-05 | 0.037575 | 1e-05 | 0.954588 | 0.007807 |
| DBG-4A | 0.079603 | 1e-05 | 1e-05 | 1e-05 | 0.920357 | 1e-05 |
| DBG-5A | 1e-05 | 1e-05 | 1e-05 | 1e-05 | 0.99995 | 1e-05 |
| DBG-6A | 1e-05 | 0.017357 | 1e-05 | 1e-05 | 0.935867 | 0.046747 |
| DBG-7A | 1e-05 | 1.3e-05 | 1e-05 | 1e-05 | 0.981896 | 0.018061 |
| DBG-8A | 0.021401 | 0.031889 | 1e-05 | 1e-05 | 0.946679 | 1e-05 |
| DBG-9A | 0.046896 | 0.013895 | 1e-05 | 1e-05 | 0.913569 | 0.02562 |
| TEG-10A | 1e-05 | 1e-05 | 1e-05 | 1e-05 | 0.99995 | 1e-05 |
| TEG-1A | 0.028332 | 1e-05 | 0.036854 | 0.066776 | 0.868018 | 1e-05 |
| TEG-2A | 1e-05 | 1e-05 | 0.157922 | 1e-05 | 0.838916 | 0.003131 |
| TEG-3A | 1e-05 | 1e-05 | 0.049539 | 1e-05 | 0.950421 | 1e-05 |
| TEG-4A | 0.029155 | 1e-05 | 0.025104 | 1e-05 | 0.945711 | 1e-05 |
| TEG-5A | 1e-05 | 1.2e-05 | 0.015522 | 1e-05 | 0.984436 | 1e-05 |
| TEG-6A | 1e-05 | 0.012288 | 0.154475 | 1e-05 | 0.833207 | 1e-05 |
| TEG-7A | 1e-05 | 1e-05 | 0.168486 | 0.052435 | 0.779049 | 1e-05 |
| TEG-8A | 1e-05 | 1e-05 | 0.107884 | 1e-05 | 0.892076 | 1e-05 |
| TEG-9A | 1e-05 | 1e-05 | 0.062596 | 0.004058 | 0.933316 | 1e-05 |
| ZPZ-10A | 1e-05 | 1e-05 | 1e-05 | 1e-05 | 0.99995 | 1e-05 |
| ZPZ-1A | 1e-05 | 1e-05 | 1e-05 | 1e-05 | 0.99995 | 1e-05 |
| ZPZ-2A | 1e-05 | 1e-05 | 0.03416 | 1e-05 | 0.9658 | 1e-05 |
| ZPZ-3A | 1e-05 | 1e-05 | 0.137948 | 1e-05 | 0.862012 | 1e-05 |
| ZPZ-4A | 1e-05 | 1e-05 | 1e-05 | 1e-05 | 0.99995 | 1e-05 |
| ZPZ-5A | 1e-05 | 1e-05 | 0.053699 | 1e-05 | 0.946261 | 1e-05 |
| ZPZ-6A | 1e-05 | 1e-05 | 0.306538 | 8.8e-05 | 0.693344 | 1e-05 |
| ZPZ-7A | 1e-05 | 1e-05 | 0.120876 | 1e-05 | 0.879084 | 1e-05 |
| ZPZ-8A | 1e-05 | 1e-05 | 1e-05 | 1e-05 | 0.99995 | 1e-05 |
| ZPZ-9A | 1e-05 | 1e-05 | 1.1e-05 | 1e-05 | 0.999949 | 1e-05 |
| YLJ1-1A | 1e-05 | 1e-05 | 1e-05 | 1e-05 | 0.99995 | 1e-05 |
| YLJ1-2A | 1e-05 | 1e-05 | 1e-05 | 1e-05 | 0.99995 | 1e-05 |
| YLJ1-3A | 1e-05 | 1e-05 | 1e-05 | 1e-05 | 0.99995 | 1e-05 |
| YLJ1-4A | 1e-05 | 1e-05 | 1e-05 | 1e-05 | 0.99995 | 1e-05 |
| YLJ1-5A | 1e-05 | 1e-05 | 1e-05 | 1e-05 | 0.99995 | 1e-05 |
| YLJ1-6A | 1e-05 | 1e-05 | 0.03896 | 1e-05 | 0.961 | 1e-05 |
| YLJ1-7A | 1e-05 | 1e-05 | 1e-05 | 1e-05 | 0.99995 | 1e-05 |
| YLJ1-8A | 1e-05 | 1e-05 | 0.140157 | 1e-05 | 0.859803 | 1e-05 |
| YLJ2-1A | 1e-05 | 1e-05 | 0.027925 | 1e-05 | 0.972035 | 1e-05 |
| YLJ2-2A | 1e-05 | 1e-05 | 1e-05 | 1e-05 | 0.99995 | 1e-05 |
| YLJ2-3A | 1e-05 | 1e-05 | 0.028307 | 1e-05 | 0.971653 | 1e-05 |
| YLJ2-4A | 1e-05 | 1e-05 | 0.212337 | 1e-05 | 0.787623 | 1e-05 |
| YLJ2-5A | 1e-05 | 1e-05 | 1e-05 | 1e-05 | 0.99995 | 1e-05 |
| YLJ2-6B | 1e-05 | 1e-05 | 0.033008 | 1e-05 | 0.966952 | 1e-05 |
| YLJ2-7A | 1e-05 | 1e-05 | 1e-05 | 1e-05 | 0.99995 | 1e-05 |
| YLJ2-8A | 1e-05 | 1e-05 | 0.228995 | 1e-05 | 0.770965 | 1e-05 |
| LM-10A | 1e-05 | 1e-05 | 0.99995 | 1e-05 | 1e-05 | 1e-05 |
| LM-1A | 1e-05 | 1e-05 | 0.99995 | 1e-05 | 1e-05 | 1e-05 |
| LM-2A | 1e-05 | 1e-05 | 0.99995 | 1e-05 | 1e-05 | 1e-05 |
| LM-3A | 1e-05 | 1e-05 | 0.999941 | 1.9e-05 | 1e-05 | 1e-05 |
| LM-4A | 1e-05 | 1e-05 | 0.99995 | 1e-05 | 1e-05 | 1e-05 |
| LM-5A | 1e-05 | 1e-05 | 0.99995 | 1e-05 | 1e-05 | 1e-05 |
| LM-6A | 1e-05 | 1e-05 | 0.99995 | 1e-05 | 1e-05 | 1e-05 |
| LM-7A | 1e-05 | 1e-05 | 0.99995 | 1e-05 | 1e-05 | 1e-05 |
| LM-8A | 1.9e-05 | 1.4e-05 | 0.999936 | 1.1e-05 | 1e-05 | 1e-05 |
| LM-9A | 1e-05 | 1e-05 | 0.99995 | 1e-05 | 1e-05 | 1e-05 |
| LM-11A | 1e-05 | 1e-05 | 0.99995 | 1e-05 | 1e-05 | 1e-05 |
| LM-12A | 1e-05 | 1e-05 | 0.99995 | 1e-05 | 1e-05 | 1e-05 |
| LM-13A | 1e-05 | 1e-05 | 0.99995 | 1e-05 | 1e-05 | 1e-05 |
| LM-14A | 1e-05 | 1e-05 | 0.99995 | 1e-05 | 1e-05 | 1e-05 |
| LM-15A | 1e-05 | 1e-05 | 0.99995 | 1e-05 | 1e-05 | 1e-05 |
| BYP1-1A | 0.071016 | 1e-05 | 0.908098 | 1e-05 | 0.020856 | 1e-05 |
| BYP1-2A | 0.133844 | 1e-05 | 0.866116 | 1e-05 | 1e-05 | 1e-05 |
| BYP1-3A | 0.031592 | 1e-05 | 0.843485 | 1e-05 | 0.124893 | 1e-05 |
| BYP1-4A | 0.097182 | 1e-05 | 0.845944 | 1e-05 | 0.056844 | 1e-05 |
| BYP1-5A | 0.097438 | 1e-05 | 0.808757 | 1e-05 | 0.093775 | 1e-05 |
| BYP1-6A | 1e-05 | 1e-05 | 0.99995 | 1e-05 | 1e-05 | 1e-05 |
| BYP1-7A | 1e-05 | 1e-05 | 0.99995 | 1e-05 | 1e-05 | 1e-05 |
| BYP1-8A | 1e-05 | 1e-05 | 0.99995 | 1e-05 | 1e-05 | 1e-05 |
| BYP1-9A | 1e-05 | 1e-05 | 0.99995 | 1e-05 | 1e-05 | 1e-05 |
| BYP1-10A | 1e-05 | 0.029296 | 0.932647 | 0.038027 | 1e-05 | 1e-05 |
| BYP2-1A | 1e-05 | 1e-05 | 0.99995 | 1e-05 | 1e-05 | 1e-05 |
| BYP2-2A | 1e-05 | 1e-05 | 0.995834 | 1e-05 | 0.004126 | 1e-05 |
| BYP2-3A | 0.293682 | 1e-05 | 0.706278 | 1e-05 | 1e-05 | 1e-05 |
| BYP2-4A | 1e-05 | 1e-05 | 0.99995 | 1e-05 | 1e-05 | 1e-05 |
| BYP2-5A | 0.247204 | 1e-05 | 0.617579 | 1e-05 | 0.135187 | 1e-05 |
| BYP2-6A | 0.119247 | 1e-05 | 0.86603 | 1e-05 | 1e-05 | 0.014694 |
| BYP2-7A | 0.066402 | 1e-05 | 0.877186 | 1e-05 | 0.056382 | 1e-05 |
| BYP2-8A | 1e-05 | 1e-05 | 0.99995 | 1e-05 | 1e-05 | 1e-05 |
| BYP3-1A | 0.11093 | 1e-05 | 0.828076 | 1e-05 | 0.060964 | 1e-05 |
| BYP3-2A | 0.123441 | 1e-05 | 0.876519 | 1e-05 | 1e-05 | 1e-05 |
| BYP3-3A | 0.12761 | 1e-05 | 0.826027 | 1e-05 | 0.046333 | 1e-05 |
| BYP3-4A | 0.091241 | 1e-05 | 0.898667 | 0.010062 | 1e-05 | 1e-05 |
| BYP3-5A | 0.150716 | 0.005692 | 0.830329 | 0.013243 | 1e-05 | 1e-05 |
| BYP3-6A | 0.088054 | 1e-05 | 0.814372 | 1e-05 | 0.097544 | 1e-05 |
| BYP3-7A | 0.10786 | 1e-05 | 0.8921 | 1e-05 | 1e-05 | 1e-05 |
| BYP3-8A | 0.103511 | 1e-05 | 0.896449 | 1e-05 | 1e-05 | 1e-05 |
| KBG-1A | 0.99995 | 1e-05 | 1e-05 | 1e-05 | 1e-05 | 1e-05 |
| KBG-2A | 0.937391 | 1e-05 | 1e-05 | 1e-05 | 0.062569 | 1e-05 |
| KBG-3A | 0.99995 | 1e-05 | 1e-05 | 1e-05 | 1e-05 | 1e-05 |
| KBG-4A | 0.99995 | 1e-05 | 1e-05 | 1e-05 | 1e-05 | 1e-05 |
| KBG-5A | 0.99995 | 1e-05 | 1e-05 | 1e-05 | 1e-05 | 1e-05 |
| QJG-1A | 0.99995 | 1e-05 | 1e-05 | 1e-05 | 1e-05 | 1e-05 |
| QJG-2A | 0.99995 | 1e-05 | 1e-05 | 1e-05 | 1e-05 | 1e-05 |
| QJG-3A | 0.99995 | 1e-05 | 1e-05 | 1e-05 | 1e-05 | 1e-05 |
| QJG-4A | 0.99995 | 1e-05 | 1e-05 | 1e-05 | 1e-05 | 1e-05 |
| QJG-5A | 0.99995 | 1e-05 | 1e-05 | 1e-05 | 1e-05 | 1e-05 |
| LTG-1A | 0.725664 | 1e-05 | 0.160716 | 1e-05 | 0.11359 | 1e-05 |
| LTG-2A | 0.873138 | 1e-05 | 0.038828 | 1e-05 | 0.052278 | 0.035736 |
| LTG-3A | 0.99995 | 1e-05 | 1e-05 | 1e-05 | 1e-05 | 1e-05 |
| LTG-4A | 0.99995 | 1e-05 | 1e-05 | 1e-05 | 1e-05 | 1e-05 |
| LTG-5A | 0.891743 | 1e-05 | 0.044703 | 1e-05 | 1e-05 | 0.063524 |
| BW-10B | 0.984396 | 1e-05 | 0.015564 | 1e-05 | 1e-05 | 1e-05 |
| BW-1B | 0.99995 | 1e-05 | 1e-05 | 1e-05 | 1e-05 | 1e-05 |
| BW-2B | 0.99995 | 1e-05 | 1e-05 | 1e-05 | 1e-05 | 1e-05 |
| BW-3B | 0.99995 | 1e-05 | 1e-05 | 1e-05 | 1e-05 | 1e-05 |
| BW-4B | 0.957181 | 1e-05 | 0.042779 | 1e-05 | 1e-05 | 1e-05 |
| BW-5B | 0.99994 | 1e-05 | 2e-05 | 1e-05 | 1e-05 | 1e-05 |
| BW-6B | 0.99995 | 1e-05 | 1e-05 | 1e-05 | 1e-05 | 1e-05 |
| BW-7A | 0.99995 | 1e-05 | 1e-05 | 1e-05 | 1e-05 | 1e-05 |
| BW-8B | 0.99995 | 1e-05 | 1e-05 | 1e-05 | 1e-05 | 1e-05 |
| BW-9A | 0.99995 | 1e-05 | 1e-05 | 1e-05 | 1e-05 | 1e-05 |
| GA-1A | 0.99995 | 1e-05 | 1e-05 | 1e-05 | 1e-05 | 1e-05 |
| GA-2A | 0.923536 | 1e-05 | 0.076424 | 1e-05 | 1e-05 | 1e-05 |
| GA-3A | 0.851685 | 1e-05 | 0.148275 | 1e-05 | 1e-05 | 1e-05 |
| MR-1A | 0.999949 | 1e-05 | 1.1e-05 | 1e-05 | 1e-05 | 1e-05 |
| MR-2A | 0.999947 | 1e-05 | 1.3e-05 | 1e-05 | 1e-05 | 1e-05 |
| MR-3A | 0.887881 | 1e-05 | 0.087554 | 0.012483 | 1e-05 | 0.012061 |
| MR-4A | 0.888525 | 1e-05 | 0.111435 | 1e-05 | 1e-05 | 1e-05 |
| MR-5A | 0.992078 | 0.007882 | 1e-05 | 1e-05 | 1e-05 | 1e-05 |
| MDG-6A | 0.830624 | 1e-05 | 1e-05 | 0.169336 | 1e-05 | 1e-05 |
| MDG-1A | 0.964521 | 1e-05 | 1e-05 | 0.035439 | 1e-05 | 1e-05 |
| MDG-2A | 0.9503 | 1e-05 | 1e-05 | 0.04966 | 1e-05 | 1e-05 |
| MDG-3A | 0.86222 | 1e-05 | 1e-05 | 0.13774 | 1e-05 | 1e-05 |
| MDG-4A | 0.821534 | 1e-05 | 1e-05 | 0.178426 | 1e-05 | 1e-05 |
| MDG-5A | 0.881662 | 1e-05 | 1e-05 | 0.118298 | 1e-05 | 1e-05 |
| AJ-10A | 0.315728 | 1e-05 | 1e-05 | 0.684232 | 1e-05 | 1e-05 |
| AJ-12A | 0.290134 | 1e-05 | 1.7e-05 | 0.709819 | 1e-05 | 1e-05 |
| AJ-13A | 0.330641 | 1e-05 | 1e-05 | 0.669319 | 1e-05 | 1e-05 |
| AJ-14A | 0.389824 | 1e-05 | 1e-05 | 0.610136 | 1e-05 | 1e-05 |
| AJ-1A | 0.352299 | 1e-05 | 1e-05 | 0.647661 | 1e-05 | 1e-05 |
| AJ-3A | 0.366046 | 1e-05 | 1e-05 | 0.633914 | 1e-05 | 1e-05 |
| AJ-4A | 0.217063 | 0.006824 | 1e-05 | 0.776083 | 1e-05 | 1e-05 |
| AJ-5A | 0.290897 | 1e-05 | 1e-05 | 0.709063 | 1e-05 | 1e-05 |
| AJ-7A | 0.376669 | 1e-05 | 1e-05 | 0.599878 | 0.023424 | 1e-05 |
| AJ-9A | 0.220648 | 1e-05 | 1e-05 | 0.779312 | 1e-05 | 1e-05 |
| XGG-1A | 1.6e-05 | 1e-05 | 1e-05 | 0.999944 | 1e-05 | 1e-05 |
| XGG-2A | 1.7e-05 | 1e-05 | 1e-05 | 0.999943 | 1e-05 | 1e-05 |
| XGG-3A | 1e-05 | 1e-05 | 1e-05 | 0.99995 | 1e-05 | 1e-05 |
| XGG-4A | 1e-05 | 1e-05 | 1e-05 | 0.99995 | 1e-05 | 1e-05 |
| XGG-5A | 0.05114 | 1e-05 | 1e-05 | 0.94882 | 1e-05 | 1e-05 |
| PSR-01A | 1.6e-05 | 1e-05 | 1e-05 | 0.999944 | 1e-05 | 1e-05 |
| PSR-04A | 1e-05 | 1e-05 | 1e-05 | 0.99995 | 1e-05 | 1e-05 |
| PSR-10A | 1e-05 | 1e-05 | 1e-05 | 0.99995 | 1e-05 | 1e-05 |
| PSR-13A | 0.000493 | 1e-05 | 0.029328 | 0.970149 | 1e-05 | 1e-05 |
| LR-10A | 1e-05 | 1e-05 | 1e-05 | 0.99995 | 1e-05 | 1e-05 |
| LR-1A | 0.05559 | 1e-05 | 1e-05 | 0.944369 | 1.1e-05 | 1e-05 |
| LR-2A | 1e-05 | 1e-05 | 1e-05 | 0.99995 | 1e-05 | 1e-05 |
| LR-3A | 1e-05 | 1e-05 | 1e-05 | 0.99995 | 1e-05 | 1e-05 |
| LR-4A | 1e-05 | 1e-05 | 1e-05 | 0.99995 | 1e-05 | 1e-05 |
| LR-5A | 1e-05 | 1e-05 | 1.5e-05 | 0.999945 | 1e-05 | 1e-05 |
| LR-6A | 0.022062 | 1e-05 | 1e-05 | 0.963167 | 0.014741 | 1e-05 |
| LR-7A | 1e-05 | 1e-05 | 1e-05 | 0.99995 | 1e-05 | 1e-05 |
| LR-8A | 1e-05 | 1e-05 | 1e-05 | 0.99995 | 1e-05 | 1e-05 |
| LR-9A | 1e-05 | 1e-05 | 1e-05 | 0.99995 | 1e-05 | 1e-05 |
| BYR-10A | 1e-05 | 1e-05 | 1e-05 | 0.99995 | 1e-05 | 1e-05 |
| BYR-1A | 1e-05 | 1e-05 | 1e-05 | 0.99995 | 1e-05 | 1e-05 |
| BYR-2A | 1e-05 | 1e-05 | 1e-05 | 0.99995 | 1e-05 | 1e-05 |
| BYR-3A | 1e-05 | 1e-05 | 1e-05 | 0.99995 | 1e-05 | 1e-05 |
| BYR-4A | 1.4e-05 | 1e-05 | 1e-05 | 0.999946 | 1e-05 | 1e-05 |
| BYR-5A | 1e-05 | 1e-05 | 1e-05 | 0.99995 | 1e-05 | 1e-05 |
| BYR-6A | 1e-05 | 1e-05 | 1e-05 | 0.99995 | 1e-05 | 1e-05 |
| BYR-7A | 0.027395 | 1e-05 | 1e-05 | 0.972565 | 1e-05 | 1e-05 |

K = 2

|  |  |  |
| --- | --- | --- |
| CDG-1A | 1e-05 | 0.99999 |
| CDG-2A | 1e-05 | 0.99999 |
| CDG-3A | 1e-05 | 0.99999 |
| CDG-4A | 1e-05 | 0.99999 |
| CDG-5A | 1e-05 | 0.99999 |
| CDG-6A | 1e-05 | 0.99999 |
| CDG-7A | 1e-05 | 0.99999 |
| CDG-8A | 1e-05 | 0.99999 |
| CDG-9A | 1e-05 | 0.99999 |
| CDG-10A | 1e-05 | 0.99999 |
| TKX-1A | 1e-05 | 0.99999 |
| TKX-2A | 1e-05 | 0.99999 |
| TKX-3A | 1e-05 | 0.99999 |
| TKX-4A | 1e-05 | 0.99999 |
| TKX-5A | 1e-05 | 0.99999 |
| TKX-6A | 1e-05 | 0.99999 |
| TKX-7A | 1e-05 | 0.99999 |
| TKX-8A | 1e-05 | 0.99999 |
| TKX-9A | 1e-05 | 0.99999 |
| GJ-1A | 0.028311 | 0.971689 |
| GJ-2A | 0.029805 | 0.970195 |
| GJ-3A | 1e-05 | 0.99999 |
| GJ-4A | 1e-05 | 0.99999 |
| GJ-5A | 1e-05 | 0.99999 |
| GJ-6A | 1e-05 | 0.99999 |
| GJ-7A | 1e-05 | 0.99999 |
| GJ-8A | 1e-05 | 0.99999 |
| GJ-9A | 1e-05 | 0.99999 |
| TZC-1A | 1e-05 | 0.99999 |
| TZC-2A | 1e-05 | 0.99999 |
| TZC-3A | 0.018809 | 0.981191 |
| TZC-4A | 1e-05 | 0.99999 |
| TZC-01A | 0.017739 | 0.982261 |
| TZC-04A | 0.041588 | 0.958412 |
| TZC-07A | 0.058203 | 0.941797 |
| TZC-11A | 1e-05 | 0.99999 |
| TZC-13A | 1e-05 | 0.99999 |
| DBG-10A | 1e-05 | 0.99999 |
| DBG-1A | 0.096486 | 0.903514 |
| DBG-2A | 0.050855 | 0.949145 |
| DBG-3A | 0.045962 | 0.954038 |
| DBG-4A | 0.04469 | 0.95531 |
| DBG-5A | 0.018422 | 0.981578 |
| DBG-6A | 0.054976 | 0.945024 |
| DBG-7A | 0.051064 | 0.948936 |
| DBG-8A | 0.023803 | 0.976197 |
| DBG-9A | 0.010101 | 0.989899 |
| TEG-10A | 0.004305 | 0.995695 |
| TEG-1A | 0.12317 | 0.87683 |
| TEG-2A | 0.024071 | 0.975929 |
| TEG-3A | 1e-05 | 0.99999 |
| TEG-4A | 1e-05 | 0.99999 |
| TEG-5A | 1e-05 | 0.99999 |
| TEG-6A | 1e-05 | 0.99999 |
| TEG-7A | 0.059972 | 0.940028 |
| TEG-8A | 1e-05 | 0.99999 |
| TEG-9A | 0.059993 | 0.940007 |
| ZPZ-10A | 1e-05 | 0.99999 |
| ZPZ-1A | 1e-05 | 0.99999 |
| ZPZ-2A | 1e-05 | 0.99999 |
| ZPZ-3A | 1e-05 | 0.99999 |
| ZPZ-4A | 1e-05 | 0.99999 |
| ZPZ-5A | 1e-05 | 0.99999 |
| ZPZ-6A | 0.022619 | 0.977381 |
| ZPZ-7A | 1e-05 | 0.99999 |
| ZPZ-8A | 1e-05 | 0.99999 |
| ZPZ-9A | 1e-05 | 0.99999 |
| YLJ1-1A | 1e-05 | 0.99999 |
| YLJ1-2A | 1e-05 | 0.99999 |
| YLJ1-3A | 1e-05 | 0.99999 |
| YLJ1-4A | 1e-05 | 0.99999 |
| YLJ1-5A | 1e-05 | 0.99999 |
| YLJ1-6A | 1e-05 | 0.99999 |
| YLJ1-7A | 1e-05 | 0.99999 |
| YLJ1-8A | 1e-05 | 0.99999 |
| YLJ2-1A | 0.009603 | 0.990397 |
| YLJ2-2A | 1e-05 | 0.99999 |
| YLJ2-3A | 0.034024 | 0.965976 |
| YLJ2-4A | 0.015566 | 0.984434 |
| YLJ2-5A | 0.014597 | 0.985403 |
| YLJ2-6B | 1e-05 | 0.99999 |
| YLJ2-7A | 1e-05 | 0.99999 |
| YLJ2-8A | 0.035092 | 0.964908 |
| LM-10A | 1e-05 | 0.99999 |
| LM-1A | 1e-05 | 0.99999 |
| LM-2A | 1e-05 | 0.99999 |
| LM-3A | 1e-05 | 0.99999 |
| LM-4A | 1e-05 | 0.99999 |
| LM-5A | 1e-05 | 0.99999 |
| LM-6A | 1e-05 | 0.99999 |
| LM-7A | 1e-05 | 0.99999 |
| LM-8A | 1e-05 | 0.99999 |
| LM-9A | 1e-05 | 0.99999 |
| LM-11A | 1e-05 | 0.99999 |
| LM-12A | 1e-05 | 0.99999 |
| LM-13A | 1e-05 | 0.99999 |
| LM-14A | 1e-05 | 0.99999 |
| LM-15A | 1e-05 | 0.99999 |
| BYP1-1A | 0.014568 | 0.985432 |
| BYP1-2A | 0.037052 | 0.962948 |
| BYP1-3A | 1e-05 | 0.99999 |
| BYP1-4A | 1e-05 | 0.99999 |
| BYP1-5A | 1e-05 | 0.99999 |
| BYP1-6A | 1e-05 | 0.99999 |
| BYP1-7A | 1e-05 | 0.99999 |
| BYP1-8A | 1e-05 | 0.99999 |
| BYP1-9A | 1e-05 | 0.99999 |
| BYP1-10A | 0.087069 | 0.912931 |
| BYP2-1A | 1e-05 | 0.99999 |
| BYP2-2A | 1e-05 | 0.99999 |
| BYP2-3A | 0.079415 | 0.920585 |
| BYP2-4A | 1e-05 | 0.99999 |
| BYP2-5A | 0.055573 | 0.944427 |
| BYP2-6A | 0.016101 | 0.983899 |
| BYP2-7A | 1e-05 | 0.99999 |
| BYP2-8A | 1e-05 | 0.99999 |
| BYP3-1A | 0.007045 | 0.992955 |
| BYP3-2A | 0.038227 | 0.961773 |
| BYP3-3A | 0.035127 | 0.964873 |
| BYP3-4A | 0.070841 | 0.929159 |
| BYP3-5A | 0.099555 | 0.900445 |
| BYP3-6A | 0.012215 | 0.987785 |
| BYP3-7A | 0.113779 | 0.886221 |
| BYP3-8A | 0.050406 | 0.949594 |
| KBG-1A | 0.34878 | 0.65122 |
| KBG-2A | 0.326623 | 0.673377 |
| KBG-3A | 0.271357 | 0.728643 |
| KBG-4A | 0.301934 | 0.698066 |
| KBG-5A | 0.398256 | 0.601744 |
| QJG-1A | 0.450923 | 0.549077 |
| QJG-2A | 0.338439 | 0.661561 |
| QJG-3A | 0.442257 | 0.557743 |
| QJG-4A | 0.422574 | 0.577426 |
| QJG-5A | 0.429487 | 0.570513 |
| LTG-1A | 0.166455 | 0.833545 |
| LTG-2A | 0.302687 | 0.697313 |
| LTG-3A | 0.41726 | 0.58274 |
| LTG-4A | 0.421055 | 0.578945 |
| LTG-5A | 0.24915 | 0.75085 |
| BW-10B | 0.189454 | 0.810546 |
| BW-1B | 0.25089 | 0.74911 |
| BW-2B | 0.221006 | 0.778994 |
| BW-3B | 0.248351 | 0.751649 |
| BW-4B | 0.226759 | 0.773241 |
| BW-5B | 0.224142 | 0.775858 |
| BW-6B | 0.218108 | 0.781892 |
| BW-7A | 0.168316 | 0.831684 |
| BW-8B | 0.255861 | 0.744139 |
| BW-9A | 0.244882 | 0.755118 |
| GA-1A | 0.365211 | 0.634789 |
| GA-2A | 0.229679 | 0.770321 |
| GA-3A | 0.182915 | 0.817085 |
| MR-1A | 0.278646 | 0.721354 |
| MR-2A | 0.272402 | 0.727598 |
| MR-3A | 0.264014 | 0.735986 |
| MR-4A | 0.30238 | 0.69762 |
| MR-5A | 0.195378 | 0.804622 |
| MDG-6A | 0.583858 | 0.416142 |
| MDG-1A | 0.555471 | 0.444529 |
| MDG-2A | 0.5487 | 0.4513 |
| MDG-3A | 0.542351 | 0.457649 |
| MDG-4A | 0.515591 | 0.484409 |
| MDG-5A | 0.48235 | 0.51765 |
| AJ-10A | 0.894744 | 0.105256 |
| AJ-12A | 0.99999 | 1e-05 |
| AJ-13A | 0.943807 | 0.056193 |
| AJ-14A | 0.931807 | 0.068193 |
| AJ-1A | 0.881414 | 0.118586 |
| AJ-3A | 0.819956 | 0.180044 |
| AJ-4A | 0.943728 | 0.056272 |
| AJ-5A | 0.906576 | 0.093424 |
| AJ-7A | 0.801404 | 0.198596 |
| AJ-9A | 0.99999 | 1e-05 |
| XGG-1A | 0.99999 | 1e-05 |
| XGG-2A | 0.99999 | 1e-05 |
| XGG-3A | 0.99999 | 1e-05 |
| XGG-4A | 0.99999 | 1e-05 |
| XGG-5A | 0.99999 | 1e-05 |
| PSR-01A | 0.99999 | 1e-05 |
| PSR-04A | 0.99999 | 1e-05 |
| PSR-10A | 0.99999 | 1e-05 |
| PSR-13A | 0.99999 | 1e-05 |
| LR-10A | 0.99999 | 1e-05 |
| LR-1A | 0.99999 | 1e-05 |
| LR-2A | 0.99999 | 1e-05 |
| LR-3A | 0.99999 | 1e-05 |
| LR-4A | 0.99999 | 1e-05 |
| LR-5A | 0.99999 | 1e-05 |
| LR-6A | 0.99999 | 1e-05 |
| LR-7A | 0.99999 | 1e-05 |
| LR-8A | 0.99999 | 1e-05 |
| LR-9A | 0.99999 | 1e-05 |
| BYR-10A | 0.99999 | 1e-05 |
| BYR-1A | 0.99999 | 1e-05 |
| BYR-2A | 0.99999 | 1e-05 |
| BYR-3A | 0.99999 | 1e-05 |
| BYR-4A | 0.99999 | 1e-05 |
| BYR-5A | 0.99999 | 1e-05 |
| BYR-6A | 0.99999 | 1e-05 |
| BYR-7A | 0.99999 | 1e-05 |

K = 7

|  |  |  |  |  |  |  |  |
| --- | --- | --- | --- | --- | --- | --- | --- |
| CDG-1A | 1e-05 | 1e-05 | 1e-05 | 1e-05 | 1e-05 | 0.99994 | 1e-05 |
| CDG-2A | 1e-05 | 1e-05 | 1e-05 | 1e-05 | 1e-05 | 0.99994 | 1e-05 |
| CDG-3A | 1e-05 | 1e-05 | 1e-05 | 1e-05 | 1e-05 | 0.99994 | 1e-05 |
| CDG-4A | 1e-05 | 1.7e-05 | 1e-05 | 1e-05 | 1e-05 | 0.999933 | 1e-05 |
| CDG-5A | 1.3e-05 | 1e-05 | 1e-05 | 1e-05 | 1e-05 | 0.999937 | 1e-05 |
| CDG-6A | 1e-05 | 1.1e-05 | 1e-05 | 1e-05 | 1e-05 | 0.999939 | 1e-05 |
| CDG-7A | 1e-05 | 1e-05 | 1e-05 | 1e-05 | 1e-05 | 0.99994 | 1e-05 |
| CDG-8A | 1e-05 | 1e-05 | 1e-05 | 1e-05 | 1e-05 | 0.99994 | 1e-05 |
| CDG-9A | 1e-05 | 1e-05 | 1e-05 | 1e-05 | 1e-05 | 0.99994 | 1e-05 |
| CDG-10A | 1e-05 | 1e-05 | 1e-05 | 1e-05 | 1e-05 | 0.99994 | 1e-05 |
| TKX-1A | 1e-05 | 1e-05 | 1e-05 | 1e-05 | 0.99994 | 1e-05 | 1e-05 |
| TKX-2A | 1e-05 | 1.6e-05 | 1.8e-05 | 1e-05 | 0.999926 | 1e-05 | 1e-05 |
| TKX-3A | 1.5e-05 | 2e-05 | 1e-05 | 1e-05 | 0.999925 | 1e-05 | 1e-05 |
| TKX-4A | 1e-05 | 1e-05 | 1e-05 | 1e-05 | 0.99994 | 1e-05 | 1e-05 |
| TKX-5A | 1e-05 | 1e-05 | 1e-05 | 1e-05 | 0.99994 | 1e-05 | 1e-05 |
| TKX-6A | 1e-05 | 1e-05 | 1.7e-05 | 1e-05 | 0.999933 | 1e-05 | 1e-05 |
| TKX-7A | 1e-05 | 1e-05 | 1e-05 | 1.2e-05 | 0.999937 | 1e-05 | 1e-05 |
| TKX-8A | 1e-05 | 2e-05 | 1e-05 | 1e-05 | 0.99993 | 1e-05 | 1e-05 |
| TKX-9A | 1e-05 | 1e-05 | 1e-05 | 1.1e-05 | 0.999939 | 1e-05 | 1e-05 |
| GJ-1A | 1e-05 | 0.99994 | 1e-05 | 1e-05 | 1e-05 | 1e-05 | 1e-05 |
| GJ-2A | 1e-05 | 0.99994 | 1e-05 | 1e-05 | 1e-05 | 1e-05 | 1e-05 |
| GJ-3A | 1e-05 | 0.99994 | 1e-05 | 1e-05 | 1e-05 | 1e-05 | 1e-05 |
| GJ-4A | 1e-05 | 0.999938 | 1e-05 | 1.2e-05 | 1e-05 | 1e-05 | 1e-05 |
| GJ-5A | 1e-05 | 0.99994 | 1e-05 | 1e-05 | 1e-05 | 1e-05 | 1e-05 |
| GJ-6A | 1e-05 | 0.99994 | 1e-05 | 1e-05 | 1e-05 | 1e-05 | 1e-05 |
| GJ-7A | 1e-05 | 0.999932 | 1.8e-05 | 1e-05 | 1e-05 | 1e-05 | 1e-05 |
| GJ-8A | 1e-05 | 0.99994 | 1e-05 | 1e-05 | 1e-05 | 1e-05 | 1e-05 |
| GJ-9A | 1e-05 | 0.999931 | 1.9e-05 | 1e-05 | 1e-05 | 1e-05 | 1e-05 |
| TZC-1A | 1e-05 | 0.99994 | 1e-05 | 1e-05 | 1e-05 | 1e-05 | 1e-05 |
| TZC-2A | 1.9e-05 | 0.999927 | 1e-05 | 1e-05 | 1.4e-05 | 1e-05 | 1e-05 |
| TZC-3A | 0.007055 | 0.888317 | 0.056859 | 0.04774 | 1e-05 | 1e-05 | 1e-05 |
| TZC-4A | 1e-05 | 0.999939 | 1.1e-05 | 1e-05 | 1e-05 | 1e-05 | 1e-05 |
| TZC-01A | 1e-05 | 0.999932 | 1e-05 | 1e-05 | 1.8e-05 | 1e-05 | 1e-05 |
| TZC-04A | 1e-05 | 0.99993 | 1.3e-05 | 1e-05 | 1.7e-05 | 1e-05 | 1e-05 |
| TZC-07A | 1e-05 | 0.99994 | 1e-05 | 1e-05 | 1e-05 | 1e-05 | 1e-05 |
| TZC-11A | 1e-05 | 0.999926 | 1e-05 | 1e-05 | 1.6e-05 | 1.9e-05 | 1e-05 |
| TZC-13A | 1e-05 | 0.99994 | 1e-05 | 1e-05 | 1e-05 | 1e-05 | 1e-05 |
| DBG-10A | 1e-05 | 0.99994 | 1e-05 | 1e-05 | 1e-05 | 1e-05 | 1e-05 |
| DBG-1A | 1e-05 | 0.84363 | 1e-05 | 0.124115 | 1e-05 | 1e-05 | 0.032216 |
| DBG-2A | 1e-05 | 0.99994 | 1e-05 | 1e-05 | 1e-05 | 1e-05 | 1e-05 |
| DBG-3A | 1e-05 | 0.99994 | 1e-05 | 1e-05 | 1e-05 | 1e-05 | 1e-05 |
| DBG-4A | 1e-05 | 0.99994 | 1e-05 | 1e-05 | 1e-05 | 1e-05 | 1e-05 |
| DBG-5A | 1.1e-05 | 0.881568 | 1e-05 | 0.118382 | 1e-05 | 1e-05 | 1e-05 |
| DBG-6A | 1e-05 | 0.999938 | 1e-05 | 1e-05 | 1.2e-05 | 1e-05 | 1e-05 |
| DBG-7A | 1e-05 | 0.99994 | 1e-05 | 1e-05 | 1e-05 | 1e-05 | 1e-05 |
| DBG-8A | 1e-05 | 0.99994 | 1e-05 | 1e-05 | 1e-05 | 1e-05 | 1e-05 |
| DBG-9A | 1e-05 | 0.932067 | 1e-05 | 0.056776 | 0.011117 | 1e-05 | 1e-05 |
| TEG-10A | 1e-05 | 0.2809 | 1e-05 | 0.71905 | 1e-05 | 1e-05 | 1e-05 |
| TEG-1A | 1e-05 | 0.354138 | 0.02738 | 0.546337 | 1e-05 | 1e-05 | 0.072115 |
| TEG-2A | 1e-05 | 0.245081 | 1e-05 | 0.731985 | 1e-05 | 1e-05 | 0.022894 |
| TEG-3A | 1e-05 | 0.369255 | 1e-05 | 0.630695 | 1e-05 | 1e-05 | 1e-05 |
| TEG-4A | 1e-05 | 0.410061 | 0.023864 | 0.566035 | 1e-05 | 1e-05 | 1e-05 |
| TEG-5A | 1e-05 | 0.397364 | 1e-05 | 0.602586 | 1e-05 | 1e-05 | 1e-05 |
| TEG-6A | 0.060005 | 0.29162 | 0.016905 | 0.617249 | 1e-05 | 0.0142 | 1e-05 |
| TEG-7A | 0.073926 | 0.28095 | 1e-05 | 0.58432 | 1e-05 | 1e-05 | 0.060774 |
| TEG-8A | 0.051324 | 0.344325 | 1e-05 | 0.604311 | 1e-05 | 1e-05 | 1e-05 |
| TEG-9A | 1e-05 | 0.328714 | 1e-05 | 0.648636 | 1e-05 | 1e-05 | 0.02261 |
| ZPZ-10A | 1e-05 | 1e-05 | 1e-05 | 0.99994 | 1e-05 | 1e-05 | 1e-05 |
| ZPZ-1A | 1e-05 | 1e-05 | 1e-05 | 0.99994 | 1e-05 | 1e-05 | 1e-05 |
| ZPZ-2A | 1e-05 | 1e-05 | 1e-05 | 0.99994 | 1e-05 | 1e-05 | 1e-05 |
| ZPZ-3A | 0.023484 | 1e-05 | 1e-05 | 0.976466 | 1e-05 | 1e-05 | 1e-05 |
| ZPZ-4A | 1e-05 | 1e-05 | 1e-05 | 0.99994 | 1e-05 | 1e-05 | 1e-05 |
| ZPZ-5A | 1e-05 | 1e-05 | 1e-05 | 0.99994 | 1e-05 | 1e-05 | 1e-05 |
| ZPZ-6A | 0.111499 | 0.091435 | 0.023576 | 0.752391 | 1e-05 | 1e-05 | 0.021078 |
| ZPZ-7A | 0.006319 | 0.134772 | 1e-05 | 0.858869 | 1e-05 | 1e-05 | 1e-05 |
| ZPZ-8A | 1e-05 | 1e-05 | 1e-05 | 0.99994 | 1e-05 | 1e-05 | 1e-05 |
| ZPZ-9A | 1e-05 | 1e-05 | 1e-05 | 0.99994 | 1e-05 | 1e-05 | 1e-05 |
| YLJ1-1A | 1e-05 | 1e-05 | 1e-05 | 0.999938 | 1e-05 | 1.2e-05 | 1e-05 |
| YLJ1-2A | 1e-05 | 1.6e-05 | 1.1e-05 | 0.999932 | 1.1e-05 | 1e-05 | 1e-05 |
| YLJ1-3A | 1e-05 | 1e-05 | 1e-05 | 0.99994 | 1e-05 | 1e-05 | 1e-05 |
| YLJ1-4A | 1e-05 | 1e-05 | 1e-05 | 0.99994 | 1e-05 | 1e-05 | 1e-05 |
| YLJ1-5A | 1e-05 | 1e-05 | 1e-05 | 0.99994 | 1e-05 | 1e-05 | 1e-05 |
| YLJ1-6A | 1e-05 | 1e-05 | 1e-05 | 0.977558 | 0.022392 | 1e-05 | 1e-05 |
| YLJ1-7A | 1e-05 | 1.2e-05 | 1e-05 | 0.999938 | 1e-05 | 1e-05 | 1e-05 |
| YLJ1-8A | 0.001126 | 1e-05 | 1e-05 | 0.998824 | 1e-05 | 1e-05 | 1e-05 |
| YLJ2-1A | 1e-05 | 1e-05 | 1e-05 | 0.99994 | 1e-05 | 1e-05 | 1e-05 |
| YLJ2-2A | 1e-05 | 1e-05 | 1e-05 | 0.99994 | 1e-05 | 1e-05 | 1e-05 |
| YLJ2-3A | 1e-05 | 1e-05 | 1e-05 | 0.99994 | 1e-05 | 1e-05 | 1e-05 |
| YLJ2-4A | 1e-05 | 1e-05 | 1.2e-05 | 0.999938 | 1e-05 | 1e-05 | 1e-05 |
| YLJ2-5A | 1e-05 | 1e-05 | 1e-05 | 0.99994 | 1e-05 | 1e-05 | 1e-05 |
| YLJ2-6B | 1e-05 | 1e-05 | 1e-05 | 0.99994 | 1e-05 | 1e-05 | 1e-05 |
| YLJ2-7A | 1e-05 | 1.5e-05 | 1e-05 | 0.999935 | 1e-05 | 1e-05 | 1e-05 |
| YLJ2-8A | 1e-05 | 1e-05 | 1e-05 | 0.99994 | 1e-05 | 1e-05 | 1e-05 |
| LM-10A | 0.99994 | 1e-05 | 1e-05 | 1e-05 | 1e-05 | 1e-05 | 1e-05 |
| LM-1A | 0.99994 | 1e-05 | 1e-05 | 1e-05 | 1e-05 | 1e-05 | 1e-05 |
| LM-2A | 0.999922 | 1e-05 | 1.9e-05 | 1.6e-05 | 1e-05 | 1.2e-05 | 1e-05 |
| LM-3A | 0.99994 | 1e-05 | 1e-05 | 1e-05 | 1e-05 | 1e-05 | 1e-05 |
| LM-4A | 0.999929 | 1.7e-05 | 1.4e-05 | 1e-05 | 1e-05 | 1e-05 | 1e-05 |
| LM-5A | 0.999931 | 1.7e-05 | 1.2e-05 | 1e-05 | 1e-05 | 1e-05 | 1e-05 |
| LM-6A | 0.99994 | 1e-05 | 1e-05 | 1e-05 | 1e-05 | 1e-05 | 1e-05 |
| LM-7A | 0.99994 | 1e-05 | 1e-05 | 1e-05 | 1e-05 | 1e-05 | 1e-05 |
| LM-8A | 0.99994 | 1e-05 | 1e-05 | 1e-05 | 1e-05 | 1e-05 | 1e-05 |
| LM-9A | 0.999935 | 1e-05 | 1e-05 | 1e-05 | 1.5e-05 | 1e-05 | 1e-05 |
| LM-11A | 0.99994 | 1e-05 | 1e-05 | 1e-05 | 1e-05 | 1e-05 | 1e-05 |
| LM-12A | 0.998509 | 1e-05 | 1.7e-05 | 0.001434 | 1e-05 | 1e-05 | 1e-05 |
| LM-13A | 0.99994 | 1e-05 | 1e-05 | 1e-05 | 1e-05 | 1e-05 | 1e-05 |
| LM-14A | 0.999939 | 1.1e-05 | 1e-05 | 1e-05 | 1e-05 | 1e-05 | 1e-05 |
| LM-15A | 0.99994 | 1e-05 | 1e-05 | 1e-05 | 1e-05 | 1e-05 | 1e-05 |
| BYP1-1A | 0.107885 | 1.6e-05 | 0.558584 | 0.333485 | 1e-05 | 1e-05 | 1e-05 |
| BYP1-2A | 0.116381 | 1e-05 | 0.661676 | 0.221903 | 1e-05 | 1e-05 | 1e-05 |
| BYP1-3A | 0.068559 | 1e-05 | 0.605111 | 0.32629 | 1e-05 | 1e-05 | 1e-05 |
| BYP1-4A | 0.019839 | 1e-05 | 0.687165 | 0.292956 | 1e-05 | 1e-05 | 1e-05 |
| BYP1-5A | 0.048141 | 1e-05 | 0.568017 | 0.383803 | 1e-05 | 1e-05 | 1e-05 |
| BYP1-6A | 0.032336 | 1.4e-05 | 0.836343 | 0.131277 | 1e-05 | 1e-05 | 1e-05 |
| BYP1-7A | 0.130422 | 1e-05 | 0.689244 | 0.180286 | 1.8e-05 | 1e-05 | 1e-05 |
| BYP1-8A | 0.029551 | 1e-05 | 0.836073 | 0.134335 | 1e-05 | 1e-05 | 1e-05 |
| BYP1-9A | 0.056712 | 1.4e-05 | 0.782072 | 0.161171 | 1e-05 | 1.1e-05 | 1e-05 |
| BYP1-10A | 0.141121 | 1e-05 | 0.733288 | 0.12371 | 1e-05 | 0.001851 | 1e-05 |
| BYP2-1A | 1e-05 | 1e-05 | 0.454394 | 0.545556 | 1e-05 | 1e-05 | 1e-05 |
| BYP2-2A | 0.063071 | 1e-05 | 0.411476 | 0.525414 | 1e-05 | 1e-05 | 1e-05 |
| BYP2-3A | 0.13061 | 1e-05 | 0.719586 | 0.149764 | 1e-05 | 1e-05 | 1e-05 |
| BYP2-4A | 0.000758 | 1e-05 | 0.402677 | 0.596525 | 1e-05 | 1e-05 | 1e-05 |
| BYP2-5A | 0.066932 | 1e-05 | 0.660019 | 0.273009 | 1e-05 | 1e-05 | 1e-05 |
| BYP2-6A | 0.170325 | 1e-05 | 0.53943 | 0.290205 | 1e-05 | 1e-05 | 1e-05 |
| BYP2-7A | 0.414867 | 1e-05 | 0.36292 | 0.222173 | 1e-05 | 1e-05 | 1e-05 |
| BYP2-8A | 0.040126 | 1e-05 | 0.488915 | 0.470912 | 1e-05 | 1.7e-05 | 1e-05 |
| BYP3-1A | 0.152548 | 1e-05 | 0.626341 | 0.221071 | 1e-05 | 1e-05 | 1e-05 |
| BYP3-2A | 0.066738 | 1e-05 | 0.716233 | 0.216989 | 1e-05 | 1e-05 | 1e-05 |
| BYP3-3A | 0.245412 | 1.5e-05 | 0.52827 | 0.226273 | 1e-05 | 1e-05 | 1e-05 |
| BYP3-4A | 0.185965 | 1e-05 | 0.553534 | 0.26046 | 1e-05 | 1e-05 | 1e-05 |
| BYP3-5A | 0.143376 | 1e-05 | 0.632045 | 0.224538 | 1e-05 | 1e-05 | 1e-05 |
| BYP3-6A | 0.174324 | 1e-05 | 0.512786 | 0.31285 | 1e-05 | 1e-05 | 1e-05 |
| BYP3-7A | 0.108312 | 1e-05 | 0.678462 | 0.213186 | 1e-05 | 1e-05 | 1e-05 |
| BYP3-8A | 0.089192 | 1e-05 | 0.653203 | 0.257565 | 1e-05 | 1e-05 | 1e-05 |
| KBG-1A | 1e-05 | 1e-05 | 0.99994 | 1e-05 | 1e-05 | 1e-05 | 1e-05 |
| KBG-2A | 1e-05 | 1e-05 | 0.919166 | 0.080784 | 1e-05 | 1e-05 | 1e-05 |
| KBG-3A | 1e-05 | 1e-05 | 0.999936 | 1.4e-05 | 1e-05 | 1e-05 | 1e-05 |
| KBG-4A | 1e-05 | 1e-05 | 0.999939 | 1.1e-05 | 1e-05 | 1e-05 | 1e-05 |
| KBG-5A | 1e-05 | 1e-05 | 0.980175 | 1e-05 | 1e-05 | 1e-05 | 0.019775 |
| QJG-1A | 1e-05 | 1e-05 | 0.995362 | 1e-05 | 1e-05 | 1e-05 | 0.004588 |
| QJG-2A | 1e-05 | 1e-05 | 0.99994 | 1e-05 | 1e-05 | 1e-05 | 1e-05 |
| QJG-3A | 1e-05 | 1e-05 | 0.99994 | 1e-05 | 1e-05 | 1e-05 | 1e-05 |
| QJG-4A | 1e-05 | 1e-05 | 0.99994 | 1e-05 | 1e-05 | 1e-05 | 1e-05 |
| QJG-5A | 1e-05 | 0.027539 | 0.88201 | 1e-05 | 1e-05 | 1e-05 | 0.090411 |
| LTG-1A | 0.017669 | 0.177458 | 0.804833 | 1e-05 | 1e-05 | 1e-05 | 1e-05 |
| LTG-2A | 1e-05 | 0.036025 | 0.871536 | 0.04624 | 0.038906 | 1e-05 | 0.007271 |
| LTG-3A | 1e-05 | 1e-05 | 0.99994 | 1e-05 | 1e-05 | 1e-05 | 1e-05 |
| LTG-4A | 1e-05 | 1e-05 | 0.991481 | 1e-05 | 1e-05 | 1e-05 | 0.008469 |
| LTG-5A | 0.017684 | 1e-05 | 0.909371 | 1e-05 | 0.072904 | 1e-05 | 1e-05 |
| BW-10B | 1e-05 | 1e-05 | 0.990784 | 0.009166 | 1e-05 | 1e-05 | 1e-05 |
| BW-1B | 0.013123 | 1.2e-05 | 0.986825 | 1e-05 | 1e-05 | 1e-05 | 1e-05 |
| BW-2B | 1.3e-05 | 1e-05 | 0.999937 | 1e-05 | 1e-05 | 1e-05 | 1e-05 |
| BW-3B | 1.4e-05 | 1e-05 | 0.999936 | 1e-05 | 1e-05 | 1e-05 | 1e-05 |
| BW-4B | 1e-05 | 1e-05 | 0.99994 | 1e-05 | 1e-05 | 1e-05 | 1e-05 |
| BW-5B | 1e-05 | 1e-05 | 0.99994 | 1e-05 | 1e-05 | 1e-05 | 1e-05 |
| BW-6B | 1e-05 | 1e-05 | 0.99994 | 1e-05 | 1e-05 | 1e-05 | 1e-05 |
| BW-7A | 1e-05 | 1e-05 | 0.99994 | 1e-05 | 1e-05 | 1e-05 | 1e-05 |
| BW-8B | 0.010282 | 1e-05 | 0.989668 | 1e-05 | 1e-05 | 1e-05 | 1e-05 |
| BW-9A | 1.1e-05 | 1e-05 | 0.999939 | 1e-05 | 1e-05 | 1e-05 | 1e-05 |
| GA-1A | 1e-05 | 1e-05 | 0.931772 | 1e-05 | 1e-05 | 1e-05 | 0.068178 |
| GA-2A | 0.027511 | 1e-05 | 0.958401 | 1e-05 | 1e-05 | 1e-05 | 0.014048 |
| GA-3A | 0.052829 | 1e-05 | 0.947119 | 1e-05 | 1.2e-05 | 1e-05 | 1e-05 |
| MR-1A | 1e-05 | 1e-05 | 0.951265 | 1e-05 | 1e-05 | 1e-05 | 0.048685 |
| MR-2A | 1e-05 | 1e-05 | 0.959875 | 1e-05 | 1e-05 | 0.003174 | 0.036911 |
| MR-3A | 0.081147 | 0.015049 | 0.817967 | 1e-05 | 0.018097 | 1e-05 | 0.067721 |
| MR-4A | 1e-05 | 0.00978 | 0.945851 | 1e-05 | 1e-05 | 1e-05 | 0.044329 |
| MR-5A | 1e-05 | 0.029657 | 0.958006 | 1e-05 | 1e-05 | 0.012297 | 1e-05 |
| MDG-6A | 1e-05 | 0.006935 | 0.738237 | 1e-05 | 1e-05 | 1e-05 | 0.254789 |
| MDG-1A | 1e-05 | 1e-05 | 0.861231 | 1e-05 | 1e-05 | 1e-05 | 0.138719 |
| MDG-2A | 1e-05 | 1.6e-05 | 0.841097 | 1e-05 | 1e-05 | 1e-05 | 0.158847 |
| MDG-3A | 0.008133 | 1e-05 | 0.766113 | 1e-05 | 1e-05 | 1e-05 | 0.225714 |
| MDG-4A | 1e-05 | 0.03495 | 0.724445 | 1e-05 | 1e-05 | 1e-05 | 0.240565 |
| MDG-5A | 1e-05 | 1e-05 | 0.785783 | 1e-05 | 1e-05 | 1e-05 | 0.214167 |
| AJ-10A | 1e-05 | 1e-05 | 0.261341 | 1e-05 | 1e-05 | 1e-05 | 0.738609 |
| AJ-12A | 1e-05 | 1e-05 | 0.225249 | 1e-05 | 1e-05 | 1e-05 | 0.774701 |
| AJ-13A | 1e-05 | 1e-05 | 0.26709 | 1e-05 | 1e-05 | 1e-05 | 0.73286 |
| AJ-14A | 1e-05 | 1e-05 | 0.318478 | 1e-05 | 1e-05 | 1e-05 | 0.681472 |
| AJ-1A | 1e-05 | 1e-05 | 0.294091 | 1e-05 | 1e-05 | 1e-05 | 0.705859 |
| AJ-3A | 1e-05 | 1e-05 | 0.304407 | 0.012832 | 1e-05 | 1e-05 | 0.682721 |
| AJ-4A | 1e-05 | 0.001883 | 0.171834 | 1e-05 | 1e-05 | 0.008954 | 0.817299 |
| AJ-5A | 1e-05 | 1e-05 | 0.254275 | 1e-05 | 1e-05 | 1e-05 | 0.745675 |
| AJ-7A | 1e-05 | 0.045081 | 0.290809 | 0.004679 | 1e-05 | 1e-05 | 0.659401 |
| AJ-9A | 1e-05 | 1.1e-05 | 0.150321 | 1.1e-05 | 1.5e-05 | 1e-05 | 0.849622 |
| XGG-1A | 1e-05 | 1e-05 | 1.1e-05 | 1e-05 | 1e-05 | 1e-05 | 0.999939 |
| XGG-2A | 1e-05 | 1e-05 | 1e-05 | 1e-05 | 1e-05 | 1e-05 | 0.99994 |
| XGG-3A | 1e-05 | 1e-05 | 1e-05 | 1e-05 | 1e-05 | 1e-05 | 0.99994 |
| XGG-4A | 1e-05 | 1e-05 | 1e-05 | 1e-05 | 1e-05 | 1e-05 | 0.99994 |
| XGG-5A | 1e-05 | 1e-05 | 0.037344 | 1e-05 | 1e-05 | 1e-05 | 0.962606 |
| PSR-01A | 1e-05 | 1e-05 | 1e-05 | 1e-05 | 1e-05 | 1e-05 | 0.99994 |
| PSR-04A | 1e-05 | 1e-05 | 1e-05 | 1e-05 | 1e-05 | 1e-05 | 0.99994 |
| PSR-10A | 1e-05 | 1e-05 | 1e-05 | 1e-05 | 1e-05 | 1e-05 | 0.99994 |
| PSR-13A | 1e-05 | 1e-05 | 0.057078 | 1e-05 | 1e-05 | 1e-05 | 0.942872 |
| LR-10A | 1e-05 | 1e-05 | 1e-05 | 1e-05 | 1e-05 | 1e-05 | 0.99994 |
| LR-1A | 1e-05 | 1e-05 | 0.017823 | 1e-05 | 1e-05 | 1e-05 | 0.982127 |
| LR-2A | 1e-05 | 1e-05 | 1e-05 | 1e-05 | 1e-05 | 1e-05 | 0.99994 |
| LR-3A | 1e-05 | 1e-05 | 1e-05 | 1e-05 | 1e-05 | 1e-05 | 0.99994 |
| LR-4A | 1e-05 | 1e-05 | 1e-05 | 1e-05 | 1e-05 | 1e-05 | 0.99994 |
| LR-5A | 0.002006 | 1e-05 | 1e-05 | 1e-05 | 1e-05 | 1e-05 | 0.997944 |
| LR-6A | 1e-05 | 0.014728 | 1e-05 | 0.006948 | 1e-05 | 1e-05 | 0.978284 |
| LR-7A | 1e-05 | 1e-05 | 1e-05 | 1e-05 | 1e-05 | 1e-05 | 0.99994 |
| LR-8A | 1e-05 | 1e-05 | 1e-05 | 1e-05 | 1e-05 | 1e-05 | 0.99994 |
| LR-9A | 1e-05 | 1e-05 | 1e-05 | 1e-05 | 1e-05 | 1e-05 | 0.99994 |
| BYR-10A | 1e-05 | 1e-05 | 1e-05 | 1e-05 | 1e-05 | 1e-05 | 0.99994 |
| BYR-1A | 1e-05 | 1e-05 | 1e-05 | 1e-05 | 1e-05 | 1e-05 | 0.99994 |
| BYR-2A | 1e-05 | 1e-05 | 1e-05 | 1e-05 | 1e-05 | 1e-05 | 0.99994 |
| BYR-3A | 1e-05 | 1e-05 | 1e-05 | 1e-05 | 1e-05 | 1e-05 | 0.99994 |
| BYR-4A | 1e-05 | 1e-05 | 1.2e-05 | 1e-05 | 1e-05 | 1e-05 | 0.999938 |
| BYR-5A | 1e-05 | 1e-05 | 1e-05 | 1e-05 | 1e-05 | 1e-05 | 0.99994 |
| BYR-6A | 1e-05 | 1e-05 | 1e-05 | 1e-05 | 1e-05 | 1e-05 | 0.99994 |
| BYR-7A | 1e-05 | 0.028081 | 1e-05 | 1e-05 | 1e-05 | 1e-05 | 0.971869 |

K = 8

|  |  |  |  |  |  |  |  |  |
| --- | --- | --- | --- | --- | --- | --- | --- | --- |
| CDG-1A | 1e-05 | 1e-05 | 0.99993 | 1e-05 | 1e-05 | 1e-05 | 1e-05 | 1e-05 |
| CDG-2A | 1e-05 | 1e-05 | 0.99993 | 1e-05 | 1e-05 | 1e-05 | 1e-05 | 1e-05 |
| CDG-3A | 1e-05 | 1.3e-05 | 0.999927 | 1e-05 | 1e-05 | 1e-05 | 1e-05 | 1e-05 |
| CDG-4A | 1e-05 | 1e-05 | 0.99993 | 1e-05 | 1e-05 | 1e-05 | 1e-05 | 1e-05 |
| CDG-5A | 1e-05 | 1e-05 | 0.999927 | 1.3e-05 | 1e-05 | 1e-05 | 1e-05 | 1e-05 |
| CDG-6A | 1e-05 | 1e-05 | 0.99993 | 1e-05 | 1e-05 | 1e-05 | 1e-05 | 1e-05 |
| CDG-7A | 1e-05 | 1e-05 | 0.99993 | 1e-05 | 1e-05 | 1e-05 | 1e-05 | 1e-05 |
| CDG-8A | 1e-05 | 1e-05 | 0.99993 | 1e-05 | 1e-05 | 1e-05 | 1e-05 | 1e-05 |
| CDG-9A | 1e-05 | 1e-05 | 0.99993 | 1e-05 | 1e-05 | 1e-05 | 1e-05 | 1e-05 |
| CDG-10A | 1e-05 | 1e-05 | 0.99993 | 1e-05 | 1e-05 | 1e-05 | 1e-05 | 1e-05 |
| TKX-1A | 1e-05 | 1e-05 | 1.2e-05 | 1e-05 | 0.999928 | 1e-05 | 1e-05 | 1e-05 |
| TKX-2A | 1e-05 | 1.1e-05 | 1e-05 | 1e-05 | 0.999929 | 1e-05 | 1e-05 | 1e-05 |
| TKX-3A | 1.5e-05 | 1e-05 | 1e-05 | 1.6e-05 | 0.999918 | 1e-05 | 1e-05 | 1e-05 |
| TKX-4A | 1e-05 | 1.7e-05 | 1e-05 | 1e-05 | 0.999917 | 1.6e-05 | 1e-05 | 1e-05 |
| TKX-5A | 1e-05 | 1e-05 | 1e-05 | 1e-05 | 0.99993 | 1e-05 | 1e-05 | 1e-05 |
| TKX-6A | 1.1e-05 | 1e-05 | 1e-05 | 1e-05 | 0.99992 | 1.9e-05 | 1e-05 | 1e-05 |
| TKX-7A | 1e-05 | 1e-05 | 1e-05 | 1e-05 | 0.99993 | 1e-05 | 1e-05 | 1e-05 |
| TKX-8A | 1e-05 | 1e-05 | 1e-05 | 1e-05 | 0.999928 | 1e-05 | 1.2e-05 | 1e-05 |
| TKX-9A | 1e-05 | 1e-05 | 1e-05 | 1e-05 | 0.99993 | 1e-05 | 1e-05 | 1e-05 |
| GJ-1A | 1e-05 | 1e-05 | 1e-05 | 1e-05 | 1e-05 | 1e-05 | 0.99993 | 1e-05 |
| GJ-2A | 1e-05 | 1e-05 | 1e-05 | 1e-05 | 1e-05 | 1e-05 | 0.99993 | 1e-05 |
| GJ-3A | 1e-05 | 1e-05 | 1.7e-05 | 1e-05 | 1e-05 | 1e-05 | 0.999923 | 1e-05 |
| GJ-4A | 1.1e-05 | 1.1e-05 | 1.3e-05 | 1e-05 | 1e-05 | 1e-05 | 0.999925 | 1e-05 |
| GJ-5A | 1e-05 | 1e-05 | 1e-05 | 1e-05 | 1e-05 | 1e-05 | 0.99993 | 1e-05 |
| GJ-6A | 1e-05 | 1e-05 | 1e-05 | 1e-05 | 1e-05 | 1e-05 | 0.99993 | 1e-05 |
| GJ-7A | 1e-05 | 1.7e-05 | 1e-05 | 1e-05 | 1e-05 | 1e-05 | 0.999923 | 1e-05 |
| GJ-8A | 1e-05 | 1e-05 | 1e-05 | 1e-05 | 1e-05 | 1e-05 | 0.99993 | 1e-05 |
| GJ-9A | 1e-05 | 1.8e-05 | 1e-05 | 1e-05 | 1e-05 | 1e-05 | 0.999922 | 1e-05 |
| TZC-1A | 1e-05 | 1e-05 | 1e-05 | 1e-05 | 1e-05 | 1e-05 | 0.99993 | 1e-05 |
| TZC-2A | 1e-05 | 1e-05 | 1e-05 | 1e-05 | 1e-05 | 1e-05 | 0.99993 | 1e-05 |
| TZC-3A | 1e-05 | 1e-05 | 1e-05 | 0.098179 | 1e-05 | 1e-05 | 0.852587 | 0.049184 |
| TZC-4A | 1e-05 | 1e-05 | 1e-05 | 1e-05 | 1e-05 | 1e-05 | 0.99993 | 1e-05 |
| TZC-01A | 1e-05 | 1e-05 | 1e-05 | 1e-05 | 0.000114 | 1e-05 | 0.999826 | 1e-05 |
| TZC-04A | 1e-05 | 1e-05 | 1e-05 | 1e-05 | 1e-05 | 1e-05 | 0.99993 | 1e-05 |
| TZC-07A | 1e-05 | 1e-05 | 1e-05 | 1e-05 | 1e-05 | 1e-05 | 0.99993 | 1e-05 |
| TZC-11A | 1e-05 | 1e-05 | 1e-05 | 1e-05 | 1e-05 | 1e-05 | 0.99993 | 1e-05 |
| TZC-13A | 1.3e-05 | 1e-05 | 1e-05 | 1e-05 | 1e-05 | 1e-05 | 0.999927 | 1e-05 |
| DBG-10A | 1e-05 | 1e-05 | 1e-05 | 1e-05 | 1e-05 | 1e-05 | 0.99993 | 1e-05 |
| DBG-1A | 1e-05 | 1e-05 | 1e-05 | 1e-05 | 1e-05 | 0.052175 | 0.831363 | 0.116412 |
| DBG-2A | 0.002103 | 1e-05 | 1e-05 | 1e-05 | 1e-05 | 1e-05 | 0.997837 | 1e-05 |
| DBG-3A | 0.00674 | 1e-05 | 1e-05 | 1e-05 | 1e-05 | 1e-05 | 0.9932 | 1e-05 |
| DBG-4A | 1e-05 | 1e-05 | 1e-05 | 1e-05 | 1e-05 | 1e-05 | 0.99993 | 1e-05 |
| DBG-5A | 1e-05 | 1e-05 | 1e-05 | 1e-05 | 1e-05 | 1e-05 | 0.87571 | 0.12423 |
| DBG-6A | 1e-05 | 1e-05 | 1e-05 | 1e-05 | 0.00047 | 1e-05 | 0.99947 | 1e-05 |
| DBG-7A | 1e-05 | 1e-05 | 1e-05 | 1e-05 | 1e-05 | 1e-05 | 0.99993 | 1e-05 |
| DBG-8A | 1e-05 | 1e-05 | 1e-05 | 1e-05 | 1e-05 | 1e-05 | 0.99993 | 1e-05 |
| DBG-9A | 0.002946 | 1e-05 | 1e-05 | 1e-05 | 0.014366 | 1e-05 | 0.949403 | 0.033246 |
| TEG-10A | 1e-05 | 1e-05 | 1e-05 | 1e-05 | 1e-05 | 1e-05 | 0.273137 | 0.726803 |
| TEG-1A | 0.00113 | 0.026047 | 1e-05 | 0.00669 | 1e-05 | 0.057462 | 0.369143 | 0.539508 |
| TEG-2A | 0.063379 | 0.029419 | 1e-05 | 1e-05 | 1e-05 | 1e-05 | 0.236252 | 0.67091 |
| TEG-3A | 0.005239 | 1e-05 | 1e-05 | 1e-05 | 1e-05 | 1e-05 | 0.37781 | 0.616901 |
| TEG-4A | 1e-05 | 1e-05 | 1e-05 | 0.028011 | 1e-05 | 1e-05 | 0.413504 | 0.558435 |
| TEG-5A | 1e-05 | 1e-05 | 1e-05 | 1e-05 | 1e-05 | 1e-05 | 0.40173 | 0.59821 |
| TEG-6A | 0.111152 | 1e-05 | 0.013841 | 1e-05 | 1e-05 | 1e-05 | 0.288009 | 0.586958 |
| TEG-7A | 0.120786 | 1e-05 | 1e-05 | 1e-05 | 1e-05 | 0.047289 | 0.279788 | 0.552096 |
| TEG-8A | 0.055304 | 1e-05 | 1e-05 | 1e-05 | 1e-05 | 1e-05 | 0.350961 | 0.593686 |
| TEG-9A | 0.002732 | 0.036227 | 1e-05 | 1e-05 | 1e-05 | 1e-05 | 0.328408 | 0.632593 |
| ZPZ-10A | 1e-05 | 1.1e-05 | 1e-05 | 1e-05 | 1e-05 | 1e-05 | 1e-05 | 0.999929 |
| ZPZ-1A | 1e-05 | 1e-05 | 1.8e-05 | 1e-05 | 1e-05 | 1e-05 | 1e-05 | 0.999922 |
| ZPZ-2A | 1e-05 | 1e-05 | 1e-05 | 1e-05 | 1e-05 | 1e-05 | 1e-05 | 0.99993 |
| ZPZ-3A | 0.05803 | 1e-05 | 1e-05 | 1e-05 | 1e-05 | 1e-05 | 1e-05 | 0.94191 |
| ZPZ-4A | 1e-05 | 1e-05 | 1e-05 | 1e-05 | 1e-05 | 1e-05 | 1e-05 | 0.99993 |
| ZPZ-5A | 1e-05 | 1.1e-05 | 1e-05 | 1.4e-05 | 1e-05 | 1e-05 | 1e-05 | 0.999925 |
| ZPZ-6A | 0.231973 | 0.023367 | 1.1e-05 | 1e-05 | 1e-05 | 1e-05 | 0.057389 | 0.687229 |
| ZPZ-7A | 0.040628 | 1e-05 | 1.1e-05 | 1e-05 | 1e-05 | 1e-05 | 0.121535 | 0.837786 |
| ZPZ-8A | 1.1e-05 | 1e-05 | 1e-05 | 1.6e-05 | 1e-05 | 1e-05 | 1e-05 | 0.999923 |
| ZPZ-9A | 1e-05 | 1e-05 | 1.8e-05 | 1e-05 | 1e-05 | 1e-05 | 1e-05 | 0.999922 |
| YLJ1-1A | 1e-05 | 1e-05 | 1e-05 | 1e-05 | 1e-05 | 1e-05 | 1e-05 | 0.99993 |
| YLJ1-2A | 1e-05 | 1.8e-05 | 1.7e-05 | 1.2e-05 | 1e-05 | 1e-05 | 1e-05 | 0.999912 |
| YLJ1-3A | 1e-05 | 1e-05 | 1e-05 | 1e-05 | 1e-05 | 1e-05 | 1e-05 | 0.99993 |
| YLJ1-4A | 1.3e-05 | 1.4e-05 | 1e-05 | 1.2e-05 | 1e-05 | 1e-05 | 1e-05 | 0.999922 |
| YLJ1-5A | 1e-05 | 1e-05 | 1.2e-05 | 1e-05 | 1.1e-05 | 1e-05 | 1e-05 | 0.999927 |
| YLJ1-6A | 1e-05 | 1e-05 | 1e-05 | 1e-05 | 0.015432 | 1e-05 | 1e-05 | 0.984508 |
| YLJ1-7A | 1e-05 | 1.5e-05 | 1.1e-05 | 1e-05 | 1.2e-05 | 1e-05 | 1e-05 | 0.999922 |
| YLJ1-8A | 0.0466 | 1e-05 | 1e-05 | 1e-05 | 1e-05 | 1e-05 | 1e-05 | 0.95334 |
| YLJ2-1A | 1e-05 | 1e-05 | 1e-05 | 1e-05 | 1e-05 | 1e-05 | 1e-05 | 0.99993 |
| YLJ2-2A | 1e-05 | 1e-05 | 1e-05 | 1e-05 | 1e-05 | 1e-05 | 1e-05 | 0.99993 |
| YLJ2-3A | 1e-05 | 1.5e-05 | 1e-05 | 1e-05 | 1e-05 | 1e-05 | 1e-05 | 0.999925 |
| YLJ2-4A | 1e-05 | 1e-05 | 1e-05 | 1e-05 | 1e-05 | 1e-05 | 1e-05 | 0.99993 |
| YLJ2-5A | 1e-05 | 1e-05 | 1e-05 | 1e-05 | 1e-05 | 1e-05 | 1e-05 | 0.99993 |
| YLJ2-6B | 1e-05 | 1e-05 | 1e-05 | 1.2e-05 | 1e-05 | 1e-05 | 1e-05 | 0.999928 |
| YLJ2-7A | 1e-05 | 1e-05 | 1e-05 | 1e-05 | 1e-05 | 1e-05 | 1e-05 | 0.99993 |
| YLJ2-8A | 0.041334 | 1e-05 | 1e-05 | 1e-05 | 1e-05 | 1e-05 | 1e-05 | 0.958606 |
| LM-10A | 0.99993 | 1e-05 | 1e-05 | 1e-05 | 1e-05 | 1e-05 | 1e-05 | 1e-05 |
| LM-1A | 0.99993 | 1e-05 | 1e-05 | 1e-05 | 1e-05 | 1e-05 | 1e-05 | 1e-05 |
| LM-2A | 0.99993 | 1e-05 | 1e-05 | 1e-05 | 1e-05 | 1e-05 | 1e-05 | 1e-05 |
| LM-3A | 0.99993 | 1e-05 | 1e-05 | 1e-05 | 1e-05 | 1e-05 | 1e-05 | 1e-05 |
| LM-4A | 0.99993 | 1e-05 | 1e-05 | 1e-05 | 1e-05 | 1e-05 | 1e-05 | 1e-05 |
| LM-5A | 0.99993 | 1e-05 | 1e-05 | 1e-05 | 1e-05 | 1e-05 | 1e-05 | 1e-05 |
| LM-6A | 0.99993 | 1e-05 | 1e-05 | 1e-05 | 1e-05 | 1e-05 | 1e-05 | 1e-05 |
| LM-7A | 0.99993 | 1e-05 | 1e-05 | 1e-05 | 1e-05 | 1e-05 | 1e-05 | 1e-05 |
| LM-8A | 0.99993 | 1e-05 | 1e-05 | 1e-05 | 1e-05 | 1e-05 | 1e-05 | 1e-05 |
| LM-9A | 0.99993 | 1e-05 | 1e-05 | 1e-05 | 1e-05 | 1e-05 | 1e-05 | 1e-05 |
| LM-11A | 0.99993 | 1e-05 | 1e-05 | 1e-05 | 1e-05 | 1e-05 | 1e-05 | 1e-05 |
| LM-12A | 0.999929 | 1e-05 | 1e-05 | 1e-05 | 1e-05 | 1e-05 | 1.1e-05 | 1e-05 |
| LM-13A | 0.99993 | 1e-05 | 1e-05 | 1e-05 | 1e-05 | 1e-05 | 1e-05 | 1e-05 |
| LM-14A | 0.99993 | 1e-05 | 1e-05 | 1e-05 | 1e-05 | 1e-05 | 1e-05 | 1e-05 |
| LM-15A | 0.99993 | 1e-05 | 1e-05 | 1e-05 | 1e-05 | 1e-05 | 1e-05 | 1e-05 |
| BYP1-1A | 0.731433 | 1e-05 | 1e-05 | 0.069569 | 1e-05 | 1e-05 | 1e-05 | 0.198948 |
| BYP1-2A | 0.786527 | 1e-05 | 1e-05 | 0.141848 | 1e-05 | 1e-05 | 1e-05 | 0.071574 |
| BYP1-3A | 0.747597 | 1e-05 | 1e-05 | 0.070779 | 1e-05 | 1e-05 | 1e-05 | 0.181574 |
| BYP1-4A | 0.730869 | 1e-05 | 1e-05 | 0.109963 | 1e-05 | 1e-05 | 1e-05 | 0.159117 |
| BYP1-5A | 0.628273 | 1e-05 | 1e-05 | 0.09176 | 1e-05 | 1e-05 | 1e-05 | 0.279917 |
| BYP1-6A | 0.99993 | 1e-05 | 1e-05 | 1e-05 | 1e-05 | 1e-05 | 1e-05 | 1e-05 |
| BYP1-7A | 0.997861 | 1e-05 | 1e-05 | 0.002079 | 1e-05 | 1e-05 | 1e-05 | 1e-05 |
| BYP1-8A | 0.99993 | 1e-05 | 1e-05 | 1e-05 | 1e-05 | 1e-05 | 1e-05 | 1e-05 |
| BYP1-9A | 0.976404 | 1e-05 | 1e-05 | 0.023536 | 1e-05 | 1e-05 | 1e-05 | 1e-05 |
| BYP1-10A | 0.920323 | 0.050596 | 0.029031 | 1e-05 | 1e-05 | 1e-05 | 1e-05 | 1e-05 |
| BYP2-1A | 0.999928 | 1e-05 | 1e-05 | 1.1e-05 | 1e-05 | 1e-05 | 1e-05 | 1e-05 |
| BYP2-2A | 0.94466 | 1e-05 | 1e-05 | 1.4e-05 | 1e-05 | 1e-05 | 0.028761 | 0.026524 |
| BYP2-3A | 0.657408 | 1e-05 | 1e-05 | 0.342532 | 1e-05 | 1e-05 | 1e-05 | 1e-05 |
| BYP2-4A | 0.997897 | 1.8e-05 | 1e-05 | 1e-05 | 1.1e-05 | 1.3e-05 | 1e-05 | 0.002031 |
| BYP2-5A | 0.546337 | 1e-05 | 1e-05 | 0.282732 | 1e-05 | 1e-05 | 0.042339 | 0.128552 |
| BYP2-6A | 0.742362 | 1e-05 | 1e-05 | 0.135614 | 0.009158 | 1e-05 | 1e-05 | 0.112827 |
| BYP2-7A | 0.797573 | 1e-05 | 1e-05 | 0.086799 | 1e-05 | 1e-05 | 1e-05 | 0.115579 |
| BYP2-8A | 0.938569 | 1e-05 | 1e-05 | 1e-05 | 1e-05 | 1e-05 | 1e-05 | 0.061371 |
| BYP3-1A | 0.747541 | 1e-05 | 1e-05 | 0.145541 | 1e-05 | 1e-05 | 0.025007 | 0.081871 |
| BYP3-2A | 0.784068 | 1e-05 | 1e-05 | 0.163352 | 1e-05 | 1e-05 | 1e-05 | 0.05253 |
| BYP3-3A | 0.75788 | 1e-05 | 1e-05 | 0.160201 | 1e-05 | 1e-05 | 0.031423 | 0.050456 |
| BYP3-4A | 0.774483 | 0.031073 | 1e-05 | 0.094481 | 1e-05 | 1e-05 | 1e-05 | 0.099923 |
| BYP3-5A | 0.710208 | 0.030517 | 1e-05 | 0.162681 | 1e-05 | 1e-05 | 1e-05 | 0.096555 |
| BYP3-6A | 0.716792 | 1e-05 | 1e-05 | 0.120785 | 1e-05 | 1e-05 | 1e-05 | 0.162373 |
| BYP3-7A | 0.813732 | 1e-05 | 1e-05 | 0.140328 | 1e-05 | 0.015577 | 1e-05 | 0.030323 |
| BYP3-8A | 0.769322 | 1e-05 | 1e-05 | 0.132395 | 1e-05 | 1e-05 | 1e-05 | 0.098233 |
| KBG-1A | 1e-05 | 1e-05 | 1e-05 | 0.99993 | 1e-05 | 1e-05 | 1e-05 | 1e-05 |
| KBG-2A | 1e-05 | 1e-05 | 1e-05 | 0.925497 | 1e-05 | 1e-05 | 1e-05 | 0.074443 |
| KBG-3A | 1e-05 | 1e-05 | 1e-05 | 0.99993 | 1e-05 | 1e-05 | 1e-05 | 1e-05 |
| KBG-4A | 1e-05 | 1e-05 | 1e-05 | 0.99993 | 1e-05 | 1e-05 | 1e-05 | 1e-05 |
| KBG-5A | 1e-05 | 1e-05 | 1e-05 | 0.99993 | 1e-05 | 1e-05 | 1e-05 | 1e-05 |
| QJG-1A | 1e-05 | 1e-05 | 1e-05 | 0.99993 | 1e-05 | 1e-05 | 1e-05 | 1e-05 |
| QJG-2A | 1e-05 | 1e-05 | 1e-05 | 0.99993 | 1e-05 | 1e-05 | 1e-05 | 1e-05 |
| QJG-3A | 1.8e-05 | 1e-05 | 1e-05 | 0.999922 | 1e-05 | 1e-05 | 1e-05 | 1e-05 |
| QJG-4A | 1e-05 | 1e-05 | 1e-05 | 0.99993 | 1e-05 | 1e-05 | 1e-05 | 1e-05 |
| QJG-5A | 1e-05 | 0.007841 | 1e-05 | 0.992099 | 1e-05 | 1e-05 | 1e-05 | 1e-05 |
| LTG-1A | 0.160583 | 1e-05 | 1e-05 | 0.675845 | 1e-05 | 1e-05 | 0.163523 | 1e-05 |
| LTG-2A | 0.01887 | 1e-05 | 1e-05 | 0.88796 | 0.036084 | 1e-05 | 0.0528 | 0.004256 |
| LTG-3A | 1e-05 | 1e-05 | 1e-05 | 0.99993 | 1e-05 | 1e-05 | 1e-05 | 1e-05 |
| LTG-4A | 1e-05 | 1e-05 | 1e-05 | 0.99993 | 1e-05 | 1e-05 | 1e-05 | 1e-05 |
| LTG-5A | 0.005677 | 1e-05 | 1e-05 | 0.928471 | 0.065801 | 1e-05 | 1e-05 | 1e-05 |
| BW-10B | 1e-05 | 1e-05 | 1e-05 | 0.99993 | 1e-05 | 1e-05 | 1e-05 | 1e-05 |
| BW-1B | 1e-05 | 1e-05 | 1e-05 | 0.99993 | 1e-05 | 1e-05 | 1e-05 | 1e-05 |
| BW-2B | 1.9e-05 | 1e-05 | 1e-05 | 0.999921 | 1e-05 | 1e-05 | 1e-05 | 1e-05 |
| BW-3B | 1e-05 | 1e-05 | 1e-05 | 0.99993 | 1e-05 | 1e-05 | 1e-05 | 1e-05 |
| BW-4B | 0.010138 | 1e-05 | 1e-05 | 0.989802 | 1e-05 | 1e-05 | 1e-05 | 1e-05 |
| BW-5B | 1e-05 | 1e-05 | 1e-05 | 0.99993 | 1e-05 | 1e-05 | 1e-05 | 1e-05 |
| BW-6B | 1e-05 | 1e-05 | 1e-05 | 0.99993 | 1e-05 | 1e-05 | 1e-05 | 1e-05 |
| BW-7A | 1.7e-05 | 1e-05 | 1e-05 | 0.999923 | 1e-05 | 1e-05 | 1e-05 | 1e-05 |
| BW-8B | 1e-05 | 1e-05 | 1e-05 | 0.99993 | 1e-05 | 1e-05 | 1e-05 | 1e-05 |
| BW-9A | 1e-05 | 1e-05 | 1e-05 | 0.99993 | 1e-05 | 1e-05 | 1e-05 | 1e-05 |
| GA-1A | 1e-05 | 1e-05 | 1e-05 | 0.967253 | 1e-05 | 0.032687 | 1e-05 | 1e-05 |
| GA-2A | 0.063787 | 1e-05 | 1e-05 | 0.936153 | 1e-05 | 1e-05 | 1e-05 | 1e-05 |
| GA-3A | 0.131087 | 1e-05 | 1e-05 | 0.868853 | 1e-05 | 1e-05 | 1e-05 | 1e-05 |
| MR-1A | 1e-05 | 1e-05 | 1e-05 | 0.99993 | 1e-05 | 1e-05 | 1e-05 | 1e-05 |
| MR-2A | 1e-05 | 1e-05 | 1e-05 | 0.99993 | 1e-05 | 1e-05 | 1e-05 | 1e-05 |
| MR-3A | 0.0641 | 0.0291 | 1e-05 | 0.89832 | 0.00844 | 1e-05 | 1e-05 | 1e-05 |
| MR-4A | 0.093895 | 0.003919 | 1e-05 | 0.902136 | 1e-05 | 1e-05 | 1e-05 | 1e-05 |
| MR-5A | 1e-05 | 1e-05 | 0.006397 | 0.993543 | 1e-05 | 1e-05 | 1e-05 | 1e-05 |
| MDG-6A | 1e-05 | 1e-05 | 1e-05 | 0.782233 | 1e-05 | 0.217707 | 1e-05 | 1e-05 |
| MDG-1A | 1e-05 | 1e-05 | 1e-05 | 0.882516 | 1e-05 | 0.117424 | 1e-05 | 1e-05 |
| MDG-2A | 1e-05 | 1e-05 | 1e-05 | 0.886865 | 1e-05 | 0.113075 | 1e-05 | 1e-05 |
| MDG-3A | 0.015968 | 1e-05 | 1e-05 | 0.802136 | 1e-05 | 0.181846 | 1e-05 | 1e-05 |
| MDG-4A | 1e-05 | 0.111403 | 1e-05 | 0.76402 | 1e-05 | 0.099495 | 0.025042 | 1e-05 |
| MDG-5A | 1e-05 | 0.067842 | 1e-05 | 0.839604 | 1e-05 | 0.092504 | 1e-05 | 1e-05 |
| AJ-10A | 1e-05 | 1e-05 | 1e-05 | 0.105212 | 1e-05 | 0.86037 | 0.034368 | 1e-05 |
| AJ-12A | 1e-05 | 1e-05 | 1e-05 | 1e-05 | 1e-05 | 0.99993 | 1e-05 | 1e-05 |
| AJ-13A | 1e-05 | 1e-05 | 1e-05 | 1e-05 | 1e-05 | 0.99993 | 1e-05 | 1e-05 |
| AJ-14A | 1e-05 | 1e-05 | 1e-05 | 0.027091 | 1e-05 | 0.972849 | 1e-05 | 1e-05 |
| AJ-1A | 1e-05 | 1e-05 | 1e-05 | 0.202578 | 1e-05 | 0.797362 | 1e-05 | 1e-05 |
| AJ-3A | 0.023348 | 0.145334 | 1e-05 | 0.188016 | 1e-05 | 0.643261 | 1e-05 | 1e-05 |
| AJ-4A | 1e-05 | 0.091095 | 1e-05 | 0.052073 | 1e-05 | 0.817534 | 0.039258 | 1e-05 |
| AJ-5A | 1e-05 | 0.089557 | 1e-05 | 0.166735 | 1e-05 | 0.743659 | 1e-05 | 1e-05 |
| AJ-7A | 1e-05 | 1e-05 | 1e-05 | 0.210934 | 1e-05 | 0.726242 | 0.062773 | 1e-05 |
| AJ-9A | 1e-05 | 1e-05 | 1e-05 | 1e-05 | 1e-05 | 0.99993 | 1e-05 | 1e-05 |
| XGG-1A | 1e-05 | 0.94912 | 1e-05 | 1e-05 | 1e-05 | 0.05082 | 1e-05 | 1e-05 |
| XGG-2A | 1e-05 | 0.799168 | 1e-05 | 1e-05 | 1e-05 | 0.200772 | 1e-05 | 1e-05 |
| XGG-3A | 1e-05 | 0.832061 | 1e-05 | 1e-05 | 1e-05 | 0.167879 | 1e-05 | 1e-05 |
| XGG-4A | 1e-05 | 0.889517 | 1e-05 | 1e-05 | 1e-05 | 0.110423 | 1e-05 | 1e-05 |
| XGG-5A | 1e-05 | 0.770875 | 1e-05 | 0.032729 | 1e-05 | 0.196345 | 1e-05 | 1e-05 |
| PSR-01A | 1.8e-05 | 0.903102 | 1e-05 | 1e-05 | 1e-05 | 0.09683 | 1e-05 | 1e-05 |
| PSR-04A | 1e-05 | 0.92904 | 1e-05 | 1e-05 | 1e-05 | 0.0709 | 1e-05 | 1e-05 |
| PSR-10A | 1e-05 | 0.92131 | 1e-05 | 1e-05 | 1e-05 | 0.07863 | 1e-05 | 1e-05 |
| PSR-13A | 1e-05 | 0.97051 | 1e-05 | 0.02943 | 1e-05 | 1e-05 | 1e-05 | 1e-05 |
| LR-10A | 1e-05 | 0.99993 | 1e-05 | 1e-05 | 1e-05 | 1e-05 | 1e-05 | 1e-05 |
| LR-1A | 1e-05 | 0.98292 | 1e-05 | 0.01702 | 1e-05 | 1e-05 | 1e-05 | 1e-05 |
| LR-2A | 1e-05 | 0.99993 | 1e-05 | 1e-05 | 1e-05 | 1e-05 | 1e-05 | 1e-05 |
| LR-3A | 1e-05 | 0.999922 | 1.8e-05 | 1e-05 | 1e-05 | 1e-05 | 1e-05 | 1e-05 |
| LR-4A | 1e-05 | 0.99993 | 1e-05 | 1e-05 | 1e-05 | 1e-05 | 1e-05 | 1e-05 |
| LR-5A | 1e-05 | 0.99993 | 1e-05 | 1e-05 | 1e-05 | 1e-05 | 1e-05 | 1e-05 |
| LR-6A | 1e-05 | 0.99993 | 1e-05 | 1e-05 | 1e-05 | 1e-05 | 1e-05 | 1e-05 |
| LR-7A | 1e-05 | 0.99993 | 1e-05 | 1e-05 | 1e-05 | 1e-05 | 1e-05 | 1e-05 |
| LR-8A | 1e-05 | 0.999926 | 1.4e-05 | 1e-05 | 1e-05 | 1e-05 | 1e-05 | 1e-05 |
| LR-9A | 1e-05 | 0.999924 | 1e-05 | 1e-05 | 1.6e-05 | 1e-05 | 1e-05 | 1e-05 |
| BYR-10A | 1e-05 | 0.748406 | 1.2e-05 | 1e-05 | 1e-05 | 0.251527 | 1.5e-05 | 1e-05 |
| BYR-1A | 1e-05 | 0.99993 | 1e-05 | 1e-05 | 1e-05 | 1e-05 | 1e-05 | 1e-05 |
| BYR-2A | 1e-05 | 0.99993 | 1e-05 | 1e-05 | 1e-05 | 1e-05 | 1e-05 | 1e-05 |
| BYR-3A | 1e-05 | 0.988905 | 1e-05 | 1e-05 | 1.2e-05 | 0.011033 | 1e-05 | 1e-05 |
| BYR-4A | 1e-05 | 0.99993 | 1e-05 | 1e-05 | 1e-05 | 1e-05 | 1e-05 | 1e-05 |
| BYR-5A | 1e-05 | 0.99993 | 1e-05 | 1e-05 | 1e-05 | 1e-05 | 1e-05 | 1e-05 |
| BYR-6A | 1e-05 | 0.99993 | 1e-05 | 1e-05 | 1e-05 | 1e-05 | 1e-05 | 1e-05 |
| BYR-7A | 1e-05 | 0.850787 | 1e-05 | 1e-05 | 1e-05 | 0.149153 | 1e-05 | 1e-05 |

K = 9

|  |  |  |  |  |  |  |  |  |  |
| --- | --- | --- | --- | --- | --- | --- | --- | --- | --- |
| CDG-1A | 1.1e-05 | 1e-05 | 1.7e-05 | 1.5e-05 | 1e-05 | 1e-05 | 0.999908 | 1e-05 | 1e-05 |
| CDG-2A | 1e-05 | 1e-05 | 1e-05 | 1e-05 | 1e-05 | 1e-05 | 0.99992 | 1e-05 | 1e-05 |
| CDG-3A | 1e-05 | 1e-05 | 1e-05 | 1e-05 | 1e-05 | 1e-05 | 0.99992 | 1e-05 | 1e-05 |
| CDG-4A | 1e-05 | 1e-05 | 1.5e-05 | 1e-05 | 1.2e-05 | 1e-05 | 0.999913 | 1e-05 | 1e-05 |
| CDG-5A | 1e-05 | 1e-05 | 1e-05 | 1.3e-05 | 1e-05 | 1e-05 | 0.999907 | 2e-05 | 1e-05 |
| CDG-6A | 1e-05 | 1e-05 | 1e-05 | 1e-05 | 1e-05 | 1e-05 | 0.99992 | 1e-05 | 1e-05 |
| CDG-7A | 1e-05 | 1e-05 | 1e-05 | 1e-05 | 1e-05 | 1e-05 | 0.99992 | 1e-05 | 1e-05 |
| CDG-8A | 1.5e-05 | 1e-05 | 1e-05 | 1e-05 | 1e-05 | 1e-05 | 0.999915 | 1e-05 | 1e-05 |
| CDG-9A | 1e-05 | 1e-05 | 1e-05 | 1e-05 | 1e-05 | 1e-05 | 0.99992 | 1e-05 | 1e-05 |
| CDG-10A | 1.2e-05 | 1e-05 | 1e-05 | 1e-05 | 1e-05 | 1e-05 | 0.999918 | 1e-05 | 1e-05 |
| TKX-1A | 1.1e-05 | 0.999919 | 1e-05 | 1e-05 | 1e-05 | 1e-05 | 1e-05 | 1e-05 | 1e-05 |
| TKX-2A | 1e-05 | 0.99992 | 1e-05 | 1e-05 | 1e-05 | 1e-05 | 1e-05 | 1e-05 | 1e-05 |
| TKX-3A | 1e-05 | 0.999915 | 1.1e-05 | 1.1e-05 | 1e-05 | 1e-05 | 1.2e-05 | 1e-05 | 1e-05 |
| TKX-4A | 2e-05 | 0.99991 | 1e-05 | 1e-05 | 1e-05 | 1e-05 | 1e-05 | 1e-05 | 1e-05 |
| TKX-5A | 1e-05 | 0.99992 | 1e-05 | 1e-05 | 1e-05 | 1e-05 | 1e-05 | 1e-05 | 1e-05 |
| TKX-6A | 1e-05 | 0.99992 | 1e-05 | 1e-05 | 1e-05 | 1e-05 | 1e-05 | 1e-05 | 1e-05 |
| TKX-7A | 1e-05 | 0.999899 | 2e-05 | 1.2e-05 | 1e-05 | 1e-05 | 1.9e-05 | 1e-05 | 1e-05 |
| TKX-8A | 1e-05 | 0.99992 | 1e-05 | 1e-05 | 1e-05 | 1e-05 | 1e-05 | 1e-05 | 1e-05 |
| TKX-9A | 1e-05 | 0.99992 | 1e-05 | 1e-05 | 1e-05 | 1e-05 | 1e-05 | 1e-05 | 1e-05 |
| GJ-1A | 1e-05 | 1e-05 | 1e-05 | 1e-05 | 1e-05 | 1e-05 | 1e-05 | 0.99992 | 1e-05 |
| GJ-2A | 1e-05 | 1e-05 | 1e-05 | 1e-05 | 1e-05 | 1e-05 | 1e-05 | 0.99992 | 1e-05 |
| GJ-3A | 1e-05 | 1e-05 | 1e-05 | 1e-05 | 1e-05 | 1e-05 | 1e-05 | 0.99992 | 1e-05 |
| GJ-4A | 1e-05 | 1e-05 | 1.3e-05 | 1.9e-05 | 1e-05 | 1e-05 | 1e-05 | 0.999908 | 1e-05 |
| GJ-5A | 1e-05 | 1e-05 | 1e-05 | 1e-05 | 1e-05 | 1e-05 | 1e-05 | 0.99992 | 1e-05 |
| GJ-6A | 1e-05 | 1e-05 | 1.8e-05 | 1e-05 | 1.2e-05 | 1e-05 | 1e-05 | 0.999909 | 1e-05 |
| GJ-7A | 1e-05 | 1.4e-05 | 1.3e-05 | 1e-05 | 1e-05 | 1e-05 | 1.1e-05 | 0.999912 | 1e-05 |
| GJ-8A | 1e-05 | 1e-05 | 1e-05 | 1e-05 | 1e-05 | 1e-05 | 1e-05 | 0.99992 | 1e-05 |
| GJ-9A | 1e-05 | 1e-05 | 1e-05 | 1e-05 | 1.1e-05 | 1e-05 | 1e-05 | 0.999919 | 1e-05 |
| TZC-1A | 1e-05 | 1e-05 | 1e-05 | 1e-05 | 1e-05 | 1e-05 | 1e-05 | 0.99992 | 1e-05 |
| TZC-2A | 1e-05 | 1e-05 | 1e-05 | 1e-05 | 1e-05 | 1e-05 | 1e-05 | 0.99992 | 1e-05 |
| TZC-3A | 1e-05 | 1e-05 | 1e-05 | 0.105128 | 1e-05 | 1e-05 | 1e-05 | 0.87763 | 0.017182 |
| TZC-4A | 1e-05 | 1e-05 | 1e-05 | 1e-05 | 1e-05 | 1e-05 | 1e-05 | 0.99992 | 1e-05 |
| TZC-01A | 1e-05 | 1e-05 | 1e-05 | 1e-05 | 1e-05 | 1e-05 | 1e-05 | 0.99992 | 1e-05 |
| TZC-04A | 1e-05 | 1e-05 | 1e-05 | 1e-05 | 1e-05 | 1e-05 | 1e-05 | 0.99992 | 1e-05 |
| TZC-07A | 1e-05 | 1e-05 | 1e-05 | 1e-05 | 1e-05 | 1e-05 | 1e-05 | 0.99992 | 1e-05 |
| TZC-11A | 1e-05 | 1e-05 | 1e-05 | 1e-05 | 1e-05 | 1e-05 | 1e-05 | 0.99992 | 1e-05 |
| TZC-13A | 1e-05 | 1e-05 | 1e-05 | 1e-05 | 1e-05 | 1e-05 | 1e-05 | 0.99992 | 1e-05 |
| DBG-10A | 1e-05 | 1e-05 | 1e-05 | 1e-05 | 1e-05 | 1e-05 | 1e-05 | 0.981288 | 0.018642 |
| DBG-1A | 1e-05 | 1e-05 | 0.072725 | 1e-05 | 1e-05 | 0.048941 | 1e-05 | 0.802592 | 0.075692 |
| DBG-2A | 1e-05 | 1e-05 | 0.098887 | 1e-05 | 1e-05 | 1e-05 | 1e-05 | 0.901043 | 1e-05 |
| DBG-3A | 1e-05 | 1e-05 | 0.140235 | 1e-05 | 1e-05 | 1e-05 | 1e-05 | 0.859695 | 1e-05 |
| DBG-4A | 1e-05 | 1e-05 | 0.052745 | 1.5e-05 | 1e-05 | 1e-05 | 1e-05 | 0.94718 | 1e-05 |
| DBG-5A | 1e-05 | 1e-05 | 1e-05 | 1e-05 | 1e-05 | 1e-05 | 1e-05 | 0.872134 | 0.127796 |
| DBG-6A | 1e-05 | 0.008975 | 1e-05 | 1e-05 | 1e-05 | 1e-05 | 1e-05 | 0.990955 | 1e-05 |
| DBG-7A | 1e-05 | 1e-05 | 1e-05 | 1e-05 | 1e-05 | 1e-05 | 1e-05 | 0.99992 | 1e-05 |
| DBG-8A | 1e-05 | 1e-05 | 1.1e-05 | 1e-05 | 1e-05 | 1e-05 | 1e-05 | 0.999919 | 1e-05 |
| DBG-9A | 1e-05 | 0.023041 | 0.092627 | 1e-05 | 1e-05 | 1e-05 | 1e-05 | 0.884272 | 1e-05 |
| TEG-10A | 1e-05 | 1e-05 | 1e-05 | 1e-05 | 0.015658 | 1e-05 | 1e-05 | 0.422735 | 0.561547 |
| TEG-1A | 0.018792 | 1e-05 | 0.160204 | 1e-05 | 1e-05 | 0.043773 | 1e-05 | 0.471179 | 0.306012 |
| TEG-2A | 0.00997 | 1e-05 | 0.24074 | 1e-05 | 1e-05 | 1e-05 | 1e-05 | 0.420838 | 0.328402 |
| TEG-3A | 1e-05 | 1.4e-05 | 0.132742 | 1e-05 | 1e-05 | 1e-05 | 1e-05 | 0.533324 | 0.33387 |
| TEG-4A | 1e-05 | 1e-05 | 0.152724 | 1e-05 | 1.3e-05 | 1e-05 | 1e-05 | 0.578463 | 0.268749 |
| TEG-5A | 1e-05 | 1e-05 | 0.16978 | 1.1e-05 | 1e-05 | 1e-05 | 1e-05 | 0.541236 | 0.288924 |
| TEG-6A | 1e-05 | 1e-05 | 0.228058 | 1e-05 | 0.005854 | 1e-05 | 0.011739 | 0.461107 | 0.293203 |
| TEG-7A | 1e-05 | 1e-05 | 0.145858 | 1e-05 | 0.06143 | 0.045353 | 1e-05 | 0.419016 | 0.328304 |
| TEG-8A | 1e-05 | 1e-05 | 0.076693 | 1.2e-05 | 0.059807 | 1e-05 | 1e-05 | 0.491588 | 0.37186 |
| TEG-9A | 0.017775 | 1.7e-05 | 0.166736 | 1.2e-05 | 1e-05 | 1e-05 | 1e-05 | 0.501141 | 0.31429 |
| ZPZ-10A | 1e-05 | 1e-05 | 1e-05 | 1e-05 | 1e-05 | 1e-05 | 1e-05 | 1e-05 | 0.999919 |
| ZPZ-1A | 1.2e-05 | 1e-05 | 1e-05 | 1e-05 | 1e-05 | 1e-05 | 1e-05 | 1e-05 | 0.999918 |
| ZPZ-2A | 1e-05 | 1e-05 | 1e-05 | 1e-05 | 1e-05 | 1e-05 | 1e-05 | 1e-05 | 0.99992 |
| ZPZ-3A | 1e-05 | 1e-05 | 0.034445 | 1e-05 | 0.038473 | 1e-05 | 1e-05 | 1e-05 | 0.927022 |
| ZPZ-4A | 1.1e-05 | 1e-05 | 1.4e-05 | 1e-05 | 1.3e-05 | 1e-05 | 1e-05 | 1e-05 | 0.999912 |
| ZPZ-5A | 1e-05 | 1e-05 | 1e-05 | 1e-05 | 1e-05 | 1e-05 | 1e-05 | 1e-05 | 0.99992 |
| ZPZ-6A | 0.034258 | 1e-05 | 0.245452 | 1e-05 | 0.052244 | 1e-05 | 1e-05 | 0.037071 | 0.630935 |
| ZPZ-7A | 1e-05 | 1e-05 | 0.100703 | 1e-05 | 1.2e-05 | 1e-05 | 1e-05 | 0.126744 | 0.772492 |
| ZPZ-8A | 1.1e-05 | 1e-05 | 1e-05 | 1e-05 | 1e-05 | 1e-05 | 1e-05 | 1e-05 | 0.999919 |
| ZPZ-9A | 1e-05 | 1e-05 | 1e-05 | 1e-05 | 1e-05 | 1e-05 | 1e-05 | 1e-05 | 0.99992 |
| YLJ1-1A | 1.1e-05 | 1e-05 | 1e-05 | 1.2e-05 | 1.1e-05 | 1e-05 | 1e-05 | 1e-05 | 0.999916 |
| YLJ1-2A | 1.1e-05 | 1.1e-05 | 1.1e-05 | 1.2e-05 | 1e-05 | 1e-05 | 1e-05 | 1e-05 | 0.999915 |
| YLJ1-3A | 1e-05 | 1e-05 | 1e-05 | 1.1e-05 | 1e-05 | 1e-05 | 1e-05 | 1e-05 | 0.999919 |
| YLJ1-4A | 1e-05 | 1.1e-05 | 1.1e-05 | 1e-05 | 1e-05 | 1e-05 | 1.1e-05 | 1e-05 | 0.999917 |
| YLJ1-5A | 1.1e-05 | 1e-05 | 1e-05 | 1.1e-05 | 1e-05 | 1e-05 | 1e-05 | 1e-05 | 0.999918 |
| YLJ1-6A | 1e-05 | 0.010266 | 1e-05 | 1e-05 | 1e-05 | 1e-05 | 1e-05 | 1e-05 | 0.989664 |
| YLJ1-7A | 1e-05 | 1e-05 | 1.1e-05 | 1e-05 | 1e-05 | 1e-05 | 1.1e-05 | 1e-05 | 0.999919 |
| YLJ1-8A | 1.6e-05 | 1e-05 | 0.081448 | 1.1e-05 | 0.001586 | 1e-05 | 1e-05 | 1e-05 | 0.916899 |
| YLJ2-1A | 1e-05 | 1e-05 | 1e-05 | 1e-05 | 1e-05 | 1e-05 | 1e-05 | 1e-05 | 0.99992 |
| YLJ2-2A | 1e-05 | 1e-05 | 1e-05 | 1e-05 | 1e-05 | 1e-05 | 1e-05 | 1e-05 | 0.99992 |
| YLJ2-3A | 1.6e-05 | 1e-05 | 1e-05 | 1e-05 | 0.009822 | 1e-05 | 1e-05 | 1e-05 | 0.990102 |
| YLJ2-4A | 1e-05 | 1e-05 | 0.155634 | 1e-05 | 1.1e-05 | 1e-05 | 1e-05 | 1e-05 | 0.844296 |
| YLJ2-5A | 1e-05 | 1e-05 | 1e-05 | 1e-05 | 1e-05 | 1e-05 | 1e-05 | 1e-05 | 0.99992 |
| YLJ2-6B | 1e-05 | 1e-05 | 1e-05 | 1e-05 | 1e-05 | 1e-05 | 1e-05 | 1e-05 | 0.99992 |
| YLJ2-7A | 1e-05 | 1e-05 | 1.5e-05 | 1e-05 | 1e-05 | 1e-05 | 1e-05 | 1e-05 | 0.999915 |
| YLJ2-8A | 1e-05 | 1e-05 | 0.164052 | 1e-05 | 1e-05 | 1e-05 | 1e-05 | 1e-05 | 0.835878 |
| LM-10A | 1e-05 | 1e-05 | 1e-05 | 1e-05 | 0.99992 | 1e-05 | 1e-05 | 1e-05 | 1e-05 |
| LM-1A | 1.1e-05 | 1e-05 | 1e-05 | 1e-05 | 0.999919 | 1e-05 | 1e-05 | 1e-05 | 1e-05 |
| LM-2A | 1e-05 | 1e-05 | 1e-05 | 1e-05 | 0.99992 | 1e-05 | 1e-05 | 1e-05 | 1e-05 |
| LM-3A | 1e-05 | 1e-05 | 1e-05 | 1e-05 | 0.999919 | 1.1e-05 | 1e-05 | 1e-05 | 1e-05 |
| LM-4A | 1e-05 | 1e-05 | 1.5e-05 | 1e-05 | 0.999915 | 1e-05 | 1e-05 | 1e-05 | 1e-05 |
| LM-5A | 1e-05 | 1e-05 | 1e-05 | 1e-05 | 0.99992 | 1e-05 | 1e-05 | 1e-05 | 1e-05 |
| LM-6A | 1e-05 | 1e-05 | 1e-05 | 1e-05 | 0.99992 | 1e-05 | 1e-05 | 1e-05 | 1e-05 |
| LM-7A | 1e-05 | 1e-05 | 1.7e-05 | 1e-05 | 0.999913 | 1e-05 | 1e-05 | 1e-05 | 1e-05 |
| LM-8A | 1e-05 | 1e-05 | 1.3e-05 | 1e-05 | 0.999917 | 1e-05 | 1e-05 | 1e-05 | 1e-05 |
| LM-9A | 1e-05 | 1e-05 | 1e-05 | 1e-05 | 0.999912 | 1.8e-05 | 1e-05 | 1e-05 | 1e-05 |
| LM-11A | 1e-05 | 1e-05 | 1e-05 | 1e-05 | 0.999917 | 1.3e-05 | 1e-05 | 1e-05 | 1e-05 |
| LM-12A | 1e-05 | 1e-05 | 1.1e-05 | 1e-05 | 0.994294 | 1e-05 | 1e-05 | 1e-05 | 0.005635 |
| LM-13A | 1e-05 | 1e-05 | 1e-05 | 1e-05 | 0.99992 | 1e-05 | 1e-05 | 1e-05 | 1e-05 |
| LM-14A | 1e-05 | 1e-05 | 1.2e-05 | 1e-05 | 0.999918 | 1e-05 | 1e-05 | 1e-05 | 1e-05 |
| LM-15A | 1e-05 | 1e-05 | 1e-05 | 1e-05 | 0.999919 | 1e-05 | 1.1e-05 | 1e-05 | 1e-05 |
| BYP1-1A | 1e-05 | 1e-05 | 0.99992 | 1e-05 | 1e-05 | 1e-05 | 1e-05 | 1e-05 | 1e-05 |
| BYP1-2A | 1e-05 | 1.1e-05 | 0.963423 | 0.0162 | 0.020316 | 1e-05 | 1e-05 | 1e-05 | 1e-05 |
| BYP1-3A | 1e-05 | 1e-05 | 0.962657 | 1e-05 | 1e-05 | 1e-05 | 1e-05 | 1e-05 | 0.037273 |
| BYP1-4A | 1e-05 | 1.2e-05 | 0.999915 | 1e-05 | 1e-05 | 1e-05 | 1e-05 | 1.2e-05 | 1e-05 |
| BYP1-5A | 1e-05 | 1e-05 | 0.896967 | 1e-05 | 1.8e-05 | 1e-05 | 1e-05 | 1e-05 | 0.102955 |
| BYP1-6A | 1e-05 | 1e-05 | 0.99992 | 1e-05 | 1e-05 | 1e-05 | 1e-05 | 1e-05 | 1e-05 |
| BYP1-7A | 1e-05 | 1e-05 | 0.999915 | 1e-05 | 1.4e-05 | 1e-05 | 1e-05 | 1.1e-05 | 1e-05 |
| BYP1-8A | 1e-05 | 1e-05 | 0.99992 | 1e-05 | 1e-05 | 1e-05 | 1e-05 | 1e-05 | 1e-05 |
| BYP1-9A | 1e-05 | 1e-05 | 0.99992 | 1e-05 | 1e-05 | 1e-05 | 1e-05 | 1e-05 | 1e-05 |
| BYP1-10A | 1e-05 | 1e-05 | 0.992279 | 1e-05 | 1e-05 | 1e-05 | 0.007651 | 1e-05 | 1e-05 |
| BYP2-1A | 1e-05 | 1e-05 | 0.99992 | 1e-05 | 1e-05 | 1e-05 | 1e-05 | 1e-05 | 1e-05 |
| BYP2-2A | 1e-05 | 1e-05 | 0.999917 | 1.2e-05 | 1e-05 | 1e-05 | 1e-05 | 1e-05 | 1e-05 |
| BYP2-3A | 1e-05 | 1e-05 | 0.950568 | 1.3e-05 | 0.049359 | 1e-05 | 1e-05 | 1e-05 | 1e-05 |
| BYP2-4A | 1e-05 | 1e-05 | 0.99992 | 1e-05 | 1e-05 | 1e-05 | 1e-05 | 1e-05 | 1e-05 |
| BYP2-5A | 1e-05 | 1e-05 | 0.834048 | 0.102487 | 1e-05 | 1e-05 | 1e-05 | 0.053376 | 0.010039 |
| BYP2-6A | 1e-05 | 1e-05 | 0.959692 | 1e-05 | 0.040238 | 1e-05 | 1e-05 | 1e-05 | 1e-05 |
| BYP2-7A | 1e-05 | 1e-05 | 0.544677 | 0.004635 | 0.374323 | 1e-05 | 1e-05 | 1e-05 | 0.076316 |
| BYP2-8A | 1e-05 | 1e-05 | 0.99992 | 1e-05 | 1e-05 | 1e-05 | 1e-05 | 1e-05 | 1e-05 |
| BYP3-1A | 1e-05 | 1e-05 | 0.999918 | 1.2e-05 | 1e-05 | 1e-05 | 1e-05 | 1e-05 | 1e-05 |
| BYP3-2A | 1e-05 | 1e-05 | 0.99992 | 1e-05 | 1e-05 | 1e-05 | 1e-05 | 1e-05 | 1e-05 |
| BYP3-3A | 1e-05 | 1e-05 | 0.912426 | 1e-05 | 0.087504 | 1e-05 | 1e-05 | 1e-05 | 1e-05 |
| BYP3-4A | 1e-05 | 1e-05 | 0.99992 | 1e-05 | 1e-05 | 1e-05 | 1e-05 | 1e-05 | 1e-05 |
| BYP3-5A | 1e-05 | 1e-05 | 0.987182 | 1e-05 | 0.012748 | 1e-05 | 1e-05 | 1e-05 | 1e-05 |
| BYP3-6A | 1e-05 | 1e-05 | 0.886722 | 1e-05 | 0.107734 | 1e-05 | 1e-05 | 1e-05 | 0.005483 |
| BYP3-7A | 1e-05 | 1.1e-05 | 0.999919 | 1e-05 | 1e-05 | 1e-05 | 1e-05 | 1e-05 | 1e-05 |
| BYP3-8A | 1.1e-05 | 1e-05 | 0.999918 | 1e-05 | 1e-05 | 1e-05 | 1e-05 | 1e-05 | 1e-05 |
| KBG-1A | 1e-05 | 1e-05 | 1e-05 | 0.99992 | 1e-05 | 1e-05 | 1e-05 | 1e-05 | 1e-05 |
| KBG-2A | 1e-05 | 1e-05 | 1.7e-05 | 0.923258 | 1e-05 | 1e-05 | 1e-05 | 1e-05 | 0.076665 |
| KBG-3A | 1e-05 | 1e-05 | 0.007023 | 0.992907 | 1.1e-05 | 1e-05 | 1e-05 | 1e-05 | 1e-05 |
| KBG-4A | 1e-05 | 1.2e-05 | 0.043316 | 0.956609 | 1.3e-05 | 1e-05 | 1e-05 | 1e-05 | 1e-05 |
| KBG-5A | 1e-05 | 1e-05 | 1.1e-05 | 0.999912 | 1e-05 | 1.7e-05 | 1e-05 | 1e-05 | 1e-05 |
| QJG-1A | 1e-05 | 1e-05 | 1e-05 | 0.999918 | 1e-05 | 1.2e-05 | 1e-05 | 1e-05 | 1e-05 |
| QJG-2A | 1e-05 | 1e-05 | 0.006654 | 0.968941 | 1e-05 | 1e-05 | 1e-05 | 1e-05 | 0.024345 |
| QJG-3A | 1e-05 | 1e-05 | 1e-05 | 0.99992 | 1e-05 | 1e-05 | 1e-05 | 1e-05 | 1e-05 |
| QJG-4A | 1e-05 | 1e-05 | 1e-05 | 0.999917 | 1.1e-05 | 1.2e-05 | 1e-05 | 1e-05 | 1e-05 |
| QJG-5A | 0.000952 | 1e-05 | 1e-05 | 0.998978 | 1e-05 | 1e-05 | 1e-05 | 1e-05 | 1e-05 |
| LTG-1A | 1e-05 | 1e-05 | 0.202805 | 0.672222 | 1e-05 | 1e-05 | 1e-05 | 0.124912 | 1e-05 |
| LTG-2A | 1e-05 | 0.017899 | 0.156241 | 0.809274 | 1e-05 | 1e-05 | 1e-05 | 0.016536 | 1e-05 |
| LTG-3A | 1e-05 | 1e-05 | 1e-05 | 0.99992 | 1e-05 | 1e-05 | 1e-05 | 1e-05 | 1e-05 |
| LTG-4A | 1e-05 | 1e-05 | 1.1e-05 | 0.999919 | 1e-05 | 1e-05 | 1e-05 | 1e-05 | 1e-05 |
| LTG-5A | 1e-05 | 0.050403 | 0.097652 | 0.851883 | 1.1e-05 | 1e-05 | 1e-05 | 1e-05 | 1e-05 |
| BW-10B | 1e-05 | 1e-05 | 1e-05 | 0.986555 | 1e-05 | 1e-05 | 1e-05 | 1e-05 | 0.013375 |
| BW-1B | 1e-05 | 1e-05 | 1e-05 | 0.99992 | 1e-05 | 1e-05 | 1e-05 | 1e-05 | 1e-05 |
| BW-2B | 1e-05 | 1e-05 | 1e-05 | 0.999919 | 1e-05 | 1e-05 | 1e-05 | 1e-05 | 1e-05 |
| BW-3B | 1e-05 | 1e-05 | 1e-05 | 0.99992 | 1e-05 | 1e-05 | 1e-05 | 1e-05 | 1e-05 |
| BW-4B | 1e-05 | 1e-05 | 0.065084 | 0.934846 | 1e-05 | 1e-05 | 1e-05 | 1e-05 | 1e-05 |
| BW-5B | 1e-05 | 1e-05 | 1e-05 | 0.99992 | 1e-05 | 1e-05 | 1e-05 | 1e-05 | 1e-05 |
| BW-6B | 1.1e-05 | 1.2e-05 | 1e-05 | 0.999917 | 1e-05 | 1e-05 | 1e-05 | 1e-05 | 1e-05 |
| BW-7A | 1e-05 | 1e-05 | 1e-05 | 0.99992 | 1e-05 | 1e-05 | 1e-05 | 1e-05 | 1e-05 |
| BW-8B | 1e-05 | 1e-05 | 1e-05 | 0.985635 | 0.014295 | 1e-05 | 1e-05 | 1e-05 | 1e-05 |
| BW-9A | 1e-05 | 1e-05 | 1e-05 | 0.99992 | 1e-05 | 1e-05 | 1e-05 | 1e-05 | 1e-05 |
| GA-1A | 1e-05 | 1e-05 | 1e-05 | 0.958173 | 1e-05 | 0.032002 | 1e-05 | 1e-05 | 0.009765 |
| GA-2A | 1e-05 | 1e-05 | 0.104973 | 0.888688 | 0.006279 | 1e-05 | 1e-05 | 1e-05 | 1e-05 |
| GA-3A | 1e-05 | 1e-05 | 0.218915 | 0.781015 | 1e-05 | 1e-05 | 1e-05 | 1e-05 | 1e-05 |
| MR-1A | 1.6e-05 | 1e-05 | 1e-05 | 0.999914 | 1e-05 | 1e-05 | 1e-05 | 1e-05 | 1e-05 |
| MR-2A | 1e-05 | 1e-05 | 1.1e-05 | 0.999919 | 1e-05 | 1e-05 | 1e-05 | 1e-05 | 1e-05 |
| MR-3A | 0.0279 | 0.003861 | 1e-05 | 0.887915 | 0.080274 | 1e-05 | 1e-05 | 1e-05 | 1e-05 |
| MR-4A | 0.000849 | 1e-05 | 0.132812 | 0.86628 | 1e-05 | 1e-05 | 1e-05 | 1e-05 | 1e-05 |
| MR-5A | 1e-05 | 1e-05 | 1e-05 | 0.993757 | 1e-05 | 1e-05 | 0.006173 | 1e-05 | 1e-05 |
| MDG-6A | 1e-05 | 1e-05 | 1e-05 | 0.786107 | 1.2e-05 | 0.213818 | 1.2e-05 | 1e-05 | 1e-05 |
| MDG-1A | 1.1e-05 | 1e-05 | 1e-05 | 0.88845 | 1e-05 | 0.111478 | 1e-05 | 1e-05 | 1e-05 |
| MDG-2A | 1e-05 | 1e-05 | 1e-05 | 0.894848 | 1e-05 | 0.105082 | 1e-05 | 1e-05 | 1e-05 |
| MDG-3A | 1e-05 | 1e-05 | 1e-05 | 0.788802 | 0.033896 | 0.177241 | 1e-05 | 1.1e-05 | 1e-05 |
| MDG-4A | 0.116671 | 1e-05 | 1e-05 | 0.772963 | 1e-05 | 0.09177 | 1e-05 | 0.018546 | 1e-05 |
| MDG-5A | 0.072162 | 1e-05 | 1e-05 | 0.838929 | 0.00466 | 0.084199 | 1e-05 | 1e-05 | 1e-05 |
| AJ-10A | 1e-05 | 1e-05 | 1e-05 | 0.107646 | 0.009226 | 0.858562 | 1e-05 | 0.024517 | 1e-05 |
| AJ-12A | 1e-05 | 1e-05 | 1e-05 | 1e-05 | 1e-05 | 0.999918 | 1.2e-05 | 1e-05 | 1e-05 |
| AJ-13A | 1e-05 | 1e-05 | 1e-05 | 1e-05 | 1e-05 | 0.99992 | 1e-05 | 1e-05 | 1e-05 |
| AJ-14A | 1.9e-05 | 1e-05 | 1e-05 | 0.035957 | 1e-05 | 0.963964 | 1e-05 | 1e-05 | 1e-05 |
| AJ-1A | 1e-05 | 1e-05 | 1e-05 | 0.205772 | 1e-05 | 0.794158 | 1e-05 | 1e-05 | 1e-05 |
| AJ-3A | 0.140776 | 1e-05 | 1e-05 | 0.172112 | 0.047192 | 0.639871 | 1e-05 | 1e-05 | 1e-05 |
| AJ-4A | 0.08995 | 1e-05 | 1e-05 | 0.06423 | 1e-05 | 0.815381 | 1e-05 | 0.030389 | 1e-05 |
| AJ-5A | 0.089605 | 1e-05 | 1e-05 | 0.166003 | 1e-05 | 0.744332 | 1e-05 | 1e-05 | 1e-05 |
| AJ-7A | 1e-05 | 1e-05 | 1e-05 | 0.219041 | 1e-05 | 0.722352 | 1.3e-05 | 0.058544 | 1e-05 |
| AJ-9A | 1e-05 | 1e-05 | 1e-05 | 1e-05 | 1e-05 | 0.99992 | 1e-05 | 1e-05 | 1e-05 |
| XGG-1A | 0.949282 | 1e-05 | 1e-05 | 1e-05 | 1e-05 | 0.050648 | 1e-05 | 1e-05 | 1e-05 |
| XGG-2A | 0.799093 | 1e-05 | 1e-05 | 1.8e-05 | 1.2e-05 | 0.200826 | 1e-05 | 1e-05 | 1e-05 |
| XGG-3A | 0.830797 | 1e-05 | 1e-05 | 1e-05 | 1e-05 | 0.169131 | 1.1e-05 | 1e-05 | 1e-05 |
| XGG-4A | 0.890412 | 1e-05 | 1e-05 | 1e-05 | 1e-05 | 0.109518 | 1e-05 | 1e-05 | 1e-05 |
| XGG-5A | 0.771812 | 1e-05 | 1e-05 | 0.030143 | 1e-05 | 0.197985 | 1e-05 | 1e-05 | 1e-05 |
| PSR-01A | 0.902619 | 1e-05 | 1.1e-05 | 1e-05 | 1e-05 | 0.09731 | 1e-05 | 1e-05 | 1e-05 |
| PSR-04A | 0.929771 | 1e-05 | 1e-05 | 1.9e-05 | 1e-05 | 0.070149 | 1e-05 | 1e-05 | 1e-05 |
| PSR-10A | 0.921223 | 1e-05 | 1e-05 | 1e-05 | 1e-05 | 0.078706 | 1e-05 | 1e-05 | 1e-05 |
| PSR-13A | 0.949606 | 1e-05 | 0.050323 | 1e-05 | 1.2e-05 | 1e-05 | 1e-05 | 1e-05 | 1e-05 |
| LR-10A | 0.99992 | 1e-05 | 1e-05 | 1e-05 | 1e-05 | 1e-05 | 1e-05 | 1e-05 | 1e-05 |
| LR-1A | 0.976065 | 1.2e-05 | 1e-05 | 0.023862 | 1e-05 | 1e-05 | 1.1e-05 | 1e-05 | 1e-05 |
| LR-2A | 0.99992 | 1e-05 | 1e-05 | 1e-05 | 1e-05 | 1e-05 | 1e-05 | 1e-05 | 1e-05 |
| LR-3A | 0.99992 | 1e-05 | 1e-05 | 1e-05 | 1e-05 | 1e-05 | 1e-05 | 1e-05 | 1e-05 |
| LR-4A | 0.999915 | 1.2e-05 | 1e-05 | 1e-05 | 1e-05 | 1.3e-05 | 1e-05 | 1e-05 | 1e-05 |
| LR-5A | 0.999919 | 1e-05 | 1e-05 | 1e-05 | 1e-05 | 1e-05 | 1e-05 | 1.1e-05 | 1e-05 |
| LR-6A | 0.99992 | 1e-05 | 1e-05 | 1e-05 | 1e-05 | 1e-05 | 1e-05 | 1e-05 | 1e-05 |
| LR-7A | 0.99992 | 1e-05 | 1e-05 | 1e-05 | 1e-05 | 1e-05 | 1e-05 | 1e-05 | 1e-05 |
| LR-8A | 0.99992 | 1e-05 | 1e-05 | 1e-05 | 1e-05 | 1e-05 | 1e-05 | 1e-05 | 1e-05 |
| LR-9A | 0.99992 | 1e-05 | 1e-05 | 1e-05 | 1e-05 | 1e-05 | 1e-05 | 1e-05 | 1e-05 |
| BYR-10A | 0.748791 | 1.1e-05 | 1e-05 | 1.2e-05 | 1e-05 | 0.251135 | 1e-05 | 1.1e-05 | 1e-05 |
| BYR-1A | 0.999916 | 1.2e-05 | 1e-05 | 1e-05 | 1e-05 | 1e-05 | 1e-05 | 1.2e-05 | 1e-05 |
| BYR-2A | 0.999919 | 1e-05 | 1.1e-05 | 1e-05 | 1e-05 | 1e-05 | 1e-05 | 1e-05 | 1e-05 |
| BYR-3A | 0.988623 | 1e-05 | 1.2e-05 | 1.3e-05 | 1.2e-05 | 0.011299 | 1e-05 | 1e-05 | 1e-05 |
| BYR-4A | 0.99992 | 1e-05 | 1e-05 | 1e-05 | 1e-05 | 1e-05 | 1e-05 | 1e-05 | 1e-05 |
| BYR-5A | 0.99992 | 1e-05 | 1e-05 | 1e-05 | 1e-05 | 1e-05 | 1e-05 | 1e-05 | 1e-05 |
| BYR-6A | 0.99992 | 1e-05 | 1e-05 | 1e-05 | 1e-05 | 1e-05 | 1e-05 | 1e-05 | 1e-05 |
| BYR-7A | 0.851364 | 1e-05 | 1e-05 | 1.4e-05 | 1e-05 | 0.14856 | 1e-05 | 1.2e-05 | 1e-05 |

**Figure S11** Population structure of 227 *A. pentaphyllum* individuals at *K* = 2 to *K* =9.

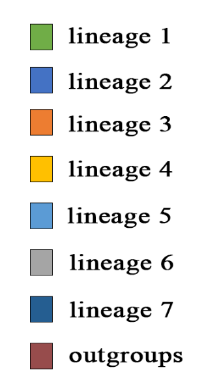

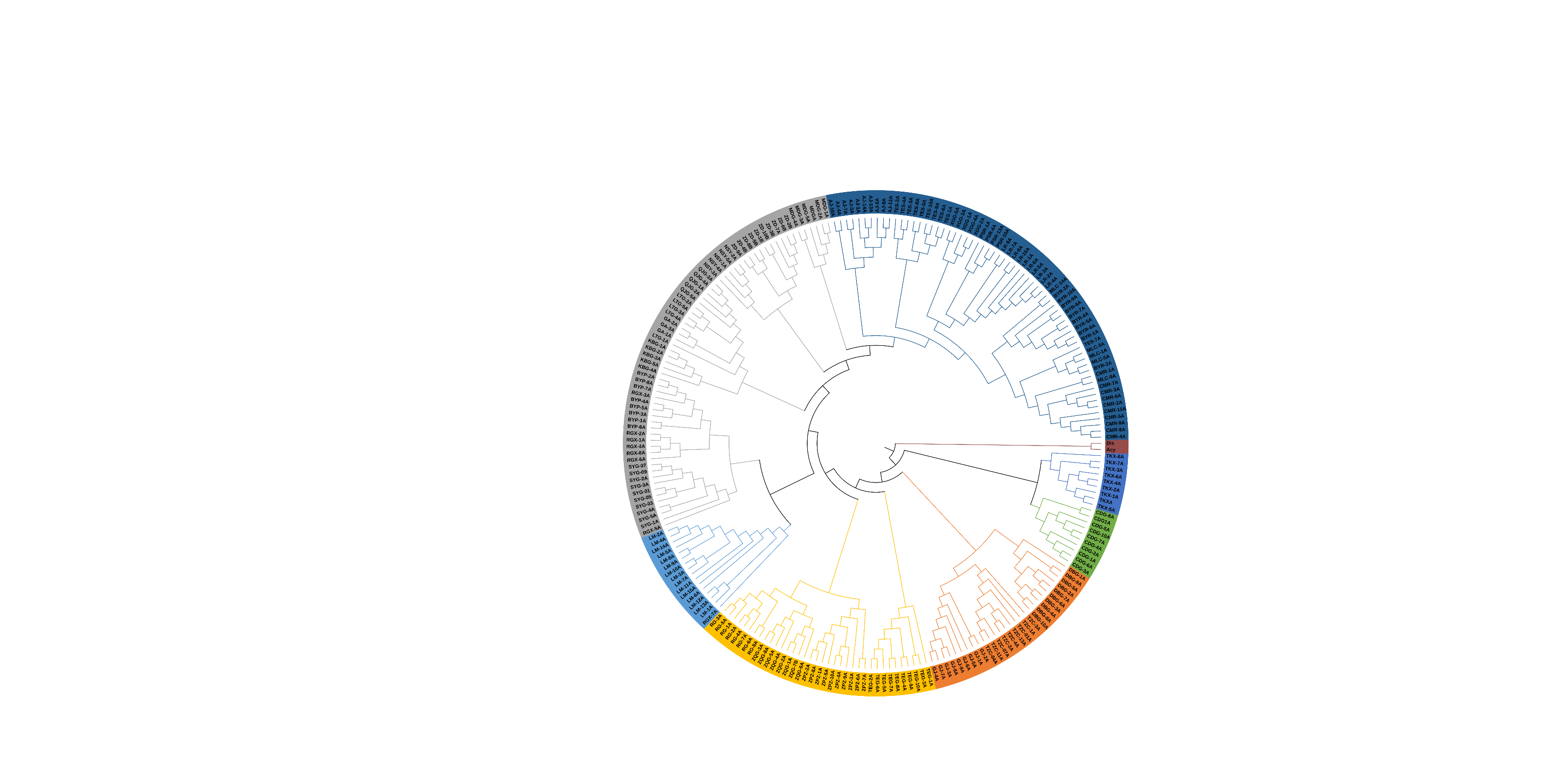

**Figure S12** The phylogenetic tree of *A. pentaphyllum* based on SNP Dataset 4 (different colors represent different clusters at *K* = 7).

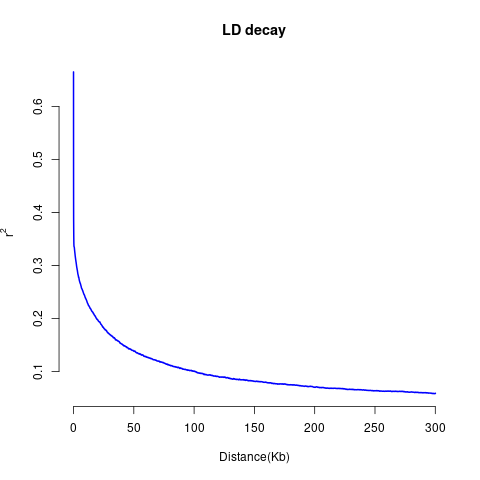

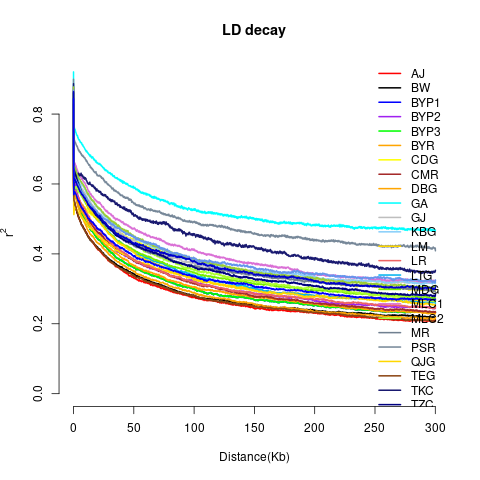

(b)

(a)

**Figure S13** Genome-wide linkage disequilibrium (LD) decay in *A. pentaphyllum* among 28 separate populations (a) and considering the ten populations as a whole (b).

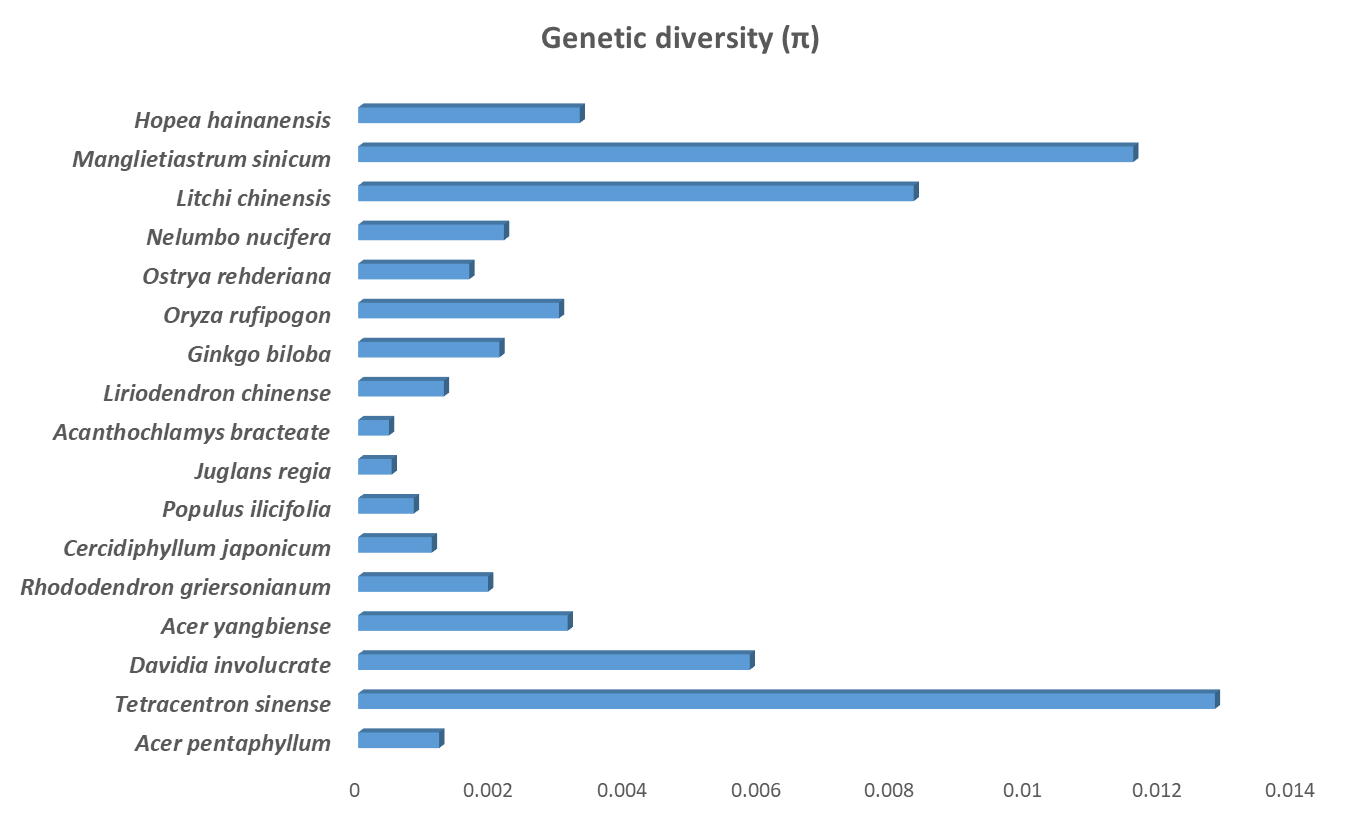

**Figure S14** Genome-wide sequence diversity (π) for 17 threatened woody species.

Geographic distance vs Genetic distance

Genetic distance

| Geographic distance | Y |
| --- | --- |
| 25.7987 | 0.330978 |
| 32.1556 | 0.33411 |
| 6.56475 | 0.197734 |
| 33.5972 | 0.2939 |
| 8.65186 | 0.176202 |
| 6.21723 | 0.143647 |
| 45.1271 | 0.327736 |
| 19.7886 | 0.196129 |
| 14.6917 | 0.17436 |
| 11.5304 | 0.152889 |
| 54.0549 | 0.340997 |
| 29.0446 | 0.25672 |
| 24.0181 | 0.217658 |
| 20.5814 | 0.17551 |
| 9.34343 | 0.158982 |
| 59.6865 | 0.368324 |
| 34.1761 | 0.298966 |
| 28.5166 | 0.254078 |
| 26.0972 | 0.21338 |
| 14.5715 | 0.201257 |
| 6.80525 | 0.176309 |
| 60.9999 | 0.370195 |
| 35.6068 | 0.278878 |
| 30.0693 | 0.233201 |
| 27.405 | 0.191335 |
| 15.8828 | 0.150079 |
| 7.4106 | 0.163156 |
| 1.87801 | 0.193288 |
| 81.0052 | 0.375695 |
| 64.1663 | 0.272986 |
| 62.6388 | 0.25275 |
| 56.6687 | 0.235828 |
| 50.931 | 0.201745 |
| 44.9899 | 0.230151 |
| 47.3986 | 0.259166 |
| 45.8943 | 0.238848 |
| 68.4648 | 0.324165 |
| 44.1671 | 0.232564 |
| 39.4344 | 0.20096 |
| 35.5544 | 0.155866 |
| 24.7437 | 0.148623 |
| 15.4635 | 0.157647 |
| 13.4475 | 0.199436 |
| 11.5892 | 0.159555 |
| 36.137 | 0.187753 |
| 68.7985 | 0.326337 |
| 44.9478 | 0.225585 |
| 40.5006 | 0.20242 |
| 36.2998 | 0.142039 |
| 25.8686 | 0.140433 |
| 16.7868 | 0.154105 |
| 15.6554 | 0.185835 |
| 13.8411 | 0.160919 |
| 33.4085 | 0.17592 |
| 2.72864 | 0.085858 |
| 71.0009 | 0.327074 |
| 47.3814 | 0.221681 |
| 43.0397 | 0.198489 |
| 38.7295 | 0.144852 |
| 28.441 | 0.120231 |
| 19.418 | 0.157204 |
| 18.2529 | 0.197627 |
| 16.4207 | 0.160714 |
| 31.6218 | 0.172146 |
| 4.94762 | 0.072197 |
| 2.6653 | 0.057717 |
| 84.1939 | 0.33147 |
| 59.9195 | 0.220957 |
| 55.0363 | 0.210202 |
| 51.3218 | 0.130384 |
| 40.3619 | 0.159752 |
| 31.021 | 0.196693 |
| 27.4996 | 0.245832 |
| 25.6798 | 0.189143 |
| 36.3809 | 0.218756 |
| 15.7799 | 0.126004 |
| 15.4928 | 0.116932 |
| 13.6297 | 0.112649 |
| 81.1532 | 0.348745 |
| 56.9736 | 0.264765 |
| 52.1735 | 0.236009 |
| 48.3608 | 0.167948 |
| 37.4857 | 0.177546 |
| 28.1576 | 0.22316 |
| 24.9464 | 0.266153 |
| 23.0993 | 0.212659 |
| 35.0543 | 0.227919 |
| 12.8069 | 0.143591 |
| 12.4134 | 0.144885 |
| 10.5461 | 0.134638 |
| 3.09005 | 0.060661 |
| 79.4701 | 0.335675 |
| 54.9918 | 0.253104 |
| 50.017 | 0.233894 |
| 46.4227 | 0.166542 |
| 35.3625 | 0.180507 |
| 26.0194 | 0.199728 |
| 22.3895 | 0.240066 |
| 20.5694 | 0.215687 |
| 37.1051 | 0.226863 |
| 11.018 | 0.128593 |
| 11.2146 | 0.127373 |
| 9.89401 | 0.124069 |
| 5.11052 | 0.095344 |
| 2.97144 | 0.093071 |
| 79.7263 | 0.309562 |
| 54.5663 | 0.208978 |
| 49.1 | 0.18798 |
| 46.2215 | 0.160229 |
| 34.782 | 0.160335 |
| 25.6725 | 0.196731 |
| 20.6271 | 0.227288 |
| 19.0307 | 0.19902 |
| 43.4921 | 0.182985 |
| 13.0273 | 0.133962 |
| 14.4689 | 0.121424 |
| 14.234 | 0.121734 |
| 9.71514 | 0.103915 |
| 9.11648 | 0.111056 |
| 6.49092 | 0.115815 |
| 83.4622 | 0.363445 |
| 58.4906 | 0.274554 |
| 53.1484 | 0.253094 |
| 50.0642 | 0.154433 |
| 38.7066 | 0.182905 |
| 29.4828 | 0.222966 |
| 24.774 | 0.273846 |
| 23.1146 | 0.2205 |
| 42.3447 | 0.240355 |
| 15.6867 | 0.134413 |
| 16.5152 | 0.132108 |
| 15.5908 | 0.121882 |
| 6.53791 | 0.10115 |
| 7.30031 | 0.116975 |
| 5.91994 | 0.135349 |
| 4.37124 | 0.129763 |
| 80.7777 | 0.338074 |
| 55.4441 | 0.218717 |
| 49.8265 | 0.195352 |
| 47.206 | 0.154662 |
| 35.7053 | 0.150655 |
| 26.7663 | 0.2103 |
| 21.3133 | 0.25113 |
| 19.8378 | 0.212126 |
| 45.8313 | 0.201697 |
| 15.0027 | 0.126853 |
| 16.6249 | 0.133424 |
| 16.5325 | 0.117525 |
| 11.4017 | 0.120363 |
| 11.2234 | 0.134374 |
| 8.75982 | 0.140545 |
| 2.38342 | 0.088679 |
| 5.21637 | 0.120194 |
| 93.9132 | 0.355441 |
| 69.9013 | 0.232926 |
| 65.1064 | 0.210328 |
| 61.2801 | 0.167042 |
| 50.4215 | 0.17691 |
| 41.0885 | 0.229569 |
| 37.5744 | 0.271084 |
| 35.7662 | 0.216518 |
| 37.8789 | 0.229325 |
| 25.7368 | 0.147501 |
| 25.1239 | 0.157151 |
| 22.937 | 0.15211 |
| 10.1086 | 0.115408 |
| 12.9394 | 0.138385 |
| 15.2068 | 0.157822 |
| 18.4594 | 0.140055 |
| 14.208 | 0.157282 |
| 19.3459 | 0.133062 |
| 106.94 | 0.335166 |
| 82.2152 | 0.253303 |
| 76.9418 | 0.231848 |
| 73.7269 | 0.178781 |
| 62.4525 | 0.188713 |
| 53.1723 | 0.236054 |
| 48.5555 | 0.257887 |
| 46.9116 | 0.213976 |
| 51.1869 | 0.250425 |
| 38.5113 | 0.173535 |
| 38.476 | 0.184657 |
| 36.6172 | 0.167223 |
| 23.0046 | 0.150796 |
| 26.0946 | 0.148165 |
| 27.4935 | 0.160225 |
| 27.9504 | 0.164662 |
| 23.7987 | 0.156118 |
| 27.6323 | 0.157462 |
| 14.6785 | 0.138762 |
| 178.871 | 0.410655 |
| 155.174 | 0.334307 |
| 150.303 | 0.307172 |
| 146.55 | 0.261909 |
| 135.644 | 0.259165 |
| 126.301 | 0.313621 |
| 122.248 | 0.356002 |
| 120.532 | 0.295593 |
| 106.683 | 0.321306 |
| 111.009 | 0.25336 |

**Figure S15** Mental test plot of genetic and geographic distances for *A. pentaphyllum.*

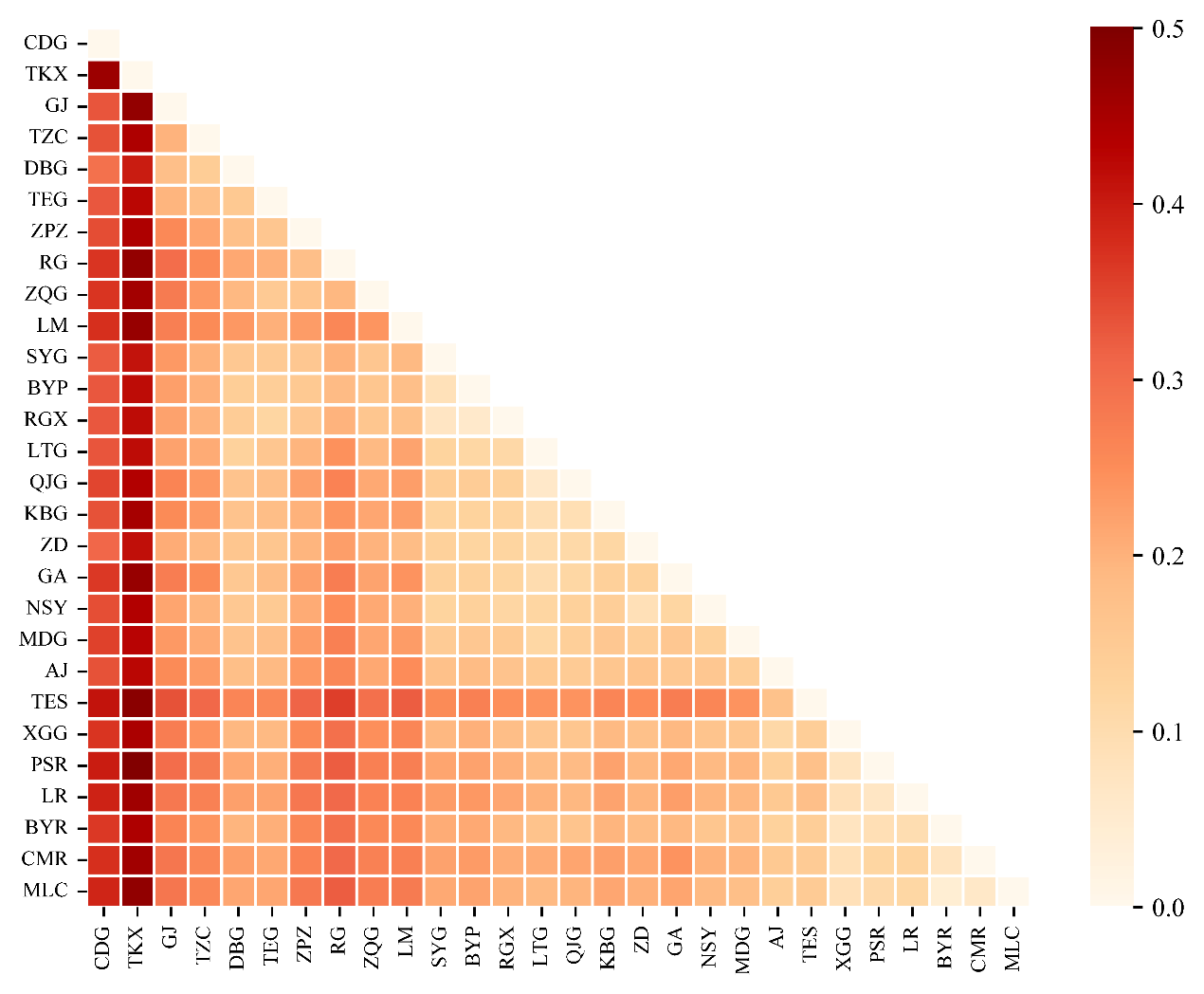

**Figure S16** Genetic differentiation level (*F*_st_) of each population for *A. pentaphyllum*

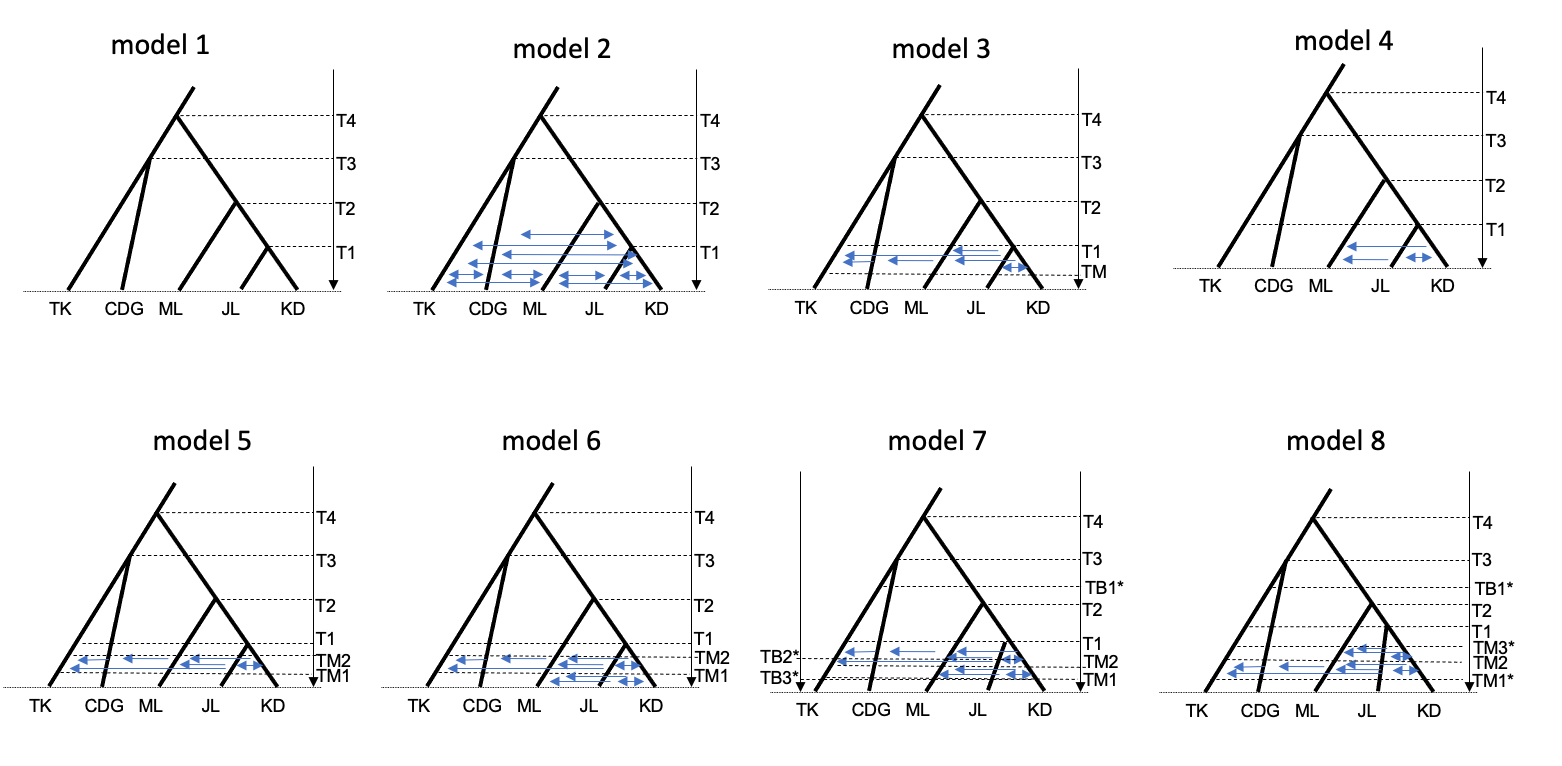

**Figure S17** Representative scenarios tested in this study. Asterisks besides parameters represent changes in population sizes.

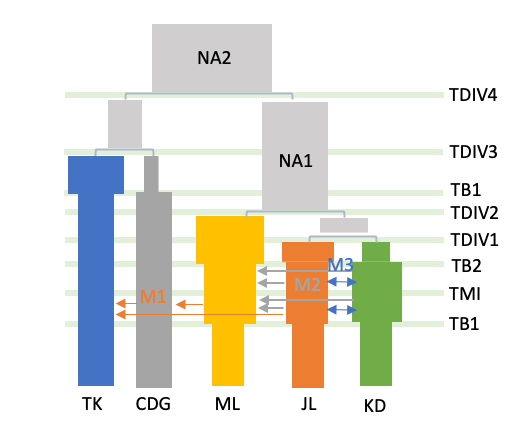

**Figure S18** Illustration of the best-fit model with parameters names.

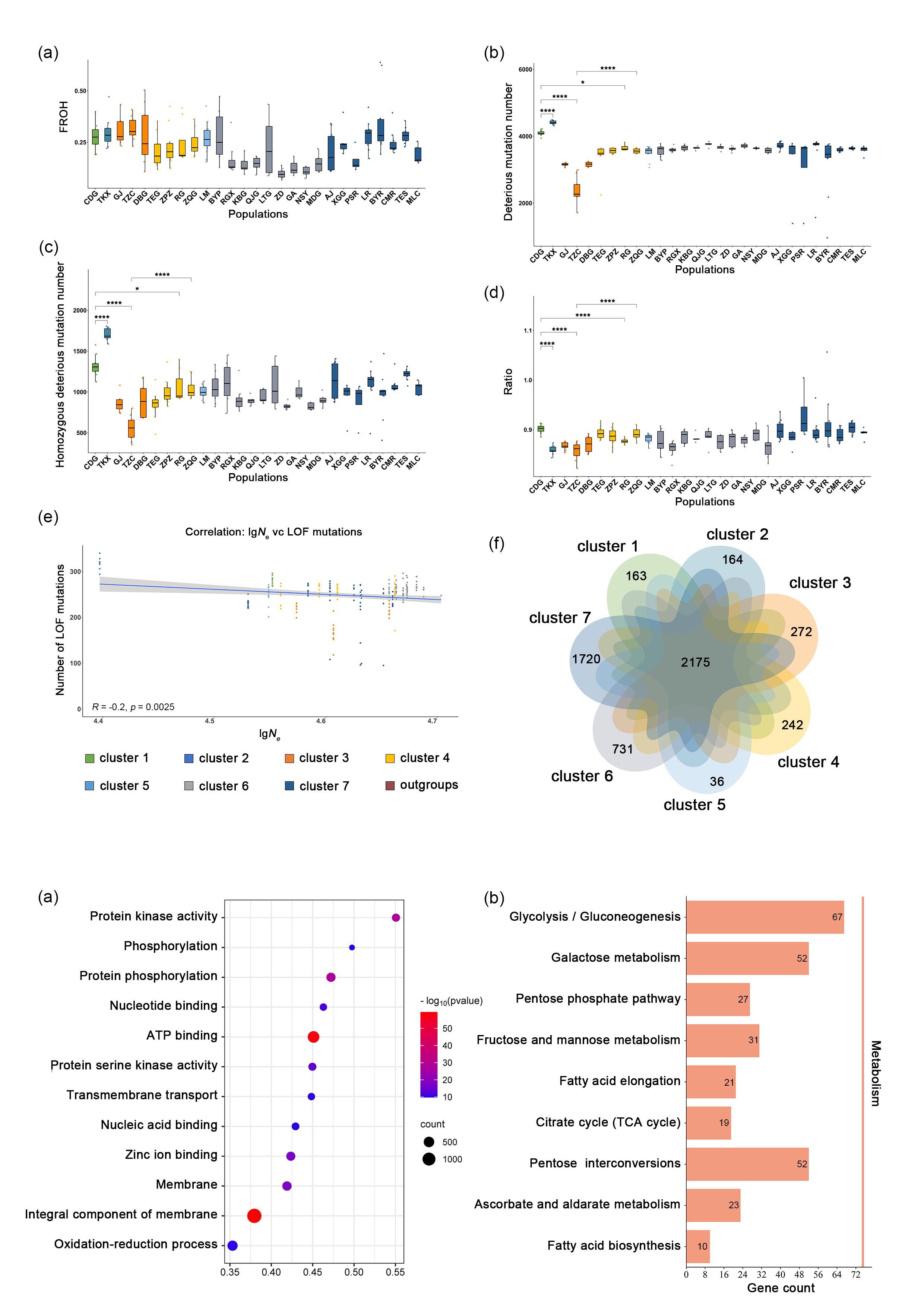

**Figure S19** Correlation analysis between the number of extreme deleterious mutations (LOF) and the effective population size (*N*_e_) across the 28 populations of *A. pentaphyllum.*

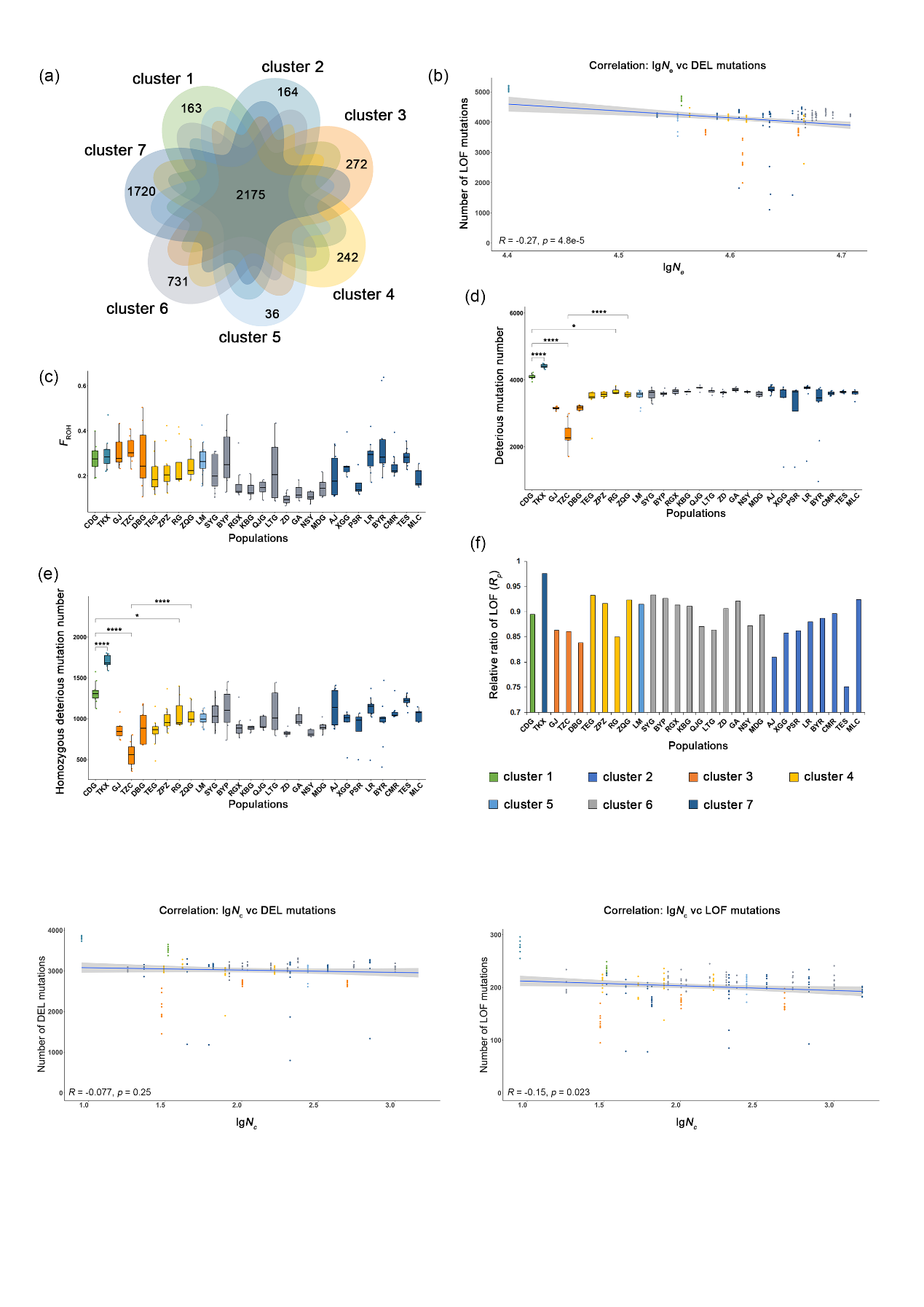

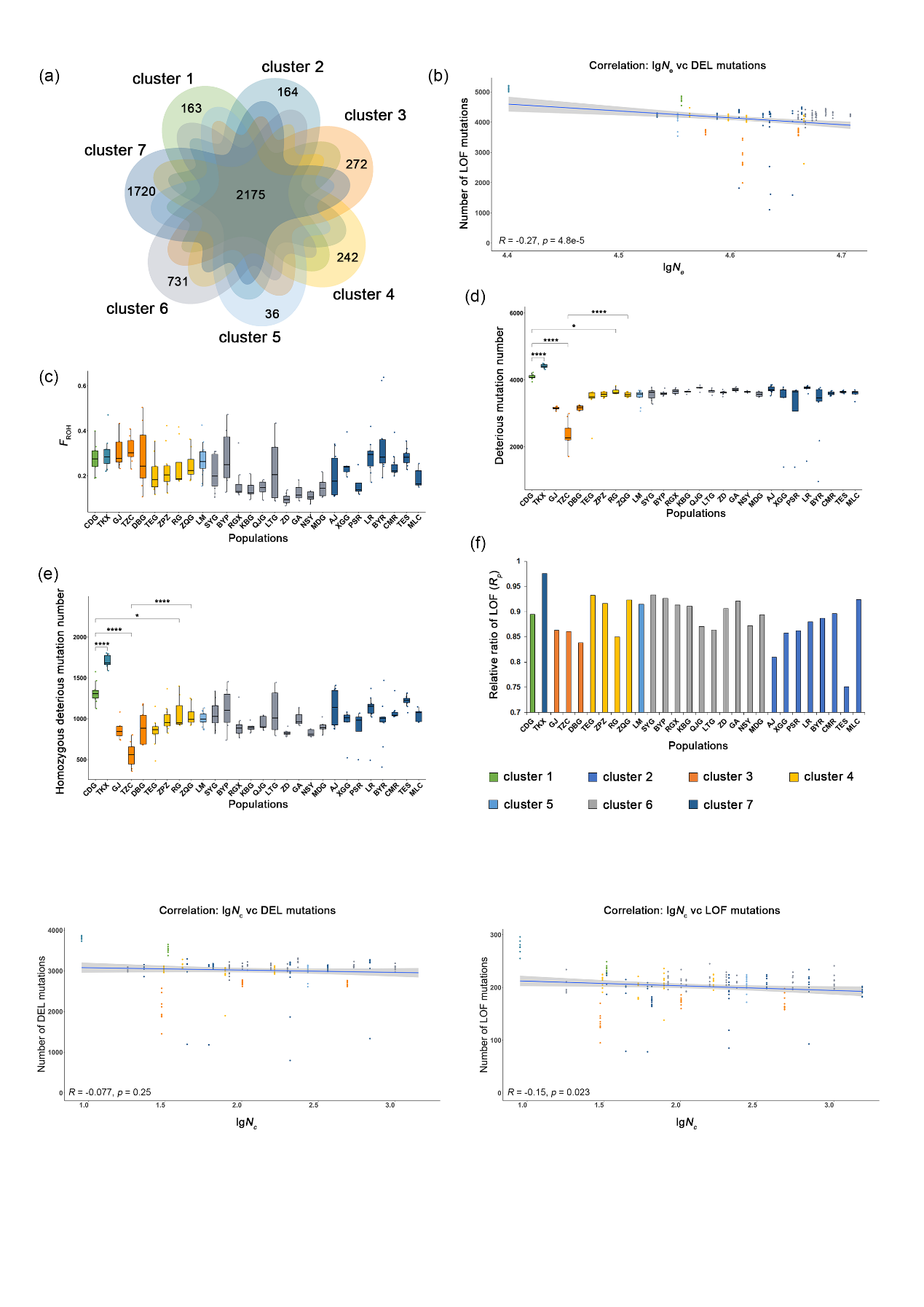

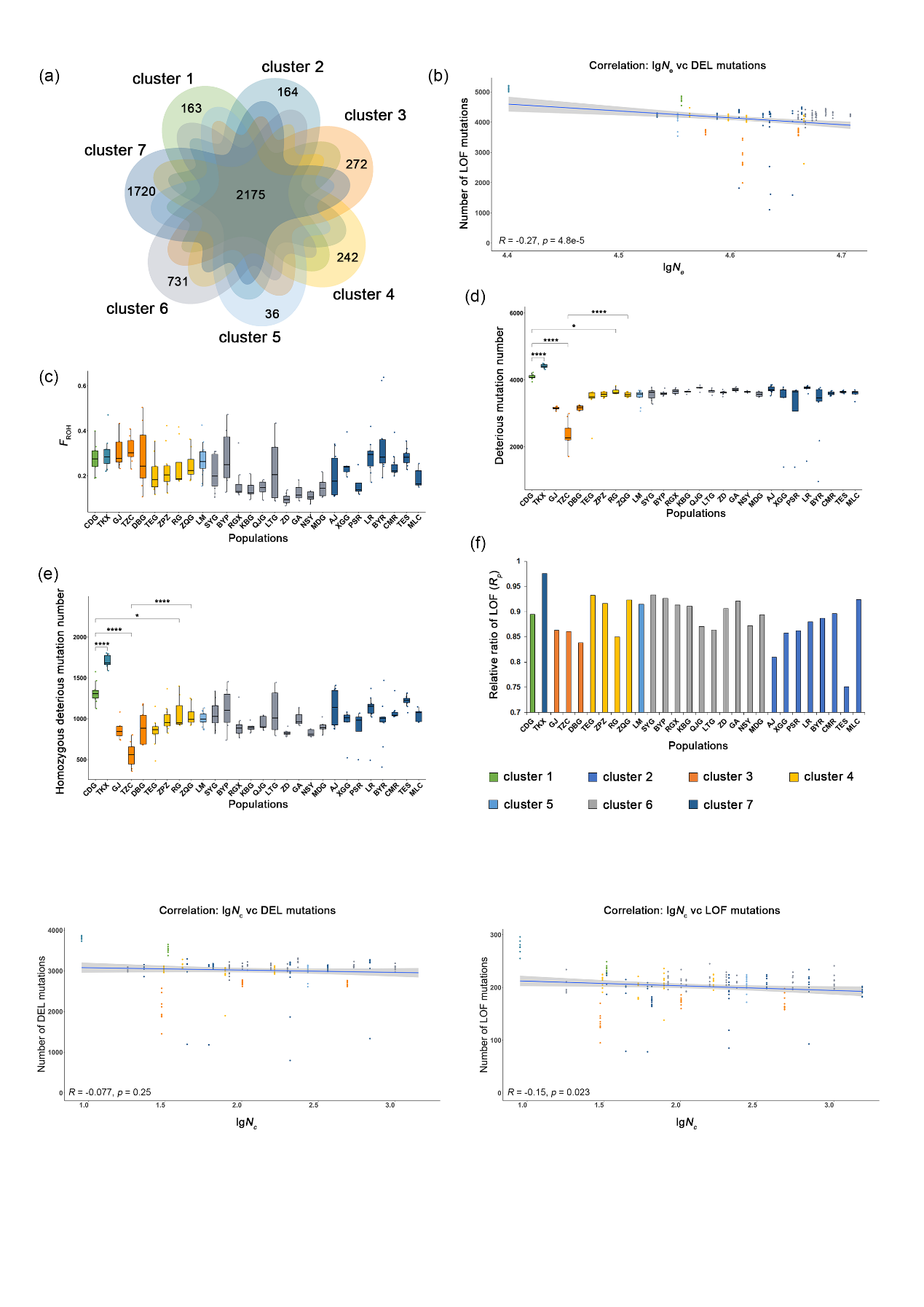

(b)
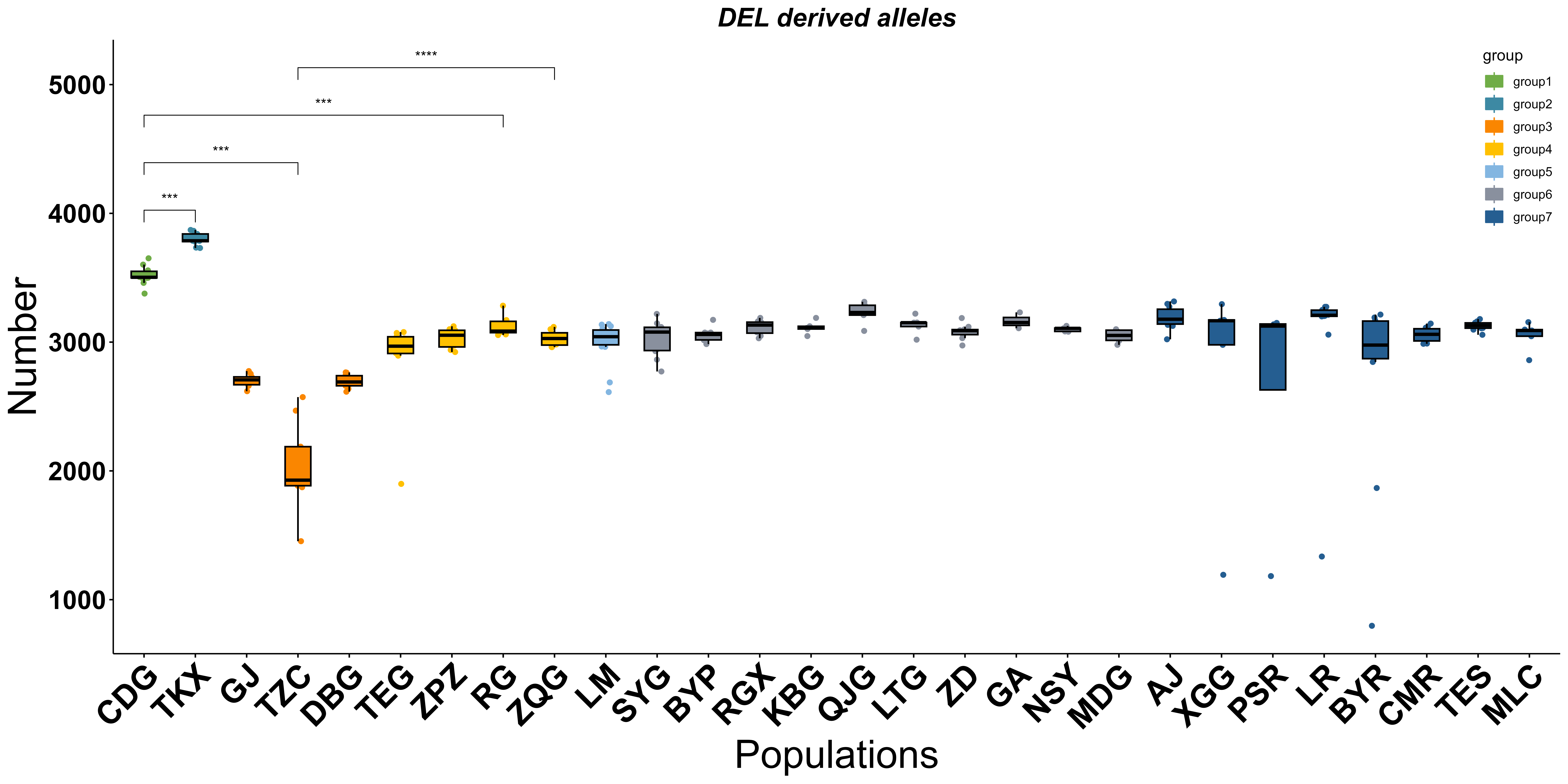

(a)
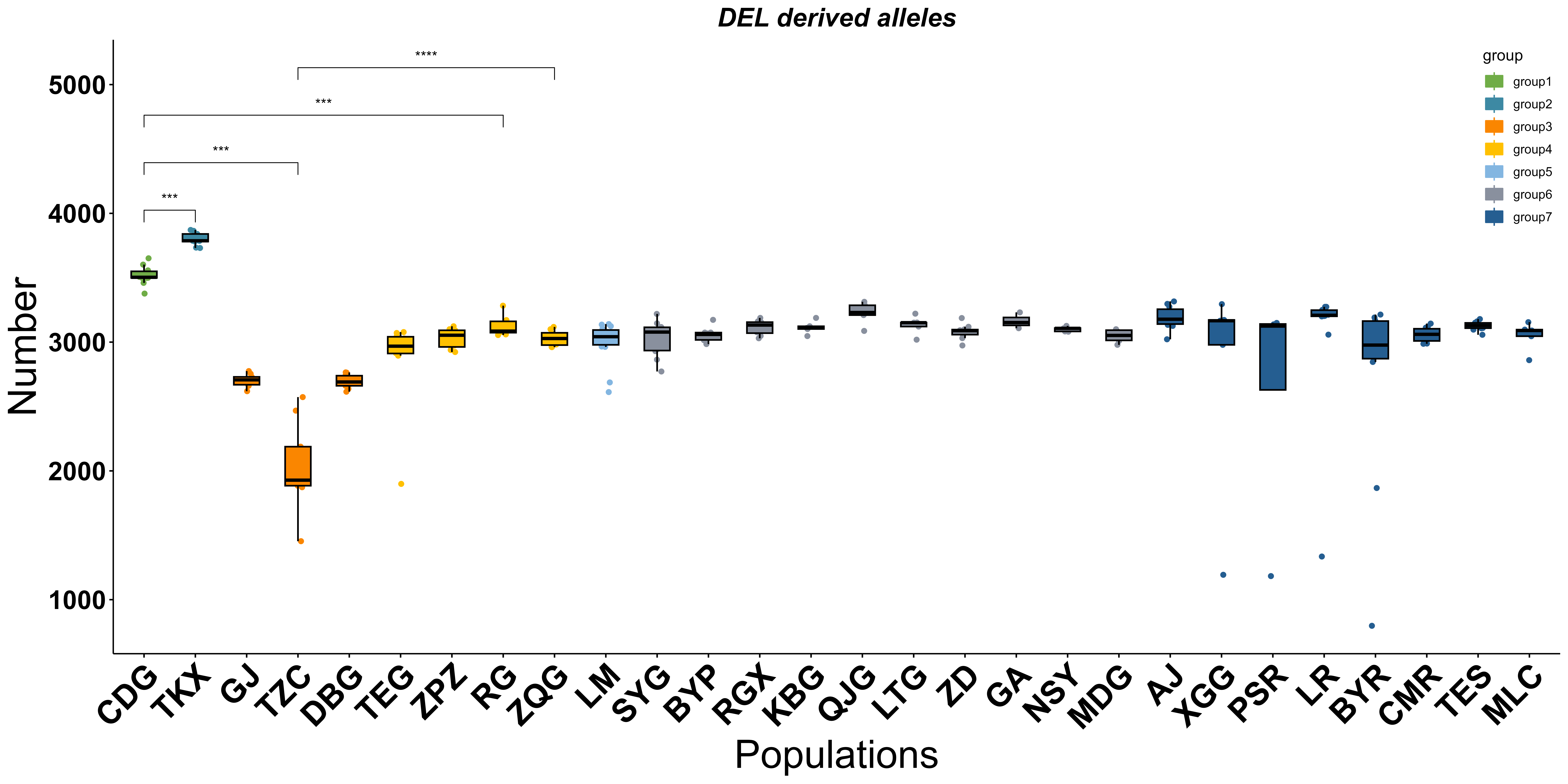

**Figure S20** Correlation analysis between the census population size (*N_c_*) and the number of deleterious mutations of (a) DEL and (b) LOF across the 28 populations of *A. pentaphyllum.*

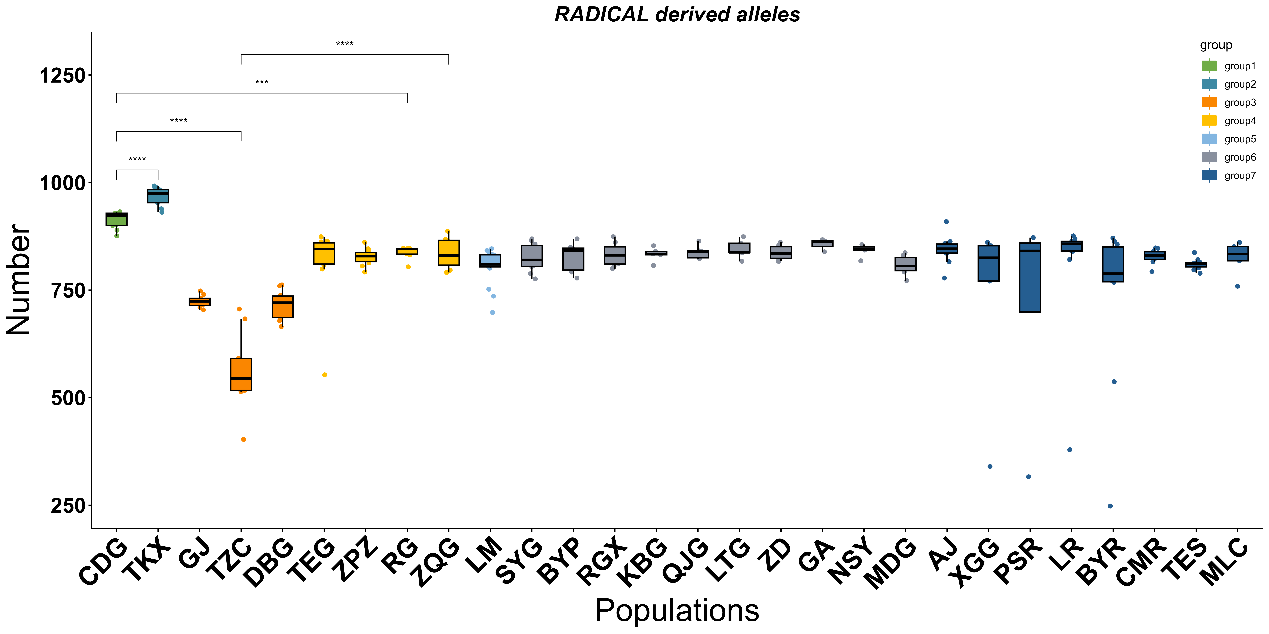

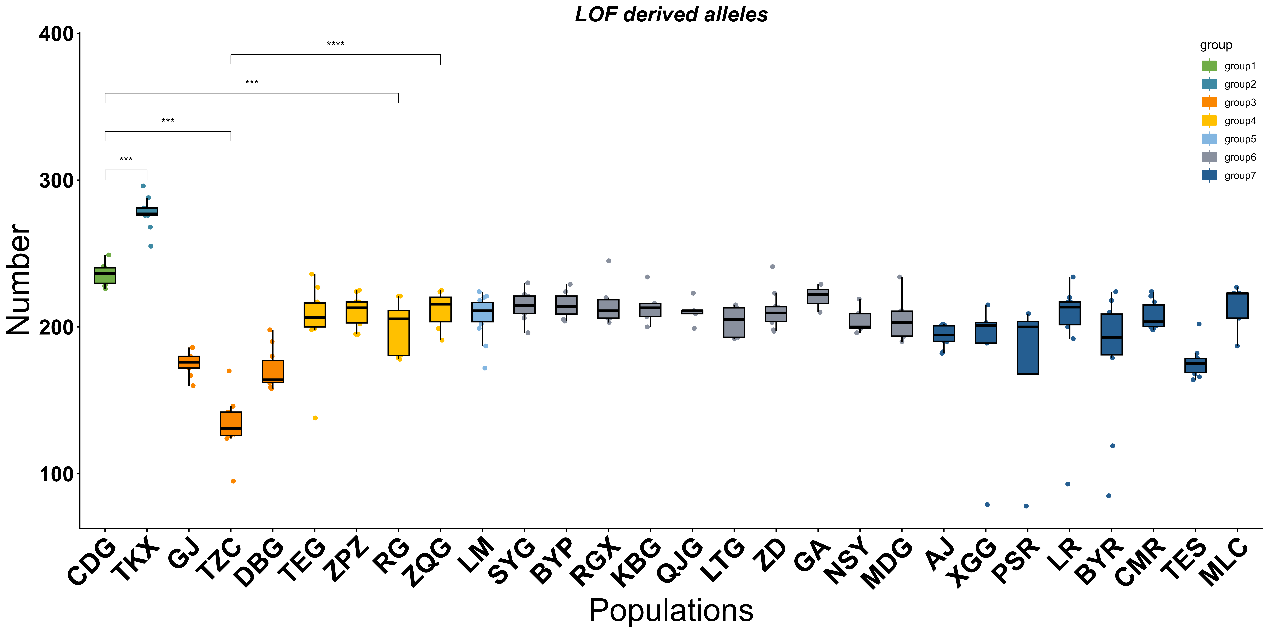

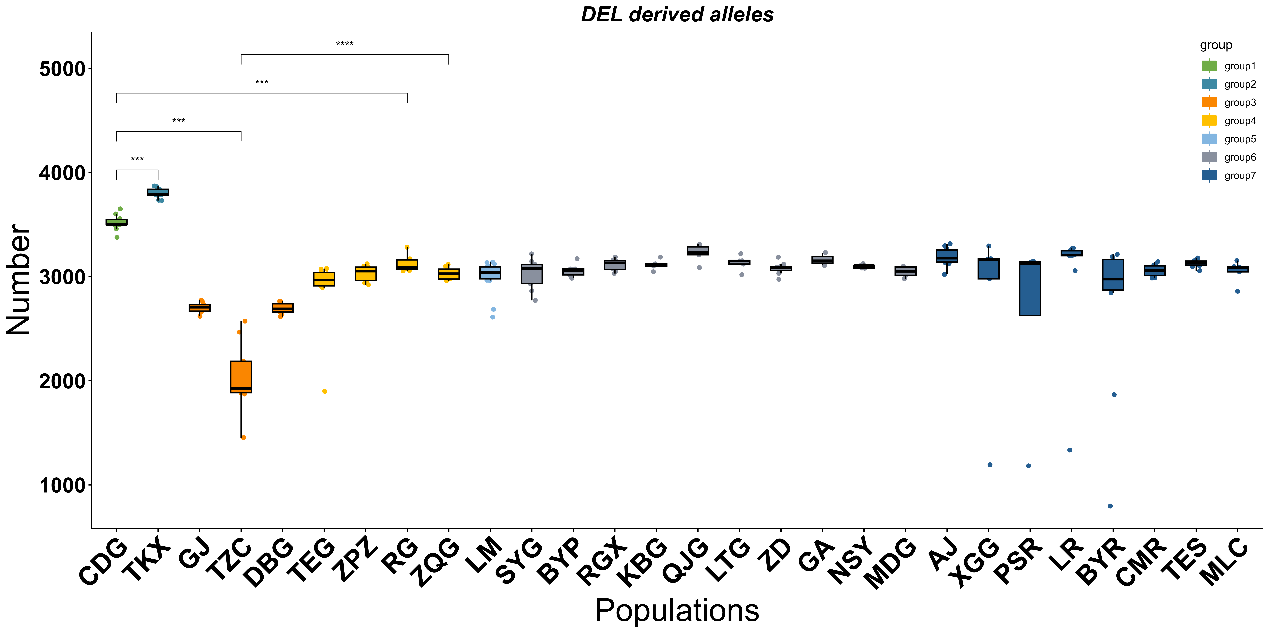
**Figure S21** Number of (a) DEL, (b) LOF, and (c) RADICAL mutations among 28 populations of *A. pentaphyllum.*

(b)

(a)
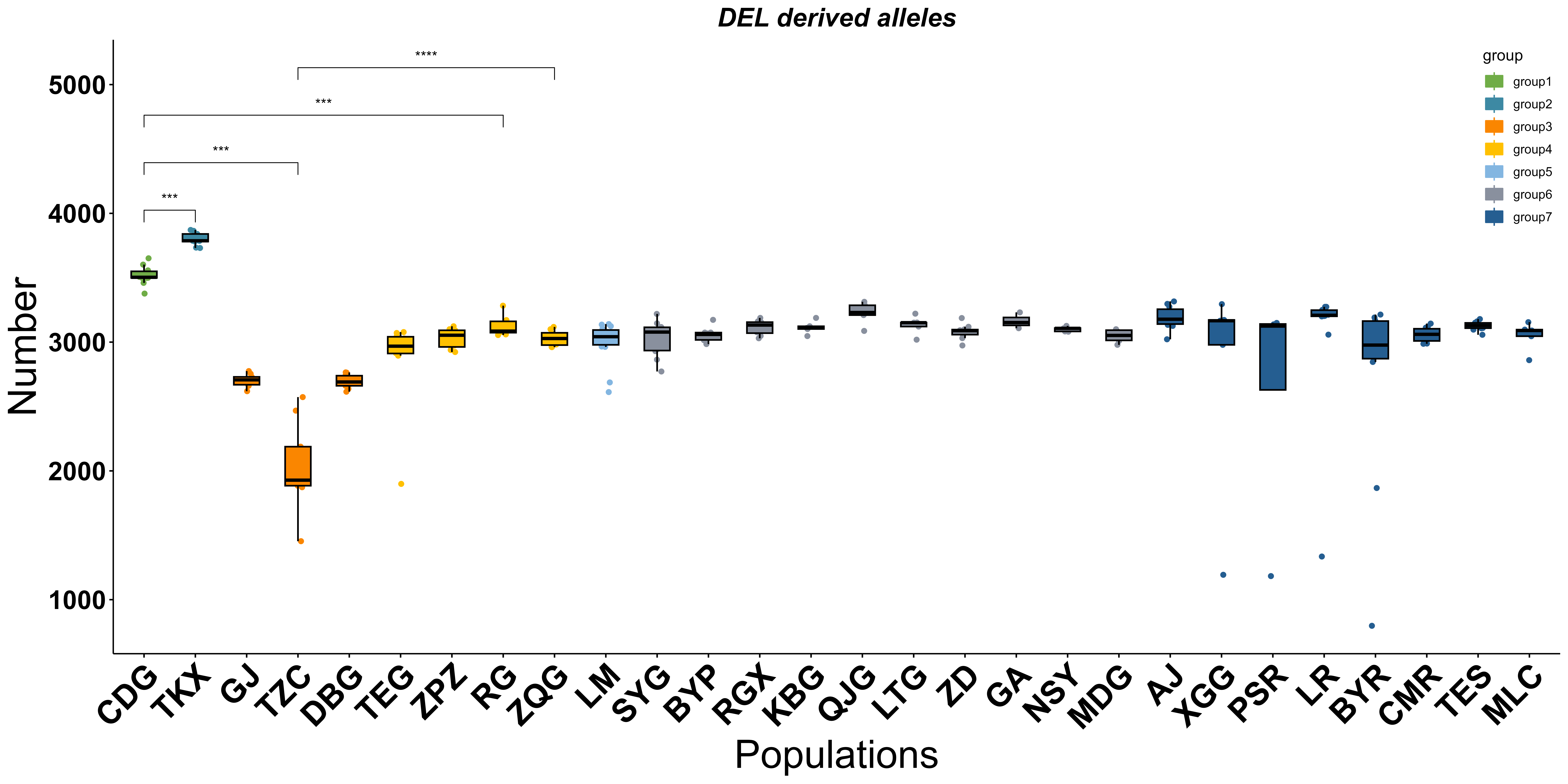

(c)

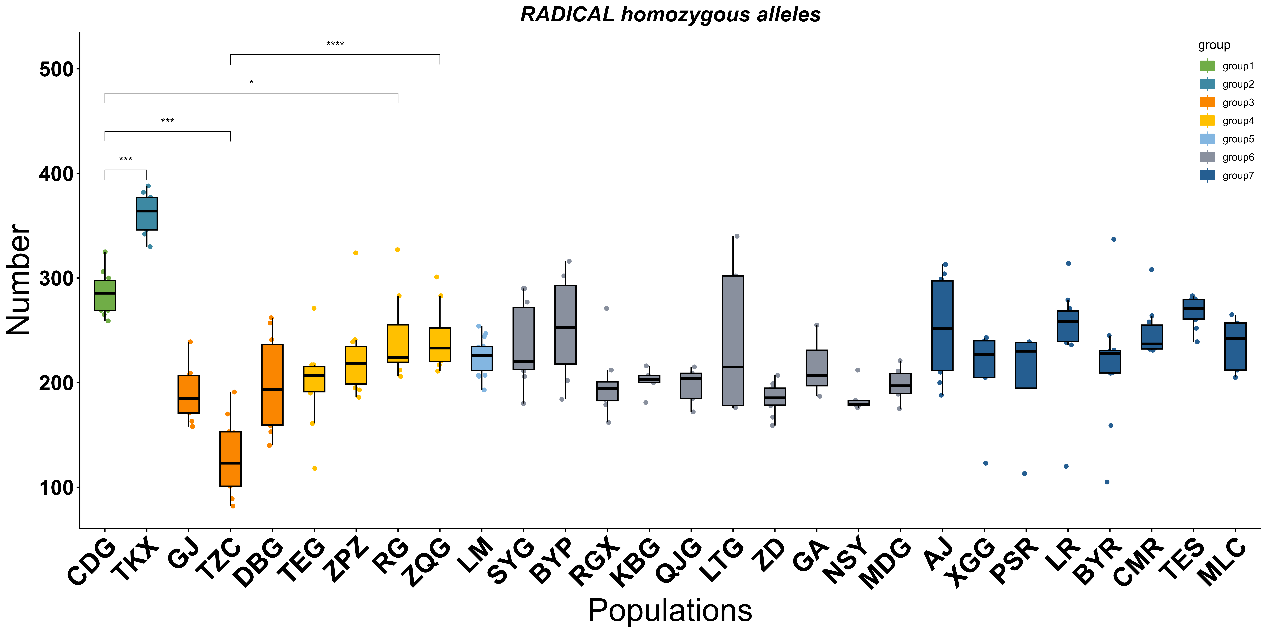

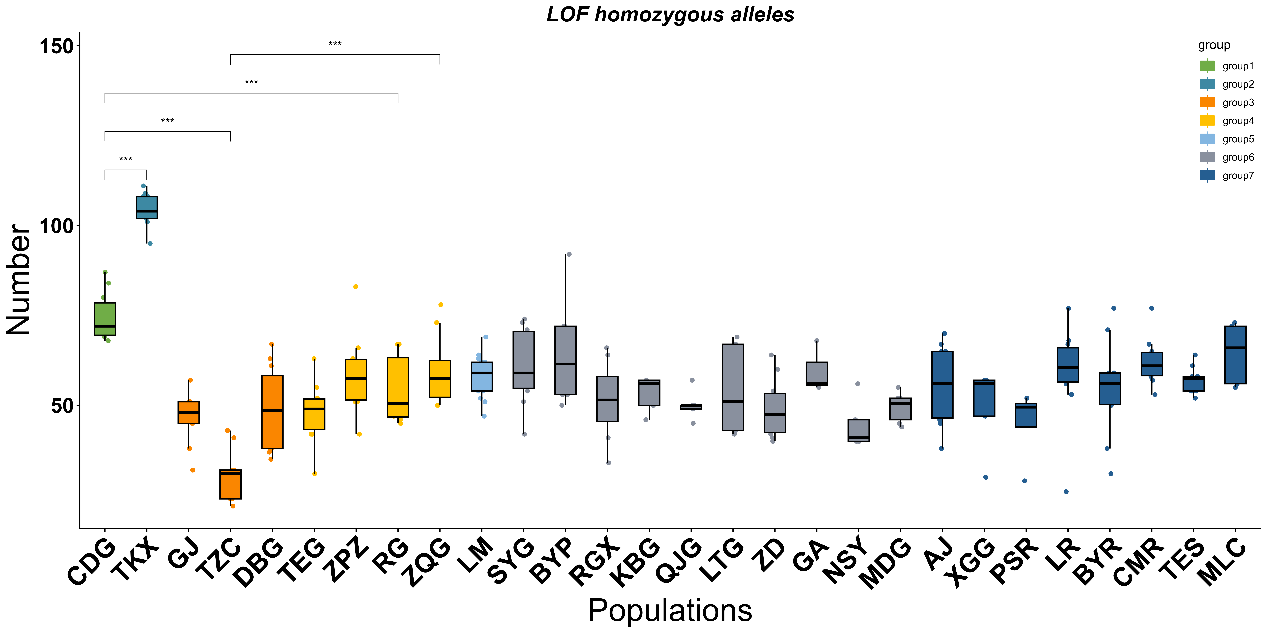

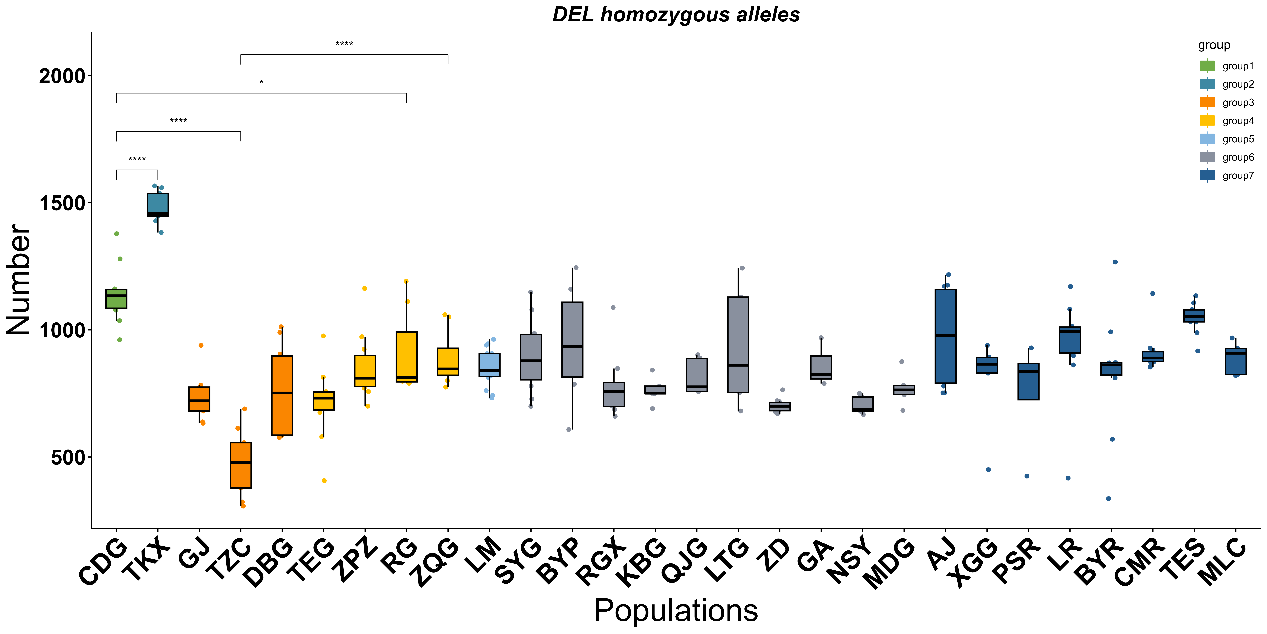
**Figure S22** Number of homozygous (a) DEL, (b) LOF, and (c) RADICAL mutations among 28 populations of *A. pentaphyllum.*

(a)

(b)

(c)

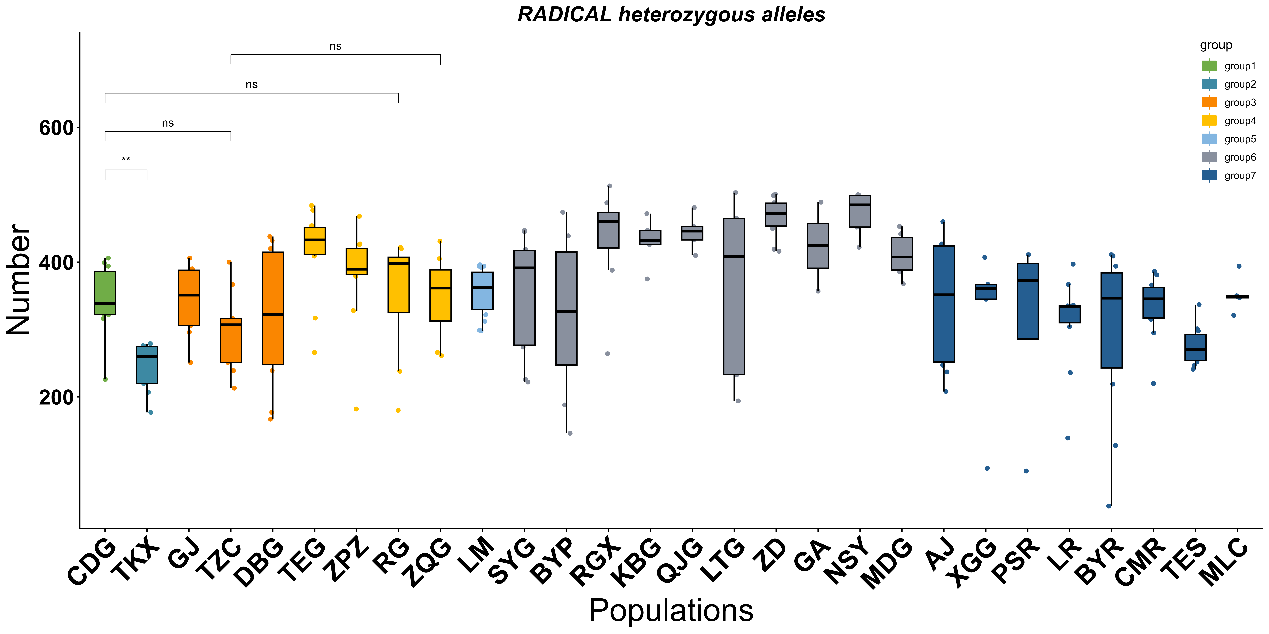

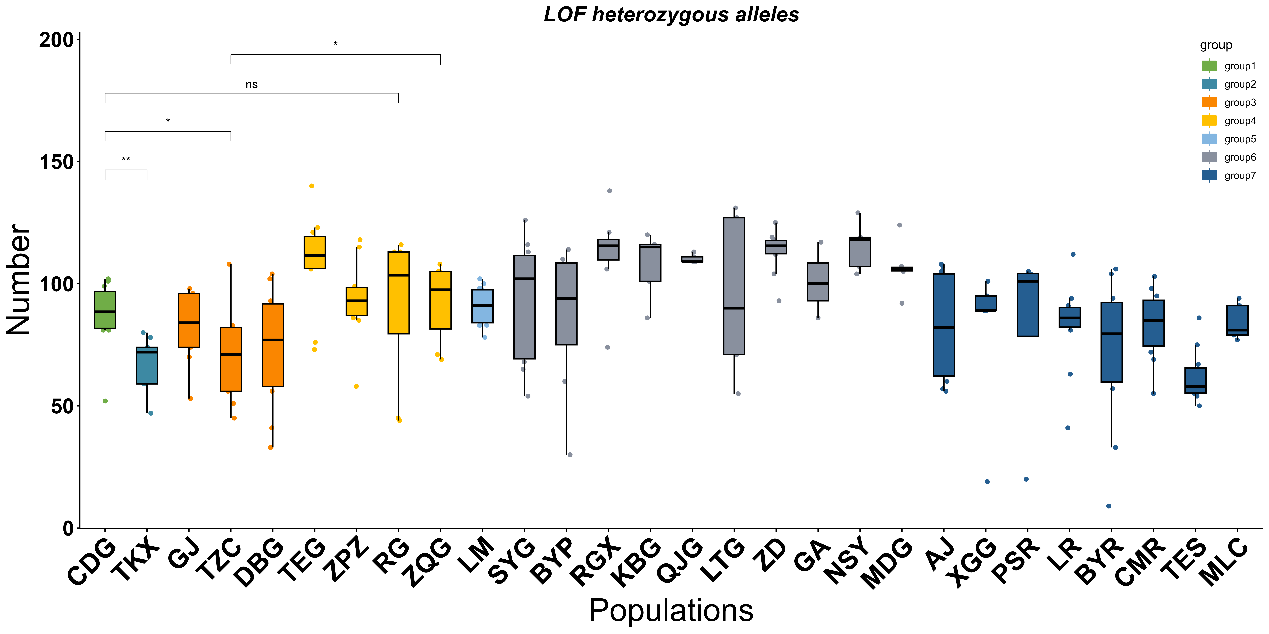

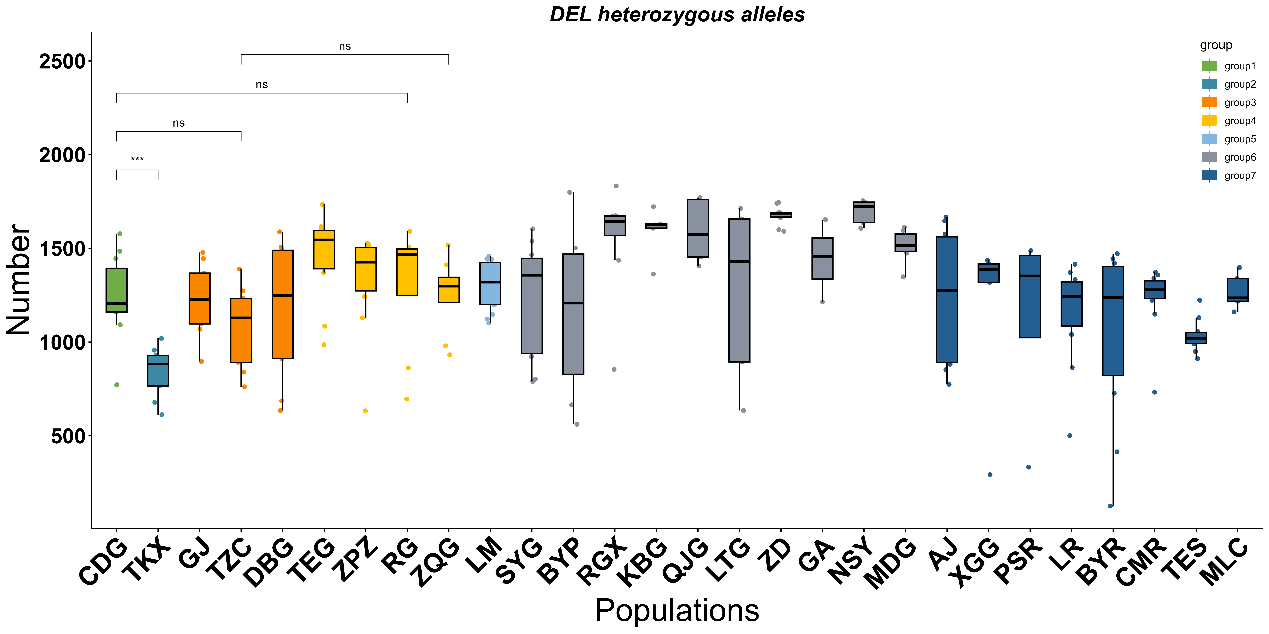
**Figure S23** Number of heterozygous (a) DEL, (b) LOF, and (c) SYN mutations among 28 populations of *A. pentaphyllum.*

(a)

(b)

(c)

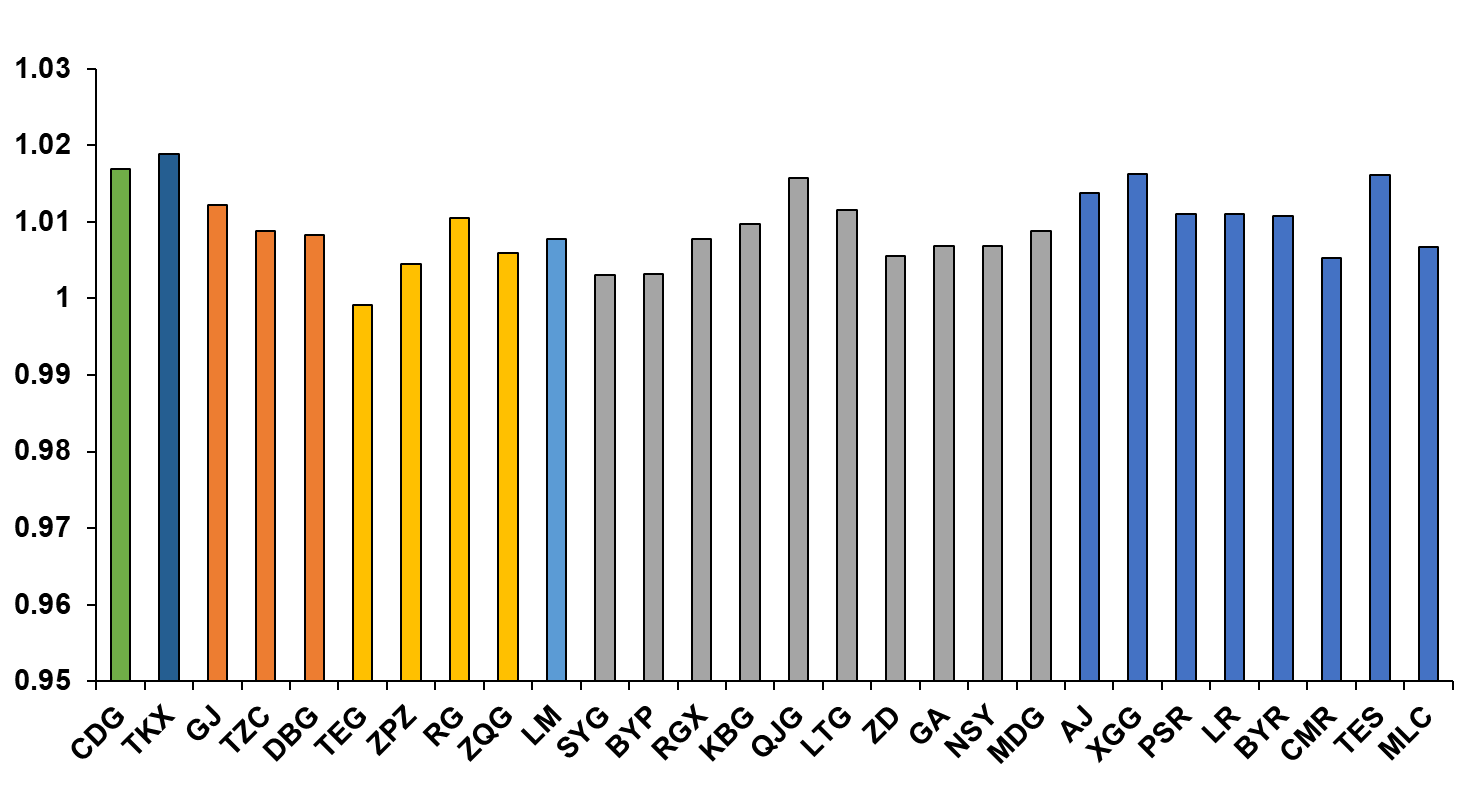

**Relative ratio of RADICAL (*R_p_*)**

**Populations**

(a)

(b)

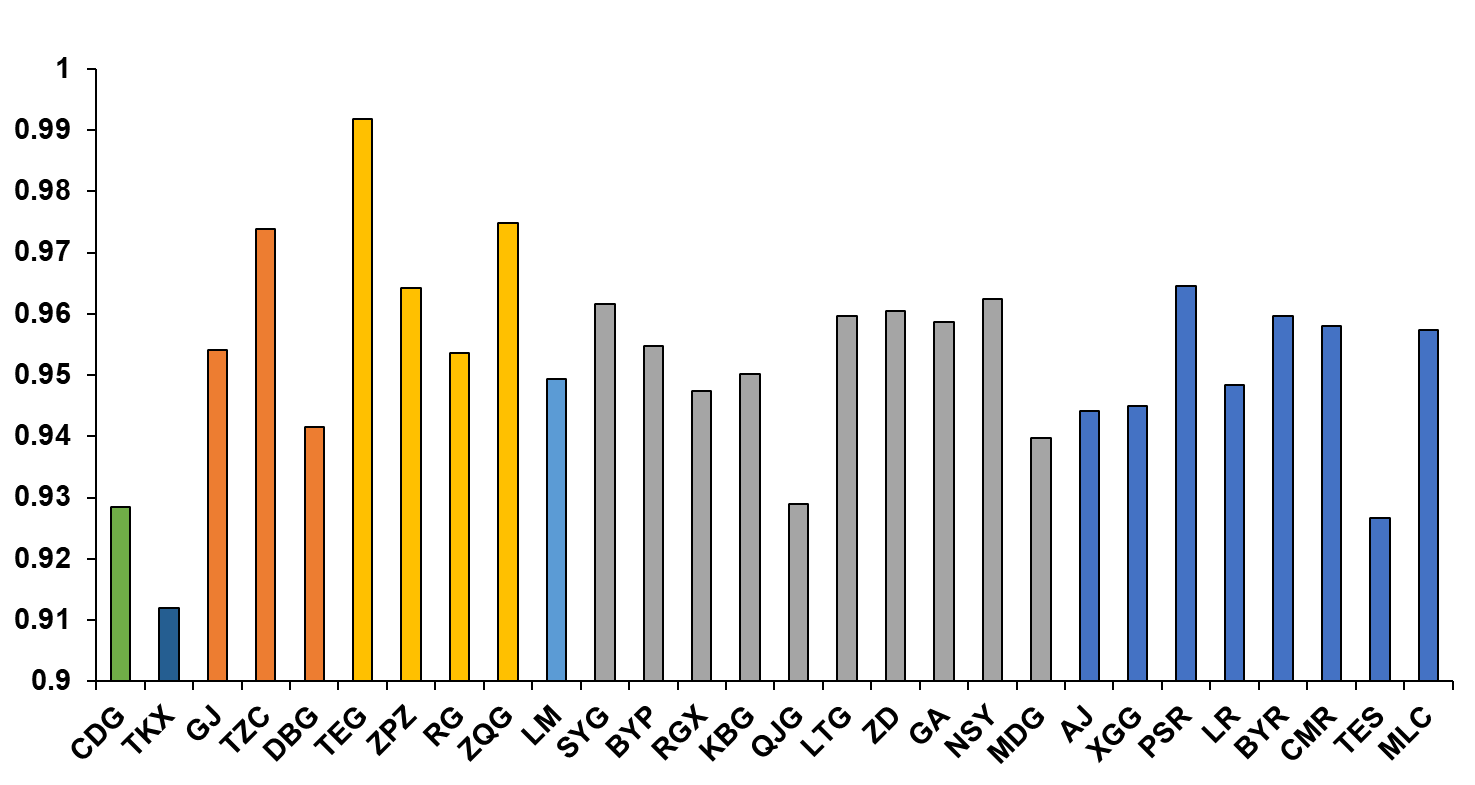

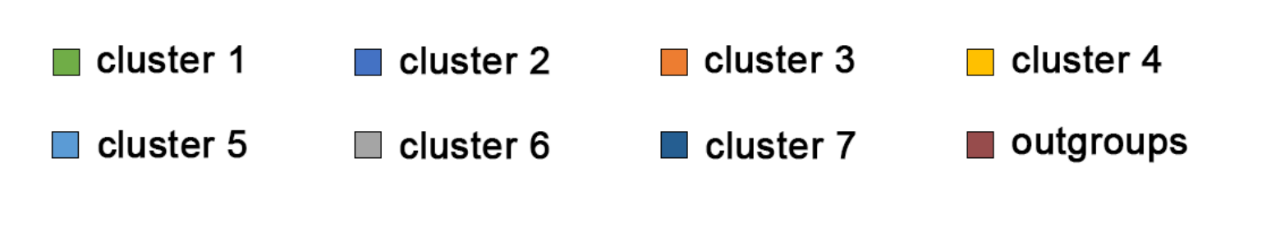

**Populations**

**Relative ratio of DEL (*R_p_*)**

**Figure S24**
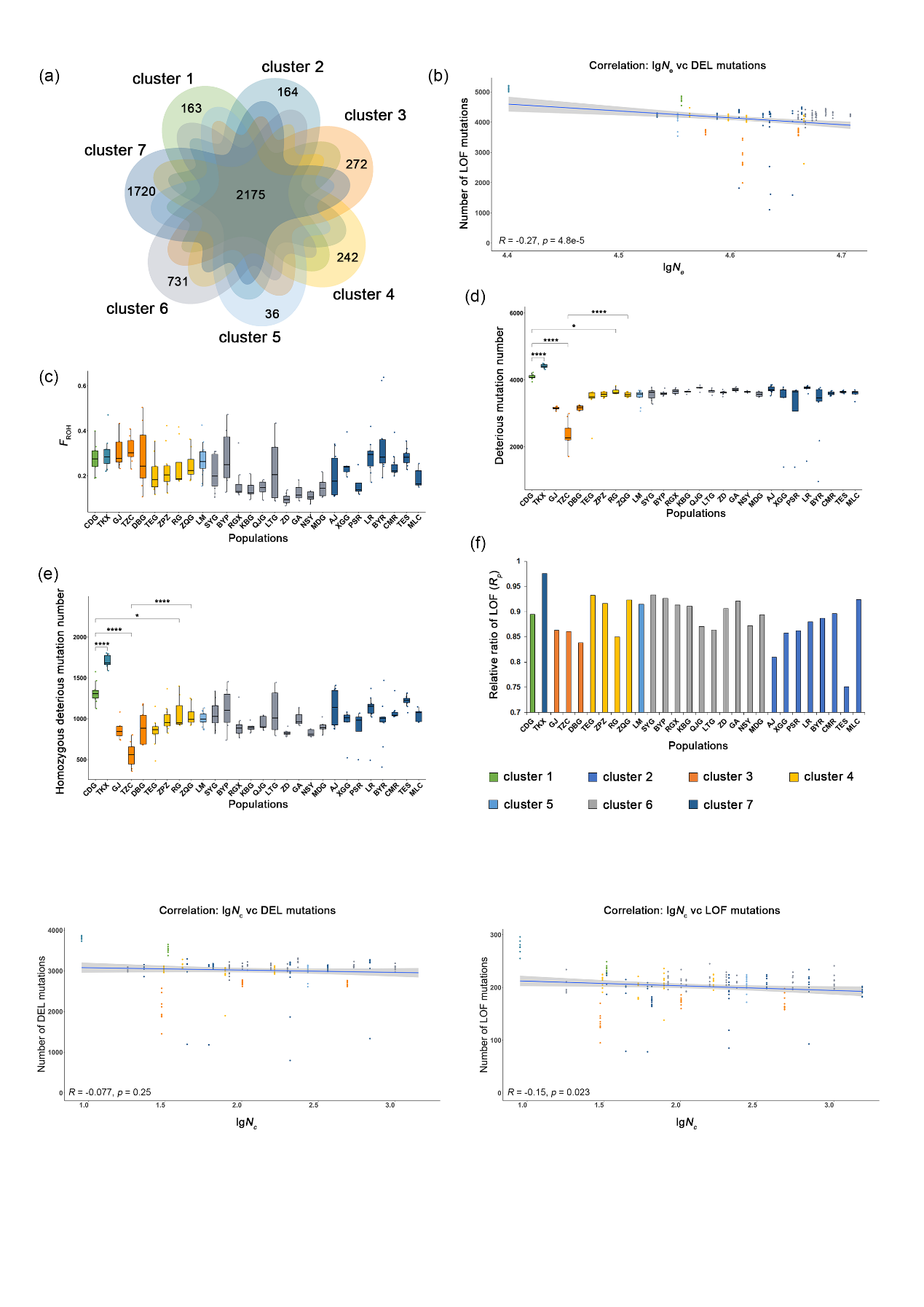
 The relative ratio ($R_{‾p}$) of the mean derived alleles frequency () for (a) DEL and (b) RADICAL mutations.
