## Supplementary material for "High-Resolution Genome Assembly and Population Genetic Study of the Endangered Maple *Acer pentaphyllum* (Sapindaceae): Implications for Conservation Strategies": Supplementary_methods_20240801.docx

### Appendix S1 Estimation of genome features

We determined the genome size and C-value (chromatin-value) using the DNA flow cytometry method (Doležel *et al.*, 2007). By comparing the fluorescence peak value of *A. pentaphyllum* with an internal control species (*Oryza Sativa* ssp. *japonica*, ~430 Mb), the genome size can be calculated using the formula: DNA size of sample = DNA size of internal control species × (fluorescence intensity of sample/fluorescence intensity of internal control species). Also, the k-mer analysis (Marcais and Kingsford, 2011) was used to estimate the genome characteristics of *A. pentaphyllum* using the data from Illumine sequencing. First, to acquire high-quality sequences, we filtered out all low-quality reads with adapters, 20% bases having Phred quality < 5 and > 10% N content using fastp. Then, jellyfish v 2.3.0 (Marcais and Kingsford, 2011) was used to calculate the number and depth of k-mers with the parameter ‘-m 17’. Finally, GenomeScope (Vurture *et al.*, 2017) was used to estimate the genome size and other features, including repeat contents, GC contents, and heterozygosity.

### Appendix S2 Genome annotation

#### Repeat annotation

We employed two strategies to annotate repeat sequences across the *A. pentaphyllum* genome: homology-based and *de novo* prediction. Firstly, MITE-Hunter software (Han and Wessler, 2010) was utilized to identify the Miniature Inverted Repeat Transposable Elements (MITEs), which belong to Class II transposable elements (TEs) and are widely distributed throughout the genome. Secondly, LTRharvest (Ellinghaus *et al.*, 2008) and LTR_FINDER (Xu and Wang, 2007) were employed for identifying the highest proportion of long terminal repeat-retrotransposons (LTR-RTs) in the *A. pentaphyllum* genome, and LTR_retriever (Ou and Jiang, 2018) was then used to integrate the predictions from two software, constructing a high-quality LTR library. In addition, the assembly was compared with the [Repbase](https://mailsucasaccn-my.sharepoint.com/personal/lixiong20_mails_ucas_ac_cn/Documents/五小叶槭基因组和群体分析/文章/Rfam) database (https://www.girinst.org/repbase/) using RepeatMasker (Tarailo-Graovac and Chen, 2009) to search for consensus sequences. Finally, repeat sequences were identified and classified *de novo* using RepeatModeler2 (Flynn *et al.*, 2020). The above results were further integrated to perform repeat prediction for *A. pentaphyllum*.

#### Gene structure prediction

Transcriptome sequences, homologous proteins, and *de novo* predictions were utilized to estimate the gene structure of the genome, involving the following steps: (1) The transcriptome sequence data from different organs of *A. pentaphyllum* were processed using HISAT2 (Kim *et al.*, 2019). The generated full-length transcripts were then aligned to the genome using PASA (Haas *et al.*, 2003), through which corresponding prediction open reading frame (ORF) models were obtained; (2) 140,473 nonredundant protein sequence data from closely related species, including *A. yangbiense*, *A. pseudosieboldianum*, *A. truncatum*, *A. catalpifolium*, and *Sapindus mukorossi* Gaertn, were compared with the reference genome of *A. pentaphyllum* using GeMoMa (Keilwagen *et al.*, 2019). Subsequently, the alignment results from all homologous species were combined to determine the exon and intron boundaries for the predicted genes; (3) For *de novo* estimation based on the genes assembled from the transcriptome data, reliable genes were chosen to train models using AUGUSTUS (Stanke *et al.*, 2004), SNAP (Korf, 2004), GlimmerHMM (Majoros *et al.*, 2004), and GeneMark-ET (Hoff *et al.*, 2016) software to predict gene structure in the genome, obtaining an *A. pentaphyllum* prediction model. Following the integrated gene prediction results obtained by the three methods above using the EvidenceModeler (Haas *et al.*, 2008), the initial prediction genomic gene set of *A. pentaphyllum* was obtained. Finally, UTR regions and alternative splicing on the genome were annotated using PASA based on the comparison of transcripts with the predicted genes, after which the final gene set was obtained for subsequent analysis.

#### Non-coding RNA annotation

Non-coding RNAs (ncRNAs) are functional RNA molecules that cannot be translated into proteins, including tRNAs, rRNAs, and small RNAs such as microRNAs, siRNAs, and so on. Compared to traditional translatable mRNA, ncRNAs are shorter and less abundance within cells but play crucial roles in t gene expression regulation. tRNAscan-SE v2.0 (Lowe and Eddy, 1997) was employed to *de novo* predict tRNAs in the genome of *A. pentaphyllum*, while RNAmmer (Lagesen *et al.*, 2007) was used to predict the rRNA and subunits. Additionally, the reference genome was compared with the [Rfam](https://mailsucasaccn-my.sharepoint.com/personal/lixiong20_mails_ucas_ac_cn/Documents/五小叶槭基因组和群体分析/文章/Rfam) database (<http://rfam.xfam.org/>) using Infernal software (Nawrocki and Eddy, 2013) to annotate other non-coding RNA. The above results were further integrated to perform ncRNA prediction for *A. pentaphyllum*.

#### Gene function annotation

Gene function annotation was performed for five public protein databases via BLASTP (https://blast.ncbi.nlm.nih.gov/Blast.cgi?PROGRAM=blastp), including Non-Reduntant Protein Database (NR, https://ftp.ncbi.nlm.nih.gov/blast/db/FASTA/), Gene Ontology (GO, http://geneontology.org/), Kyoto Encyclopedia of Gene and Genomes (KEGG, https://www.kegg.jp/), evolutionary genealogy of genes: Non-supervised Orthologous Groups 5.0 (eggNOG, <http://eggnog5.embl.de/download/eggn> og5.0/), and Swissport (<https://www.uniprot.org/>).

### Appendix S3 Positive selection analysis

PAL2NAL (Suyama *et al.*, 2006) was employed to convert single-copy orthologous protein sequences of 15 species into codon-based alignment CDS sequences, serving as input files for the CODEML program in PAML (Yang, 2007). The specific steps of positive selection analysis are as follows: the null hypothesis analysis was initiated by setting fix_omega = 1 and omega = 1, yielding likelihood value l0, then the alte hypothesis analysis was repeated by setting fix_omega = 0 and omega = 1.5, yielding l1. After conducting a chi-square test using the command "chi2 1 2Δ(l1-l0)", genes with p-value <0.05 were considered under positive selection. GO functional annotations were performed on these genes to infer their potential functions.

### Appendix S4 Lineage divergence and inference of demographic history

To infer lineage divergence times within *A. pentaphyllum,* we extracted 225 single-copy orthologous genes from five representative individuals and dated the phylogeny including six *Acer* species and two *Dipteornia* species using MCMCTREE (Yang, 2007) with the same calibrations used in the comparative genome analysis. These inferred divergence times were used to set the prior distribution of divergence in the FASTSIMCOAL2 (Excoffier *et al*., 2013) simulations.

In addition, to recover the demographic history and explore the role of gene flow in the process of speciation and lineage divergence of *A. pentaphyllum*, we conducted composite maximum likelihood (ML) inference as complementary based on site frequency spectrum (SFS). To generate unfolded site frequency spectrum (u-SFS), we inferred the probability of the derived versus ancestral allelic state using est-sfs v2.03 (Pickrell and Pritchard, 2012). One individual of *D. sinensis* and *A. yangbiense* was used as the outgroups in the est-sfs analysis, leaving 679,558 ancestral state sites (Polarized SNPs). The joint unfolded 2D-SFS were built using neutral SNPs with using easySFS.py (https:// github.com/isaacovercast/easySFS). The SNP data of each cluster were down-projected to an SFS with relatively similar sampling sizes (18 and 20) across groups to decrease the effect of different levels of missing data between clusters.

Eight representative scenarios were set focusing on changes in population sizes and occurrence and magnitude of gene flow among populations grouped based on genetic structure and geographical distributions (i.e., TK, CDG, ML, JL, KD) of *A. pentaphyllum* (Figure S17). We calculated the likelihood function for different demographic scenarios using the software FASTSIMCOAL2 v2.6 (Excoffier et al. 2013). For each scenario, 100,000 coalescent simulations per likelihood estimation (-n 100,000) and 40 expectation-conditional maximization (ECM) cycles (-L40) were used as the command line parameters for each run. The Akaike information criterion (AIC) was used to compare different models. In this case, AIC = 2k–2ln(MaxEstLhood), where k is the number of parameters estimated by each model, and MaxEstLhood is the ML function value for each model. Moreover, when searching for a maximum likelihood value, FASTSIMCOAL2 may reach a local optimum instead of a global optimum. Thus, we repeated each step at least twice (results not shown), to ensure we were not ending in a local optimum, thereby getting better estimates of the global optimum.

### References

**Doležel, J., Greilhuber, J. and Suda, J.** (2007) Flow cytometry with plants: an overview. *Flow cytometry with plant cells: analysis of genes, chromosomes and genomes*, 41-65.

**Ellinghaus, D., Kurtz, S. and Willhoeft, U.** (2008) LTRharvest, an efficient and flexible software for de novo detection of LTR retrotransposons. *BMC Bioinformatics*, **9**, 18.

**Excoffier, L., Dupanloup, I., Huerta-Sánchez, E., Sousa, V.C. and Foll, M.** (2013) Robust Demographic Inference from Genomic and SNP Data. *PLOS Genetics*, **9**.

**Flynn, J.M., Hubley, R., Goubert, C., Rosen, J., Clark, A.G., Feschotte, C. and Smit, A.F.** (2020) RepeatModeler2 for automated genomic discovery of transposable element families. *Proceedings of the National Academy of Sciences of the United States of America*, **117**, 9451-9457.

**Han, Y. and Wessler, S.R.** (2010) MITE-Hunter: a program for discovering miniature inverted-repeat transposable elements from genomic sequences. *Nucleic Acids Research*, **38**, e199.

**Haas, B.J., Delcher, A.L., Mount, S.M., Wortman, J.R., Smith, R.K., Hannick, L.I., Maiti, R., Ronning, C.M., Rusch, D.B., Town, C.D., Salzberg, S.L. and White, O.** (2003) Improving the genome annotation using maximal transcript alignment assemblies. *Nucleic Acids Research*, **31**, 5654-5666.

**Haas, B.J., Salzberg, S.L., Zhu, W., Pertea, M., Allen, J.E., Orvis, J., White, O., Buell, C.R. and Wortman, J.R.** (2008) Automated eukaryotic gene structure annotation using EVidenceModeler and the program to assemble spliced alignments. *Genome Biology*, **9**.

**Hoff, K.J., Lange, S., Lomsadze, A., Borodovsky, M. and Stanke, M.** (2016) BRAKER1: Unsupervised RNA-Seq-Based Genome Annotation with GeneMark-ET and AUGUSTUS. *Bioinformatics*, **32**, 767-769.

**Keilwagen, J., Hartung, F. and Grau, J.** (2019) GeMoMa: Homology-Based Gene Prediction Utilizing Intron Position Conservation and RNA-seq Data. *Methods in Molecular Biology*. 1962, 161-177.

**Kim, D., Paggi, J.M., Park, C., Bennett, C. and Salzberg, S.L.** (2019) Graph-based genome alignment and genotyping with HISAT2 and HISAT-genotype. *Nature Biotechnology*, **37**, 907-+.

**Korf, I.** (2004) Gene finding in novel genomes. *Bmc Bioinformatics*, **5**.

**Lagesen, K., Hallin, P., Rodland, E.A., Stærfeldt, H.H., Rognes, T. and Ussery, D.W.** (2007) RNAmmer: consistent and rapid annotation of ribosomal RNA genes. *Nucleic Acids Research*, **35**, 3100-3108.

**Lowe, T.M. and Eddy, S.R.** (1997) tRNAscan-SE: A program for improved detection of transfer RNA genes in genomic sequence. *Nucleic Acids Research*, **25**, 955-964.

**Marcais, G. and Kingsford, C.** (2011) A fast, lock-free approach for efficient parallel counting of occurrences of k-mers. *Bioinformatics*, **27**, 764-770.

**Majoros, W.H., Pertea, M. and Salzberg, S.L.** (2004) TigrScan and GlimmerHMM: two open source eukaryotic gene-finders. *Bioinformatics*, **20**, 2878-2879.

**Nawrocki, E.P. and Eddy, S.R.** (2013) Infernal 1.1: 100-fold faster RNA homology searches. *Bioinformatics*, **29**, 2933-2935.

**Ou, S. and Jiang, N.** (2018) LTR_retriever: A Highly Accurate and Sensitive Program for Identification of Long Terminal Repeat Retrotransposons. *Plant Physiology*, **176**, 1410-1422.

**Pickrell, J.K. and Pritchard, J.K.** (2012) Inference of population splits and mixtures from genome-wide allele frequency data. *PLOS Genetics*, **8**, e1002967.

**Stanke, M., Steinkamp, R., Waack, S. and Morgenstern, B.** (2004) AUGUSTUS: a web server for gene finding in eukaryotes. *Nucleic Acids Research*, **32**, W309-W312.

**Suyama, M., Torrents, D. and Bork, P.** (2006) PAL2NAL: robust conversion of protein sequence alignments into the corresponding codon alignments. *Nucleic Acids Research*, **34**, W609-W612.

**Tarailo-Graovac, M. and Chen, N.** (2009) Using RepeatMasker to Identify Repetitive Elements in Genomic Sequences. *Curr Protoc Bioinformatics*, **25**, Chapter 4:Unit 4.10.

**Vurture, G.W., Sedlazeck, F.J., Nattestad, M., Underwood, C.J., Fang, H., Gurtowski, J. and Schatz, M.C.** (2017) GenomeScope: fast reference-free genome profiling from short reads. *Bioinformatics*, **33**, 2202-2204.

**Xu, Z. and Wang, H.** (2007) LTR_FINDER: an efficient tool for the prediction of full-length LTR retrotransposons. *Nucleic Acids Research*, **35**, W265-268.

**Yang, Z.** (2007) PAML 4: phylogenetic analysis by maximum likelihood. *Molecular Biology and Evolution*, **24**, 1586-1591.
